# Foliar application of nano urea results in higher biomass, chlorophyll, and nitrogen content than equimolar bulk urea through differential gene regulation in *Arabidopsis thaliana*

**DOI:** 10.1101/2024.09.03.611005

**Authors:** Arpan Dey, Neelam Jangir, Devanshu Verma, Rajveer Singh Shekhawat, Pankaj Yadav, Ayan Sadhukhan

## Abstract

Indian Farmers Fertilizer Cooperative (IFFCO)’s liquid nano urea formulation (NUF) was applied to one-month-old *Arabidopsis thaliana* plants grown in vermiculite as a 0.4% foliar spray twice at an interval of 10 days and compared with sprays of equimolar bulk urea. NUF resulted in a 51 ± 14.9% increase in biomass, 29.5 ± 9.1% in chlorophyll, 8.4 ± 3.1% in nitrogen, and 4.5 ± 0.3% in amino acid content of the leaves, compared to bulk urea. NUF’s zeta potential of -54.7 mV and particle size of ≃27.7 nm, measured by dynamic light scattering and transmission electron microscopy, make it suitable for stomatal uptake. We conducted a differential gene expression analysis by mRNA sequencing to understand the molecular basis of the phenotypic gains under NUF rather than urea. NUF resulted in significantly higher expression levels of 211 genes (log_2_fold-change > 0.5, *FDR* < 0.05) involved in the biosynthesis of carbohydrates, amino acids, nucleotides, lipids, phytohormones, and secondary metabolites, cell wall biosynthesis and modification, growth and developmental processes, cell cycle, and stress response than bulk urea. On the other hand, 1,286 genes (log_2_fold-change < -0.5) involved in cell death, abscission, senescence, nitrogen transport and metabolism, and biotic stress response showed lower expression levels upon NUF application than bulk urea. Our results suggest that although NUF foliar spray suppresses nitrogen uptake genes, possibly due to nitrogen excess, it enhances growth by up-regulating the synthesis of essential biomolecules and growth-promoting genes, compared to bulk urea.

## Introduction

Depletion of soil nutrients is the prime concern directly associated with food insecurity in underdeveloped and developing countries due to the overuse of land for agriculture (Dimkpa et al. 2023). The input of chemical fertilizers is the current prominent solution to maintain the soil nutrient balance and enhance food productivity without expanding the existing agricultural area. Applying chemical fertilizers has substantially enhanced crop production and ensured food security (Bockman et al. 1990). Although chemical fertilizers are promising for reaching the targets of ever-expanding crop productivity demands, their overuse is posing severe threats to the environment polluting air, water, and soil (Kumar et al. 2019). Research indicates that 40-70% of nitrogen fertilizers are lost to the environment by volatilization or leaching and not used by crops (Guo et al. 2005). The solution to this problem demands a fertilizer that can be uptaken by crops easily, and fewer amounts of the fertilizer than conventional chemical fertilizers can fulfill the plant’s nutrient requirements, resulting in enhanced productivity. Over the past ten years, nano fertilizers have drawn the interest of environmentalists and soil scientists because of their potential for precise and controlled delivery of nutrients, boosting production, enhancing soil fertility, lowering pollution, and creating an environment that is conducive to beneficial soil microbes (Ahmed et al. 2012; Seleiman et al. 2020; Kumar et al. 2019). At a size smaller than the stomatal or cuticular apertures of the plant leaves, nano fertilizers effectively penetrate plant tissues, leading to increased uptake and nutrient use efficiency (Dimkpa et al. 2015; Qureshi et al. 2018; Reddy et al. 2024).

A liquid nano urea formulation (NUF) was recently patented (patent no. 508189) by the Indian Farmers Fertiliser Cooperative (IFFCO). Field trials have demonstrated higher efficiency of NUF over conventional bulk urea (Kumar et al. 2019). The recommended application of this formulation has shown promising results in various crops, enhancing their productivity. For example, combining two foliar sprays of NUF and prilled urea led to a higher yield of maize under 25% reduced nitrogen application (Rawat et al. 2024). However, the molecular mechanisms behind the action of NUF remain largely unknown. Molecular biological studies in various plants revealed differential gene regulation by nano versus bulk/ ionic forms of different nutrients, leading to higher growth (Sun et al. 2020; Ghosh and Bera, 2021) or stress tolerance (Shang et al. 2020) by the nanofertilizers. For example, nano urea led to higher induction of the urea transporter gene *CsDUR3* in cucumber, leading to a higher and extended-duration uptake than bulk urea (Feil et al. 2021). Combinatorial application of nano fertilizers such as nano calcium carbonate or nano carbon synergists together with bulk urea led to a higher induction of the nitrate transporter and assimilation genes (*NRT*), and glutamine synthase (*GS*), compared to the application of only bulk urea (Yang et al. 2023). The *AQP2* aquaporin expression was much higher in tomato roots under nanocarbon application in tissue culture than under traditional carbon (Khodakovskaya et al. 2010). ZnO nanoparticles increased the expression of the antioxidant *SOD* and *GPX* genes in tomato plants (Alharby et al. 2016). Magnetite nanoparticle application in hydroponic media resulted in a dramatic increase in the photosynthesis genes of barley (Tombuloglu et al. 2019). Additionally, photosynthesis and the activities of several vital nitrogen assimilation enzymes in spinach, such as glutamic-pyruvic transaminase, glutamine synthase, nitrate reductase, and glutamate dehydrogenase, were improved by seed treatment with nano-anatase TiO_2_ (Yang et al. 2006). Foliar spray application of FeS nanoparticles induced ribulose-1,5-bisphosphate carboxylase/oxygenase (*rubisco*), *GS,* and glutamate synthase (*GOGAT*) genes in *Brassica juncea* (Rawat et al. 2017).

*A. thaliana* is the laboratory model representing higher plants, including all crops, with many genomic resources available, making it suitable for the mechanistic investigation of NUF foliar spray. In this study, we aim to compare the effect of foliar sprays of NUF versus equimolar bulk urea on *Arabidopsis thaliana*’s vegetative growth and unravel the molecular mechanisms responsible for the growth patterns observed through comparative transcriptome analysis, providing insights into the differential mode of action of the nano versus conventional form of urea in plants.

## Materials and Methods

### Particle size determination of NUF

NUF was purchased from the market (batch number IPNU291440224; manufactured on July 13, 2023). A 0.4% NUF aqueous solution of NUF was prepared, as recommended by IFFCO, for all physical characterizations and subsequent biological experiments. The particle size of 1 mL of a 0.4% aqueous solution of NUF was estimated through dynamic light scattering (DLS) using Malvern Zetasizer Ultra Red Label (AIMIL, New Delhi, India) at a temperature of 25℃. Before measurements, the cuvettes were thoroughly cleaned with ethanol and deionized water to eliminate impurities. Each reading comprised three one-minute runs, and the mean value was recorded. The same instrument also computed the surface charge of NUF particles by measuring the zeta potential. The particle size of the 0.4% NUF solution was further investigated through transmission electron microscopy (TEM) (model LVEM-5, Delong Instruments, Brno, Czech Republic).

### Plant materials and foliar fertilization with NUF

The *A. thaliana* Col-0 ecotype seeds were incubated at 4°C for three days and germinated on nylon meshes floated on ¼ Hoagland’s solution (Hoagland and Arnon 1938), as described earlier (Jangir et al. 2024). The plants were grown for 15 d under hydroponic conditions at 22 ± 2°C and 60 ± 5% humidity and a 12 h photoperiod (light intensity: 80 µmol m^-2^ s^-1^). After 15 d, equal-sized plants were transferred with forceps from the meshes to sterilized vermiculite in 2.5” plastic pots. The potted plants were grown under the same temperature, light, and humidity for 15 d. During the growth regime, the vermiculite in each pot was watered in two-day intervals with an equal volume of ¼ Hoagland’s solution. This diluted medium contained 1 mM Ca(NO_3_)_2_.4H_2_O0.24 mM NH_4_H_2_PO_4_, and 1.5 mM KNO_3_ as nitrogen sources (Jangir et al. 2024). The pots were watered with deionized water on the remaining days. NUF contains 1.5 M urea as mentioned on the product datasheet. To prepare a urea solution of molarity equal to 0.4% NUF solution, we prepared a 1.5 M urea stock by dissolving granular urea in deionized water and diluting it to 0.4% vol/vol strength. One-month-old plants grown as described above were sprayed on both sides of the leaves with 0.4% NUF or 0.4% bulk urea twice at intervals of 10 d, and the growth was observed 20 d after the first spray.

### Phenotyping Arabidopsis

The growth attributes of potted *A. thaliana* plants 20 d after foliar sprays with NUF, bulk urea, and mock spray were evaluated by evaluating the rosette diameter and biomass. The rosette diameters were computed from photographs taken at a uniform distance using the LIA32 software as described earlier (Sadhukhan et al. 2017). Subsequently, the rosettes were excised and weighed in a digital balance (Aczet Pvt Ltd, Mumbai, India). Ten plants for each treatment were used to record the phenotype data.

### Measurement of leaf chlorophyll, nitrogen, and amino acid content

The second rosette leaves were harvested from NUF, urea, and mock-sprayed *A. thaliana* plants, and the total chlorophyll content was quantified following Arnon’s (1949) procedure. Leaves were macerated using a mortar pestle with 80% acetone, and centrifuged at 10,000 rpm for 10 minutes. A UV spectrophotometer (Shimadzu, Kyoto, Japan) was used to measure the absorbance of the samples at 663 nm and 645 nm wavelengths. The total chlorophyll (in mg) was estimated using the equation (8.2 × A_663 nm_) + (20.2 × A_645 nm_) and expressed per gram of leaf fresh weight. The total leaf nitrogen content was measured as described by Koistinen et al. (2019) and Jangir et al. (2024). One gram of leaf sample was homogenized, resuspended in 2 ml deionized water, and centrifuged at 10,000 rpm for 10 minutes. Subsequently, 5 mL of a solution containing 10 g/L K_2_S_2_O_8_ and 6 g/L H_3_BO_3_ in 75 mM NaOH was introduced to the sample. Right after, 3 mL of a reagent, prepared by mixing 40 mL of 1% sulfanilamide (solvent: 2.4 N HCl), 200 mL of 0.5% VCl₃ (solvent: 1.2 N HCl), and 40 mL of 0.07% N-(1-naphthyl)-ethylenediamine dihydrochloride, were added. Next, the sample was incubated at 45°C for 30 minutes, and finally, its absorbance was recorded at 545 nm. We interpolated the total tissue nitrogen levels from a standard curve of Na_2_EDTA. The total leaf amino acid content was quantified according to Doi et al. (1981). Samples were homogenized and resuspended in 2 mL of a 1:1:2 mixture of chloroform, methanol, and deionized water, centrifuged at 10,000 rpm for 10 minutes at 4°C. Next, the upper phase was mixed with 1 mL of ninhydrin reagent and 4 mL of deionized water. Ninhydrin reagent contained 4% ninhydrin in 2-methoxyethanol and 1.6 g/L SnCl₂·2H₂O in 0.2 M citrate buffer (pH 5.0). The mixture was heated at 90°C for 15 min and cooled down. At last, 1 mL of ethanol was added, and the absorbance was measured at 545 nm. We interpolated the milligrams of total amino acids from the absorbance using a standard curve of pure L-leucine and finally expressed per gram leaf fresh weight.

### Transcriptome analysis

The second rosette leaves of *A. thaliana* plants sprayed twice with NUF or urea were harvested 20 days after the first spray and promptly subjected to snap freezing using liquid nitrogen. RNA isolation and downstream analysis were conducted as described previously (Marik et al. 2024). The leaves were homogenized through 0.1% DEPC-treated and twice-autoclaved mortar pestles for total RNA extraction using the CTAB-LiCl method. After that, the integrity and quality of the RNA were evaluated using the Agilent 4150 TapeStation system (Agilent Technologies, California, USA) and further quantified with the Qubit 4 fluorometer from Thermo Fisher Scientific. Utilizing the NEBNext^®^ Ultra^TM^ II RNA Library Prep Kit for Illumina^®^ (New England Biolabs, Massachusetts, USA), RNA-Seq libraries were prepared and sequenced on Illumina’s advanced NovaSeq 6000 V1.5 platform (Illumina Inc., California, USA), yielding 150 bp paired-end reads. We checked the raw data quality using in-house scripts, trimmed the adapters at Q30 by AdapterRemoval v2.3.2 (Schubert et al. 2016), and aligned them to the *A. thaliana* genome (NCBI assembly ID: GCF000001735.4) using HISAT v2.2.1 (Zhang et al. 2021). We determined the mRNA levels using featureCounts v2.0.3 (Liao, Smyth, and Shi 2014) and conducted differential gene expression (DEG) analysis using edgeR version 3.42.4 (Chen, Lun, and Smyth 2016). The BAR Classification SuperViewer application was used to calculate the functional enrichment analysis (Provert and Zhu 2003).

### Real-time PCR analysis

Employing quantitative real-time quantitative PCR (qPCR), the expression levels observed in the transcriptome analysis were validated. Twenty days after the NUF or urea foliar sprays, RNA was isolated from the second rosette leaves of *A. thaliana* and converted to cDNA using the RevertAid First Strand cDNA Synthesis Kit (Thermo Fisher Scientific, Powai, India). The Primer3 web application (https://primer3.ut.ee/) was used to design the q-PCR primers.

### Statistical analysis

Ten *A. thaliana* plants as independent biological replicates were used for each treatment for all phenotypic and biochemical analyses, and mean values with the standard error were reported in all cases. The entire experiments were repeated three times. Student’s *t*-test compared two samples, whereas Tukey’s HSD test was used to compare multiple samples. Significant differences were considered at a *P*-value < 0.05. Three biological replicates were used for the transcriptome and qPCR analyses. The comparisons in the transcriptome analysis were subjected to Fisher’s exact test with the Benjamini-Hochberg *false-discovery rate* (*FDR*) correction in *P*-values.

## Results

### Particle size of NUF is suitable for foliar uptake

Particle dimension and surface charge influence plants’ ability to uptake nano fertilizers. Our DLS measurements determined the particle size of 0.4% aqueous solution of NUF, yielding a measurement of 27.77 nm. TEM also verified the size variation within the 20 to 35 nm range. These measured particle sizes suggested the suitability of foliar uptake of NUF through stomatal conduits. Moreover, the zeta potential of NUF was -54.74 mV, indicating its anionic properties in an aqueous environment, preventing agglomeration.

### Foliar sprays of NUF enhance biomass, chlorophyll, and nitrogen content of leaves than equimolar bulk urea

NUF was applied as a foliar spray to one-month-old *A. thaliana* plants grown in vermiculite, and the effects on several phenotypes, including biomass, chlorophyll content, nitrogen, and amino acid content, were measured. Vermiculite contains no nitrogen source and was chosen for uniformity in all treatments. Every alternate day the vermiculite in each pot received basal nitrogen fertilization, in the form of nitrate and ammonium, through an equal amount of ¼ Hoagland’s solution. Foliar NUF sprays in increasing concentrations of 0.3%, 0.4%, and 0.5% were tested along with equimolar doses of bulk urea and mock sprays with only deionized water (Fig. S1). After 20 days of two sprays in 10-day intervals, a significant increase in rosette diameter and biomass was observed in both NUF and urea treatments, compared to the mock treatments (Fig. 1A). The 0.4% v/v NUF dose produced the highest increase of 51 ± 14.9% in shoot biomass and 10.2 ± 2.8% in rosette diameter compared to equimolar urea. Hence, this dose was used for all further phenotyping experiments (Fig. 1B, S1). We observed a significant 29.5 ± 9.1% increase in total chlorophyll, 8.4 ± 3.1% increase in total leaf nitrogen, and 4.5 ± 0.3% increase in total amino acid levels after 0.4% NUF treatment compared to equimolar urea (Fig. 1C-E).

**Fig 1.**
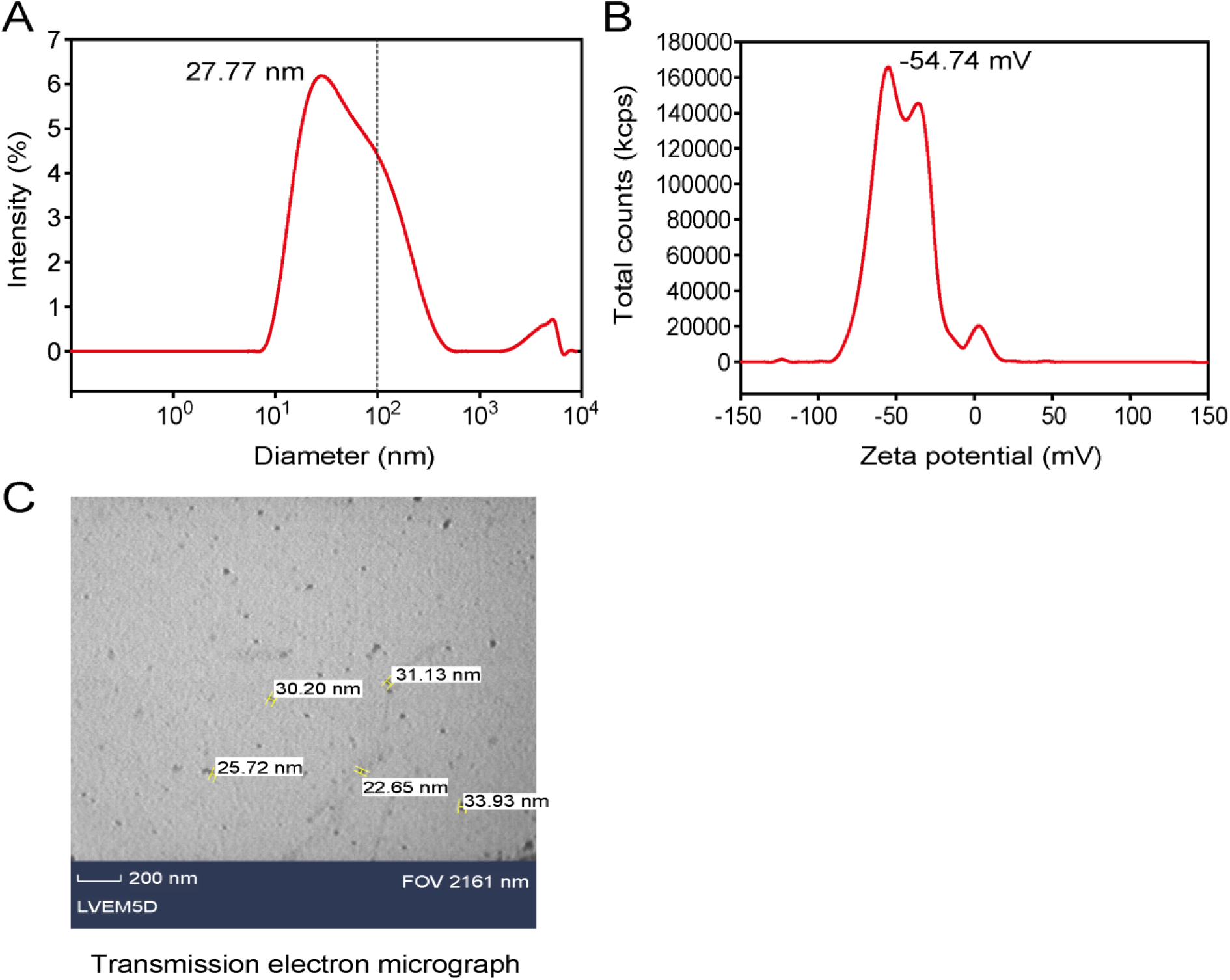
Dynamic light scattering and transmission electron microscopic analysis of IFFCO nano urea. The particle size of a 0.4% aqueous solution of IFFCO liquid nano urea formulation (NUF) was determined by dynamic light scattering (DLS) and transmission electron microscopy (TEM). The DLS results are shown as a plot of scattered light intensity and particle size in nanometers (A). The scattered photon count expressed in kilo counts per second (kcps) was plotted against the zeta potential in mV (B). A transmission electron micrograph (C). The black scale bar denotes 200 nm. Five randomly selected particles are shown with their diameters in nanometers. FOV: field of view 2161 nm.

**Fig. 2.**
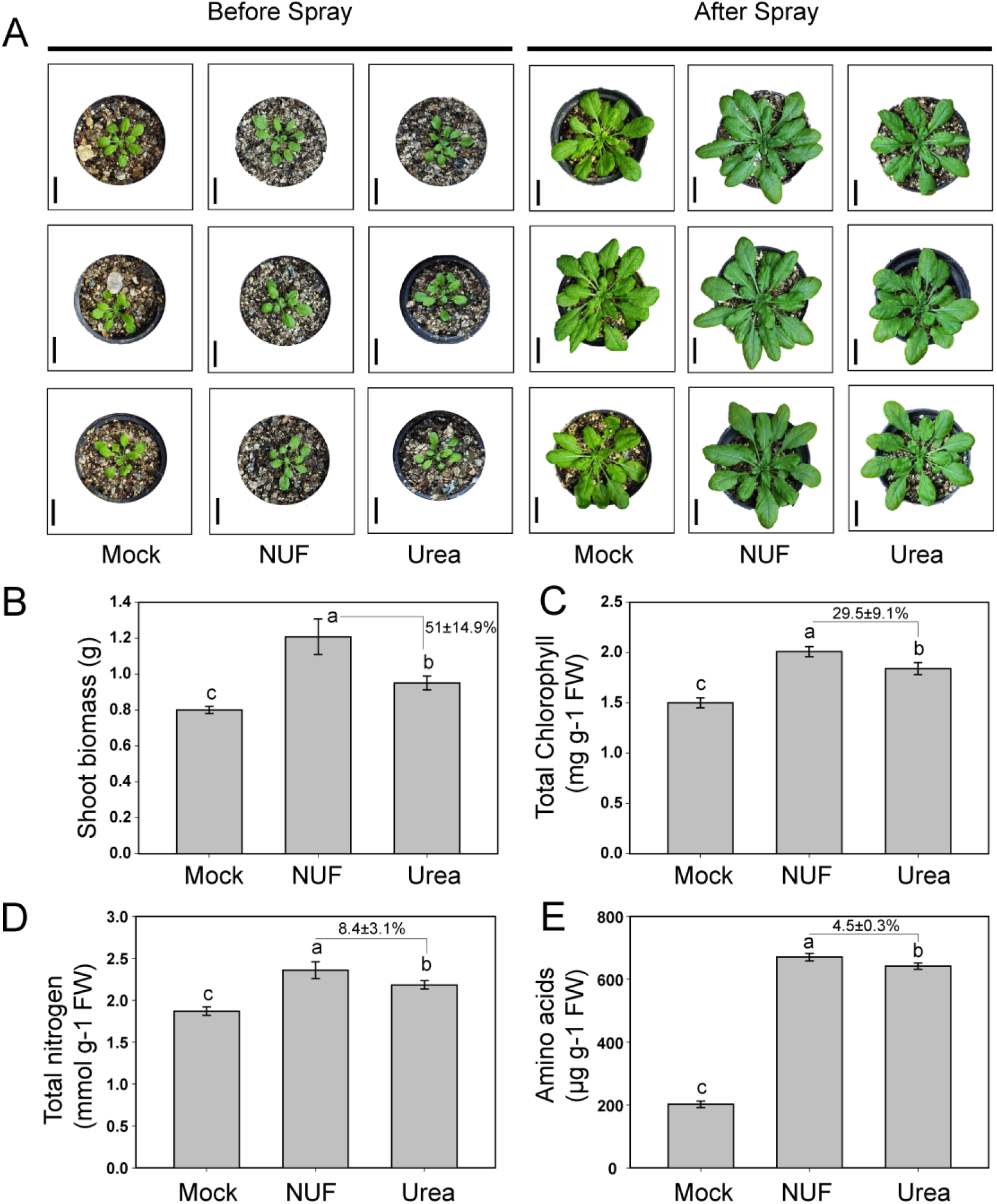
Phenotypic effects of IFFCO nano urea foliar spray on *Arabidopsis thaliana.* The effects of IFFCO liquid nano urea formulation (NUF) versus equimolar bulk urea on biomass, chlorophyll, total leaf nitrogen, and amino acid contents are shown. Arabidopsis plants were grown for a month in vermiculite in 2.5” pots watered with ¼-Hoagland’s solution every alternate day. NUF contains 1.5 M urea, as per the manufacturer. For uniformity, a stock aqueous solution of 1.5 M Urea was prepared. 0.4% dilutions were prepared from the urea and NUF stock solutions and used to spray one-month-old *A. thaliana* plants. The plants were sprayed twice in 10-d intervals with 0.4% NUF or 0.4% urea at both the abaxial and adaxial leaf surfaces. Mock plants were sprayed with distilled water and photographed. The shoots were harvested to measure biomass, total chlorophyll, total nitrogen, and total amino acid content after 20 d. The shoot fresh weights were measured after 20-d growth in mock, NUF, and urea and presented as biomass in grams (**A**). Bars indicate the average of 10 seedlings, with standard error. The total chlorophyll contents of seedling shoots in milligrams per gram of fresh tissue weight after 20-d growth in mock, NUF, and urea are shown (**B**). Bars indicate the average of 10 replicates, with standard error. The total tissue nitrogen contents (**C**) and amino acids (**D**) in shoots are presented after mock, urea, or NUF spray treatments. The shoots were separated and homogenized using a mortar and pestle, and nitrogen and amino acid contents were measured spectrophotometrically by biochemical tests. Bars indicate the average of 10 replicates with standard error. Different alphabets indicate significant differences (Tukey’s HSD test, *P* < 0.05, N = 10).

### Different gene expression patterns are observed after foliar sprays of NUF and equimolar bulk urea

To understand NUF’s and bulk urea’s differential molecular modes of action, we conducted a transcriptome analysis of *A. thaliana* rosette leaves 20 days after foliar sprays with NUF, urea, and mock sprays. NUF spray resulted in significant up-regulation of 493 genes (log_2_fold-change > 0.5, *FDR* < 0.05) than the mock spray, while significantly down-regulating 350 genes (log_2_FC < -0.5, *FDR* < 0.05) (Fig. 3A, Table S1). However, more genes were up-regulated (1393) or down-regulated (512) by urea than mock, at the same significance levels (Fig. 3B). But when compared between NUF and urea treatments, we found that 234 genes were significantly induced (log_2_FC > 0.5, *FDR* < 0.05) by NUF than urea, and 1321 genes significantly suppressed (log_2_FC < -0.5, *FDR* < 0.05) by NUF than urea (Fig. 3C). A correlation analysis of significant DEGs (-0.5 > log_2_FC > 0.5, *FDR* < 0.05) indicated that the expression patterns of all genes did not correlate between NUF and urea, each compared to the mock treatment (Fig. 3D). The common DEGs under various treatments were visualized by Venn diagrams. NUF and urea up-regulated 121 common genes compared to the mock treatment. NUF led to the induction of 61 genes compared to both the mock and urea treatments (Fig. 3E). Conversely, 69 genes were down-regulated by both NUF and urea compared to the mock treatments. NUF suppressed 209 genes compared to mock and urea (Fig. 3F).

**Fig. 3.**
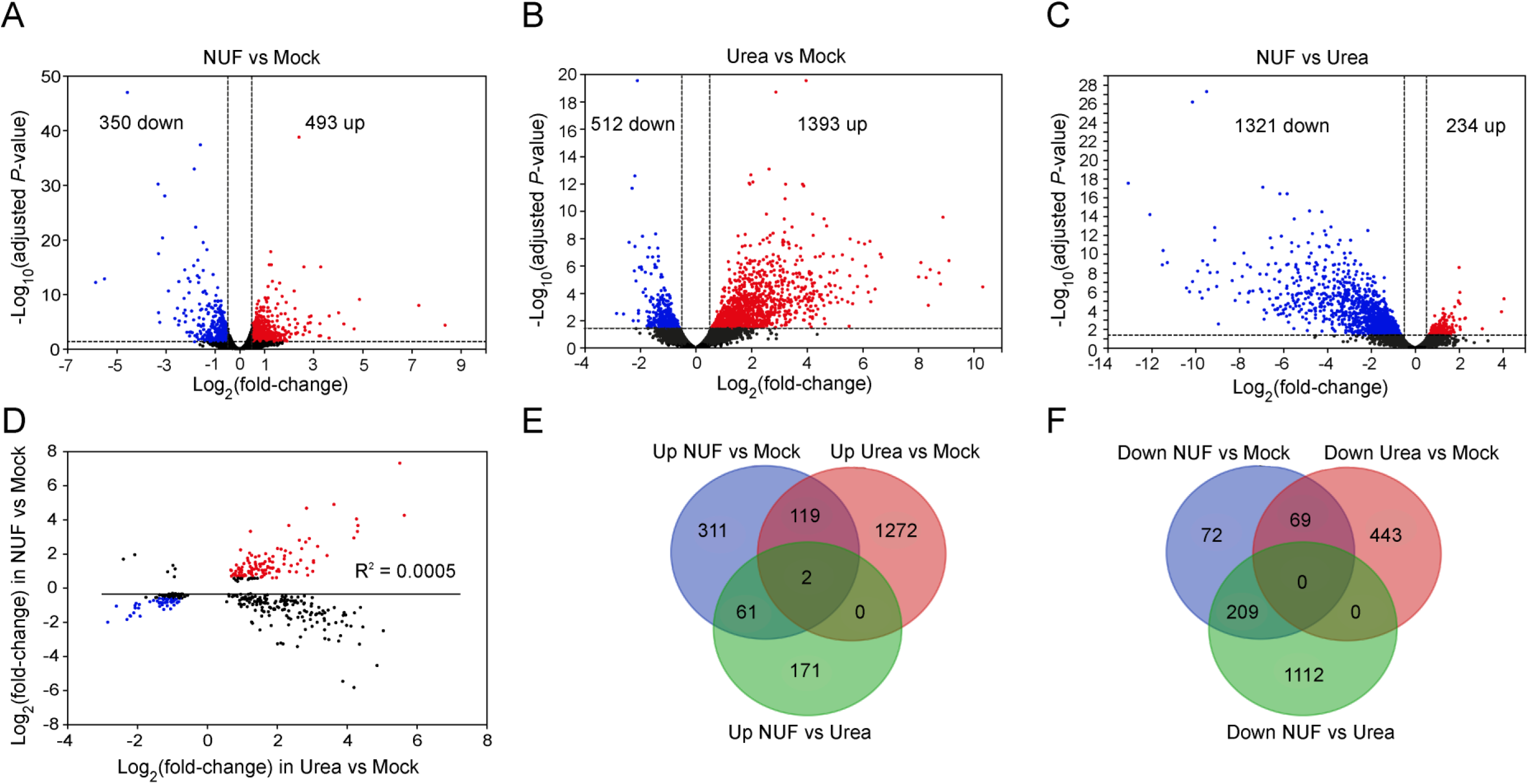
Transcriptome analysis of *Arabidopsis thaliana* leaves after foliar spray with IFFCO nano urea. One-month-old *A. thaliana* plants grown in vermiculite and fertilized every alternate day with ¼-Hoagland solution were sprayed on both sides of the leaves with 0.4% IFFCO liquid nano urea formulation (NUF) or 0.4% bulk urea twice at intervals of 10 days. After 20 days from the first spray, rosette leaves were harvested, and the transcriptome was analyzed by RNA sequencing. Mock treatment included sprays with distilled water. Volcano plots of NUF versus mock (**A**), urea versus mock (**B**), and NUF versus urea (**C**) are shown. The Y-axes present -log_10_(adjusted *P*-value), i.e., Benjamini-Hochberg *false discovery rate* (*FDR*)-adjusted *P*-values (N = 3), and X-axes present log_2_(fold-change). The horizontal dotted lines demarcate an *FDR* < 0.05, and the vertical dotted lines demarcate a log_2_(fold-change) of 0.5. Up-regulated [log_2_(fold-change) > 0.5, *FDR* < 0.05] and down-regulated [log_2_(fold-change) < -0.5, *FDR* < 0.05] genes are shown by red and blue dots, respectively. (**D**) A correlation plot of log_2_(fold-change) of differential gene expression under NUF and urea [-0.5 > log_2_(fold-change) > 0.5, *FDR* < 0.05], each compared to the mock treatment, is shown. Red and blue dots indicate genes up- or down-regulated under NUF as well as urea. Venn diagrams of genes up-regulated (**E**) or down-regulated (**F**) in the three comparisons, NUF vs. mock, urea vs. mock, and NUF versus urea, are shown.

To dissect the functional roles of the DEGs under foliar sprays of NUF versus urea (Fig. 3C), we conducted an enrichment analysis using the GO biological process and Mapman classification databases. The results pointed out that NUF treatment led to greater induction of genes involved in “growth”, “carbohydrate and lipid metabolic processes”, “cell wall”, “major CHO-metabolism”, “nucleotide metabolism”, “hormone metabolism”, “anatomical structure development”, “multicellular organism development”, “post-embryonic development”, “signal transduction”, “response to abiotic stimulus”, “response to stress”, “response to light stimulus”, etc., than equimolar bulk urea (Fig. 4A, B). On the other hand, genes down-regulated by NUF compared to urea belonged to the functional categories “cell death”, “N-metabolism”, “abscission”, “minor CHO-metabolism”, “response to biotic stimulus”, “transport”, etc. (Fig. 4C, D). Noteworthy, processes like “cell communication”, “response to stress”, “response to chemical”, “secondary metabolic process”, “signal transduction”, “response to endogenous stimulus”, “hormone metabolism”, etc. were observed in both the up and down-regulated genes, pointing to the common biological processes targetted by both urea and NUF treatments.

**Fig. 4.**
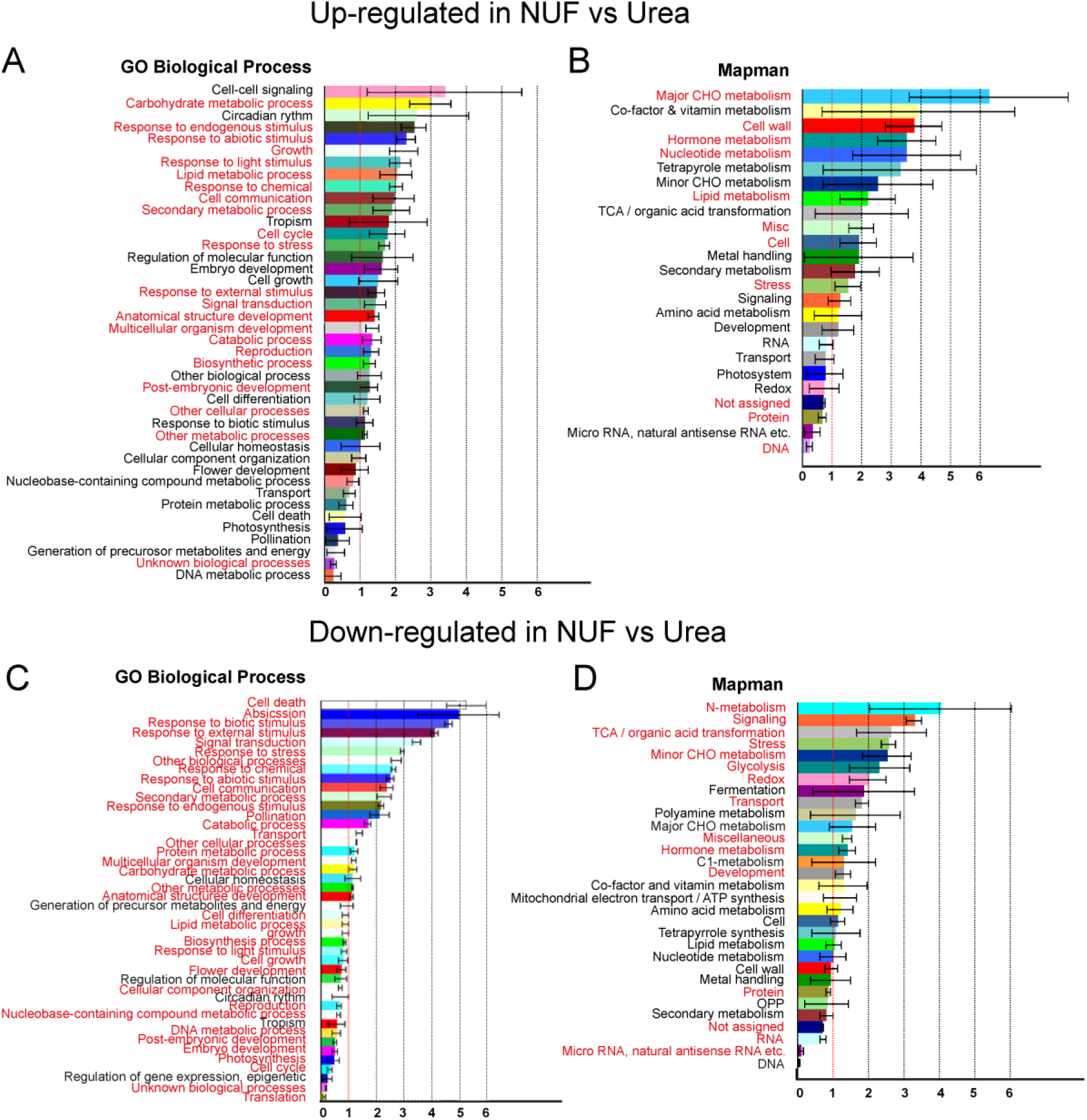
Functional enrichment analysis of *Arabidopsis thaliana* genes differentially expressed after foliar sprays with IFFCO nano urea versus equimolar bulk urea. An enrichment analysis of 234 genes up-regulated (log_2_fold-change > 0.5; *FDR* < 0.05; *N* = 3) or 1321 genes down-regulated (log_2_fold-change < -0.5; *FDR* < 0.05; *N* = 3) in *A. thaliana* leaves under IFFCO liquid nano urea formulation (NUF) versus equimolar bulk urea (see Table S1) was conducted by the BAR classification SuperViewer tool (https://bar.utoronto.ca/). The classification source was “Gene Ontology Biological Process” or “Mapman” for the up- (**A, B**) or down-regulated genes (**C, D**), respectively. The lengths of the bars along the X-axes represent normed frequencies, defined by the frequency of occurrence in the up- (234) or down-regulated (1321) gene sets versus that in genome-wide genes of *A. thaliana*, with standard deviation. The enriched categories (*FDR* < 0.05) are highlighted in red. *FDR*: Benjamini-Hochberg *false discovery rate*-corrected *P*-value.

### Nitrogen metabolism genes are differentially regulated by foliar sprays of NUF and urea

To compare the differences between NUF and urea as nitrogen fertilizers, we analyzed their effects on the expression of nitrogen uptake and metabolism genes of *A. thaliana* (Table 1). We validated the expression levels of randomly selected nitrogen transporter and metabolism genes under NUF and urea treatments by qPCR (Fig. 5; primer sequences provided in Table S2).

**Fig. 5.**
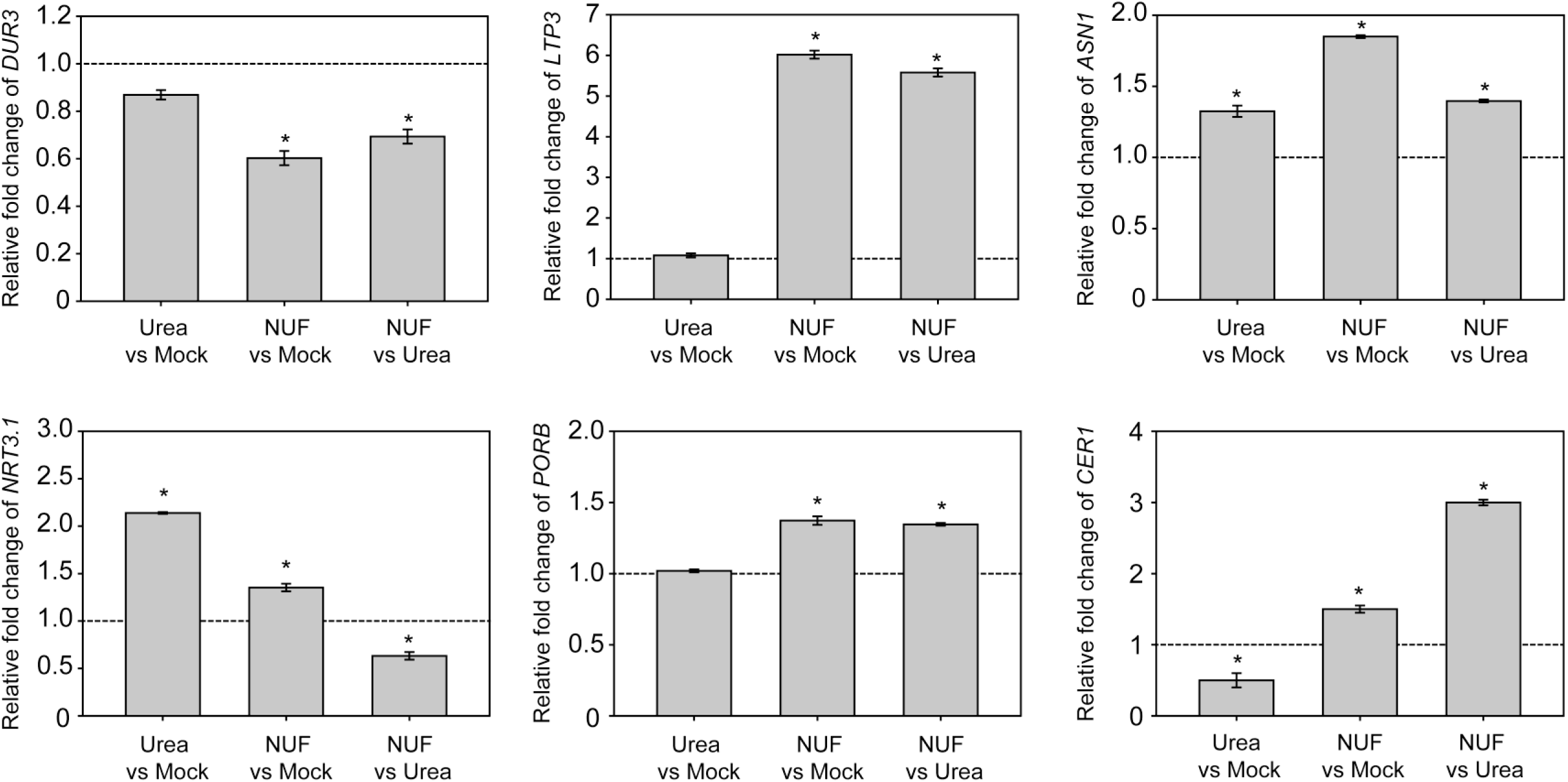
Real-time expression analysis of *Arabidopsis thaliana* genes differentially expressed after foliar sprays with IFFCO nano urea versus equimolar bulk urea. One-month-old *A. thaliana* plants grown in vermiculite and fertilized every alternate day with ¼-Hoagland solution in the pots were sprayed on both sides of the leaves with 0.4% IFFCO liquid nano urea formulation (NUF) or 0.4% bulk urea twice at intervals of 10 days. After 20 days from the first spray, rosette leaves were harvested to isolate RNA and synthesize cDNA to estimate relative expression levels using real-time PCR. The relative fold-change in expression levels of the six randomly chosen genes in the transcriptome analysis (Table 1, 2), viz., *DUR3*, *LPT3*, *ASN1*, *NRT3.1*, *PORB,* and *CER1*, between urea (treatment) and mock (control), NUF (treatment) and mock (control), and NUF (treatment) and urea (control) treated *A. thaliana* samples are shown in the graph. The expression levels in each sample were further normalized using *AtUBQ1* as the internal standard. The bars indicate an average of three biological replicates comprising six rosette leaves with standard error. The dotted lines indicate control expression levels for each comparison. Asterisks above the error bars indicate significant differences in gene expression between treatment and control samples (*P* < 0.05, Student’s *t*-test, *N* = 3 biological replicates).

**Table 1.**
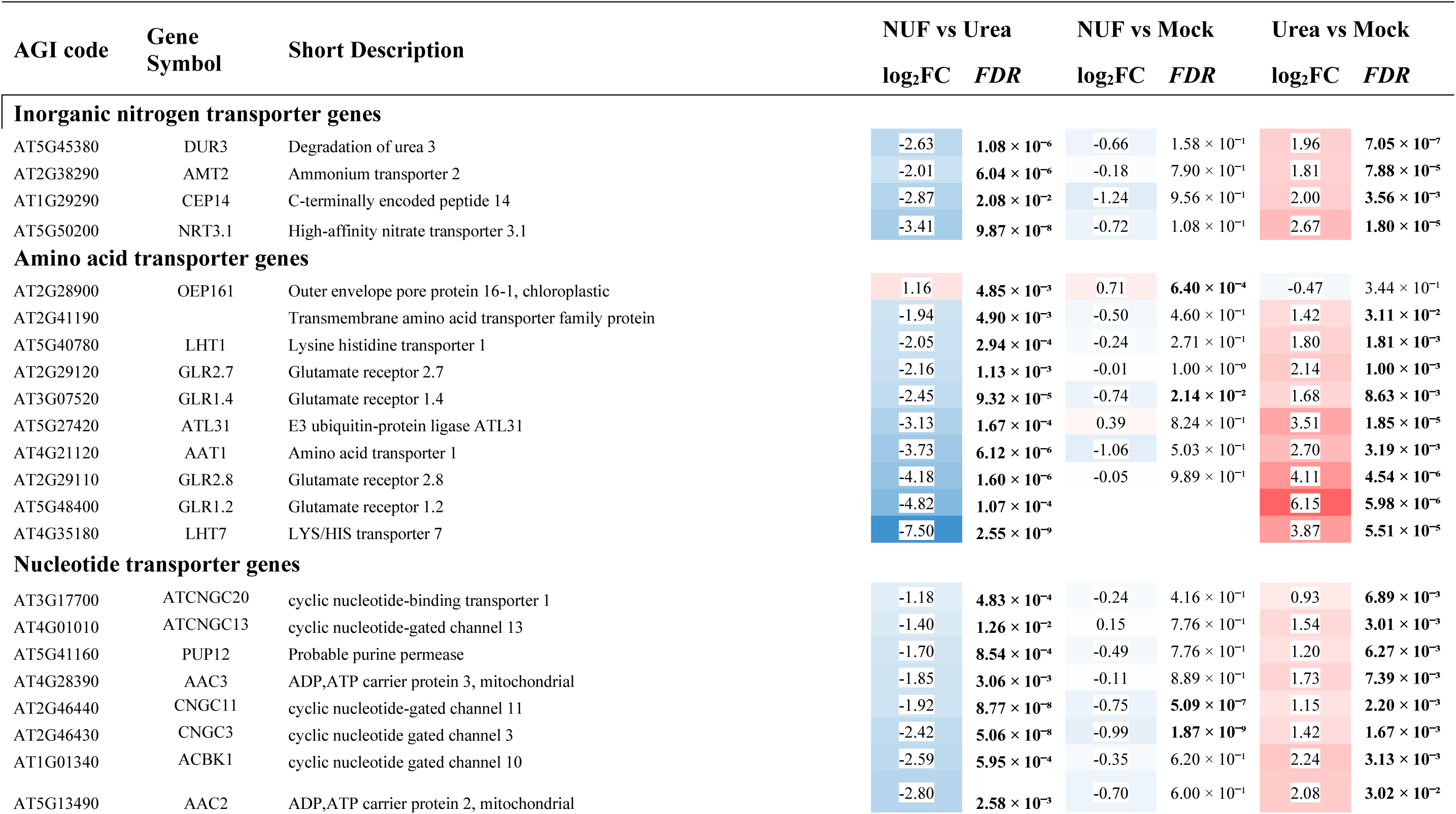
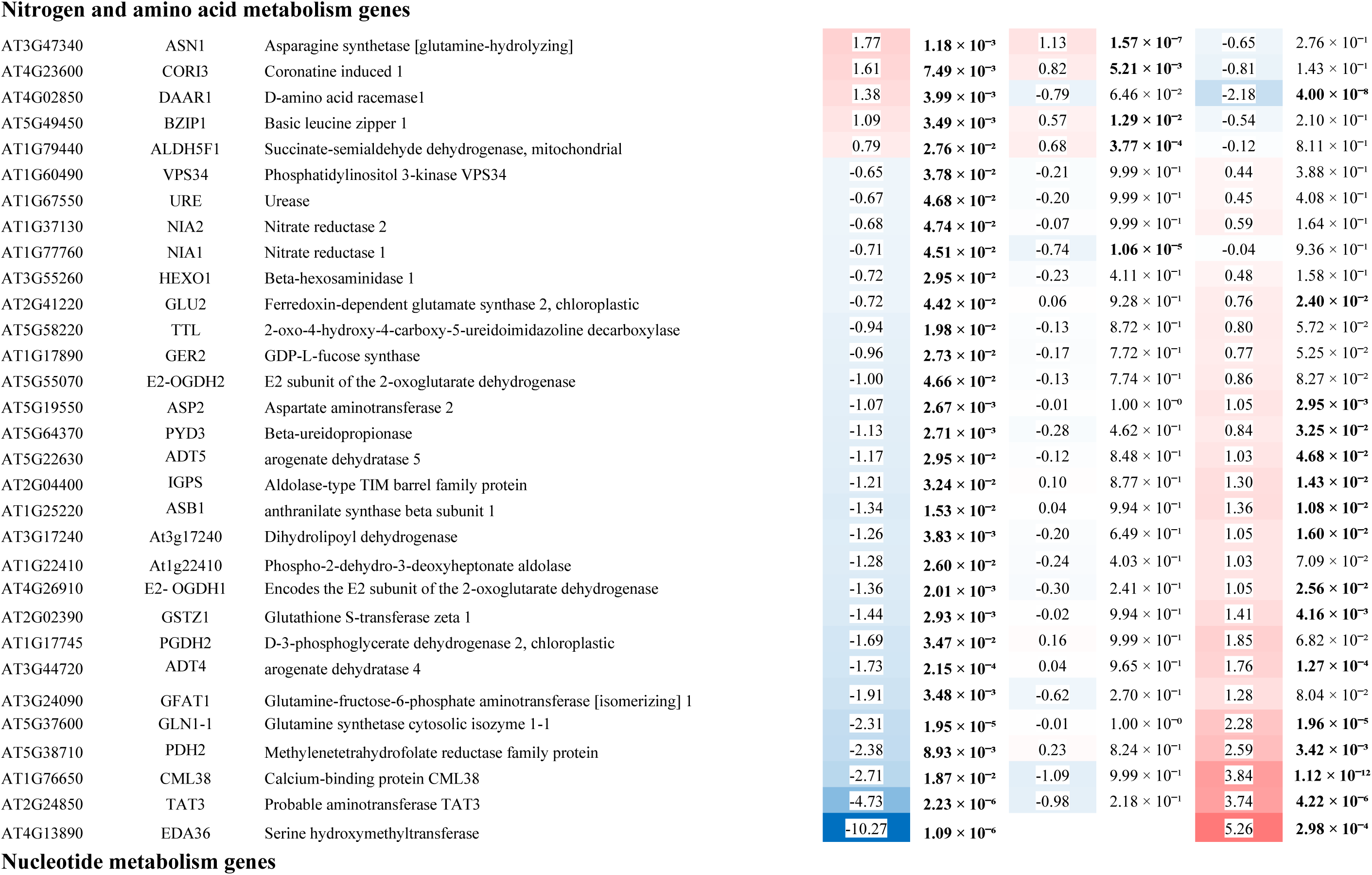
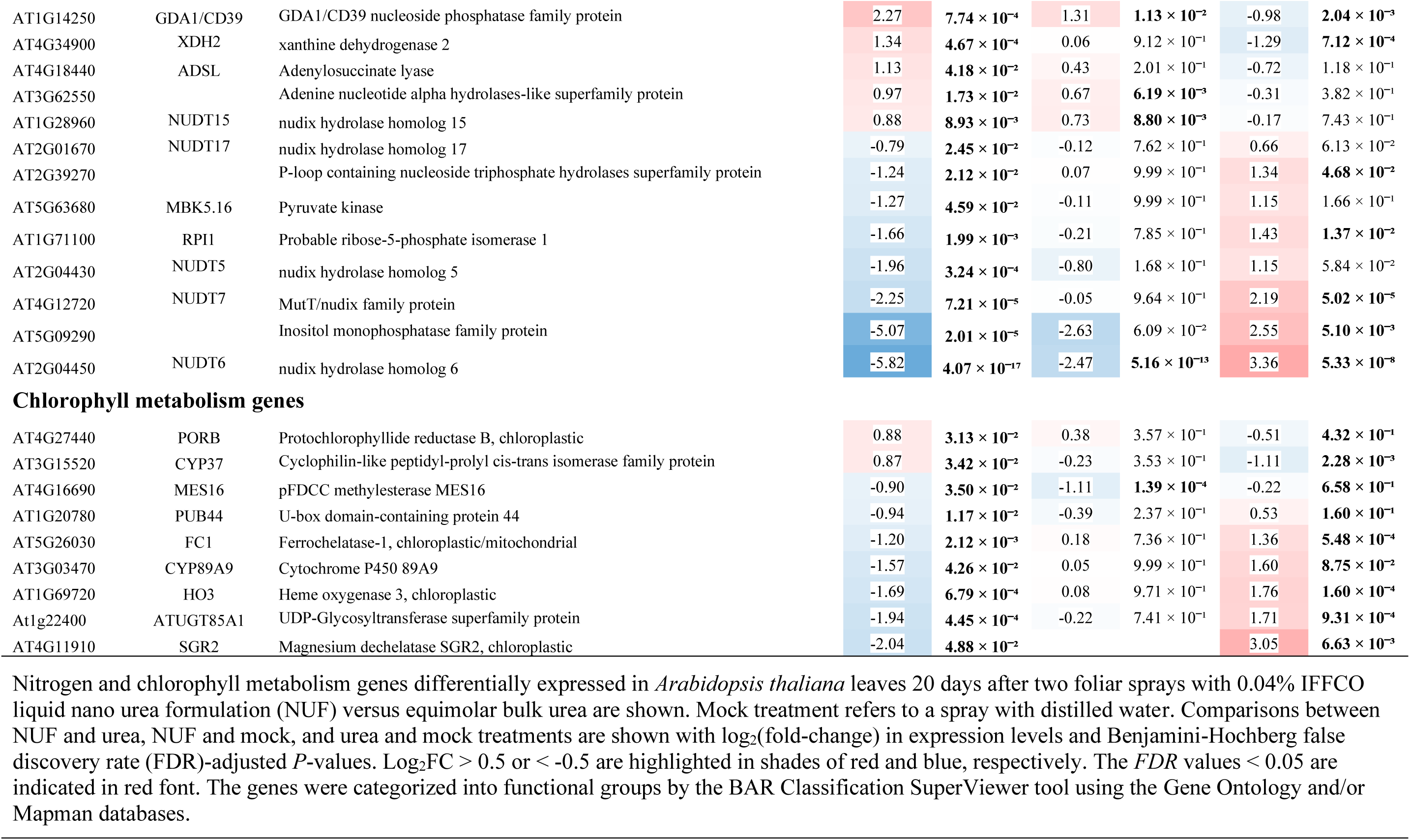
Nitrogen and chlorophyll metabolism genes differentially expressed in *Arabidopsis thaliana* leaves by foliar sprays of IFFCO nano urea versus equimolar bulk urea.

| AGI code | Gene Symbol | Short Description | NUF vs Urea |  | NUF vs Mock |  | Urea vs Mock |  |
| --- | --- | --- | --- | --- | --- | --- | --- | --- |
|  |  |  | log<sub>2</sub>FC | FDR | log<sub>2</sub>FC | FDR | log<sub>2</sub>FC | FDR |
| Inorganic nitrogen transporter genes |  |  |  |  |  |  |  |  |
| AT5G45380 | DUR3 | Degradation of urea 3 | -2.63 | 1.08 × 10<sup>-6</sup> | -0.66 | 1.58 × 10<sup>-1</sup> | 1.96 | 7.05 × 10<sup>-7</sup> |
| AT2G38290 | AMT2 | Ammonium transporter 2 | -2.01 | 6.04 × 10<sup>-6</sup> | -0.18 | 7.90 × 10<sup>-1</sup> | 1.81 | 7.88 × 10<sup>-5</sup> |
| AT1G29290 | CEP14 | C-terminally encoded peptide 14 | -2.87 | 2.08 × 10<sup>-2</sup> | -1.24 | 9.56 × 10<sup>-1</sup> | 2.00 | 3.56 × 10<sup>-3</sup> |
| AT5G50200 | NRT3.1 | High-affinity nitrate transporter 3.1 | -3.41 | 9.87 × 10<sup>-8</sup> | -0.72 | 1.08 × 10<sup>-1</sup> | 2.67 | 1.80 × 10<sup>-5</sup> |
| Amino acid transporter genes |  |  |  |  |  |  |  |  |
| AT2G28900 | OEP161 | Outer envelope pore protein 16-1, chloroplastic | 1.16 | 4.85 × 10<sup>-3</sup> | 0.71 | 6.40 × 10<sup>-4</sup> | -0.47 | 3.44 × 10<sup>-1</sup> |
| AT2G41190 |  | Transmembrane amino acid transporter family protein | -1.94 | 4.90 × 10<sup>-3</sup> | -0.50 | 4.60 × 10<sup>-1</sup> | 1.42 | 3.11 × 10<sup>-2</sup> |
| AT5G40780 | LHT1 | Lysine histidine transporter 1 | -2.05 | 2.94 × 10<sup>-4</sup> | -0.24 | 2.71 × 10<sup>-1</sup> | 1.80 | 1.81 × 10<sup>-3</sup> |
| AT2G29120 | GLR2.7 | Glutamate receptor 2.7 | -2.16 | 1.13 × 10<sup>-3</sup> | -0.01 | 1.00 × 10<sup>-0</sup> | 2.14 | 1.00 × 10<sup>-3</sup> |
| AT3G07520 | GLR1.4 | Glutamate receptor 1.4 | -2.45 | 9.32 × 10<sup>-5</sup> | -0.74 | 2.14 × 10<sup>-2</sup> | 1.68 | 8.63 × 10<sup>-3</sup> |
| AT5G27420 | ATL31 | E3 ubiquitin-protein ligase ATL31 | -3.13 | 1.67 × 10<sup>-4</sup> | 0.39 | 8.24 × 10<sup>-1</sup> | 3.51 | 1.85 × 10<sup>-5</sup> |
| AT4G21120 | AAT1 | Amino acid transporter 1 | -3.73 | 6.12 × 10<sup>-6</sup> | -1.06 | 5.03 × 10<sup>-1</sup> | 2.70 | 3.19 × 10<sup>-3</sup> |
| AT2G29110 | GLR2.8 | Glutamate receptor 2.8 | -4.18 | 1.60 × 10<sup>-6</sup> | -0.05 | 9.89 × 10<sup>-1</sup> | 4.11 | 4.54 × 10<sup>-6</sup> |
| AT5G48400 | GLR1.2 | Glutamate receptor 1.2 | -4.82 | 1.07 × 10<sup>-4</sup> |  |  | 6.15 | 5.98 × 10<sup>-6</sup> |
| AT4G35180 | LHT7 | LYS/HIS transporter 7 | -7.50 | 2.55 × 10<sup>-9</sup> |  |  | 3.87 | 5.51 × 10<sup>-5</sup> |
| Nucleotide transporter genes |  |  |  |  |  |  |  |  |
| AT3G17700 | ATCNGC20 | cyclic nucleotide-binding transporter 1 | -1.18 | 4.83 × 10<sup>-4</sup> | -0.24 | 4.16 × 10<sup>-1</sup> | 0.93 | 6.89 × 10<sup>-3</sup> |
| AT4G01010 | ATCNGC13 | cyclic nucleotide-gated channel 13 | -1.40 | 1.26 × 10<sup>-2</sup> | 0.15 | 7.76 × 10<sup>-1</sup> | 1.54 | 3.01 × 10<sup>-3</sup> |
| AT5G41160 | PUP12 | Probable purine permease | -1.70 | 8.54 × 10<sup>-4</sup> | -0.49 | 7.76 × 10<sup>-1</sup> | 1.20 | 6.27 × 10<sup>-3</sup> |
| AT4G28390 | AAC3 | ADP,ATP carrier protein 3, mitochondrial | -1.85 | 3.06 × 10<sup>-3</sup> | -0.11 | 8.89 × 10<sup>-1</sup> | 1.73 | 7.39 × 10<sup>-3</sup> |
| AT2G46440 | CNGC11 | cyclic nucleotide-gated channel 11 | -1.92 | 8.77 × 10<sup>-8</sup> | -0.75 | 5.09 × 10<sup>-7</sup> | 1.15 | 2.20 × 10<sup>-3</sup> |
| AT2G46430 | CNGC3 | cyclic nucleotide gated channel 3 | -2.42 | 5.06 × 10<sup>-8</sup> | -0.99 | 1.87 × 10<sup>-9</sup> | 1.42 | 1.67 × 10<sup>-3</sup> |
| AT1G01340 | ACBK1 | cyclic nucleotide gated channel 10 | -2.59 | 5.95 × 10<sup>-4</sup> | -0.35 | 6.20 × 10<sup>-1</sup> | 2.24 | 3.13 × 10<sup>-3</sup> |
| AT5G13490 | AAC2 | ADP,ATP carrier protein 2, mitochondrial | -2.80 | 2.58 × 10<sup>-3</sup> | -0.70 | 6.00 × 10<sup>-1</sup> | 2.08 | 3.02 × 10<sup>-2</sup> |

Nitrogen and amino acid metabolism genes
|  |  |  |
| --- | --- | --- |
| AT3G47340 | ASN1 | Asparagine synthetase [glutamine-hydrolyzing] |
| AT4G23600 | COR13 | Coronatine induced 1 |
| AT4G02850 | DAAR1 | D-amino acid racemase1 |
| AT5G49450 | BZIP1 | Basic leucine zipper 1 |
| AT1G79440 | ALDH5F1 | Succinate-semialdehyde dehydrogenase, mitochondrial |
| AT1G60490 | VPS34 | Phosphatidylinositol 3-kinase VPS34 |
| AT1G67550 | URE | Urease |
| AT1G37130 | NIA2 | Nitrate reductase 2 |
| AT1G77760 | NIA1 | Nitrate reductase 1 |
| AT3G55260 | HEXO1 | Beta-hexosaminidase 1 |
| AT2G41220 | GLU2 | Ferredoxin-dependent glutamate synthase 2, chloroplastic |
| AT5G58220 | TTL | 2-oxo-4-hydroxy-4-carboxy-5-ureidoimidazole decarboxylase |
| AT1G17890 | GER2 | GDP-L-fucose synthase |
| AT5G55070 | E2-OGDH2 | E2 subunit of the 2-oxoglutarate dehydrogenase |
| AT5G19550 | ASP2 | Aspartate aminotransferase 2 |
| AT5G64370 | PYD3 | Beta-ureidopropionase |
| AT5G22630 | ADT5 | arogenate dehydratase 5 |
| AT2G04400 | IGPS | Aldolase-type TIM barrel family protein |
| AT1G25220 | ASB1 | anthranilate synthase beta subunit 1 |
| AT3G17240 | At3g17240 | Dihydrolipoyl dehydrogenase |
| AT1G22410 | At1g22410 | Phospho-2-dehydro-3-deoxyheptonate aldolase |
| AT4G26910 | E2- OGDH1 | Encodes the E2 subunit of the 2-oxoglutarate dehydrogenase |
| AT2G02390 | GSTZ1 | Glutathione S-transferase zeta 1 |
| AT1G17745 | PGDH2 | D-3-phosphoglycerate dehydrogenase 2, chloroplastic |
| AT3G44720 | ADT4 | arogenate dehydratase 4 |
| AT3G24090 | GFAT1 | Glutamine-fructose-6-phosphate aminotransferase [isomerizing] 1 |
| AT5G37600 | GLN1-1 | Glutamine synthetase cytosolic isozyme 1-1 |
| AT5G38710 | PDH2 | Methylenetetrahydrofolate reductase family protein |
| AT1G76650 | CML38 | Calcium-binding protein CML38 |
| AT2G24850 | TAT3 | Probable aminotransferase TAT3 |
| AT4G13890 | EDA36 | Serine hydroxymethyltransferase |

|  |  |  |  |  |  |
| --- | --- | --- | --- | --- | --- |
| 1.77 | $1.18 \times 10^{-3}$ | 1.13 | $1.57 \times 10^{-7}$ | -0.65 | $2.76 \times 10^{-1}$ |
| 1.61 | $7.49 \times 10^{-3}$ | 0.82 | $5.21 \times 10^{-3}$ | -0.81 | $1.43 \times 10^{-1}$ |
| 1.38 | $3.99 \times 10^{-3}$ | -0.79 | $6.46 \times 10^{-2}$ | -2.18 | $4.00 \times 10^{-8}$ |
| 1.09 | $3.49 \times 10^{-3}$ | 0.57 | $1.29 \times 10^{-2}$ | -0.54 | $2.10 \times 10^{-1}$ |
| 0.79 | $2.76 \times 10^{-2}$ | 0.68 | $3.77 \times 10^{-4}$ | -0.12 | $8.11 \times 10^{-1}$ |
| -0.65 | $3.78 \times 10^{-2}$ | -0.21 | $9.99 \times 10^{-1}$ | 0.44 | $3.88 \times 10^{-1}$ |
| -0.67 | $4.68 \times 10^{-2}$ | -0.20 | $9.99 \times 10^{-1}$ | 0.45 | $4.08 \times 10^{-1}$ |
| -0.68 | $4.74 \times 10^{-2}$ | -0.07 | $9.99 \times 10^{-1}$ | 0.59 | $1.64 \times 10^{-1}$ |
| -0.71 | $4.51 \times 10^{-2}$ | -0.74 | $1.06 \times 10^{-5}$ | -0.04 | $9.36 \times 10^{-1}$ |
| -0.72 | $2.95 \times 10^{-2}$ | -0.23 | $4.11 \times 10^{-1}$ | 0.48 | $1.58 \times 10^{-1}$ |
| -0.72 | $4.42 \times 10^{-2}$ | 0.06 | $9.28 \times 10^{-1}$ | 0.76 | $2.40 \times 10^{-2}$ |
| -0.94 | $1.98 \times 10^{-2}$ | -0.13 | $8.72 \times 10^{-1}$ | 0.80 | $5.72 \times 10^{-2}$ |
| -0.96 | $2.73 \times 10^{-2}$ | -0.17 | $7.72 \times 10^{-1}$ | 0.77 | $5.25 \times 10^{-2}$ |
| -1.00 | $4.66 \times 10^{-2}$ | -0.13 | $7.74 \times 10^{-1}$ | 0.86 | $8.27 \times 10^{-2}$ |
| -1.07 | $2.67 \times 10^{-3}$ | -0.01 | $1.00 \times 10^{-0}$ | 1.05 | $2.95 \times 10^{-3}$ |
| -1.13 | $2.71 \times 10^{-3}$ | -0.28 | $4.62 \times 10^{-1}$ | 0.84 | $3.25 \times 10^{-2}$ |
| -1.17 | $2.95 \times 10^{-2}$ | -0.12 | $8.48 \times 10^{-1}$ | 1.03 | $4.68 \times 10^{-2}$ |
| -1.21 | $3.24 \times 10^{-2}$ | 0.10 | $8.77 \times 10^{-1}$ | 1.30 | $1.43 \times 10^{-2}$ |
| -1.34 | $1.53 \times 10^{-2}$ | 0.04 | $9.94 \times 10^{-1}$ | 1.36 | $1.08 \times 10^{-2}$ |
| -1.26 | $3.83 \times 10^{-3}$ | -0.20 | $6.49 \times 10^{-1}$ | 1.05 | $1.60 \times 10^{-2}$ |
| -1.28 | $2.60 \times 10^{-2}$ | -0.24 | $4.03 \times 10^{-1}$ | 1.03 | $7.09 \times 10^{-2}$ |
| -1.36 | $2.01 \times 10^{-3}$ | -0.30 | $2.41 \times 10^{-1}$ | 1.05 | $2.56 \times 10^{-2}$ |
| -1.44 | $2.93 \times 10^{-3}$ | -0.02 | $9.94 \times 10^{-1}$ | 1.41 | $4.16 \times 10^{-3}$ |
| -1.69 | $3.47 \times 10^{-2}$ | 0.16 | $9.99 \times 10^{-1}$ | 1.85 | $6.82 \times 10^{-2}$ |
| -1.73 | $2.15 \times 10^{-4}$ | 0.04 | $9.65 \times 10^{-1}$ | 1.76 | $1.27 \times 10^{-4}$ |
| -1.91 | $3.48 \times 10^{-3}$ | -0.62 | $2.70 \times 10^{-1}$ | 1.28 | $8.04 \times 10^{-2}$ |
| -2.31 | $1.95 \times 10^{-5}$ | -0.01 | $1.00 \times 10^{-0}$ | 2.28 | $1.96 \times 10^{-5}$ |
| -2.38 | $8.93 \times 10^{-3}$ | 0.23 | $8.24 \times 10^{-1}$ | 2.59 | $3.42 \times 10^{-3}$ |
| -2.71 | $1.87 \times 10^{-2}$ | -1.09 | $9.99 \times 10^{-1}$ | 3.84 | $1.12 \times 10^{-12}$ |
| -4.73 | $2.23 \times 10^{-6}$ | -0.98 | $2.18 \times 10^{-1}$ | 3.74 | $4.22 \times 10^{-6}$ |
| -10.27 | $1.09 \times 10^{-6}$ | | | 5.26 | $2.98 \times 10^{-4}$ |
| Nucleotide metabolism genes |  |  |  |  |  |  |  |  |
| AT1G14250 | GDA1/CD39 | GDA1/CD39 nucleoside phosphatase family protein | 2.27 | $7.74 \times 10^{-4}$ | 1.31 | $1.13 \times 10^{-2}$ | -0.98 | $2.04 \times 10^{-3}$ |
| AT4G34900 | XDH2 | xanthine dehydrogenase 2 | 1.34 | $4.67 \times 10^{-4}$ | 0.06 | $9.12 \times 10^{-1}$ | -1.29 | $7.12 \times 10^{-4}$ |
| AT4G18440 | ADSL | Adenylosuccinate lyase | 1.13 | $4.18 \times 10^{-2}$ | 0.43 | $2.01 \times 10^{-1}$ | -0.72 | $1.18 \times 10^{-1}$ |
| AT3G62550 | | Adenine nucleotide alpha hydrolases-like superfamily protein | 0.97 | $1.73 \times 10^{-2}$ | 0.67 | $6.19 \times 10^{-3}$ | -0.31 | $3.82 \times 10^{-1}$ |
| AT1G28960 | NUDT15 | nudix hydrolase homolog 15 | 0.88 | $8.93 \times 10^{-3}$ | 0.73 | $8.80 \times 10^{-3}$ | -0.17 | $7.43 \times 10^{-1}$ |
| AT2G01670 | NUDT17 | nudix hydrolase homolog 17 | -0.79 | $2.45 \times 10^{-2}$ | -0.12 | $7.62 \times 10^{-1}$ | 0.66 | $6.13 \times 10^{-2}$ |
| AT2G39270 | | P-loop containing nucleoside triphosphate hydrolases superfamily protein | -1.24 | $2.12 \times 10^{-2}$ | 0.07 | $9.99 \times 10^{-1}$ | 1.34 | $4.68 \times 10^{-2}$ |
| AT5G63680 | MBK5.16 | Pyruvate kinase | -1.27 | $4.59 \times 10^{-2}$ | -0.11 | $9.99 \times 10^{-1}$ | 1.15 | $1.66 \times 10^{-1}$ |
| AT1G71100 | RPI1 | Probable ribose-5-phosphate isomerase 1 | -1.66 | $1.99 \times 10^{-3}$ | -0.21 | $7.85 \times 10^{-1}$ | 1.43 | $1.37 \times 10^{-2}$ |
| AT2G04430 | NUDT5 | nudix hydrolase homolog 5 | -1.96 | $3.24 \times 10^{-4}$ | -0.80 | $1.68 \times 10^{-1}$ | 1.15 | $5.84 \times 10^{-2}$ |
| AT4G12720 | NUDT7 | MutT/nudix family protein | -2.25 | $7.21 \times 10^{-5}$ | -0.05 | $9.64 \times 10^{-1}$ | 2.19 | $5.02 \times 10^{-5}$ |
| AT5G09290 | | Inositol monophosphatase family protein | -5.07 | $2.01 \times 10^{-5}$ | -2.63 | $6.09 \times 10^{-2}$ | 2.55 | $5.10 \times 10^{-3}$ |
| AT2G04450 | NUDT6 | nudix hydrolase homolog 6 | -5.82 | $4.07 \times 10^{-17}$ | -2.47 | $5.16 \times 10^{-13}$ | 3.36 | $5.33 \times 10^{-8}$ |
| Chlorophyll metabolism genes |  |  |  |  |  |  |  |  |
| AT4G27440 | PORB | Protochlorophyllide reductase B, chloroplastic | 0.88 | $3.13 \times 10^{-2}$ | 0.38 | $3.57 \times 10^{-1}$ | -0.51 | $4.32 \times 10^{-1}$ |
| AT3G15520 | CYP37 | Cyclophilin-like peptidyl-prolyl cis-trans isomerase family protein | 0.87 | $3.42 \times 10^{-2}$ | -0.23 | $3.53 \times 10^{-1}$ | -1.11 | $2.28 \times 10^{-3}$ |
| AT4G16690 | MES16 | pFDCC methylsterase MES16 | -0.90 | $3.50 \times 10^{-2}$ | -1.11 | $1.39 \times 10^{-4}$ | -0.22 | $6.58 \times 10^{-1}$ |
| AT1G20780 | PUB44 | U-box domain-containing protein 44 | -0.94 | $1.17 \times 10^{-2}$ | -0.39 | $2.37 \times 10^{-1}$ | 0.53 | $1.60 \times 10^{-1}$ |
| AT5G26030 | FC1 | Ferrochelatase-1, chloroplastic/mitochondrial | -1.20 | $2.12 \times 10^{-3}$ | 0.18 | $7.36 \times 10^{-1}$ | 1.36 | $5.48 \times 10^{-4}$ |
| AT3G03470 | CYP89A9 | Cytochrome P450 89A9 | -1.57 | $4.26 \times 10^{-2}$ | 0.05 | $9.99 \times 10^{-1}$ | 1.60 | $8.75 \times 10^{-2}$ |
| AT1G69720 | HO3 | Heme oxygenase 3, chloroplastic | -1.69 | $6.79 \times 10^{-4}$ | 0.08 | $9.71 \times 10^{-1}$ | 1.76 | $1.60 \times 10^{-4}$ |
| At1g22400 | ATUGT85A1 | UDP-Glycosyltransferase superfamily protein | -1.94 | $4.45 \times 10^{-4}$ | -0.22 | $7.41 \times 10^{-1}$ | 1.71 | $9.31 \times 10^{-4}$ |
| AT4G11910 | SGR2 | Magnesium dechelataase SGR2, chloroplastic | -2.04 | $4.88 \times 10^{-2}$ | | | 3.05 | $6.63 \times 10^{-3}$ |

Nitrogen and chlorophyll metabolism genes differentially expressed in *Arabidopsis thaliana* leaves 20 days after two foliar sprays with 0.04% IFFCO liquid nano urea formulation (NUF) versus equimolar bulk urea are shown. Mock treatment refers to a spray with distilled water. Comparisons between NUF and urea, NUF and mock, and urea and mock treatments are shown with $\log_2$ (fold-change) in expression levels and Benjamini-Hochberg false discovery rate (FDR)-adjusted *P*-values. $\log_2FC > 0.5$ or $< -0.5$ are highlighted in shades of red and blue, respectively. The *FDR* values $< 0.05$ are indicated in red font. The genes were categorized into functional groups by the BAR Classification SuperViewer tool using the Gene Ontology and/or Mapman databases.

**Table 2.**
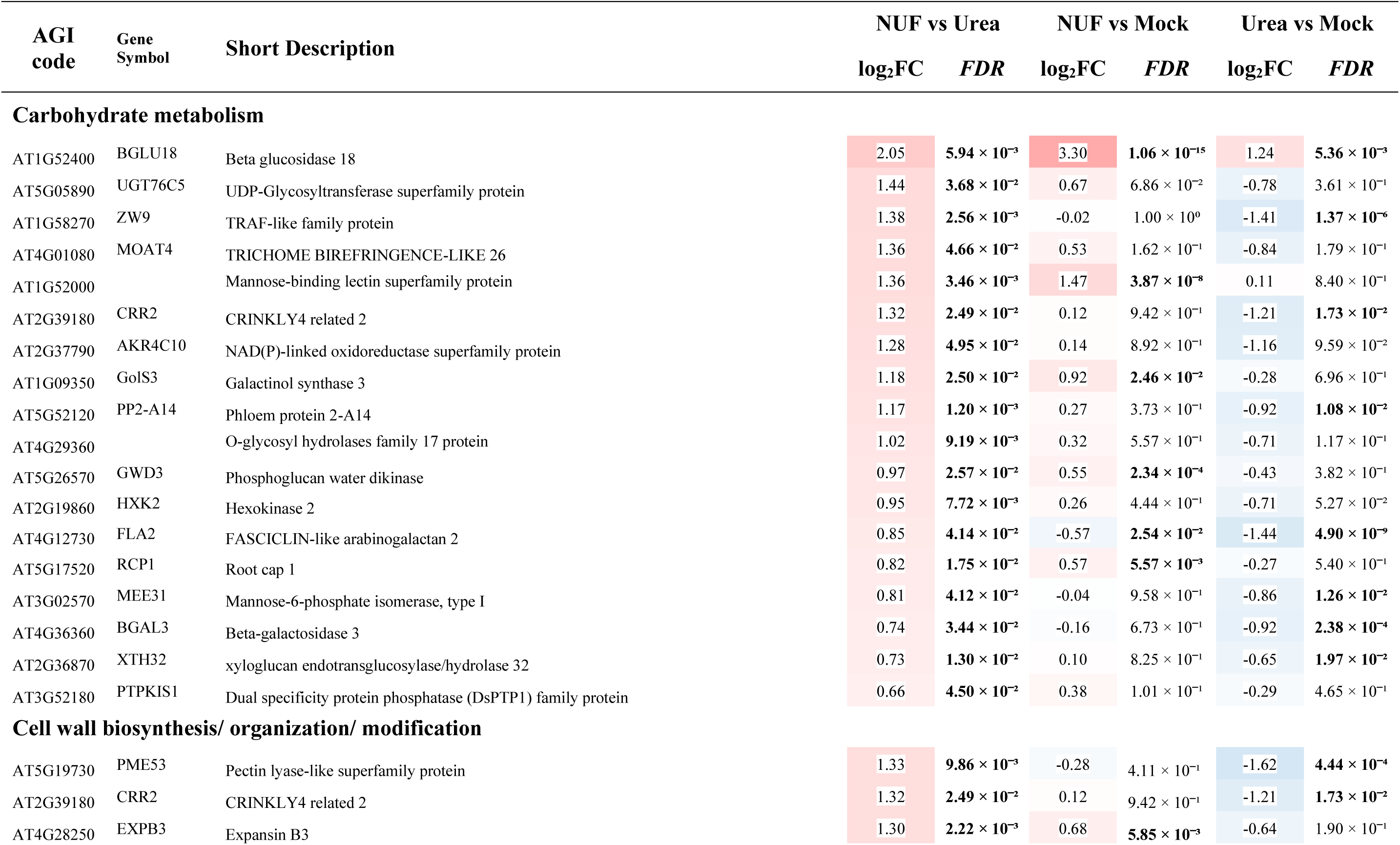
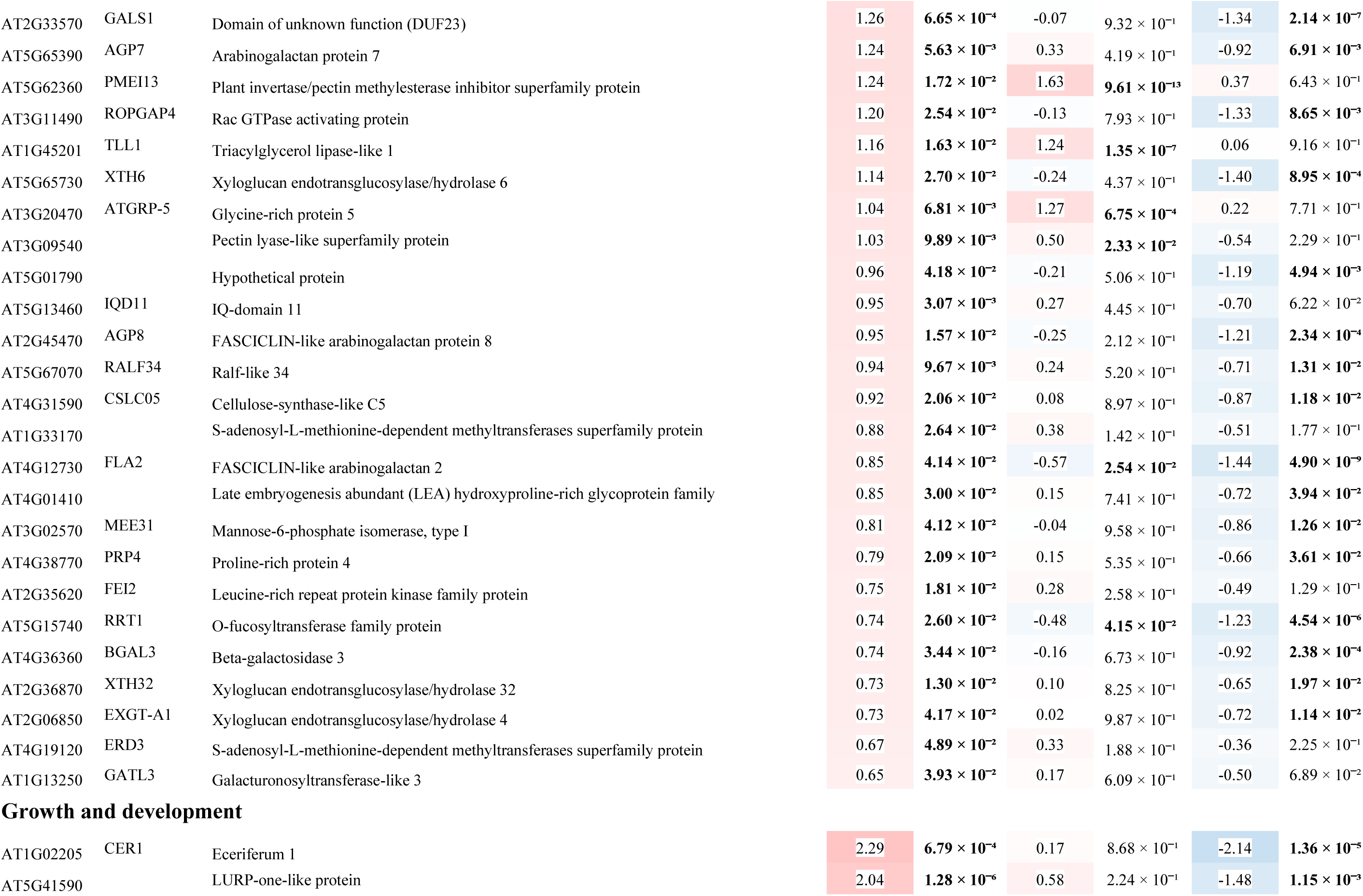
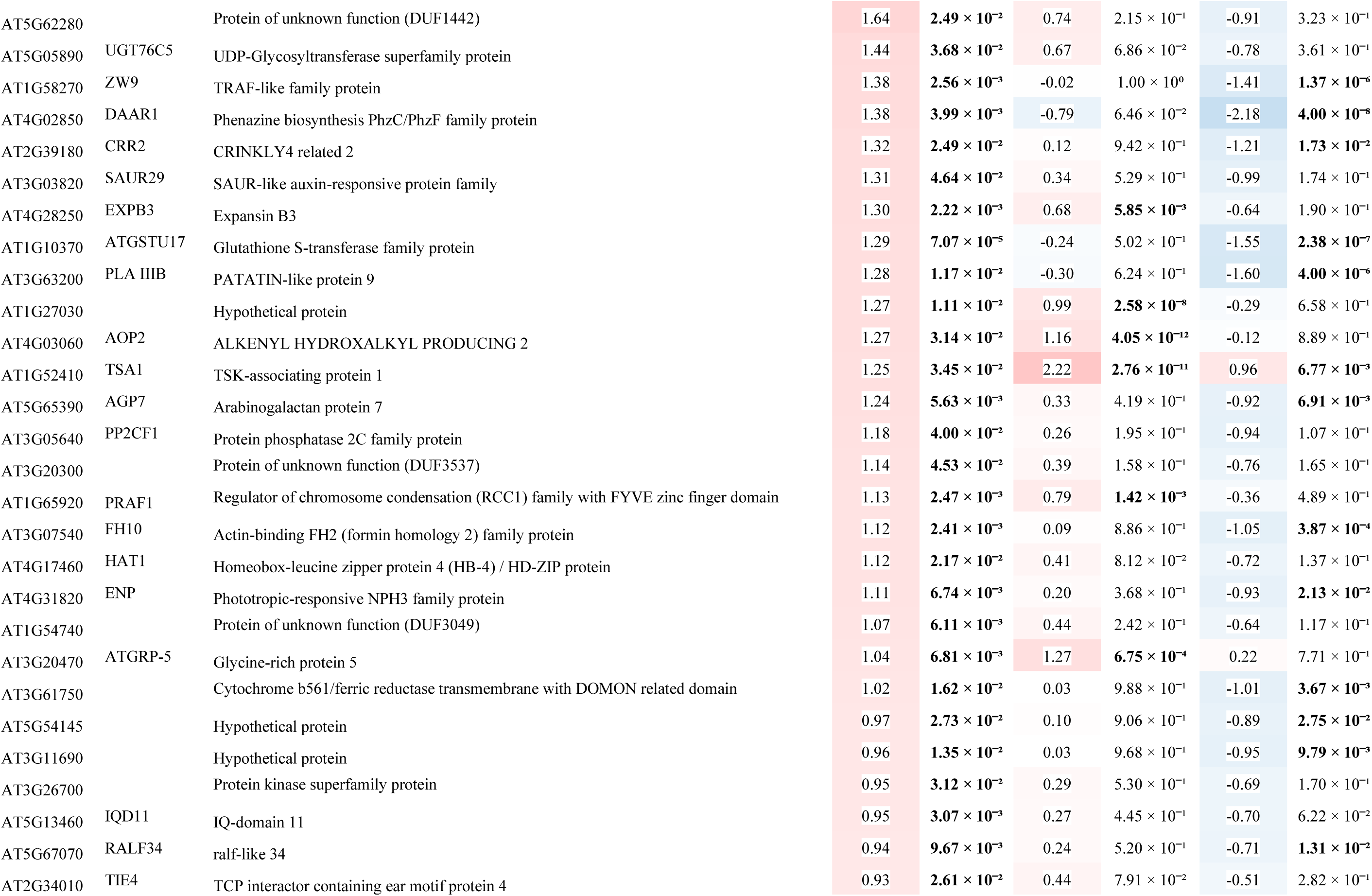
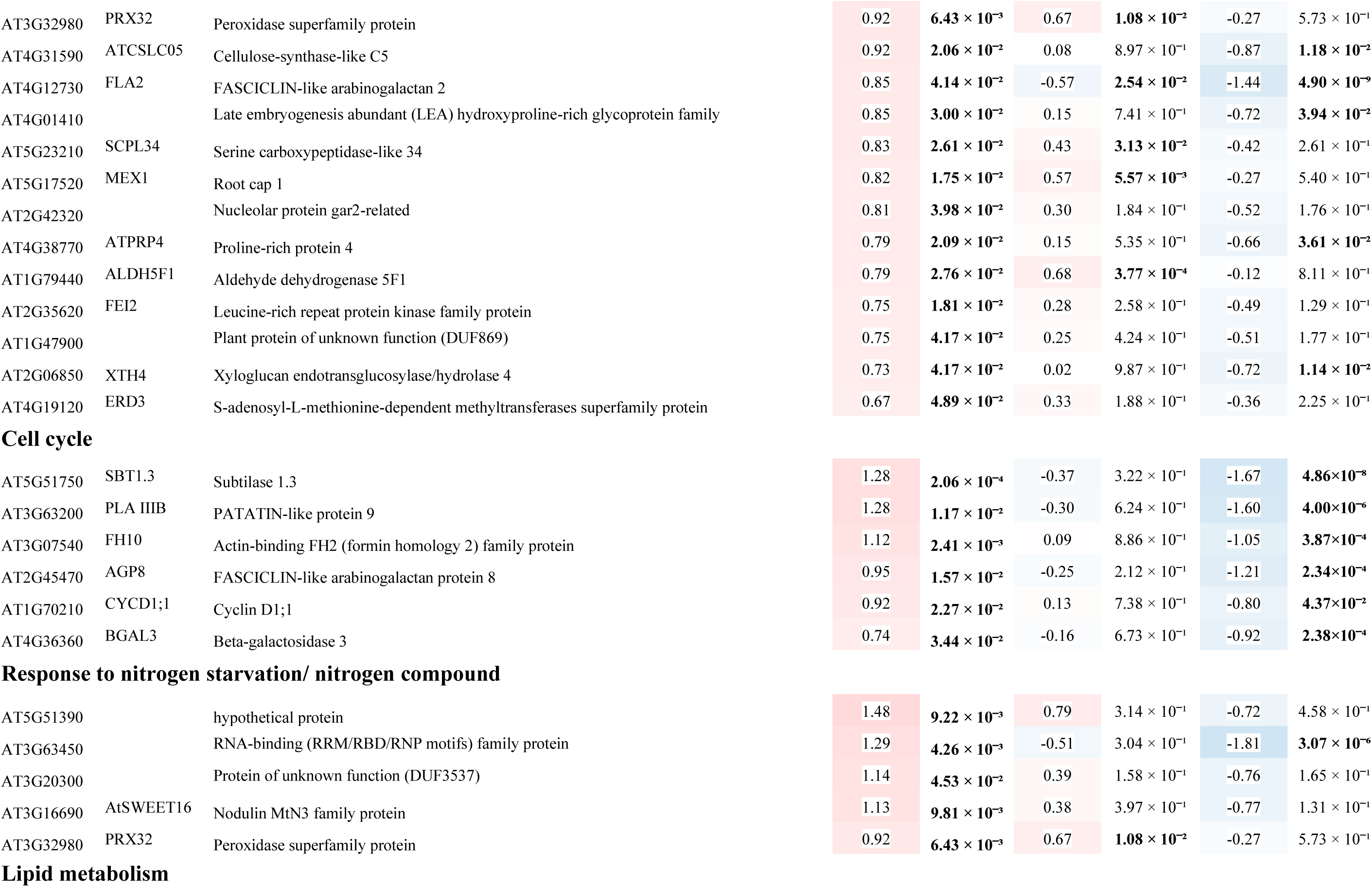
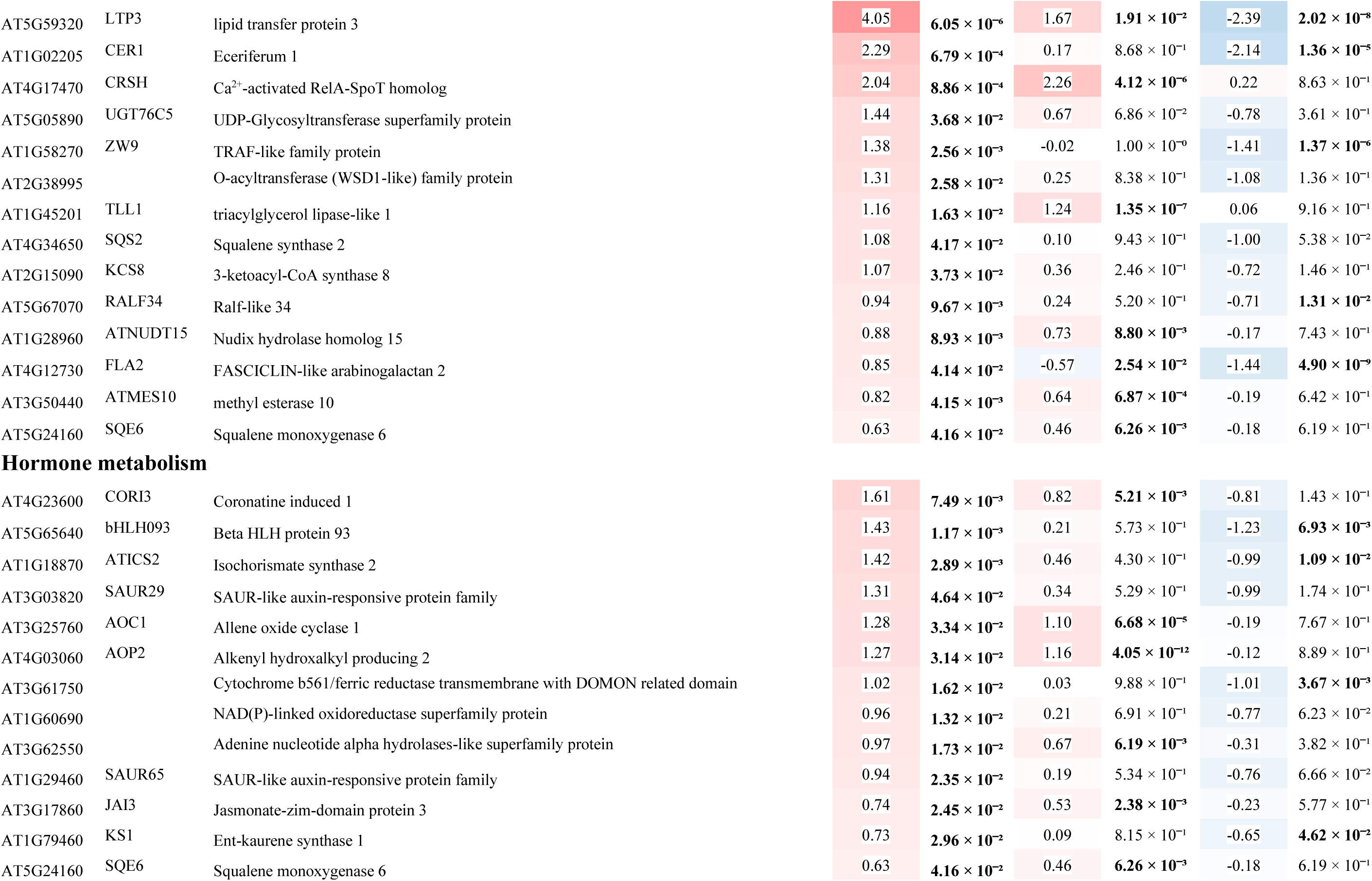
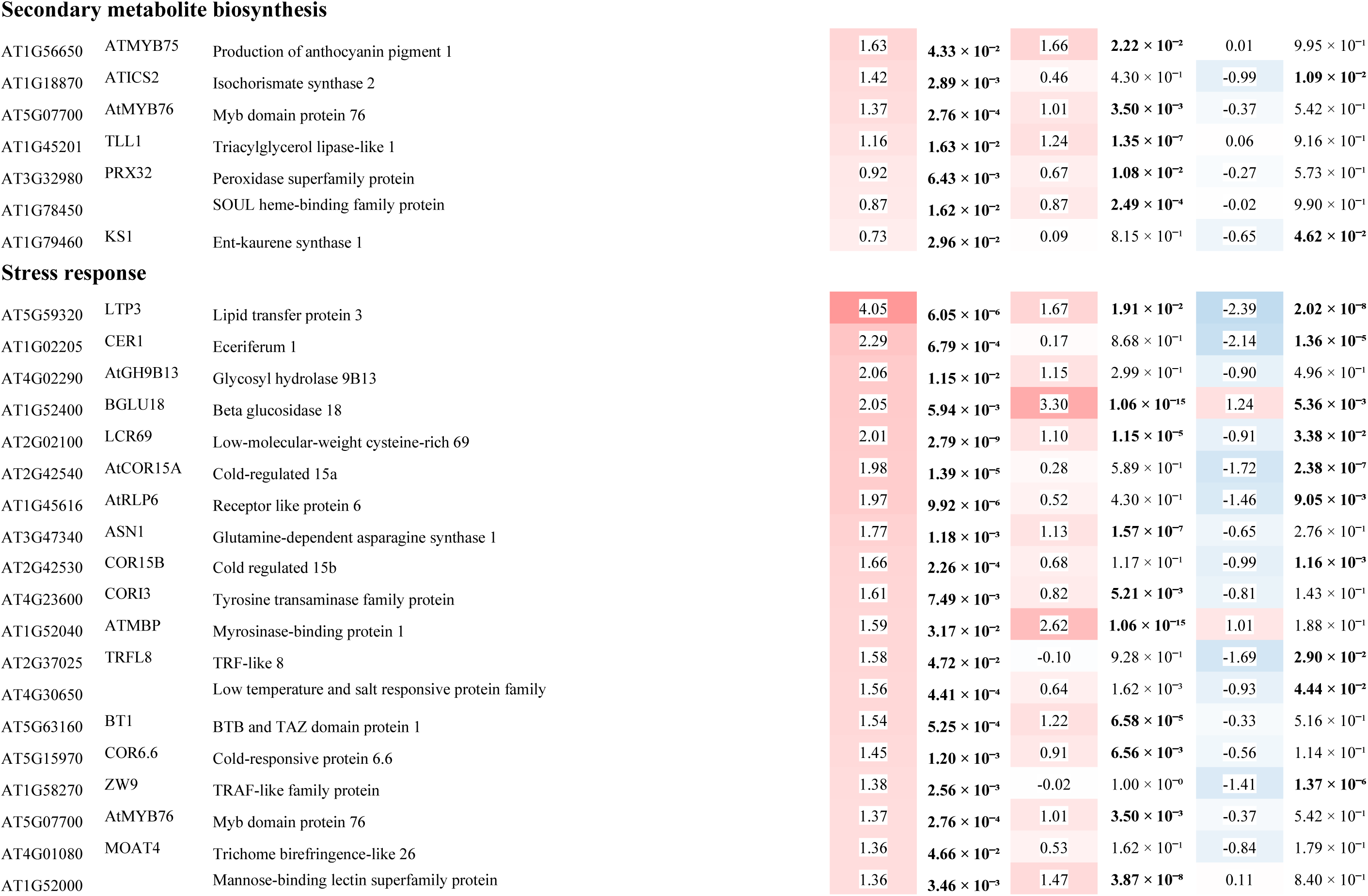
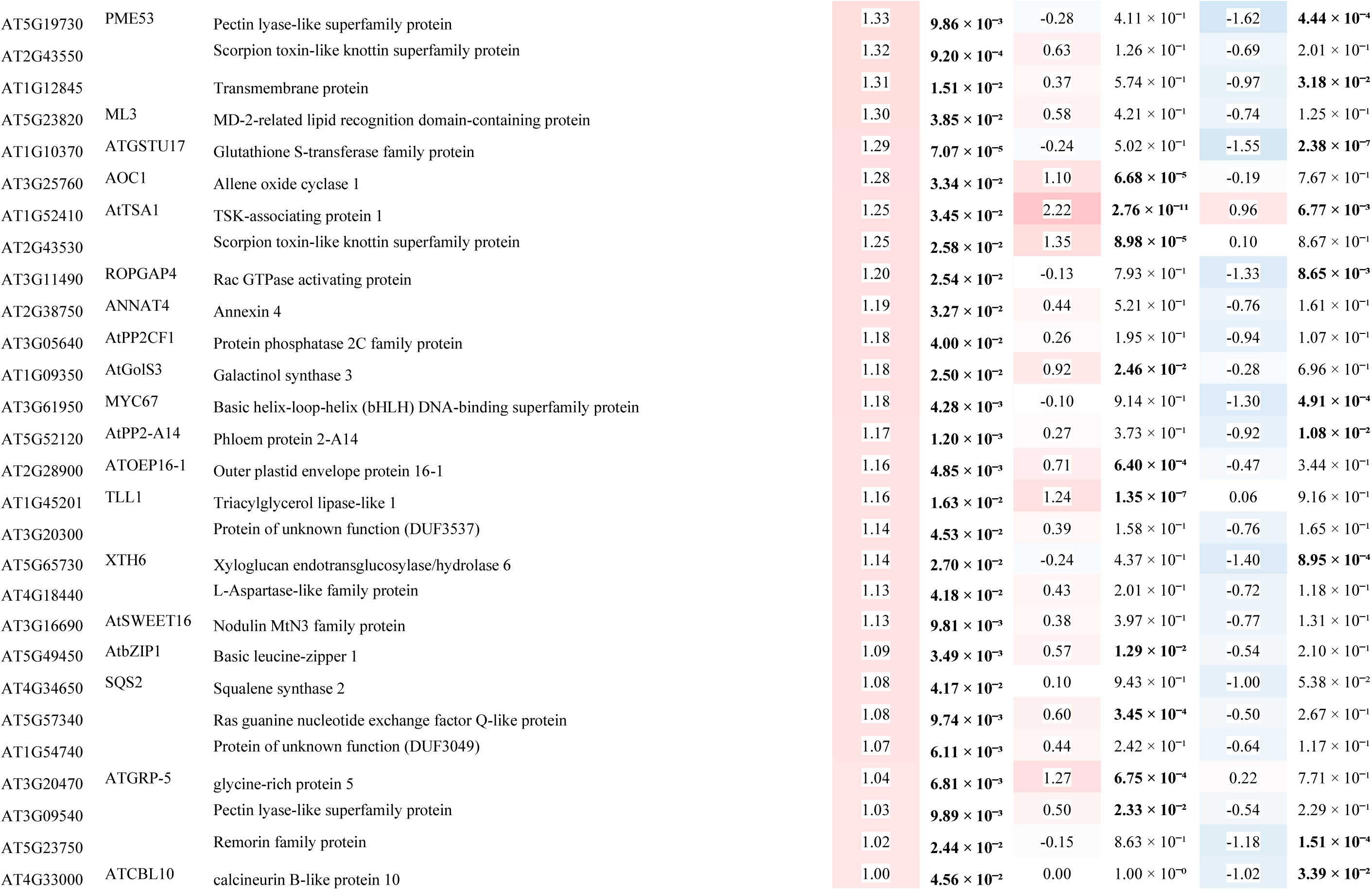
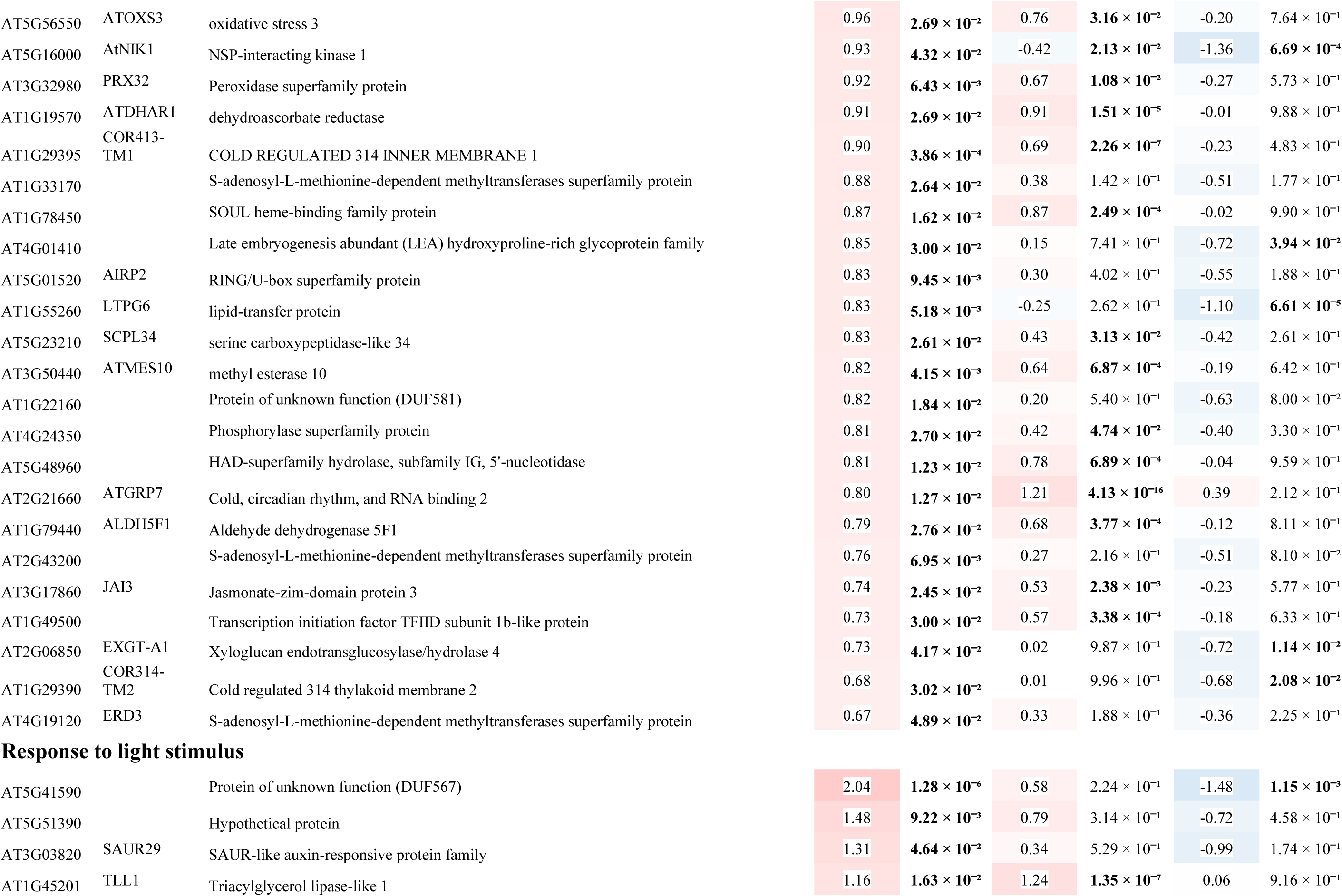
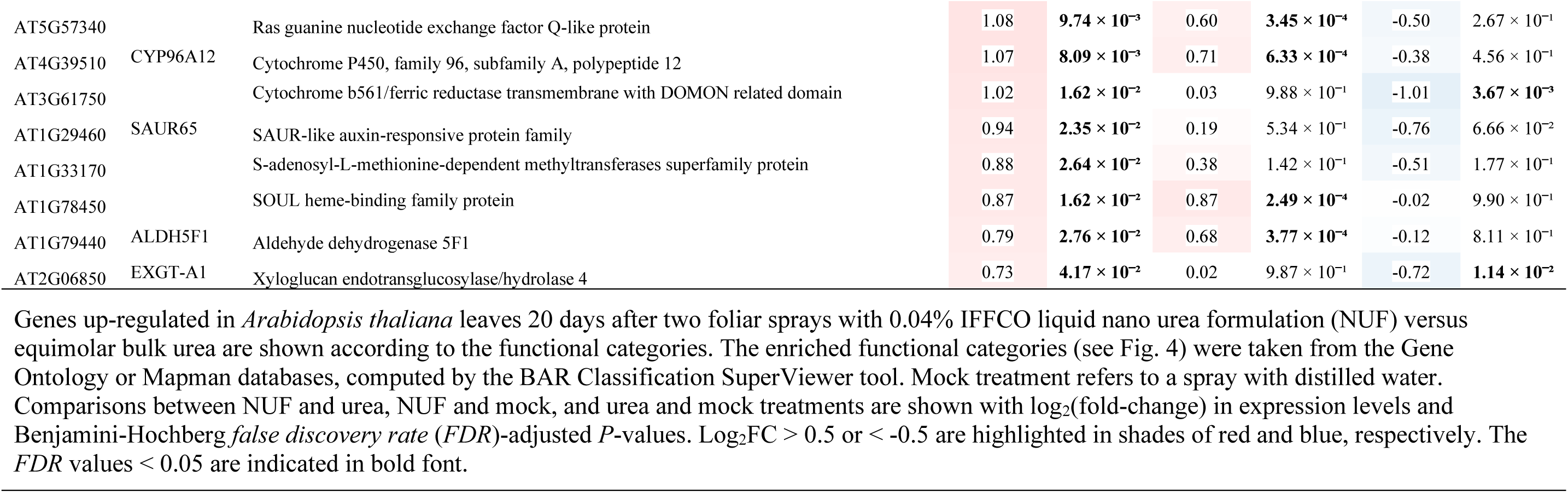
Growth-related *Arabidopsis thaliana* genes up-regulated under IFFCO nano urea versus equimolar bulk urea.

### Inorganic nitrogen, amino acid, and nucleotide transporter genes

To gain insights into the differences in the uptake modes of NUF and urea, at first we examined the expression of nitrogen transporters. Upon NUF treatment almost all the genes encoding nitrogen, amino acid, and nucleotide transporters were significantly down-regulated compared to urea treatment (Table 1, Fig. 6). Among the 22 nitrogen transporters perturbed upon NUF treatment, four inorganic nitrogen transporter genes, *degradation of urea 3* (*DUR3*; AT5G45380), *ammonium transporter 2* (*AMT2*; AT2G38290), *C-terminally encoded peptide 14* (*CEP14*; AT1G29290) and *high-affinity nitrate transporter 3.1* (*NRT3.1*; AT5G50200) were significantly down-regulated. All the amino acid transporter genes were down-regulated except *outer envelope pore protein 16-1* (*OEP161*; AT2G28900), which was up-regulated under NUF treatment. The down-regulated amino acid transporters included *transmembrane amino acid transporter family protein* (AT2G41190), *lysine histidine transporter 1* (*LHT1*; AT5G40780), *glutamate receptor 2.7* (*GLR2.7*; AT2G29120), *glutamate receptor 1.4* (*GLR1.4*; AT3G07520), *E3 ubiquitin-protein ligase* (*ATL31*; AT5G27420), *amino acid transporter 1* (*AAT1*; AT4G21120), *Glutamate receptor* (*GLR2.8*; AT2G29110), *Glutamate receptor 1.2* (*GLR1.2*; AT5G48400) and *LYS/HIS transporter 7* (*LHT7*; AT4G35180). Two genes encoding ATP carriers were significantly down-regulated which are *ADP, ATP carrier protein 2* (*AAC2*; AT5G13490) and *ADP, ATP carrier protein 3* (*AAC3*; AT4G28390). Other nucleotide transporters, which are involved in the transport of cyclic nucleotides, *cyclic nucleotide gated channel 3* (*CNGC3*; AT2G46430), *cyclic nucleotide gated channel 10* (*ACBK1*; AT1G01340) and *cyclic nucleotide-gated channel 13* (*CNGC13*; AT4G01010) also followed the trend of down-regulation under NUF treatment, compared to urea.

**Fig. 6.**
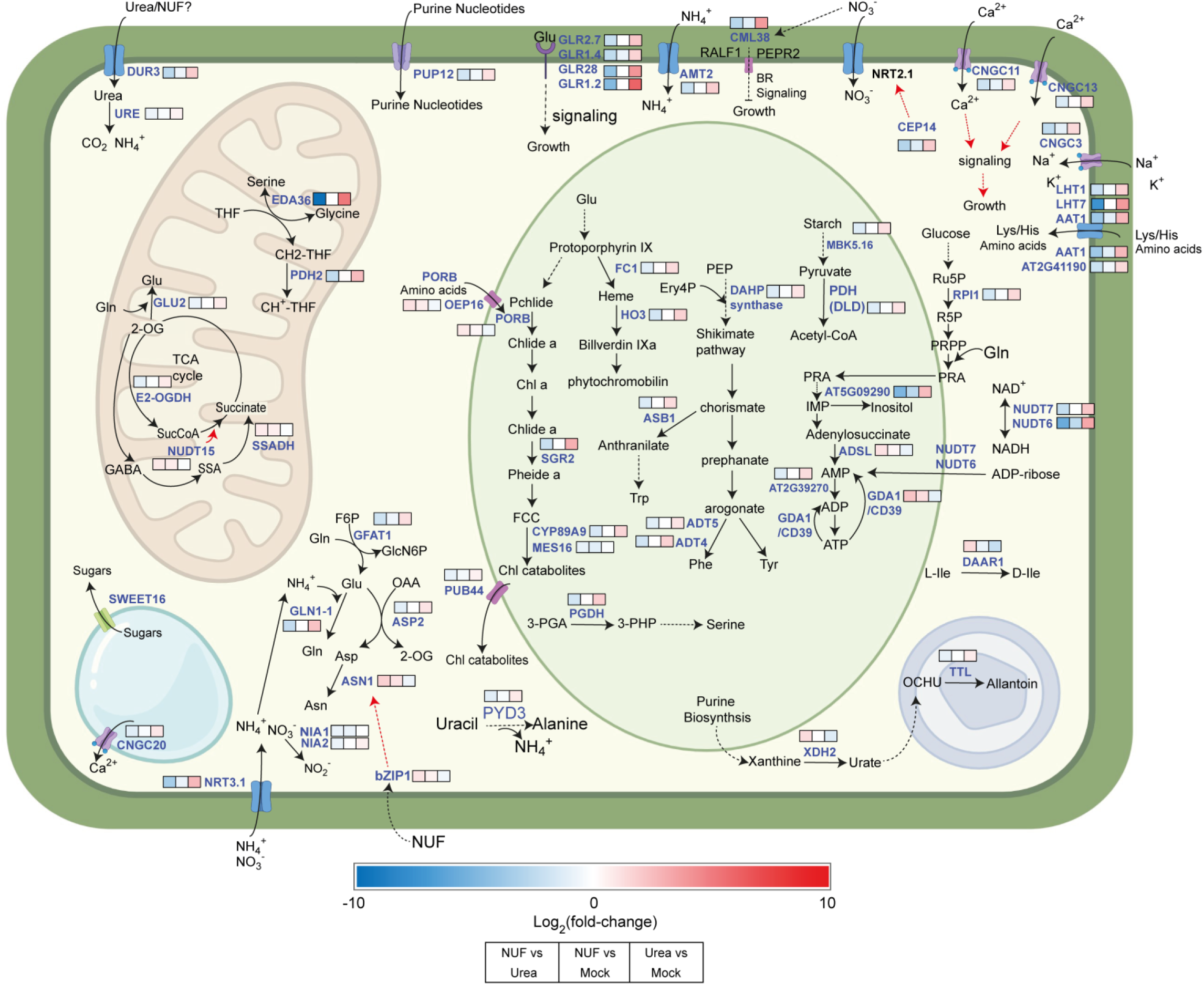
Nitrogen and chlorophyll metabolism of *Arabidopsis thaliana* perturbed by foliar administration of IFFCO nano urea and equimolar bulk urea. Biochemical conversions or transport processes are shown by black arrows and metabolites are shown in black font. Genes are indicated in blue. The three colored boxes against each gene indicate three comparisons in gene expression levels in the transcriptome: IFFCO nano urea formulation (NUF) vs urea, NUF vs mock treatment, and urea vs mock treatments (Table 1-2). In the color scale, blue indicates down-regulation (log_2_fold-change < -0.5, *P* < 0.05) and red up-regulation (log_2_FC > 0.5, *P* < 0.05). Red arrows indicate genetic regulation. Bold arrows indicate direct conversion and dotted arrows those with intermediate steps. Urea, ammonia, nitrate, and nitrite are transported into the cytosol by DUR3, AMT2, NRT2.1, and NRT3.1 transporters, where they participate in amino acid biosynthesis and key pathways like the TCA cycle in mitochondria and chlorophyll biosynthesis in chloroplasts. Glutamate (Glu) is transformed into glutamine (Gln) by GLN1-1 and into aspartate (Asp) and 2-oxoglutarate (2-OG) via ASP2. Asp is further modified into asparagine (Asn) by ASN1, regulated by the bZIP transcription factor, which shows higher expression under nano urea formulation (NUF) than urea. Gln, together with fructose-6-phosphate (F6P), forms glucosamine-6-phosphate and Glu, catalyzed by GFAT1. In the TCA cycle, 2-OG is metabolized into succinyl-CoA (SucCoA) by OGDH and converted into succinate by succinyl-CoA synthetase, regulated by NUDT15. Through the GABA shunt, SSADH facilitates the transformation of 2-OG into succinate, linking nitrogen metabolism to TCA and generating Glu via GLU2. CML38 is a secretory protein that is activated under low nitrogen conditions and, in turn, activates brassinosteroid (BR) signaling through RALF1 and PEPR2, leading to growth inhibition. However, under nitrogen excess conditions after NUF spray, *CML38* is downregulated, reducing its impact on growth inhibition. Different types of amino acids like serine, phenylalanine, tyrosine (Tyr), tryptophan (Trp), serine (Ser), alanine (Ala), and glycine (Gly) are formed by different pathways that involve different enzymes ADT5, ADT4, ASB1, PGDH, PYD3, and EDA36. Transporters like LHT1, LHT7, AAT1, and AT2G41190 transport amino acids inside and outside the cell. *De novo* biosynthesis of purines begins with ribulose-5-phosphate (Ru5P), which is converted into ribose-5-phosphate (R5P). This intermediate is further processed to form purine nucleotides. The pathway is regulated by several key enzymes, including GDA1/CD39, NUDT7, and NUDT6. Inosine monophosphate (IMP) and adenosine triphosphate (ATP) are synthesized from phosphoribosylamine (PRA) and adenylosuccinate through the action of AT5G09290 and ADSL, respectively. Purine nucleotides are degraded to form xanthine, which is then converted into urate by the enzyme XDH2. Further degradation of urate produces oxonic acid (OCHU), which is subsequently converted into allantoin by TTL. Calcium signaling pathways involve channels such as CNGC11, CNGC13, and CNGC20, which regulate calcium (Ca²⁺) influx and are crucial for plant growth and nitrogen acquisition. In the chloroplast, chlorophyll production begins with glutamate (Glu) and goes through numerous stages to protoporphyrin IX. Protoporphyrin IX diverges into two primary pathways: one for chlorophyll biosynthesis and one for phytochromobilin formation. PORB, a major enzyme in the chlorophyll biosynthesis pathway, catalyzes the conversion of protochlorophyllide (Pchlide) into chlorophyllide a (Chlide a), which is then transformed into chlorophyll. The alternate procedure employs enzymes such as FC1 and HO3 to produce heme and biliverdin IX, which are then converted to phytochromobilin. OPE16 transporter found on chloroplast membrane is responsible for transporting amino acids and PORB enzyme, which is important for chlorophyll synthesis. Under NUF treatment, certain enzymes involved in chlorophyll catabolism, specifically SGR2, CYP89A9, and MES16, which degrade chlorophyll a to chlorophyll catabolites, are not upregulated. These chlorophyll catabolites are then exported by PUB44.

### Amino acid and nucleotide metabolism genes

Nitrogen supplied in urea and nano urea is likely to be assimilated into nitrogenous organic compounds, amino acids and nucleotides, essential for plant growth. Hence, we examined the genes for nitrogen assimilation, amino acid, and nucleotide metabolism, which exhibited different expression patterns upon NUF treatment than urea (Table 1, Fig. 6). Out of them, five amino acid metabolism genes showed considerable upregulation under NUF treatment. These were *glutamine-dependent asparagine synthase 1* (*ASN1*; AT3G47340), a key enzyme in asparagine synthesis, *tyrosine transaminase family protein* (*CORI3*; AT4G23600), *D-amino acid racemase1* (*DAAR1*; AT4G02850), *basic leucine zipper 1* (*BZIP1*; AT5G49450) and *succinate-semialdehyde dehydrogenase* (*ALDH5F1*; AT1G79440), which are involved in regulation of cellular amino acid metabolic process. On the other hand, 26 genes in this category were down-regulated. These included a key enzyme in the urea catabolic process, urease (*URE*; AT1G67550), two important genes of the nitrate assimilation process *nitrate reductase 1* (*NIA1*; AT1G77760) and *nitrate reductase 2* (*NIA2*; AT1G37130), *serine hydroxymethyltransferase* (*EDA36*; AT4G13890), *glutamine synthetase cytosolic isozyme 1-1* (*GLN1-1*; AT5G37600), *tyrosine aminotransferase 3* (*TAT3*; AT2G24850) and *glutamate synthase 2* (*GLU2*; AT2G41220), which are involved in amino acid metabolism and synthesis.

Thirteen nucleotide metabolism genes showed altered expression patterns under NUF treatment. Genes induced by NUF but suppressed by urea included *adenylosuccinate lyase* (*ASL*; AT4G18440), catalyzing two important steps in purine biosynthesis, *adenine nucleotide alpha hydrolases-like superfamily protein* (AT3G62550) *nudix hydrolase homolog 15* (*NUDT15*; AT1G28960), and *xanthine dehydrogenase 2* (*XDH2*; AT4G34900). All other nucleotide metabolism genes including *inositol monophosphatase* (*IMPase*; AT5G09290), *NUDT5* (AT2G04430), *NUDT6* (AT2G04450), *ATNUDT17* (AT2G01670), *ribose-5-phosphate isomerase 1* (*RPI1*; AT1G71100) and a very important enzyme of purine metabolism *pyruvate kinase* (*MBK5.16*; AT5G63680) were considerably down-regulated under NUF but induced under urea treatment.

### Chlorophyll metabolism genes are differentially regulated by NUF and urea

Two chlorophyll biosynthesis genes including *protochlorophyllide oxidoreductase B* (*PORB*; AT4G27440) and the *Cyclophilin-like peptidyl-prolyl cis-trans isomerase family protein* (*CYP37*; AT3G15520) serving as a crucial modulator of the electron transport chain were up-regulated under NUF. Furthermore, the *outer plastid envelope protein 16-1* (*ATOEP16-1*; AT2G28900), involved in the import of protochlorophyllide oxidoreductase A, was induced by NUF (Table 2; stress-responsive genes). Furthermore, chlorophyll catabolism genes such as *pFDCC methylesterase* (*MES16*; AT4G16690) and *cytochrome P450 89A9* (*CYP89A9*; AT3G03470) were down-regulated under NUF but up-regulated by urea (Table 1).

### NUF induces higher levels of growth and development related genes than urea

Comparing the expression levels of growth-related genes after NUF and urea foliar spray treatments gave significant insights into the higher growth obtained by NUF than urea in *A. thaliana*. Genes up-regulated by NUF compared to urea (log_2_FC > 0.5; *FDR* < 0.05) belonged to the enriched categories of cell wall biosynthesis/organization/modification, carbohydrate, nucleotide, lipid, and hormone metabolism, the biological processes having well-known roles in plant growth. In addition, many of these genes were also categorized as stress-responsive genes, promoting plant growth under biotic and abiotic stresses (Table 1, 2). We validated the expression levels of randomly selected growth-promoting genes under NUF and urea treatments by qPCR (Fig. 5; Table S2).

### Cell wall biosynthesis and modification genes

We observed 28 cell wall biosynthesis and modification genes significantly induced by NUF than urea, which were involved in growth regulation processes. These included a *plant invertase/pectin methylesterase inhibitor superfamily protein* (*PMEI13*; AT5G62360), *triacylglycerol lipase-like 1* (*TLL1*; AT1G45201), *glycine-rich protein 5* (*ATGRP-5*; AT3G20470), *pectin lyase-like superfamily protein* (*ATPLL1*; AT3G09540), a kinase *CRINKLY4 related 2* (*CRR2*; AT2G39180), *arabinogalactan protein 7* (*AGP7*; AT5G65390), and a cell wall loosening and cell elongation-related *expansin B3* (*EXPB3*; AT4G28250).

### Carbohydrate and lipid metabolism genes

Eighteen carbohydrate metabolism genes exhibited higher levels of upregulation under NUF than urea treatments. Notably, a gene responsible for abscisic acid production from its glucose ester in endoplasmic reticulum bodies, *beta-glucosidase 18* (*BGLU18*; AT1G52400), which also promotes shoot development, showed a significant increase in expression by NUF. Essential carbohydrate metabolism and abiotic stress tolerance genes, *mannose-binding lectin superfamily protein* (AT1G52000), *galactinol synthase 3* (*GolS3*; AT1G09350), and *O-glycosyl hydrolases family 17 protein* (AT4G29360), were also induced higher by NUF than urea.

Fifteen genes linked to lipid metabolism showed elevated levels of expression under NUF compared to urea. Particularly, *lipid transfer protein 3* (*LTP3*; AT5G59320), critically involved in plant growth and development, was more strongly induced by NUF than urea or mock treatments (Table 2, Fig. 5). A fatty acid biosynthesis gene altering ER lipid properties under stress, *Ca^2+^-activated RelA-SpoT homolog* (*CRSH*; AT4G17470), an alkane and epicuticular wax biosynthesis gene *eceriferum 1* (*CER1*; AT1G02205), and a very long chain fatty acid biosynthesis gene, *3-ketoacyl-CoA synthase 8* (*KCS8*; AT2G15090) were considerably induced by NUF than urea. Two carbohydrate and lipid metabolism genes *UDP-Glycosyltransferase superfamily protein* (*UGT76C5*; AT5G05890) and *TRAF-like family protein* (*ZW9*; AT1G58270) were also categorized under “growth and development”.

### Cell cycle genes

Seven genes involved in the cell cycle having potential roles in plant growth showed higher expression levels under NUF than urea. Compared to the mock treatment, these genes were more down-regulated by urea, whereas NUF caused lesser suppression of these genes (Table 2). This included *subtilase 1.3* (*SBT1.3*; AT5G51750), a serine protease implicated in immune responses and programmed cell death, *Actin-binding FH2 (formin homology 2) family protein* (*FH10*; AT3G07540) which facilitates cytokinesis and cell morphogenesis, promoting growth, *FASCICLIN-like arabinogalactan protein 8* (*AGP8*; AT2G45470) which plays an assortment of roles in plant development, and *Cyclin D1;1* (*CYCD1;1*; AT1G70210), a major cell cycle regulating gene that maintains proper plant growth. Another gene in this category, *leucine-rich repeat protein kinase family protein* (*FEI2*; AT2G35620) regulates cell wall elongation.

### Hormone metabolism and signaling genes

Phytohormones like auxin, brassinosteroid (BR), gibberellic acid (GA), jasmonate (JA), and salicylate (SA) are known plant growth regulators. Thirteen genes belonging to the enriched functional category “hormone metabolism”, involved in various crucial steps of phytohormone biosynthesis and signaling, were significantly induced by NUF while being down-regulated by urea, compared to the mock treatment (Fig. 4, Table 2). The JA biosynthesis and signaling genes included *tyrosine transaminase family protei*n (*CORI3*; AT4G23600), *allene oxide cyclase 1* (*AOC1*; AT3G25760), *jasmonate-zim-domain protein 3* (*JAI3*; AT3G17860), and *alkenyl hydroxalkyl producing 2* (*AOP2*; AT4G03060). Furthermore, a jasmonic acid-induced *Tonsoku* (*TSK)-associated protein 1* (*AtTSA1*; AT1G52410) leads to the formation of endoplasmic reticulum (ER) body formation as a stress response. Similarly, a higher induction of the GA metabolism-regulating genes *Beta HLH protein 93* (*bHLH093*; AT5G65640) and the *Ent-kaurene synthase 1* (*KS1*; AT1G79460), the BR biosynthesis gene *squalene monoxygenase 6* (*SQE6*; AT5G24160), and the SA biosynthesis gene, *isochorismate synthase 2* (*ICS2*; AT1G18870) under NUF may explain the higher growth observed under NUF than urea. The auxin signaling genes included two paralogs of the *SAUR-like auxin-responsive protein family* (*SAUR29*; AT3G03820 and *SAUR65*; AT1G29460). *AOP2* and *SAUR29* were also categorized under “growth and development”. Moreover, NUF-induced *protein phosphatase 2C family protein* (*PP2CF1*; AT3G05640) interacts within the abscisic acid (ABA) signaling pathway and mitigates osmotic stress.

### Light-responsive genes

The *SAUR* genes induced at higher levels by NUF additionally fall under the enriched category “response to light stimulus” (Table 2). Other light-response genes implicated in plant growth and stress response were induced at higher levels by NUF. These included *triacylglycerol lipase-like 1* (*TLL1*; AT1G45201) catalyzing triacylglycerol breakdown, c*ytochrome b561/ferric reductase transmembrane with DOMON related domain* (AT3G61750), *cytochrome P450, family 96, subfamily A, polypeptide 12* (*CYP96A12*; AT4G39510), *SOUL heme-binding family protein* (AT1G78450), and *aldehyde dehydrogenase 5F1* (*ALDH5F1*; AT1G79440). Other photomorphogenesis genes included an auxin-transporting *phototropic-responsive NPH3 family protein* (*ENP*; AT4G31820).

### Other stress-responsive growth-promoting genes

NUF induced 70 genes within the biological process category “stress response” compared to urea. Most of these genes were up-regulated by NUF while down-regulated under urea compared to the mock treatment. These included a seedling growth-promoting *myrosinase-binding protein 1* (*ATMBP*; AT1G52040), *glycine-rich protein 5* (*GRP-5*; AT3G20470) which regulate cell and organ growth as well as stress adaptation, *calcineurin B-like protein 10* (*CBL1*0; AT4G33000), a vital ion homeostasis protein that boosts the shoot growth, the transcription factor *basic leucine-zipper 1* (*bZIP1*; AT5G49450), having roles in multiple aspects of plant development and stress tolerance. Antioxidant-encoding stress-responsive genes included *glutathione S-transferase family protein* (*ATGSTU17*; AT1G10370), *peroxidase superfamily protein* (*PRX32*; AT3G32980), *galactinol synthase 3* (*AtGolS3*; AT1G09350), which synthesizes sugars to protect against abiotic and oxidative stresses. The pathogen defense and plant development-regulating genes included *LURP-one-like protein* (*AT5G41590*), *NSP-interacting kinase 1* (*NIK1*; AT5G16000), *BTB and TAZ domain protein 1* (*BT1*; AT5G63160) were found to be up-regulated under NUF treatment. Moreover, *annexin 4* (*ANNAT4*; AT2G38750) promotes plant growth by mitigating multiple environmental stresses.

### NUF suppresses genes for cell death, abscission, and senescence in contrast to urea

We analyzed the genes suppressed by NUF but up-regulated by urea to explore their possible roles in the higher growth observed under NUF compared to urea. These genes mostly belonged to the functional categories of cell death, abscission, senescence, cellular catabolism, biotic stress response, and signal transduction (Table S3). Among them, two genes for cell death, an alpha/beta-hydrolases superfamily protein *MAGL13* (AT5G11650) and a Fe(II)- and 2-oxoglutarate-dependent dioxygenase family gene *F6’H1* (AT3G13610) were suppressed by NUF compared to urea (Table S3). Moreover, 56 genes for senescence and 6 genes regulating abscission were down-regulated by NUF. These included abscission regulating *receptor-like protein kinase 5* (*RLK5*: AT4G28490) and *mitogen-activated protein kinase kinase* 4 (*MKK4*; AT1G51660), *protein kinase superfamily protein* (*CST;* AT4G35600), leaf senescence-regulating *NAC domain containing protein 42* (*NAC042*; AT2G43000), *senescence-associated gene 101* (*SAG101*; AT5G14930), *NDR1/HIN1-like protein 10* (*NHL10*; AT2G35980), *WRKY transcription factor 53* (*WRKY53*; AT4G23810), and *DMR6-LIKE OXYGENASE 1* (DLO1; AT4G10500). Moreover, a growth restricting regulatory protein *DA1* (AT1G19270) was also down-regulated.

## Discussion

Our physical measurements using DLS and TEM have corroborated that NUF particle size ranges between 20 to 35 nm, accompanied by a -54.74 mV charge, indicating that NUF is suitable for entering the stomatal apertures in the μm range (Scarpeci, Zanor, and Valle 2017). Reportedly, nanoparticles with sizes from 3 to 95 nm, and possessing negative charges within the range of -31.1 to -69.2 mV hold the capacity to disruptively penetrate the leaf cuticle partly and mainly enter through the stomata, leading to an effective translocation to other plant parts (Avellan et al. 2021; Lian et al. 2021). The stomatal route is more plausible for water-suspended nanoparticles with a size above 10 nm, as cuticular channels are smaller, 2-4.8 nm (Eichert et al. 2008).

We cultivated *A. thaliana* in a soil-less vermiculite medium and applied NUF as foliar sprays to compare its effect on plant growth with equimolar urea and mock sprays with deionized water. In addition, the vermiculite received background fertilization with a dilute ¼-Hoagland’s solution containing low concentrations of nitrogen and other essential macro and micronutrients (Jangir et al. 2024). Under these conditions, foliar spray administration of urea and NUF led to increased growth than the mock spray. Moreover, NUF outperformed urea in enhancing leaf biomass, chlorophyll, and nitrogen content, indicating an advantage of NUF over equimolar bulk urea (Fig. 1). We conducted a comparative transcriptome analysis of the leaves to gain insights into the differential regulation of genes leading to a higher growth promotion by NUF compared to urea.

Almost all the nitrogen transporter genes were up-regulated under urea while down-regulated under NUF. *DUR3*, *AMT2* and *NRT3.1* are high-affinity nitrogen transporters and play a critical role under low nitrogen conditions (Buoso et. al, 2021; Zhang et. al, 2022). Down-regulation of these transporters under NUF treatment suggests ample supply of their substrates, urea, ammonium, and nitrate respectively, while induction of these genes under urea suggests nitrogen insufficiency. This is also supported by our observed lower nitrogen levels under urea treatment than NUF (Fig. 1D). *DUR3* transcript levels increased under nitrogen starvation while under adequate nitrogen supply low levels of transcripts were observed (Liu et al. 2003). Under low nitrogen conditions most of the *NRT* and *AMT* paralogs were up-regulated while a reverse trend of expression of these genes was observed under high nitrogen conditions (Sun et al. 2021). Previous studies in *Malus domestica*, *Cucumis sativus*, *Oryza sativa* and *A. thaliana* suggested that *CEP* genes are also induced in response to nitrogen deficiency (Li et al. 2018; Liu et al. 2021; Sui et al. 2016 and Chu et al. 2021). *NIN-like protein* (*NLP*) transcription factors regulate the suppression of *CEP* expression under high nitrogen availability (Luo et al. 2022). Amino acids are involved in sensing plant nitrogen status and maintaining nitrogen to carbon ratios in the plant. Glutamine and glutamate levels increase under high nitrogen availability and endogenous glutamine is known to play a role in feedback inhibition of nitrogen uptake through the suppression of genes encoding inorganic nitrogen transporters (Davenport, 2002; Xin et al. 2019). *LHT1*, a high-affinity amino acid transporter, was significantly up-regulated under nitrogen starvation conditions for maintaining N homeostasis in the plants (Huang et al. 2024). Down-regulation of *LHT1* under NUF treatment seems to prevent entry of amino acids into the cells and thus maintaining the N balance. Another NUF-down-regulated gene *AAT1* also regulates the N uptake and utilization efficiency (Guo et al. 2021). *ATL31* plays a very crucial role in maintaining the carbon and nitrogen balance, controling the stability of 14-3-3 chaperones at the same time. Under low nitrogen availability, *ATL31* was transcriptionally induced and involved in senescence progression in plants (Aoyama et al. 2014). Calcium is a very crucial element for plant growth as it plays a role in both nitrogen absorption and assimilation (Xing et al. 2021). *CNGC11*, *CNGC13* and *CNGC20* are the transporters which increase the cellular calcium concentration by transporting it from the extracellular environment or the vacuole to the cytoplasm. These genes were up-regulated under urea suggesting that there is a need for more absorption and assimilation of nitrogen, while NUF suppressed these paralogs in different degrees. *PUP*, involved in uptake of nitrogenous compounds like allantoin/ thiamine/ purine, is again induced under nitrogen deficient conditions, while suppressed by NUF, possibly due to nitrogen excess (Izaguirre-Mayoral et al. 2018).

On the other hand, higher induction of amino acid assimilation genes was observed under NUF than urea, which correlates with our estimated higher amino acid content in NUF-treated leaves (Table1, Fig. 1, 6). *ASN1* is a key enzyme in asparagine synthesis from glutamine for nitrogen storage and transport (Lam et al. 1998). High nitrogen availability contributes to accumulation of glutamate and glutamine, increasing the substrate for *ASN1*, which explains the upregulation of *ASN1* under NUF treatment. The *bZIP1* transcription factor is induced by nitrogen treatment (Obertello et al., 2010). NUF-induced *bZIP1* induces the expression of *ASN1, GDH2* and *ASP3* enhancing amino acid assimilation (Dietrich et al., 2011). *CML38* encodes a secretory protein that is activated under low nitrogen conditions and, in turn, activates brassinosteroid signaling through RALF1 and PEPR2, leading to growth inhibition. However, under nitrogen excess conditions after NUF spray, *CML38* is down-regulated, reducing its impact on growth inhibition (Song et al. 2021). Expression of tyrosine aminotransferase gene (*TAT2/3*) decreased under low nitrogen availability (Zhang et al., 2018). Similarly, *CORI3*, another tyrosine transaminase for tyrosine catabolism into secondary metabolites for plant defense and nutrient recycling, was induced by NUF but suppressed by urea (Prabhu and Hudson 2010). Deficiency of amino acids or glucose induces *GFAT1*, a rate limiting enzyme of hexosamine biosynthetic pathway (Chaveroux et al. 2016). Up-regulation of *GFAT1* under urea treatment but down-regulation under NUF corroborates the higher leaf amino acid content under NUF than urea (Fig. 1E). Application of urea and NUF increased the leaf chlorophyll content of *A. thaliana*. However,

NUF treated leaves showed significantly higher chlorophyll content than urea (Fig. 1C). The application of other nanofertilizers like TiO_2_ and Fe_2_O_3_ are known to perturb chlorophyll metabolism (Farahi et al. 2023; Rui et al. 2016). We observed that application of NUF induced *PORB* higher than urea which may explain higher chlorophyll levels under NUF compared to urea (Table 1, Fig. 6). PORB mediates a pivotal late-stage reaction in chlorophyll biosynthesis, the NADPH-dependent, light-induced reduction of protochlorophyllide (PChlide) to chlorophyllide (Chlide) (Garrone et al. 2015). Higher expression of other genes helped to increase chlorophyll biosynthesis or provide greater photoprotection under NUF. *ATOEP16-1* encodes a transporter which imports the precursor of PORA into the plastids and governs chloroplast formation at the time of greening (Samol et al. 2010). Thylakoid localized *CYP37* maintains the cytochrome b6/f (Cyt b6/f) complex under high light and thus directs the ETC to protect against light-induced stress (Yang et al. 2023). In addition NUF down-regulated the chlorophyll catabolism genes at greater levels than urea, thereby maintaining higher chlorophyll levels than urea. The chlorophyll catabolism gene *MES16* demethylates pheophorbide, an early chlorophyll breakdown intermediate, and fluorescent chlorophyll catabolites (FCCs) (Christ et al. 2011). On the other hand, *CYP89A9*, catalyzing the conversion of dioxobilin-type FCCs (DFCCs) from FCCs in the catabolism pathway, was induced under urea (Aubry et al. 2021).

Other than influencing the expression of nitrogen metabolism genes, NUF led to a significantly higher induction of other genes associated with growth and development, including cell wall biosynthesis or modification, carbohydrate, lipid, and hormone metabolism, which can explain the higher biomass obtained after foliar sprays with NUF compared to equimolar bulk urea. Other nano fertilizers enhance plant productivity by targeting hormone metabolism, facilitating cell wall formation, promoting cell division, and driving the synthesis of carbohydrates and lipids (Yadav et al. 2023).

Modification of the cell wall is crucial for cell growth and differentiation. The significantly higher induction of cell wall biosynthesis and modification genes can explain higher growth observed under NUF treatment than urea. NUF-induced *PMEI13* functions as a mediator of cellular adhesion, modulates cell wall permeability and enhances its flexibility (Coculo and Lionetti, 2022). Similarly, *PLL1* is implicated in numerous dimensions of growth and development, encompassing both processes dependent on and independent of cellular abscission, as well as contributing to the modulation of cambial activity and xylem differentiation (Sun and Nocker, 2010; Bush et al. 2022). Furthermore, a glycine-rich protein *GRP-5* regulates the development of the secondary cell wall framework (Mangeon et al. 2010). A cell wall glycoprotein, *AGP7*, significantly directs cell wall expansion, and signal transduction, while also alleviating various environmental stresses, thus fostering plant development (Ma and Johnson, 2023). *EXPB3* loosens the rigid carbohydrate matrix of the cell wall, modifying it during growth and vascular differentiation (Sampedro and Cosgrove, 2005). *CRR2*, a receptor-like kinase, is involved in various plant developmental processes, including the modulation of stem cell identity, differentiation in the root tip, and development of epidermal cells (Pu et al. 2012; Nikonorova et al. 2015).

Carbohydrates and lipids are fundamental building blocks and energy sources for plants, exerting an essential aspect in numerous biological processes such as photosynthesis, defense mechanisms, and cellular signaling. Their metabolism underpins the efficient storage of carbon and energy, facilitating overall plant growth and development (Gu et al. 2020; Du et al. 2021). Micronutrient nanofertilizers, especially zinc, govern the synthesis of carbohydrates (Broadley et al. 2007) while selenium nanoparticles stimulate the synthesis of lipids necessary for plant development (Zhang et al. 2024). NUF induced several carbohydrate and lipid metabolism genes while these were mostly down-regulated under urea. Out of these, *UGT76C5* leads to the production of secondary metabolites like flavonoids and terpenoids, influencing the growth and development of plant organs (Hoffmann et al. 2023; Wang et al. 2023). *BGLU18* involves the synthesis of ABA via a single-step cleavage of the inactive ABA-glucose ester, a process vital for improving resistance to abiotic stress (Han et al. 2019). Moreover, NUF-induced *LTP3* is involved in transporting cutin monomers and wax across the cell wall helps to maintain the cuticular layer structure (Pyee and Kolattukudy, 1995). Additionally, it positively regulates ABA biosynthesis (Gao et al. 2016). *CER1*, encoding an alkane biosynthesis enzyme, and *KCS8*, linked with the biosynthesis of very long-chain fatty acids and waxes within vegetative tissues, are essential for growth (Bourdenx et al. 2011; Bach and Faure 2010).

Cell division plays a crucial role in growth processes. Interestingly, various genes of the cell cycle regulation showed higher expression levels under NUF compared to urea. Previously the application of silver nanoparticles regulated cell division resulting in accelerated Arabidopsis growth (Syu et al. 2014). NUF caused a lesser suppression of *AGP8*, a glycoprotein integral to cellular expansion and division, and xylem differentiation, regulating overall plant growth and development, than urea (Liu et al. 2020). A pivotal plant growth-promoting gene *FH10* modulates cell morphogenesis by regulating cellular architecture and expansion as well as assisting in the localization and alignment of cell plates during cytokinesis (Cvrckova, 2012). *CYCD1;1* regulates cell cycle progression from the G1 to the S phase and is also involved in leaf development (Cho et al. 2004). Suppression of *FH10* and *CYCD1;1* under urea but no change under NUF compared to the mock treatment points to the differential mode of action of the nano form and may contribute to the observed higher leaf biomass under NUF treatment than urea.

Increased expression levels of some phytohormone biosynthesis and signaling genes under NUF correlated with significantly higher plant biomass than urea. Phytohormone pathway genes were triggered in response to various nanoparticle exposures such as CuNPs, AgNPs, AuNPs, ZnONPs, and FeNPs (Tripathi et al. 2022). NUF-induced *AOC1* and *AOP2* regulate JA biosynthesis and signaling and ensure plant survival under biotic and abiotic stressors (Burow et al. 2015; Li et al. 2019). *AtTSA1* is induced by JA leading to the formation of ER bodies in leaf and root tissues to counter stress (Geem et al. 2019). Moreover, *bHLH093* and *KS1*, act as crucial enzymes for gibberellins (GA) biosynthesis and regulate plant growth and development, cell elongation, foliage expansion, and leafy crown formation (Ritonga et al. 2023; Oshikawa et al. 2024). The expression of *ICS2*, an essential enzyme in the SA biosynthesis pathway, converts chorismate into isochorismate and SA functions as an essential intermediary in photosynthesis by contributing to stomatal closure, affecting chlorophyll and carotenoid levels, and promoting the activity of RuBisCO (Vicente and Plasencia, 2011; Guo et al. 2023). NUF-induced *SAUR29* and *SAUR65* of the auxin signaling pathway promote cell elongation and serve a pivotal role in governing hypocotyl development (Ren and Gray, 2015; Stortenbeker and Bemer, 2019).

We observed that many stress-responsive genes exhibited higher induction under NUF treatment than urea. Nanofertilizers generally trigger stress-responsive genes and greatly improve plant tolerance to biotic and abiotic stresses by modulating diverse physiological, biochemical, and molecular mechanisms (Khalid et al. 2022; Saadony et al. 2022). These stress-responsive genes enhance plant growth and development by interacting with growth pathways eventually boosting physiological processes like photosynthesis, hormonal regulation, and production of cellular protective molecules like soluble sugars and antioxidants (Zhang et al. 2020; Azmat et al. 2022). NUF-induced *CBL10* regulates Na^+^ homeostasis and plant growth under salt stress (Monihan et al. 2016; Plasencia et al. 2021). NUF-induced stress-responsive *bZIP1* plays a role in biomass enhancement through sugar signaling (Sulpice et al. 2009). *ANNAT4* proteins are crucial in biological processes such as vesicular trafficking, cytoskeletal dynamics, cell cycle regulation, and ion flux modulation, which contribute to cellular homeostasis and promote plant growth and development (Yadav et al. 2018). Furthermore, induction of c*ytochrome b561/ferric reductase transmembrane with DOMON related domain* (AT3G61750) promotes the meristematic cell division and positively modulates root development facilitating iron uptake by plants (Clúa et al. 2024). Nanoparticles trigger the production of reactive oxygen species (ROS) within cells (Manke et al. 2013). Hence, we observed NUF triggered higher expression levels of genes encoding antioxidant enzymes, *GSTU17* and *PRX32*, and the raffinose-producing gene *GolS3* which protects plants from oxidative stress (Estévez et al. 2020; Liu et al. 2021; Martins et al. 2022).

Interestingly urea foliar application induced the abscission and senescence genes, whereas NUF suppressed these genes (Table S3). Urea is viewed as a metabolic marker for senescence, being produced in high concentrations in senescent leaves, and is translocated out of aging tissues by enhancing the expression of *DUR3* (Bohner et al. 2015). Application of various nanoparticles like CoNPs and AgNPs alter the abscission and senescence in plants (Ha et al. 2020). NUF possibly enhances growth greater than urea through delaying abscission and senescence through down-regulation of the genes involved. For example, down-regulation of *RLK5* and its interaction with *CST* results in delaying abscission via a signal transduction cascade involving the peptide ligand IDA, the *receptor-like kinases* (RLKs) *HAE* and *HSL2*, and a subsequent MAP kinase (MAPK) pathway (Burr et al. 2011; Meng et al. 2016). Reduced expression of *MKK4* under NUF dampens the cellular responses that facilitate the separation of cells at the abscission zone (Trigo et al. 2024). High expression levels of *SAG101* disrupts the membrane structure and releases fatty acids to accelerate senescence (He and Gan, 2002). However, NUF-directed suppression of *SAG101* delays the senescence procedure. Similarly, negative regulation of *WRKY53* by NUF shuts off its epigenetic regulation to delay senescence (Bakshi and Oelmüller, 2014). *DLO1* regulates the timing and rate of leaf senescence and also prevents the SA buildup which in turn leads to the overstimulation of plant own defenses and negatively impacts growth and development (Zeilmaker et al. 2014). NUF-directed down-regulation of *DLO1* enhanced growth in this fashion. In a similar vein, suppression of many defense response genes by NUF may add to its growth promotion in *A. thaliana*, compared to urea which induced these genes (Table S3). In addition, the inhibition of *DA1* by NUF leads to unrestricted cell division, reducing leaf senescence and enhancing leaf growth (Du et al. 2014; Vanhaeren et al. 2017).

In conclusion, our results indicate foliar application of NUF is advantageous compared to equimolar urea for the vegetative growth and leaf nitrogen, chlorophyll, and amino acid content of the model organism *A. thaliana* grown in a low nitrogen background. At the molecular level, a comparative transcriptome between NUF and urea indicated that NUF upregulated genes associated with growth promotion, specifically those involved in cell wall biosynthesis, cell division, amino acid assimilation, and carbohydrate, and lipid metabolism. In contrast, it down-regulated genes linked to cell death, senescence, and abscission. Down-regulation of genes for nitrogen uptake under NUF was typical of a nitrogen-excess condition. However, these findings in the laboratory must be verified in the field conditions. Nevertheless, our results indicate considerable promise of NUF for improving crop biomass yield, with the capacity to revolutionize the agricultural sector.

## Data availability statement

The transcriptome raw data are available in the NCBI SRA database with BioProject accession number PRJNA1129292.

## Author contributions

AD, NJ, and DV conducted most experiments and prepared the original draft. RSS and PY conducted transcriptome data analysis. AS conceived the study, acquired funding, designed and supervised the experiments, curated the data, and edited the original draft.

## Acknowledgments

The authors thank the Department of Bioscience and Bioengineering (BSBE), IIT Jodhpur, and the Nano Biotechnology Research Center, Kalol, for providing laboratory facilities for the study. The guidance of Prof. Mitali Mukerji, Head, BSBE, and Prof. Prabuddha Ganguli, Advisor and Adjunct Faculty, School of Management and Entrepreneurship, is gratefully acknowledged.

