## Supplemental files for "Foliar application of nano urea results in higher biomass, chlorophyll, and nitrogen content than equimolar bulk urea through differential gene regulation in *Arabidopsis thaliana*"

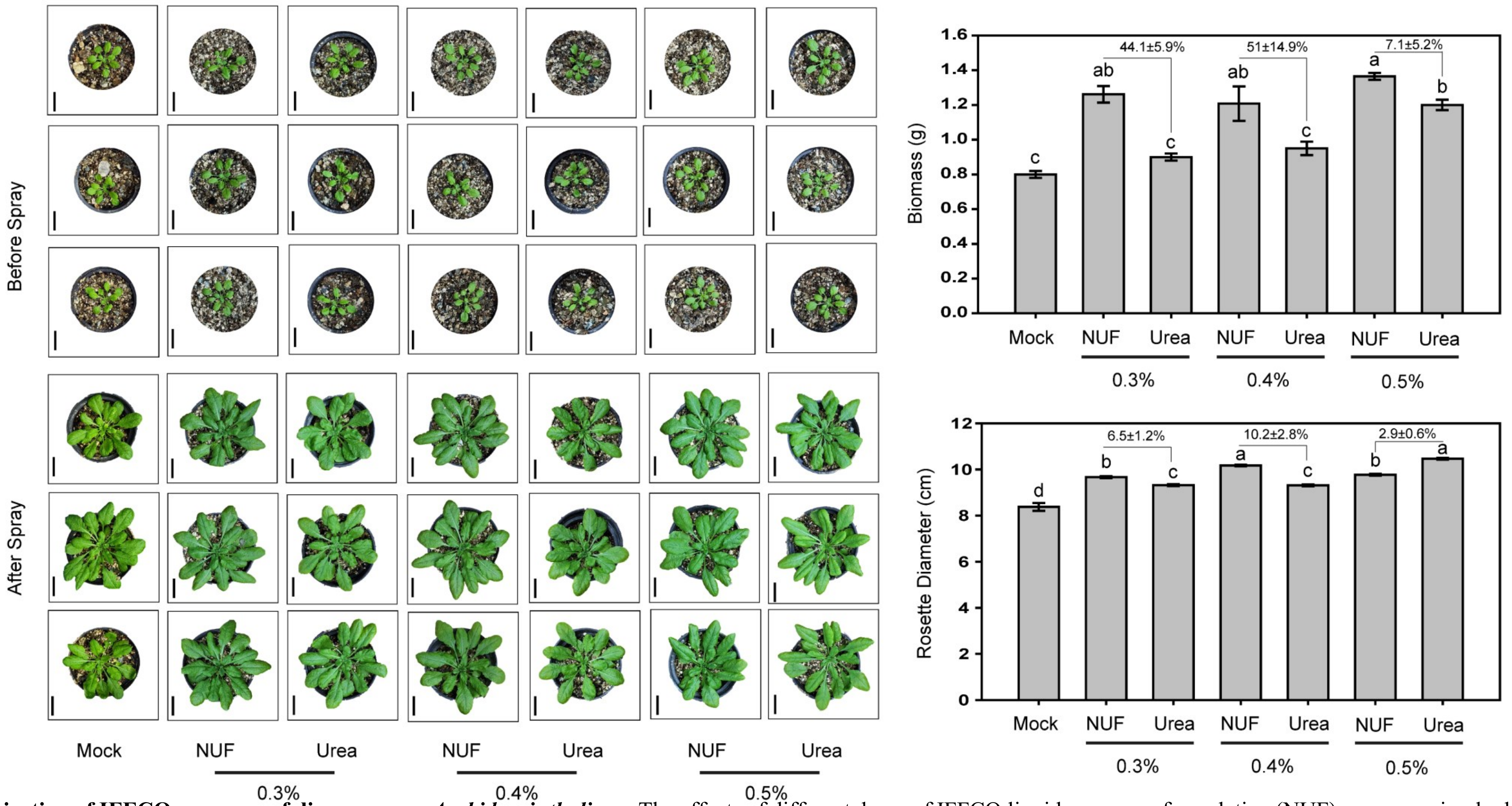

**Fig.S1. Dose optimization of IFFCO nano urea foliar spray on *Arabidopsis thaliana*.** The effects of different doses of IFFCO liquid nano urea formulation (NUF) versus equimolar bulk urea on biomass are shown. Arabidopsis plants were grown for a month in vermiculite in 2.5" pots watered with ¼-Hoagland's solution every alternate day. NUF contains 1.5 M urea, as per the manufacturer. For uniformity, a stock aqueous solution of 1.5 M Urea was prepared. 0.3%, 0.4% and 0.5% dilutions were prepared from the urea and NUF stock solutions and used to spray one-month-old *A. thaliana* plants. The plants were sprayed twice in 10-d intervals with 0.4% NUF or 0.4% urea at both the abaxial and adaxial leaf surfaces. Mock plants were sprayed with distilled water and photographed. The shoots were harvested to measure biomass, total chlorophyll, total nitrogen, and total amino acid content after 20 d. The shoot fresh weights were measured after 20-d growth in mock, NUF, and urea and presented as biomass in grams. Bars indicate the average of 10 seedlings, with standard error. Different alphabets indicate significant differences (Tukey's HSD test, $P <$ 0.05, N = 10).

**Table S1.** ***Arabidopsis thaliana*** **genes differentially expressed between IFFCO nano urea and equimolar urea foliar spray treatments.**

| Gene ID | NUF vs Urea | | NUF vs Mock | | Urea vs Mock | | Gene Symbol | TAIR Short Description |
|---|---|---|---|---|---|---|---|---|
| | $log_2FC$ | *FDR* | $log_2FC$ | *FDR* | $log_2FC$ | *FDR* | | |
| AT5G59320 | 4.05 | 6.05E-06 | 1.67 | 1.91E-02 | -2.39 | 2.02E-08 | LTP3 | Non-specific lipid-transfer protein 3 |
| AT4G11320 | 3.94 | 1.45E-04 | 1.93 | 4.09E-02 | -2.07 | 3.91E-02 | RDL5 | Probable cysteine protease RDL5 |
| AT5G24770 | 3.06 | 9.40E-03 | 2.85 | 8.59E-04 | -0.23 | 6.64E-01 | VSP2 | Vegetative storage protein 2 |
| AT1G02205 | 2.29 | 6.79E-04 | 0.17 | 8.68E-01 | -2.14 | 1.36E-05 | CER1 | Fatty acid hydroxylase superfamily |
| AT1G14250 | 2.27 | 7.74E-04 | 1.31 | 1.13E-02 | -0.98 | 2.04E-03 | APY5 | Probable apyrase 5 |
| AT4G02290 | 2.06 | 1.15E-02 | 1.15 | 2.99E-01 | -0.90 | 4.96E-01 | GH9B13 | Endoglucanase |
| AT1G52400 | 2.05 | 5.94E-03 | 3.30 | 1.06E-15 | 1.24 | 5.36E-03 | BGLU18 | Beta-D-glucopyranosyl abscisate beta-glucosidase |
| AT1G69140 | 2.04 | 2.01E-05 | 0.93 | 1.23E-02 | -1.13 | 1.08E-02 | | pseudogene |
| AT5G41590 | 2.04 | 1.28E-06 | 0.58 | 2.24E-01 | -1.48 | 1.15E-03 | At5g41590 | Uncharacterized protein |
| AT4G17470 | 2.04 | 8.86E-04 | 2.26 | 4.12E-06 | 0.22 | 8.63E-01 | At4g17470 | Alpha/beta-Hydrolases superfamily protein |
| AT2G02100 | 2.01 | 2.79E-09 | 1.10 | 1.15E-05 | -0.91 | 3.38E-02 | PDF2.2 | Defensin-like protein 2 |
| AT2G42540 | 1.98 | 1.39E-05 | 0.28 | 5.89E-01 | -1.72 | 2.38E-07 | COR15A | Cold-regulated 15a |
| AT1G45616 | 1.97 | 9.92E-06 | 0.52 | 4.30E-01 | -1.46 | 9.05E-03 | RLP6 | Receptor-like protein 6 |
| AT5G45830 | 1.87 | 4.40E-03 | 0.53 | 2.45E-01 | -1.35 | 6.36E-02 | DOG1 | Delay of germination 1 |
| AT3G43715 | 1.81 | 4.29E-05 | 0.29 | 4.79E-01 | -1.53 | 7.36E-04 | | transposable element gene |
| AT3G15950 | 1.78 | 2.17E-02 | 2.84 | 1.42E-06 | 1.05 | 2.36E-01 | NAI2 | DNA topoisomerase-like protein |
| AT3G47340 | 1.77 | 1.18E-03 | 1.13 | 1.57E-07 | -0.65 | 2.76E-01 | ASN1 | Asparagine synthetase |
| AT3G09162 | 1.74 | 1.23E-02 | 0.32 | 7.29E-01 | -1.42 | 5.69E-02 | At3g09162 | At3g09162 |
| AT3G27250 | 1.73 | 5.52E-04 | -0.07 | 9.81E-01 | -1.81 | 5.26E-05 | At3g27250 | hypothetical protein |
| AT5G35777 | 1.70 | 1.63E-02 | 1.76 | 2.44E-04 | 0.07 | 9.65E-01 | | transposable_element_gene |
| AT1G78170 | 1.70 | 6.56E-03 | 0.02 | 9.88E-01 | -1.69 | 5.58E-03 | At1g78170 | Uncharacterized protein |
| AT5G49525 | 1.70 | 3.33E-03 | 2.10 | 1.98E-03 | 0.39 | 8.03E-01 | At5g49525 | At5g49525 |
| AT2G42170 | 1.67 | 2.04E-02 | 0.24 | 8.62E-01 | -1.46 | 2.92E-02 | At2g42170 | Actin family protein |
| AT2G42530 | 1.66 | 2.26E-04 | 0.68 | 1.17E-01 | -0.99 | 1.16E-03 | COR15B | Protein COLD-REGULATED 15B, chloroplastic |
| AT5G62280 | 1.64 | 2.49E-02 | 0.74 | 2.15E-01 | -0.91 | 3.23E-01 | At5g62280 | |
| AT1G56650 | 1.63 | 4.33E-02 | 1.66 | 2.22E-02 | 0.01 | 9.95E-01 | MYB75 | Transcription factor MYB75 |
| AT4G23600 | 1.61 | 7.49E-03 | 0.82 | 5.21E-03 | -0.81 | 1.43E-01 | CORI3 | Tyrosine transaminase family protein |
| AT5G01370 | 1.61 | 1.69E-02 | 0.57 | 5.71E-01 | -1.05 | 1.93E-01 | ACI1 | ALC-interacting protein 1 |
| AT1G52040 | 1.59 | 3.17E-02 | 2.62 | 1.06E-15 | 1.01 | 1.88E-01 | MBP1 | Myrosinase-binding protein 1 |
| AT2G37025 | 1.58 | 4.72E-02 | -0.10 | 9.28E-01 | -1.69 | 2.90E-02 | TRFL8 | TRF-like 8 |
| AT4G30650 | 1.56 | 4.41E-04 | 0.64 | 1.62E-03 | -0.93 | 4.44E-02 | At4g30650 | UPF0057 membrane protein At4g30650 |
| AT2G29680 | 1.56 | 5.93E-03 | 0.20 | 9.03E-01 | -1.38 | 3.87E-02 | CDC6 | Cell division control protein 6 homolog |
| AT5G62920 | 1.55 | 2.17E-02 | -0.04 | 9.68E-01 | -1.60 | 1.43E-02 | ARR6 | Two-component response regulator ARR6 |
| AT4G03610 | 1.55 | 3.00E-02 | 1.17 | 4.87E-02 | -0.39 | 7.70E-01 | At4g03610 | Metallo-hydrolase/oxidoreductase superfamily protein |
| AT5G63160 | 1.54 | 5.25E-04 | 1.22 | 6.58E-05 | -0.33 | 5.16E-01 | BT1 | BTB and TAZ domain protein 1 |
| AT4G08100 | 1.50 | 2.17E-02 | 0.01 | 1.00E+00 | -1.51 | 1.61E-02 | | transposable_element_gene |
| AT1G62975 | 1.50 | 4.46E-02 | 0.54 | 6.46E-01 | -0.96 | 2.97E-01 | At1g62975 | Basic helix-loop-helix |
| AT1G75240 | 1.50 | 2.26E-02 | 1.19 | 3.71E-03 | -0.33 | 7.34E-01 | ZHD5 | Zinc-finger homeodomain protein 5 |
| AT5G51390 | 1.48 | 9.22E-03 | 0.79 | 3.14E-01 | -0.72 | 4.58E-01 | At5g51390 | At5g51390 |
| AT5G44010 | 1.46 | 1.92E-02 | 0.25 | 8.39E-01 | -1.22 | 2.91E-02 | MRH10.12 | Fanconi anemia group F protein |
| AT5G15970 | 1.45 | 1.20E-03 | 0.91 | 6.56E-03 | -0.56 | 1.14E-01 | KIN2 | Stress-induced protein KIN2 |
| AT4G08115 | 1.45 | 1.79E-02 | -0.29 | 7.72E-01 | -1.75 | 2.32E-03 | | |
| AT5G05890 | 1.44 | 3.68E-02 | 0.67 | 6.86E-02 | -0.78 | 3.61E-01 | UGT76C5 | UDP-glycosyltransferase 76C5 |
| AT4G03355 | 1.44 | 4.53E-02 | 0.07 | 9.82E-01 | -1.39 | 4.46E-02 | | |
| AT5G65640 | 1.43 | 1.17E-03 | 0.21 | 5.73E-01 | -1.23 | 6.93E-03 | BHLH93 | Transcription factor bHLH93 |
| AT1G80570 | 1.43 | 1.32E-02 | 0.88 | 1.62E-01 | -0.57 | 5.41E-01 | FBL14 | F-box/LRR-repeat protein 14 |
| AT4G21970 | 1.43 | 3.03E-02 | 0.18 | 8.91E-01 | -1.27 | 5.83E-02 | At4g21970 | Uncharacterized protein AT4g21970 |
| AT1G18870 | 1.42 | 2.89E-03 | 0.46 | 4.30E-01 | -0.99 | 1.09E-02 | ICS2 | isochorismate synthase |
| AT2G16750 | 1.42 | 4.83E-03 | 0.25 | 6.85E-01 | -1.18 | 3.46E-02 | STU | Protein kinase STUNTED |
| AT5G35670 | 1.40 | 2.79E-03 | 0.72 | 3.47E-01 | -0.69 | 4.22E-01 | IQD33 | Protein IQ-DOMAIN 33 |
| AT1G33615 | 1.38 | 2.54E-02 | -0.09 | 9.65E-01 | -1.48 | 2.53E-02 | | |
| AT1G58270 | 1.38 | 2.56E-03 | -0.02 | 1.00E+00 | -1.41 | 1.37E-06 | At1g58270 | Expressed protein |
| AT4G02850 | 1.38 | 3.99E-03 | -0.79 | 6.46E-02 | -2.18 | 4.00E-08 | DAAR1 | At4g02850 |
| AT1G56670 | 1.38 | 3.97E-02 | 0.65 | 1.65E-01 | -0.74 | 3.40E-01 | At1g56670 | GDSL-like Lipase/Acylhydrolase superfamily protein |
| AT5G34795 | 1.37 | 1.20E-03 | 0.41 | 3.64E-01 | -0.98 | 2.38E-02 | | pseudogene |
| AT5G07700 | 1.37 | 2.76E-04 | 1.01 | 3.50E-03 | -0.37 | 5.42E-01 | MYB76 | Transcription factor MYB76 |
| AT2G23000 | 1.36 | 2.37E-02 | -0.28 | 3.51E-01 | -1.65 | 2.59E-03 | SCPL10 | Serine carboxypeptidase-like 10 |
| AT4G01080 | 1.36 | 4.66E-02 | 0.53 | 1.62E-01 | -0.84 | 1.79E-01 | TBL26 | Protein trichome birefringence-like 26 |
| AT1G52000 | 1.36 | 3.46E-03 | 1.47 | 3.87E-08 | 0.11 | 8.40E-01 | JAL5 | Jacalin-related lectin 5 |
| AT2G23010 | 1.35 | 4.66E-03 | 1.11 | 2.40E-02 | -0.26 | 7.27E-01 | SCPL9 | Serine carboxypeptidase-like 9 |
| AT3G14450 | 1.35 | 5.18E-03 | 0.25 | 7.49E-01 | -1.10 | 2.56E-02 | CID9 | Polyadenylate-binding protein-interacting protein 9 |
| AT2G01760 | 1.34 | 1.71E-02 | 0.59 | 1.06E-01 | -0.77 | 2.72E-01 | RR14 | Response regulator 14 |
| AT4G34900 | 1.34 | 4.67E-04 | 0.06 | 9.12E-01 | -1.29 | 7.12E-04 | XDH2 | Xanthine dehydrogenase 2 |
| AT5G22930 | 1.33 | 3.19E-02 | 0.47 | 6.06E-01 | -0.87 | 2.09E-01 | At5g22930 | Enabled-like protein |
| AT5G19730 | 1.33 | 9.86E-03 | -0.28 | 4.11E-01 | -1.62 | 4.44E-04 | At5g19730 | Pectinesterase |
| AT2G39180 | 1.32 | 2.49E-02 | 0.12 | 9.42E-01 | -1.21 | 1.73E-02 | CCR2 | Serine/threonine-protein kinase-like protein CCR2 |
| AT2G43550 | 1.32 | 9.20E-04 | 0.63 | 1.26E-01 | -0.69 | 2.01E-01 | At2g43550 | Knottin scorpion toxin-like domain-containing protein |
| AT1G12845 | 1.31 | 1.51E-02 | 0.37 | 5.74E-01 | -0.97 | 3.18E-02 | F13K23.10 | F13K23.10 protein |
| AT3G56810 | 1.31 | 1.67E-02 | 0.15 | 9.47E-01 | -1.18 | 9.20E-02 | At3g56810 | Uncharacterized protein |
| AT2G38995 | 1.31 | 2.58E-02 | 0.25 | 8.38E-01 | -1.08 | 1.36E-01 | At2g38995 | O-acyltransferase |
| AT3G03820 | 1.31 | 4.64E-02 | 0.34 | 5.29E-01 | -0.99 | 1.74E-01 | SAUR29 | Auxin-responsive protein SAUR29 |
| AT2G39240 | 1.30 | 4.71E-02 | 0.51 | 6.44E-01 | -0.81 | 3.06E-01 | At2g39240 | RNA polymerase I specific transcription initiation factor RRN3 family protein |
| AT5G23820 | 1.30 | 3.85E-02 | 0.58 | 4.21E-01 | -0.74 | 1.25E-01 | ML3 | MD-2-related lipid-recognition protein 3 |

| | | | | | | | | |
|---|---|---|---|---|---|---|---|---|
| AT4G28250 | 1.30 | 2.22E-03 | 0.68 | 5.85E-03 | -0.64 | 1.90E-01 | At4g28250 | Putative beta-expansin/allergen protein |
| AT3G54910 | 1.30 | 4.67E-02 | 0.21 | 6.94E-01 | -1.11 | 1.10E-01 | F28P10.110 | Uncharacterized protein F28P10.110 |
| AT1G10370 | 1.29 | 7.07E-05 | -0.24 | 5.02E-01 | -1.55 | 2.38E-07 | GSTU17 | Glutathione S-transferase U17 |
| AT3G63450 | 1.29 | 4.26E-03 | -0.51 | 3.04E-01 | -1.81 | 3.07E-06 | At3g63450 | At3g63450 |
| AT3G25760 | 1.28 | 3.34E-02 | 1.10 | 6.68E-05 | -0.19 | 7.67E-01 | AOC1 | Allene oxide cyclase 1, chloroplastic |
| AT3G63200 | 1.28 | 1.17E-02 | -0.30 | 6.24E-01 | -1.60 | 4.00E-06 | PLP9 | Probable inactive patatin-like protein 9 |
| AT2G37790 | 1.28 | 4.95E-02 | 0.14 | 8.92E-01 | -1.16 | 9.59E-02 | AKR4C10 | Aldo-keto reductase family 4 member C10 |
| AT5G51750 | 1.28 | 2.06E-04 | -0.37 | 3.22E-01 | -1.67 | 4.86E-08 | SBT1.3 | Subtilisin-like protease SBT1.3 |
| AT4G08114 | 1.28 | 2.72E-02 | 0.51 | 4.67E-01 | -0.79 | 2.62E-01 | | |
| AT2G41050 | 1.28 | 4.06E-02 | 0.02 | 1.00E+00 | -1.26 | 1.16E-01 | At2g41050 | PQ-loop repeat family protein / transmembrane family protein |
| AT1G27030 | 1.27 | 1.11E-02 | 0.99 | 2.58E-08 | -0.29 | 6.58E-01 | At1g27030 | 2-oxoadipate dioxygenase/decarboxylase |
| AT1G05420 | 1.27 | 2.88E-02 | 0.85 | 1.23E-01 | -0.44 | 5.54E-01 | At1g05420 | Transcription repressor |
| AT4G01335 | 1.27 | 1.93E-02 | 0.39 | 6.65E-01 | -0.89 | 1.54E-01 | At4g01335 | TATA box-binding protein associated factor RNA polymerase I subunit B-like protein |
| AT4G03060 | 1.27 | 3.14E-02 | 1.16 | 4.05E-12 | -0.12 | 8.89E-01 | AOP2 | Probable inactive 2-oxoglutarate-dependent dioxygenase AOP2 |
| AT2G16630 | 1.26 | 3.22E-02 | 0.03 | 1.00E+00 | -1.25 | 2.28E-02 | At2g16630 | Uncharacterized protein At2g16630 |
| AT2G33570 | 1.26 | 6.65E-04 | -0.07 | 9.32E-01 | -1.34 | 2.14E-07 | GALS1 | Galactan beta-1,4-galactosyltransferase GALS1 |
| AT1G52410 | 1.25 | 3.45E-02 | 2.22 | 2.76E-11 | 0.96 | 6.77E-03 | TSA1 | TSK-associating protein 1 |
| AT2G43530 | 1.25 | 2.58E-02 | 1.35 | 8.98E-05 | 0.10 | 8.67E-01 | ATTI3 | Defensin-like protein 194 |
| AT5G65390 | 1.24 | 5.63E-03 | 0.33 | 4.19E-01 | -0.92 | 6.91E-03 | AGP7 | Classical arabinogalactan protein 7 |
| AT5G62360 | 1.24 | 1.72E-02 | 1.63 | 9.61E-13 | 0.37 | 6.43E-01 | At5g62360 | Pectinesterase inhibitor domain-containing protein |
| AT1G68810 | 1.22 | 4.86E-02 | 0.43 | 5.13E-01 | -0.81 | 1.56E-01 | BHLH30 | Transcription factor bHLH30 |
| AT3G11490 | 1.20 | 2.54E-02 | -0.13 | 7.93E-01 | -1.33 | 8.65E-03 | ROPGAP4 | Rho GTPase-activating protein 4 |
| AT2G38750 | 1.19 | 3.27E-02 | 0.44 | 5.21E-01 | -0.76 | 1.61E-01 | At2g38750 | AT2G38750 protein |
| AT3G05640 | 1.18 | 4.00E-02 | 0.26 | 1.95E-01 | -0.94 | 1.07E-01 | At3g05640 | Probable protein phosphatase 2C 34 |
| AT1G09350 | 1.18 | 2.50E-02 | 0.92 | 2.46E-02 | -0.28 | 6.96E-01 | GOLS3 | Hexosyltransferase |
| AT5G06670 | 1.18 | 3.96E-02 | 0.21 | 6.24E-01 | -0.99 | 7.60E-02 | KIN75 | Kinesin-like protein |
| AT3G61950 | 1.18 | 4.28E-03 | -0.10 | 9.14E-01 | -1.30 | 4.91E-04 | MYC67 | Basic helix-loop-helix |
| AT1G77960 | 1.18 | 4.71E-02 | 0.49 | 1.85E-02 | -0.70 | 2.49E-01 | At1g77960 | Repressor ROX1-like protein |
| AT4G40065 | 1.18 | 8.17E-03 | 0.65 | 8.74E-02 | -0.54 | 3.15E-01 | | |
| AT5G52120 | 1.17 | 1.20E-03 | 0.27 | 3.73E-01 | -0.92 | 1.08E-02 | At5g52120 | F-box domain-containing protein |
| AT5G61290 | 1.16 | 3.42E-02 | 0.67 | 2.42E-01 | -0.52 | 3.57E-01 | At5g61290 | Uncharacterized protein |
| AT2G20670 | 1.16 | 1.85E-02 | 0.98 | 2.41E-08 | -0.20 | 7.53E-01 | At2g20670 | |
| AT2G28900 | 1.16 | 4.85E-03 | 0.71 | 6.40E-04 | -0.47 | 3.44E-01 | OEP161 | Outer envelope pore protein 16-1, chloroplastic |
| AT4G12900 | 1.16 | 3.30E-02 | 0.11 | 8.23E-01 | -1.07 | 4.70E-02 | At4g12900 | Gamma interferon responsive lysosomal thiol |
| AT1G45201 | 1.16 | 1.63E-02 | 1.24 | 1.35E-07 | 0.06 | 9.16E-01 | TLL1 | At1g45200 |
| AT1G56720 | 1.16 | 4.97E-03 | 0.07 | 9.19E-01 | -1.10 | 7.33E-03 | At1g56720 | Protein kinase domain-containing protein |
| AT1G75190 | 1.14 | 3.40E-02 | 0.56 | 3.31E-01 | -0.60 | 3.38E-01 | At1g75190 | AT1G75190 protein |
| AT3G20300 | 1.14 | 4.53E-02 | 0.39 | 1.58E-01 | -0.76 | 1.65E-01 | At3g20300 | Uncharacterized protein At3g20300 |
| AT5G65730 | 1.14 | 2.70E-02 | -0.24 | 4.37E-01 | -1.40 | 8.95E-04 | XTH6 | Probable xyloglucan endotransglucosylase/hydrolase protein 6 |
| AT1G77580 | 1.14 | 3.85E-02 | 0.31 | 4.41E-01 | -0.83 | 1.08E-01 | VETH3 | Filament-like protein |
| AT5G01075 | 1.13 | 5.73E-03 | -0.32 | 5.12E-01 | -1.46 | 1.40E-06 | TWS1 | At5g01075 |
| AT4G18440 | 1.13 | 4.18E-02 | 0.43 | 2.01E-01 | -0.72 | 1.18E-01 | At4g18440 | Adenylosuccinate lyase |
| AT5G55250 | 1.13 | 4.22E-02 | 0.87 | 1.72E-01 | -0.26 | 8.00E-01 | IAMT1 | Indole-3-acetate O-methyltransferase 1 |
| AT1G65920 | 1.13 | 2.47E-03 | 0.79 | 1.42E-03 | -0.36 | 4.89E-01 | At1g65920 | F12P19.9 protein |
| AT1G02340 | 1.13 | 1.71E-02 | -0.20 | 7.87E-01 | -1.34 | 4.56E-07 | HFR1 | Basic helix-loop-helix |
| AT3G16690 | 1.13 | 9.81E-03 | 0.38 | 3.97E-01 | -0.77 | 1.31E-01 | SWEET16 | Bidirectional sugar transporter SWEET |
| AT5G23980 | 1.12 | 1.43E-03 | 0.70 | 1.41E-03 | -0.43 | 2.80E-01 | FRO4 | Ferric reduction oxidase 4 |
| AT3G07540 | 1.12 | 2.41E-03 | 0.09 | 8.86E-01 | -1.05 | 3.87E-04 | FH10 | Formin-like protein 10 |
| AT4G17460 | 1.12 | 2.17E-02 | 0.41 | 8.12E-02 | -0.72 | 1.37E-01 | HAT1 | Homeobox-leucine zipper protein HAT1 |
| AT5G45560 | 1.12 | 2.26E-02 | 0.52 | 3.20E-01 | -0.62 | 3.56E-01 | MFC19.23 | Pleckstrin homology |
| AT4G38370 | 1.11 | 3.17E-02 | 0.43 | 5.30E-01 | -0.69 | 2.24E-01 | At4g38370 | Phosphoglycerate mutase family protein |
| AT4G31820 | 1.11 | 6.74E-03 | 0.20 | 3.68E-01 | -0.93 | 2.13E-02 | ENP | Phototropic-responsive NPH3 family protein |
| AT3G12110 | 1.10 | 4.23E-02 | -0.92 | 2.09E-04 | -2.03 | 1.37E-06 | ACT11 | Actin-11 |
| AT5G20740 | 1.09 | 4.86E-02 | -0.20 | 8.60E-01 | -1.31 | 1.74E-02 | At5g20740 | Plant invertase/pectin methylesterase inhibitor superfamily protein |
| AT5G49450 | 1.09 | 3.49E-03 | 0.57 | 1.29E-02 | -0.54 | 2.10E-01 | BZIP1 | Basic leucine zipper 1 |
| AT4G34650 | 1.08 | 4.17E-02 | 0.10 | 9.43E-01 | -1.00 | 5.38E-02 | SQS2 | Squalene synthase 2 |
| AT1G12060 | 1.08 | 4.70E-02 | 0.07 | 9.72E-01 | -1.02 | 3.35E-02 | BAG5 | BAG family molecular chaperone regulator 5, mitochondrial |
| AT5G57340 | 1.08 | 9.74E-03 | 0.60 | 3.45E-04 | -0.50 | 2.67E-01 | MJB24.15 | Ras guanine nucleotide exchange factor Q-like protein |
| AT1G54740 | 1.07 | 6.11E-03 | 0.44 | 2.42E-01 | -0.64 | 1.17E-01 | T22H22.16 | T22H22.16 protein |
| AT2G15090 | 1.07 | 3.73E-02 | 0.36 | 2.46E-01 | -0.72 | 1.46E-01 | KCS8 | 3-ketoacyl-CoA synthase 8 |
| AT4G39510 | 1.07 | 8.09E-03 | 0.71 | 6.33E-04 | -0.38 | 4.56E-01 | CYP96A12 | AT4g39510/F23K16_140 |
| AT1G65900 | 1.06 | 4.86E-02 | -0.01 | 1.00E+00 | -1.08 | 4.23E-02 | At1g65900 | At1g65900/F12P19_7 |
| AT5G53880 | 1.06 | 1.44E-02 | 0.48 | 2.81E-01 | -0.59 | 1.62E-01 | At5g53880 | Uncharacterized protein |
| AT3G20470 | 1.04 | 6.81E-03 | 1.27 | 6.75E-04 | 0.22 | 7.71E-01 | At3g20470 | Uncharacterized protein |
| AT5G13400 | 1.03 | 4.94E-02 | 0.02 | 9.88E-01 | -1.03 | 4.00E-02 | NPF6.1 | Protein NRT1/ PTR FAMILY 6.1 |
| AT3G09540 | 1.03 | 9.89E-03 | 0.50 | 2.33E-02 | -0.54 | 2.29E-01 | At3g09540 | Pectate lyase |
| AT4G29360 | 1.02 | 9.19E-03 | 0.32 | 5.57E-01 | -0.71 | 1.17E-01 | At4g29360 | glucan endo-1,3-beta-D-glucosidase |
| AT5G23750 | 1.02 | 2.44E-02 | -0.15 | 8.63E-01 | -1.18 | 1.51E-04 | MRO11.21 | Remorin family protein |
| AT3G61750 | 1.02 | 1.62E-02 | 0.03 | 9.88E-01 | -1.01 | 3.67E-03 | At3g61750 | Cytochrome b561/ferric reductase transmembrane with DOMON related domain-containing protein |
| AT4G33000 | 1.00 | 4.56E-02 | 0.00 | 1.00E+00 | -1.02 | 3.39E-02 | At4g33000 | Calcineurin B-like protein |
| AT1G04240 | 1.00 | 2.52E-02 | 0.90 | 1.29E-04 | -0.11 | 8.74E-01 | SHY2 | Auxin-responsive protein |
| AT1G18650 | 1.00 | 4.86E-02 | -0.08 | 8.74E-01 | -1.09 | 2.23E-02 | PDCB3 | Plasmodesmata callose-binding protein 3 |
| AT4G13990 | 0.99 | 2.76E-02 | 0.54 | 3.72E-01 | -0.47 | 4.19E-01 | GT14 | Probable xyloglucan galactosyltransferase GT14 |
| AT5G07570 | 0.98 | 1.79E-02 | 0.02 | 1.00E+00 | -0.98 | 5.19E-03 | At5g07570 | Glycine/proline-rich protein |
| AT3G62550 | 0.97 | 1.73E-02 | 0.67 | 6.19E-03 | -0.31 | 3.82E-01 | At3g62550 | |
| AT5G26570 | 0.97 | 2.57E-02 | 0.55 | 2.34E-04 | -0.43 | 3.82E-01 | PWD | Chloroplastidic phosphoglucan, water dikinase |
| AT5G54145 | 0.97 | 2.73E-02 | 0.10 | 9.06E-01 | -0.89 | 2.75E-02 | At5g54145 | Transmembrane protein |
| AT3G11690 | 0.96 | 1.35E-02 | 0.03 | 9.68E-01 | -0.95 | 9.79E-03 | At3g11690 | |
| AT1G60690 | 0.96 | 1.32E-02 | 0.21 | 6.91E-01 | -0.77 | 6.23E-02 | At1g60690 | Probable aldo-keto reductase 3 |

| | | | | | | | | |
|---|---|---|---|---|---|---|---|---|
| AT5G01790 | 0.96 | 4.18E-02 | -0.21 | 5.06E-01 | -1.19 | 4.94E-03 | At5g01790 | At5g01790 |
| AT5G56550 | 0.96 | 2.69E-02 | 0.76 | 3.16E-02 | -0.20 | 7.64E-01 | OXS3 | Protein OXIDATIVE STRESS 3 |
| AT3G26700 | 0.95 | 3.12E-02 | 0.29 | 5.30E-01 | -0.69 | 1.70E-01 | At3g26700 | non-specific serine/threonine protein kinase |
| AT2G19860 | 0.95 | 7.72E-03 | 0.26 | 4.44E-01 | -0.71 | 5.27E-02 | HXK2 | Phosphotransferase |
| AT5G13460 | 0.95 | 3.07E-03 | 0.27 | 4.45E-01 | -0.70 | 6.22E-02 | IQD11 | Protein IQ-DOMAIN 11 |
| AT1G20650 | 0.95 | 1.57E-02 | 0.25 | 3.48E-01 | -0.71 | 9.61E-02 | PBL21 | Probable serine/threonine-protein kinase PBL21 |
| AT2G45470 | 0.95 | 1.57E-02 | -0.25 | 2.12E-01 | -1.21 | 2.34E-04 | FLA8 | Fasciclin-like arabinogalactan protein 8 |
| AT4G17180 | 0.94 | 3.66E-02 | 0.37 | 5.30E-01 | -0.59 | 2.32E-01 | DL4625W | glucan endo-1,3-beta-D-glucosidase |
| AT5G67070 | 0.94 | 9.67E-03 | 0.24 | 5.20E-01 | -0.71 | 1.31E-02 | RALFL34 | Protein RALF-like 34 |
| AT1G08890 | 0.94 | 3.31E-03 | 0.90 | 8.35E-08 | -0.05 | 9.34E-01 | At1g08890 | AT1G08890 protein |
| AT1G29460 | 0.94 | 2.35E-02 | 0.19 | 5.34E-01 | -0.76 | 6.66E-02 | SAUR65 | SAUR-like auxin-responsive protein family |
| AT2G30790 | 0.94 | 1.58E-02 | 0.63 | 1.28E-02 | -0.32 | 5.48E-01 | PSBP2 | Putative oxygen-evolving enhancer protein 2-2 |
| AT2G34010 | 0.93 | 2.61E-02 | 0.44 | 7.91E-02 | -0.51 | 2.82E-01 | TIE4 | Verprolin |
| AT5G16000 | 0.93 | 4.32E-02 | -0.42 | 2.13E-02 | -1.36 | 6.69E-04 | NIK1 | Protein NSP-INTERACTING KINASE 1 |
| AT4G24900 | 0.93 | 3.29E-02 | 0.25 | 6.78E-01 | -0.68 | 1.63E-01 | TTL | Coiled coil protein |
| AT3G32980 | 0.92 | 6.43E-03 | 0.67 | 1.08E-02 | -0.27 | 5.73E-01 | PER32 | Peroxidase 32 |
| AT4G31590 | 0.92 | 2.06E-02 | 0.08 | 8.97E-01 | -0.87 | 1.18E-02 | CSLC5 | Probable xyloglucan glycosyltransferase 5 |
| AT4G29140 | 0.92 | 3.73E-02 | 0.79 | 1.91E-02 | -0.15 | 8.22E-01 | DTX51 | Protein DETOXIFICATION 51 |
| AT1G70210 | 0.92 | 2.27E-02 | 0.13 | 7.38E-01 | -0.80 | 4.37E-02 | At1g70210 | CYCD1 |
| AT1G19570 | 0.91 | 2.69E-02 | 0.91 | 1.51E-05 | -0.01 | 9.88E-01 | DHAR1 | Glutathione S-transferase DHAR1, mitochondrial |
| AT1G29395 | 0.90 | 3.86E-04 | 0.69 | 2.26E-07 | -0.23 | 4.83E-01 | At1g29390 | Uncharacterized protein |
| AT4G15450 | 0.89 | 1.38E-02 | 0.92 | 8.58E-04 | 0.01 | 1.00E+00 | DL3768W | At4g15450 |
| AT4G27440 | 0.88 | 3.13E-02 | 0.38 | 3.57E-01 | -0.51 | 4.32E-01 | PORB | protochlorophyllide oxidoreductase B |
| AT1G28960 | 0.88 | 8.93E-03 | 0.73 | 8.80E-03 | -0.17 | 7.43E-01 | At1g28960 | Nudix hydrolase domain-containing protein |
| AT5G06270 | 0.88 | 4.17E-02 | 0.29 | 5.94E-01 | -0.60 | 1.16E-01 | GIR1 | Protein GL2-INTERACTING REPRESSOR 1 |
| AT5G48360 | 0.88 | 2.13E-02 | 0.12 | 8.28E-01 | -0.77 | 3.22E-02 | FH9 | Formin-like protein 9 |
| AT1G33170 | 0.88 | 2.64E-02 | 0.38 | 1.42E-01 | -0.51 | 1.77E-01 | At1g33170 | Methyltransferase |
| AT3G15520 | 0.87 | 3.42E-02 | -0.23 | 3.53E-01 | -1.11 | 2.28E-03 | At3g15520 | Cyclophilin-like peptidyl-prolyl cis-trans isomerase family protein |
| AT1G78450 | 0.87 | 1.62E-02 | 0.87 | 2.49E-04 | -0.02 | 9.90E-01 | At1g78450 | Uncharacterized protein At1g78450 |
| AT1G72970 | 0.87 | 2.21E-02 | -0.58 | 2.37E-03 | -1.47 | 8.45E-07 | At1g72970 | Uncharacterized protein At1g72970 |
| AT4G13560 | 0.87 | 2.19E-02 | -0.32 | 2.62E-01 | -1.20 | 6.88E-06 | UNE15 | Late embryogenesis abundant protein |
| AT4G12730 | 0.85 | 4.14E-02 | -0.57 | 2.54E-02 | -1.44 | 4.90E-09 | FLA2 | Fasciclin-like arabinogalactan protein 2 |
| AT1G68220 | 0.85 | 2.84E-02 | 0.98 | 3.55E-03 | 0.11 | 8.86E-01 | At1g68220 | |
| AT4G22200 | 0.85 | 2.07E-02 | 0.00 | 1.00E+00 | -0.86 | 7.22E-03 | KT2/3 | Potassium channel |
| AT4G01410 | 0.85 | 3.00E-02 | 0.15 | 7.41E-01 | -0.72 | 3.94E-02 | At4g01410 | At4g01410 |
| AT5G38865 | 0.84 | 3.40E-02 | 0.40 | 9.77E-02 | -0.46 | 2.29E-01 | | |
| AT4G33960 | 0.84 | 1.20E-02 | 0.41 | 3.09E-01 | -0.44 | 3.64E-01 | At4g33960 | Transmembrane protein |
| AT5G01520 | 0.83 | 9.45E-03 | 0.30 | 4.02E-01 | -0.55 | 1.88E-01 | AIRP2 | E3 ubiquitin-protein ligase AIRP2 |
| AT1G55260 | 0.83 | 5.18E-03 | -0.25 | 2.62E-01 | -1.10 | 6.61E-05 | LTPG6 | Bifunctional inhibitor/lipid-transfer protein/seed storage 2S albumin superfamily protein |
| AT5G23210 | 0.83 | 2.61E-02 | 0.43 | 3.13E-02 | -0.42 | 2.61E-01 | SCPL34 | Carboxypeptidase |
| AT4G10150 | 0.82 | 4.17E-02 | 0.13 | 8.40E-01 | -0.71 | 8.20E-02 | ATL7 | RING-H2 finger protein ATL7 |
| AT5G17520 | 0.82 | 1.75E-02 | 0.57 | 5.57E-03 | -0.27 | 5.40E-01 | RCP1 | Root cap 1 |
| AT3G50440 | 0.82 | 4.15E-03 | 0.64 | 6.87E-04 | -0.19 | 6.42E-01 | At3g50440 | Methylesterase |
| AT4G37590 | 0.82 | 4.44E-02 | 0.25 | 3.36E-01 | -0.58 | 1.64E-01 | At4g37590 | BTB/POZ domain-containing protein NPY5 |
| AT1G22160 | 0.82 | 1.84E-02 | 0.20 | 5.40E-01 | -0.63 | 8.00E-02 | FLZ5 | FCS-Like Zinc finger 5 |
| AT2G42320 | 0.81 | 3.98E-02 | 0.30 | 1.84E-01 | -0.52 | 1.76E-01 | MHK10.4 | Nucleolar protein gar2-like protein |
| AT4G24350 | 0.81 | 2.70E-02 | 0.42 | 4.74E-02 | -0.40 | 3.30E-01 | T22A6.180 | Uncharacterized protein AT4g24350 |
| AT3G02570 | 0.81 | 4.12E-02 | -0.04 | 9.58E-01 | -0.86 | 1.26E-02 | PMI1 | Mannose-6-phosphate isomerase 1 |
| AT5G48960 | 0.81 | 1.23E-02 | 0.78 | 6.89E-04 | -0.04 | 9.59E-01 | At5g48960 | HAD-superfamily hydrolase, subfamily IG, 5'-nucleotidase |
| AT2G21660 | 0.80 | 1.27E-02 | 1.21 | 4.13E-16 | 0.39 | 2.12E-01 | CCR2 | Cold, circadian rhythm, and rna binding 2 |
| AT5G50450 | 0.80 | 4.50E-02 | 1.03 | 1.42E-03 | 0.21 | 7.03E-01 | At5g50450 | F-box protein At5g50450 |
| AT2G28930 | 0.80 | 2.04E-02 | 0.45 | 5.02E-02 | -0.36 | 3.42E-01 | At2g28930 | Uncharacterized protein At2g28930 |
| AT5G46570 | 0.80 | 4.32E-02 | -0.16 | 7.02E-01 | -0.97 | 5.09E-03 | BSK2 | Serine/threonine-protein kinase BSK2 |
| AT1G78100 | 0.79 | 8.95E-03 | 0.00 | 1.00E+00 | -0.80 | 4.39E-03 | AUF1 | F-box protein AUF1 |
| AT4G38770 | 0.79 | 2.09E-02 | 0.15 | 5.35E-01 | -0.66 | 3.61E-02 | At4g38770 | Proline-rich protein 4 |
| AT1G79440 | 0.79 | 2.76E-02 | 0.68 | 3.77E-04 | -0.12 | 8.11E-01 | ALDH5F1 | Succinate-semialdehyde dehydrogenase, mitochondrial |
| AT5G38870 | 0.78 | 3.24E-02 | 0.39 | 3.18E-02 | -0.40 | 2.67E-01 | | |
| AT1G03930 | 0.78 | 4.86E-02 | 0.00 | 1.00E+00 | -0.79 | 3.61E-02 | CKL9 | Casein kinase 1-like protein 9 |
| AT2G43200 | 0.76 | 6.95E-03 | 0.27 | 2.16E-01 | -0.51 | 8.10E-02 | At2g43200 | Probable methyltransferase PMT19 |
| AT2G15070 | 0.76 | 4.86E-02 | 0.33 | 4.07E-01 | -0.43 | 3.36E-01 | | |
| AT2G35620 | 0.75 | 1.81E-02 | 0.28 | 2.58E-01 | -0.49 | 1.29E-01 | FEI2 | LRR receptor-like serine/threonine-protein kinase FEI 2 |
| AT1G47900 | 0.75 | 4.17E-02 | 0.25 | 4.24E-01 | -0.51 | 1.77E-01 | At1g47900 | Filament-like protein |
| AT5G39320 | 0.75 | 4.17E-02 | -0.24 | 5.28E-01 | -1.00 | 7.36E-04 | UGD4 | UDP-glucose 6-dehydrogenase 4 |
| AT5G47370 | 0.75 | 4.53E-02 | 0.39 | 3.00E-01 | -0.37 | 4.43E-01 | HAT2 | Homeobox-leucine zipper protein HAT2 |
| AT3G17860 | 0.74 | 2.45E-02 | 0.53 | 2.38E-03 | -0.23 | 5.77E-01 | JAZ3 | Protein TIFY |
| AT5G15740 | 0.74 | 2.60E-02 | -0.48 | 4.15E-02 | -1.23 | 4.54E-06 | At5g15740 | O-fucosyltransferase family protein |
| AT4G36360 | 0.74 | 3.44E-02 | -0.16 | 6.73E-01 | -0.92 | 2.38E-04 | At4g36360 | Beta-galactosidase like protein |
| AT1G78680 | 0.73 | 2.61E-02 | 0.19 | 5.48E-01 | -0.56 | 4.33E-02 | GGH2 | folate gamma-glutamyl hydrolase |
| AT3G05030 | 0.73 | 4.67E-02 | 0.16 | 6.49E-01 | -0.58 | 7.32E-02 | NHX2 | Sodium hydrogen exchanger 2 |
| AT1G49500 | 0.73 | 3.00E-02 | 0.57 | 3.38E-04 | -0.18 | 6.33E-01 | At1g49500 | Uncharacterized protein |
| AT1G79460 | 0.73 | 2.96E-02 | 0.09 | 8.15E-01 | -0.65 | 4.62E-02 | GA2 | Ent-kaur-16-ene synthase, chloroplastic |
| AT2G36870 | 0.73 | 1.30E-02 | 0.10 | 8.25E-01 | -0.65 | 1.97E-02 | XTH32 | Xyloglucan endotransglucosylase/hydrolase 32 |
| AT2G06850 | 0.73 | 4.17E-02 | 0.02 | 9.87E-01 | -0.72 | 1.14E-02 | XTH4 | Xyloglucan endotransglucosylase/hydrolase |
| AT5G62350 | 0.71 | 3.37E-02 | 0.70 | 3.89E-08 | -0.03 | 9.64E-01 | At5g62350 | |
| AT4G14560 | 0.71 | 3.46E-02 | 0.31 | 3.44E-01 | -0.41 | 2.48E-01 | IAA1 | Auxin-responsive protein IAA1 |
| AT5G61130 | 0.70 | 4.88E-02 | -0.13 | 8.16E-01 | -0.84 | 9.15E-03 | At5g61130 | PDCB1 |
| AT1G10870 | 0.70 | 4.67E-02 | -0.01 | 1.00E+00 | -0.72 | 1.84E-02 | AGD4 | ADP-ribosylation factor GTPase-activating protein |
| AT1G29390 | 0.68 | 3.02E-02 | 0.01 | 9.96E-01 | -0.68 | 2.08E-02 | At1g29390 | Uncharacterized protein |
| AT4G19120 | 0.67 | 4.89E-02 | 0.33 | 1.88E-01 | -0.36 | 2.25E-01 | ERD3 | Methyltransferase |

| | | | | | | | | |
|---|---|---|---|---|---|---|---|---|
| AT1G72275 | 0.67 | 4.49E-02 | 0.46 | 1.30E-01 | -0.23 | 6.06E-01 | T9N14.23 | Uncharacterized protein T9N14.23 |
| AT3G52180 | 0.66 | 4.50E-02 | 0.38 | 1.01E-01 | -0.29 | 4.65E-01 | At3g52180 | Uncharacterized protein At3g52180 |
| AT1G13250 | 0.65 | 3.93E-02 | 0.17 | 6.09E-01 | -0.50 | 6.89E-02 | GATL3 | Probable galacturonosyltransferase-like 3 |
| AT5G24160 | 0.63 | 4.16E-02 | 0.46 | 6.26E-03 | -0.18 | 6.19E-01 | At5g24160 | Squalene monooxygenase |
| AT4G12010 | -0.64 | 4.80E-02 | -0.43 | 3.61E-02 | 0.20 | 5.87E-01 | DSC1 | Disease resistance-like protein DSC1 |
| AT1G60490 | -0.65 | 3.78E-02 | -0.21 | 9.99E-01 | 0.44 | 3.88E-01 | VPS34 | Phosphatidylinositol 3-kinase VPS34 |
| AT3G61860 | -0.66 | 3.60E-02 | 0.04 | 9.70E-01 | 0.68 | 2.94E-02 | RS31 | Serine/arginine-rich splicing factor RS31 |
| AT1G67550 | -0.67 | 4.68E-02 | -0.20 | 9.99E-01 | 0.45 | 4.08E-01 | URE | Urease |
| AT5G04920 | -0.68 | 4.42E-02 | -0.36 | 1.64E-01 | 0.30 | 4.14E-01 | VPS36 | Vacuolar protein sorting-associated protein 36 |
| AT1G37130 | -0.68 | 4.74E-02 | -0.07 | 9.99E-01 | 0.59 | 1.64E-01 | NIA2 | Nitrate reductase 2 |
| AT1G74170 | -0.68 | 4.01E-02 | -0.46 | 3.64E-02 | 0.21 | 6.28E-01 | RLP13 | Receptor like protein 13 |
| AT1G20100 | -0.68 | 3.02E-02 | -0.25 | 4.18E-01 | 0.42 | 1.79E-01 | At1g20100 | DNA ligase-like protein |
| AT4G25940 | -0.68 | 3.14E-02 | -0.10 | 7.90E-01 | 0.57 | 5.01E-02 | At4g25940 | Putative clathrin assembly protein At4g25940 |
| AT2G39840 | -0.69 | 2.64E-02 | -0.02 | 1.00E+00 | 0.66 | 2.41E-02 | TOPP4 | Serine/threonine-protein phosphatase PP1 isozyme 4 |
| AT1G79380 | -0.70 | 3.27E-02 | -0.33 | 4.09E-02 | 0.36 | 2.98E-01 | RGLG4 | E3 ubiquitin-protein ligase RGLG4 |
| AT1G78920 | -0.70 | 2.49E-02 | -0.15 | 6.11E-01 | 0.54 | 8.72E-02 | AVPL1 | Pyrophosphate-energized membrane proton pump 2 |
| AT3G11280 | -0.71 | 2.86E-02 | 0.33 | 3.51E-01 | 1.02 | 8.62E-04 | F11B9.1 | Duplicated homeodomain-like superfamily protein |
| AT1G50380 | -0.71 | 3.85E-02 | -0.18 | 6.63E-01 | 0.51 | 1.39E-01 | At1g50380 | Prolyl endopeptidase |
| AT4G31210 | -0.71 | 4.20E-02 | -0.82 | 1.86E-04 | -0.13 | 7.63E-01 | At4g31210 | DNA topoisomerase |
| AT1G77760 | -0.71 | 4.51E-02 | -0.74 | 1.06E-05 | -0.04 | 9.36E-01 | NIA1 | Nitrate reductase [NADH] 1 |
| AT3G26180 | -0.72 | 4.97E-02 | -0.05 | 9.20E-01 | 0.65 | 5.20E-02 | CYP71B20 | Cytochrome P450, family 71, subfamily B, polypeptide 20 |
| AT3G12360 | -0.72 | 3.17E-02 | -0.16 | 5.88E-01 | 0.55 | 1.02E-01 | ITN1 | Ankyrin repeat family protein |
| AT3G55260 | -0.72 | 2.95E-02 | -0.23 | 4.11E-01 | 0.48 | 1.58E-01 | HEXO1 | Beta-hexosaminidase 1 |
| AT2G44500 | -0.72 | 4.86E-02 | 0.47 | 4.28E-02 | 1.18 | 5.23E-07 | At2g44500 | O-fucosyltransferase family protein |
| AT2G41220 | -0.72 | 4.42E-02 | 0.06 | 9.28E-01 | 0.76 | 2.40E-02 | GLU2 | Ferredoxin-dependent glutamate synthase 2, chloroplastic |
| AT1G10920 | -0.72 | 4.92E-02 | -0.37 | 2.46E-01 | 0.34 | 3.68E-01 | LOV1 | NB-ARC domain-containing disease resistance protein |
| AT3G59080 | -0.72 | 4.10E-02 | -0.17 | 7.12E-01 | 0.54 | 1.31E-01 | F17J16_130 | AT3g59080/F17J16_130 |
| AT2G14740 | -0.73 | 2.65E-02 | -0.23 | 4.22E-01 | 0.48 | 1.71E-01 | VSR3 | Vacuolar-sorting receptor 3 |
| AT5G62610 | -0.73 | 3.84E-02 | -0.18 | 6.32E-01 | 0.54 | 1.45E-01 | BHLH79 | Transcription factor bHLH79 |
| AT5G57580 | -0.73 | 2.71E-02 | -0.17 | 5.34E-01 | 0.54 | 1.02E-01 | CBP60B | Calmodulin-binding protein 60 B |
| AT1G03740 | -0.73 | 4.29E-02 | 0.13 | 6.85E-01 | 0.85 | 5.58E-03 | At1g03740 | F21B7.34 |
| AT3G52240 | -0.73 | 1.41E-02 | -0.32 | 2.31E-01 | 0.40 | 2.43E-01 | At3g52240 | Uncharacterized protein |
| AT5G09450 | -0.73 | 4.74E-02 | -0.66 | 6.38E-02 | 0.06 | 9.25E-01 | At5g09450 | Pentatricopeptide repeat-containing protein At5g09450, mitochondrial |
| AT2G29400 | -0.74 | 1.35E-02 | -0.08 | 8.82E-01 | 0.65 | 2.71E-02 | TOPP1 | Serine/threonine-protein phosphatase PP1 isozyme 1 |
| AT1G78290 | -0.74 | 4.40E-02 | -0.20 | 6.31E-01 | 0.52 | 1.69E-01 | SRK2C | Serine/threonine-protein kinase SRK2C |
| AT3G18370 | -0.75 | 2.13E-02 | -0.11 | 7.82E-01 | 0.62 | 5.68E-02 | ATSYTF | AT3g18370/MYF24_8 |
| AT4G00330 | -0.75 | 4.17E-02 | -0.17 | 5.10E-01 | 0.57 | 1.11E-01 | CRCK2 | Calmodulin-binding receptor-like cytoplasmic kinase 2 |
| AT4G02370 | -0.76 | 4.49E-02 | -0.10 | 8.11E-01 | 0.64 | 7.34E-02 | At4g02370 | AT4g02370 protein |
| AT5G44580 | -0.76 | 2.13E-02 | 0.07 | 8.72E-01 | 0.82 | 4.00E-03 | PROSCOOP1( | Serine rich endogenous peptide 10 |
| AT1G75020 | -0.76 | 3.64E-02 | -0.21 | 5.96E-01 | 0.53 | 1.66E-01 | LPAT4 | Probable 1-acyl-sn-glycerol-3-phosphate acyltransferase 4 |
| AT1G54115 | -0.76 | 1.94E-02 | 0.09 | 9.00E-01 | 0.84 | 4.04E-03 | CCX4 | Cation/calcium exchanger 4 |
| AT1G09430 | -0.77 | 4.80E-03 | -0.14 | 6.68E-01 | 0.61 | 2.42E-02 | ACLA-3 | ATP-citrate synthase alpha chain protein 3 |
| AT4G10970 | -0.77 | 2.26E-02 | -0.19 | 6.61E-01 | 0.57 | 8.54E-02 | UIEF1 | Ribosome maturation factor |
| AT1G12990 | -0.78 | 1.23E-02 | -0.15 | 6.41E-01 | 0.61 | 4.12E-02 | F3F19.2 | F3F19.2 protein |
| AT5G56260 | -0.78 | 1.57E-02 | -0.23 | 4.56E-01 | 0.53 | 1.15E-01 | At5g56260 | 4-hydroxy-4-methyl-2-oxoglutarate aldolase |
| AT1G71880 | -0.78 | 3.40E-02 | -0.05 | 9.42E-01 | 0.72 | 2.35E-02 | SUC1 | Sucrose transport protein SUC1 |
| AT1G69440 | -0.78 | 4.58E-02 | -0.17 | 6.23E-01 | 0.60 | 1.31E-01 | AGO7 | Protein argonaute 7 |
| AT3G52800 | -0.78 | 4.31E-02 | 0.05 | 9.25E-01 | 0.82 | 1.57E-02 | SAP6 | Zinc finger A20 and AN1 domain-containing stress-associated protein 6 |
| AT1G75130 | -0.78 | 3.21E-02 | -0.04 | 9.73E-01 | 0.73 | 2.74E-02 | CYP721A1 | Cytochrome P450, family 721, subfamily A, polypeptide 1 |
| AT4G23260 | -0.79 | 1.39E-02 | 0.22 | 4.76E-01 | 0.99 | 5.39E-05 | CRK18 | Cysteine-rich RLK |
| AT3G13810 | -0.79 | 1.53E-02 | -0.23 | 3.43E-01 | 0.54 | 9.88E-02 | IDD11 | Indeterminate |
| AT1G04530 | -0.79 | 6.95E-03 | -0.70 | 1.16E-06 | 0.08 | 8.55E-01 | TPR4 | Tetratricopeptide repeat |
| AT2G40940 | -0.79 | 3.50E-02 | -0.20 | 5.31E-01 | 0.57 | 1.23E-01 | ERS1 | Ethylene response sensor 1 |
| AT3G15430 | -0.79 | 4.33E-02 | -0.14 | 7.87E-01 | 0.64 | 7.75E-02 | At3g15430 | Regulator of chromosome condensation |
| AT2G17200 | -0.79 | 4.58E-02 | -0.07 | 8.88E-01 | 0.71 | 6.15E-02 | At2g17200 | DSK2b |
| AT5G56660 | -0.79 | 1.81E-02 | -0.10 | 8.78E-01 | 0.68 | 4.24E-02 | ILL2 | IAA-amino acid hydrolase ILR1-like 2 |
| AT2G01670 | -0.79 | 2.45E-02 | -0.12 | 7.62E-01 | 0.66 | 6.13E-02 | NUDT17 | Nudix hydrolase 17, mitochondrial |
| AT1G28280 | -0.79 | 3.75E-02 | -0.14 | 6.24E-01 | 0.63 | 8.34E-02 | MVQ1 | VQ motif-containing protein |
| AT1G61140 | -0.80 | 2.38E-02 | -0.18 | 5.32E-01 | 0.60 | 9.18E-02 | EDA16 | SNF2 domain-containing protein / helicase domain-containing protein / zinc finger protein-like protein |
| AT3G19190 | -0.80 | 1.53E-02 | -0.02 | 9.76E-01 | 0.76 | 1.35E-02 | ATG2 | Autophagy-related protein 2 |
| AT1G17420 | -0.80 | 3.29E-02 | 0.00 | 1.00E+00 | 0.79 | 1.93E-02 | At1g17420 | Lipoxygenase |
| AT1G05630 | -0.80 | 2.72E-02 | -0.63 | 2.50E-04 | 0.16 | 7.18E-01 | 5PTASE13 | Endonuclease/exonuclease/phosphatase family protein |
| AT3G46510 | -0.80 | 4.18E-02 | 0.11 | 7.25E-01 | 0.90 | 9.74E-03 | PUB13 | U-box domain-containing protein 13 |
| AT1G10050 | -0.81 | 2.19E-02 | -0.16 | 7.03E-01 | 0.62 | 8.28E-02 | ATXYN2 | Glycosyl hydrolase family 10 protein / carbohydrate-binding domain-containing protein |
| AT5G45410 | -0.81 | 2.24E-02 | -0.26 | 5.64E-01 | 0.53 | 2.31E-01 | ATNHR2A | AT5G45410 protein |
| AT5G11030 | -0.81 | 1.32E-02 | -0.33 | 4.55E-01 | 0.47 | 2.34E-01 | ALF4 | Aberrant root formation protein |
| AT5G51830 | -0.81 | 4.49E-02 | -0.26 | 6.90E-01 | 0.53 | 2.82E-01 | At5g51830 | Probable fructokinase-7 |
| AT3G07090 | -0.81 | 2.93E-02 | -0.28 | 5.30E-01 | 0.52 | 1.23E-01 | T1B9.26 | PPPDE putative thiol peptidase family protein |
| AT5G61530 | -0.81 | 1.90E-02 | 0.13 | 7.73E-01 | 0.93 | 7.46E-03 | At5g61530 | Small G protein family protein / RhoGAP family protein |
| AT4G34460 | -0.81 | 2.86E-02 | -0.02 | 1.00E+00 | 0.77 | 5.35E-02 | At4g34460 | At4g34460 |
| AT1G07650 | -0.81 | 4.40E-02 | -0.12 | 7.16E-01 | 0.68 | 6.94E-02 | At1g07650 | Uncharacterized protein At1g07655 |
| AT4G15260 | -0.82 | 4.37E-02 | -0.29 | 5.28E-01 | 0.50 | 2.46E-01 | DL3675W | At4g15260 |
| AT5G66675 | -0.82 | 4.58E-02 | -0.03 | 9.89E-01 | 0.77 | 1.61E-02 | At5g66675 | Uncharacterized protein |
| AT3G06530 | -0.82 | 8.92E-03 | -0.73 | 4.79E-05 | 0.07 | 8.90E-01 | At3g06530 | At3g06530 |
| AT3G49060 | -0.82 | 1.75E-02 | -0.24 | 3.74E-01 | 0.56 | 1.23E-01 | At3g49060 | U-box domain-containing protein kinase family protein |
| AT1G56520 | -0.82 | 2.76E-02 | -0.17 | 6.21E-01 | 0.64 | 7.77E-02 | F13N6.13 | Disease resistance protein |
| AT1G28380 | -0.82 | 1.53E-02 | -0.14 | 6.86E-01 | 0.67 | 5.69E-02 | NSL1 | MACPF domain-containing protein NSL1 |
| AT1G56460 | -0.83 | 3.17E-02 | 0.01 | 9.97E-01 | 0.82 | 2.21E-02 | At1g56460 | HIT zinc finger and PAPA-1-like domain-containing protein |
| AT5G01980 | -0.83 | 3.70E-02 | -0.12 | 8.77E-01 | 0.69 | 5.36E-02 | At5g01980 | |

| | | | | | | | | |
|---|---|---|---|---|---|---|---|---|
| AT4G27740 | -0.83 | 3.30E-02 | 0.24 | 7.25E-01 | 1.06 | 1.10E-02 | At4g27740 | Protein yippee-like At4g27740 |
| AT1G68585 | -0.83 | 4.53E-02 | -0.49 | 4.14E-01 | 0.33 | 5.59E-01 | At1g68585 | |
| AT2G40830 | -0.83 | 1.46E-02 | -0.44 | 4.33E-02 | 0.38 | 3.60E-01 | RHC1A | Probable E3 ubiquitin-protein ligase RHC1A |
| AT3G55880 | -0.83 | 3.85E-02 | -0.14 | 7.87E-01 | 0.67 | 7.30E-02 | SUE4 | Alpha/beta hydrolase related protein |
| AT3G17410 | -0.83 | 2.73E-02 | -0.01 | 1.00E+00 | 0.81 | 1.80E-02 | CARK1 | Receptor-like cytoplasmic kinase 1 |
| AT5G23520 | -0.83 | 1.81E-02 | -0.60 | 1.30E-02 | 0.22 | 6.10E-01 | At5g23520 | Smr domain-containing protein |
| AT1G54360 | -0.84 | 6.47E-03 | -0.25 | 3.83E-01 | 0.57 | 7.66E-02 | TAF6B | TBP-ASSOCIATED FACTOR 6B |
| AT5G18750 | -0.84 | 3.50E-02 | -0.19 | 7.77E-01 | 0.63 | 1.43E-01 | At5g18750 | J domain-containing protein |
| AT5G37480 | -0.84 | 2.63E-02 | -0.27 | 6.12E-01 | 0.56 | 1.93E-01 | At5g37480 | Transmembrane protein |
| AT3G27610 | -0.84 | 2.30E-02 | -0.65 | 3.37E-02 | 0.18 | 6.88E-01 | At3g27610 | Nucleotidylyl transferase superfamily protein |
| AT2G17420 | -0.84 | 2.29E-02 | -0.15 | 7.52E-01 | 0.68 | 8.86E-02 | NTRA | Thioredoxin reductase |
| AT5G46260 | -0.85 | 2.65E-02 | -0.05 | 9.65E-01 | 0.78 | 1.65E-02 | MPL12.4 | ADP-ribosyl cyclase/cyclic ADP-ribose hydrolase |
| AT1G52290 | -0.85 | 1.66E-02 | -0.45 | 6.17E-03 | 0.38 | 3.16E-01 | PERK15 | non-specific serine/threonine protein kinase |
| AT5G55130 | -0.85 | 2.70E-02 | 0.01 | 1.00E+00 | 0.85 | 2.94E-02 | CNX5 | Co-factor for nitrate, reductase and xanthine dehydrogenase 5 |
| AT4G03080 | -0.85 | 5.47E-03 | -0.25 | 2.58E-01 | 0.59 | 5.69E-02 | BSL1 | Serine/threonine-protein phosphatase BSL1 |
| AT3G10640 | -0.85 | 4.32E-02 | -0.16 | 7.03E-01 | 0.68 | 1.16E-01 | VPS601 | SNF7 family protein |
| AT3G23170 | -0.85 | 2.23E-02 | 0.39 | 5.21E-01 | 1.23 | 4.07E-04 | PRP | At3g23170 |
| AT2G24360 | -0.86 | 3.73E-02 | -0.07 | 8.77E-01 | 0.78 | 4.97E-02 | STY13 | Serine/threonine-protein kinase STY13 |
| AT2G35810 | -0.86 | 4.39E-02 | -0.13 | 8.68E-01 | 0.71 | 9.16E-02 | At2g35810 | Expressed protein |
| AT5G46490 | -0.86 | 1.85E-02 | -0.08 | 8.72E-01 | 0.76 | 1.84E-02 | At5g46490 | Disease resistance protein |
| AT3G51920 | -0.86 | 3.41E-03 | -0.28 | 3.14E-01 | 0.56 | 7.41E-02 | CML9 | Calmodulin-like protein 9 |
| AT5G39030 | -0.86 | 2.38E-02 | 0.03 | 9.73E-01 | 0.87 | 1.44E-02 | MDS4 | Protein kinase superfamily protein |
| AT5G50400 | -0.86 | 4.77E-03 | 0.20 | 5.15E-01 | 1.04 | 7.33E-05 | PAP27 | Purple acid phosphatase |
| AT4G23850 | -0.86 | 3.02E-02 | -0.11 | 7.38E-01 | 0.74 | 5.08E-02 | LACS4 | Long chain acyl-CoA synthetase 4 |
| AT3G60260 | -0.86 | 4.42E-02 | -0.22 | 7.36E-01 | 0.62 | 1.79E-01 | At3g60260 | |
| AT4G11900 | -0.86 | 1.67E-02 | -0.40 | 7.06E-02 | 0.45 | 2.12E-01 | At4g11900 | Receptor-like serine/threonine-protein kinase |
| AT1G14790 | -0.86 | 4.70E-03 | -0.07 | 8.71E-01 | 0.78 | 8.41E-03 | RDR1 | RNA-dependent RNA polymerase 1 |
| AT2G40600 | -0.86 | 2.64E-02 | -0.45 | 8.15E-02 | 0.41 | 3.05E-01 | At2g40600 | Expressed protein |
| AT3G49530 | -0.87 | 3.00E-02 | 0.21 | 6.06E-01 | 1.06 | 2.51E-03 | NAC062 | NAC domain containing protein 62 |
| AT2G44420 | -0.87 | 3.29E-02 | -0.31 | 6.13E-01 | 0.55 | 1.12E-01 | NTAN1 | Protein N-terminal asparagine amidohydrolase family protein |
| AT1G10590 | -0.87 | 2.90E-02 | -0.02 | 1.00E+00 | 0.84 | 2.48E-02 | At1g10590 | Nucleic acid-binding, OB-fold-like protein |
| AT1G80290 | -0.87 | 5.00E-02 | -0.16 | 7.43E-01 | 0.69 | 1.15E-01 | At1g80290 | Glycosyltransferase family protein 64 C3 |
| AT1G59620 | -0.87 | 1.43E-02 | -0.13 | 8.40E-01 | 0.73 | 1.65E-02 | CW9 | Disease resistance protein |
| AT5G43100 | -0.87 | 3.53E-02 | -0.04 | 9.55E-01 | 0.82 | 3.44E-02 | MMG4.12 | Eukaryotic aspartyl protease family protein |
| AT5G61060 | -0.87 | 9.19E-03 | -0.06 | 9.22E-01 | 0.80 | 1.45E-02 | HDA05 | Histone deacetylase 5 |
| AT1G26130 | -0.87 | 3.72E-02 | -0.10 | 8.76E-01 | 0.76 | 4.64E-02 | At1g26130 | Phospholipid-transporting ATPase |
| AT4G29900 | -0.87 | 7.68E-03 | -0.25 | 2.92E-01 | 0.61 | 5.42E-02 | At4g29900 | Calcium-transporting ATPase |
| AT2G44600 | -0.87 | 4.39E-02 | -0.90 | 2.13E-02 | -0.04 | 9.58E-01 | At2g44600 | |
| AT5G11560 | -0.88 | 2.91E-02 | -0.29 | 7.67E-02 | 0.57 | 1.68E-01 | At5g11560 | ER membrane protein complex subunit 1 |
| AT5G62570 | -0.88 | 4.56E-03 | -0.12 | 8.12E-01 | 0.74 | 9.89E-03 | CBP60A | Calmodulin binding protein-like protein |
| AT4G24040 | -0.88 | 2.02E-02 | -0.13 | 9.04E-01 | 0.74 | 6.41E-02 | At4g24040 | Trehalase |
| AT3G53270 | -0.88 | 8.52E-03 | -0.29 | 4.76E-01 | 0.57 | 1.26E-01 | At3g53270 | Small nuclear RNA activating complex |
| AT4G13810 | -0.88 | 2.77E-03 | -0.49 | 3.22E-03 | 0.37 | 2.66E-01 | RLP47 | Receptor like protein |
| AT2G36360 | -0.88 | 7.49E-03 | -0.31 | 4.59E-01 | 0.55 | 1.41E-01 | At2g36360 | AT2G36360 protein |
| AT5G58350 | -0.88 | 3.99E-02 | 0.11 | 8.57E-01 | 0.97 | 1.28E-02 | WNK4 | Probable serine/threonine-protein kinase WNK4 |
| AT5G22350 | -0.89 | 4.04E-02 | 0.04 | 9.61E-01 | 0.91 | 2.41E-02 | ELM1 | Mitochondrial fission protein ELM1 |
| AT1G55120 | -0.89 | 3.64E-02 | -0.05 | 9.65E-01 | 0.82 | 5.35E-02 | FRUCT5 | Beta-fructofuranosidase 5 |
| AT4G15070 | -0.89 | 4.82E-02 | -0.34 | 6.17E-01 | 0.53 | 3.04E-01 | dl3580c | Cysteine/Histidine-rich C1 domain family protein |
| AT2G21500 | -0.89 | 3.95E-02 | 0.23 | 5.49E-01 | 1.10 | 2.04E-03 | At2g21500 | Uncharacterized protein At2g21500 |
| AT4G08500 | -0.89 | 3.75E-02 | -0.08 | 9.26E-01 | 0.80 | 3.27E-02 | MEKK1 | Mitogen-activated protein kinase kinase kinase 1 |
| AT1G78620 | -0.89 | 2.30E-02 | -0.22 | 5.10E-01 | 0.66 | 1.13E-01 | VTE6 | Protein VTE6, chloroplastic |
| AT1G56510 | -0.89 | 1.51E-02 | -0.20 | 6.02E-01 | 0.68 | 1.63E-02 | ADR2 | Disease resistance protein ADR2 |
| AT4G28710 | -0.89 | 1.14E-02 | -0.19 | 4.11E-01 | 0.69 | 5.93E-02 | XI-H | Myosin-14 |
| AT4G29740 | -0.90 | 1.04E-02 | -0.68 | 4.45E-04 | 0.20 | 6.69E-01 | CKX4 | cytokinin dehydrogenase |
| AT1G30410 | -0.90 | 1.46E-03 | -0.47 | 2.61E-02 | 0.41 | 1.75E-01 | ABCC12 | ABC-type xenobiotic transporter |
| AT4G24940 | -0.90 | 2.30E-02 | -0.55 | 2.58E-01 | 0.33 | 4.84E-01 | SAE1A | SUMO-activating enzyme subunit 1A |
| AT4G16690 | -0.90 | 3.50E-02 | -1.11 | 1.39E-04 | -0.22 | 6.58E-01 | MES16 | pFDCC methylesterase MES16 |
| AT2G32520 | -0.90 | 1.15E-02 | -0.27 | 5.76E-01 | 0.62 | 4.23E-02 | At2g32520 | Alpha/beta-Hydrolases superfamily protein |
| AT1G76700 | -0.90 | 2.66E-02 | -0.14 | 8.26E-01 | 0.75 | 7.03E-02 | ATJ10 | Chaperone protein dnaJ 10 |
| AT3G27890 | -0.91 | 2.64E-02 | -0.29 | 2.52E-01 | 0.60 | 1.56E-01 | NQR | NADPH:quinone oxidoreductase |
| AT2G31870 | -0.91 | 5.75E-03 | -0.27 | 5.93E-01 | 0.62 | 6.11E-02 | TEJ | poly |
| AT4G33920 | -0.91 | 4.67E-02 | -0.07 | 9.59E-01 | 0.83 | 4.40E-02 | APD5 | Protein phosphatase 2C family protein |
| AT1G63350 | -0.91 | 1.57E-02 | 0.32 | 5.87E-01 | 1.22 | 1.22E-03 | At1g63350 | Disease resistance protein |
| AT1G07670 | -0.91 | 2.26E-02 | -0.18 | 5.39E-01 | 0.71 | 7.60E-02 | ECA4 | Calcium-transporting ATPase 4, endoplasmic reticulum-type |
| AT1G65540 | -0.91 | 3.70E-02 | -0.27 | 3.41E-01 | 0.63 | 1.54E-01 | F5I14.7 | F5I14.7 protein |
| AT3G29400 | -0.91 | 1.81E-02 | -0.13 | 7.48E-01 | 0.77 | 4.49E-02 | EXO70E1 | Exocyst subunit Exo70 family protein |
| AT1G22650 | -0.91 | 3.74E-02 | -0.42 | 9.03E-02 | 0.47 | 3.26E-01 | A/N-INVD | Alkaline/neutral invertase |
| AT5G20910 | -0.92 | 1.57E-02 | 0.08 | 9.51E-01 | 0.97 | 7.04E-03 | AIP2 | E3 ubiquitin-protein ligase AIP2 |
| AT2G27920 | -0.92 | 1.38E-02 | 0.27 | 6.44E-01 | 1.17 | 1.05E-03 | SCPL51 | Carboxypeptidase |
| AT3G07060 | -0.92 | 4.80E-02 | -0.60 | 3.53E-01 | 0.30 | 6.30E-01 | EMB1974 | NHL domain-containing protein |
| AT1G25390 | -0.92 | 1.04E-02 | -0.13 | 7.29E-01 | 0.77 | 2.99E-02 | LRK10L-1.1 | LEAF RUST 10 DISEASE-RESISTANCE LOCUS RECEPTOR-LIKE PROTEIN KINASE-like 1.1 |
| AT2G45030 | -0.92 | 4.67E-02 | -0.32 | 3.91E-01 | 0.59 | 2.03E-01 | MEFG2 | Elongation factor G-2, mitochondrial |
| AT5G42140 | -0.92 | 1.72E-02 | -0.21 | 6.97E-01 | 0.70 | 4.61E-02 | MJC20.25 | Regulator of chromosome condensation |
| AT3G20410 | -0.93 | 4.40E-02 | 0.22 | 6.75E-01 | 1.13 | 4.79E-03 | CPK9 | Calcium-dependent protein kinase 9 |
| AT3G14990 | -0.93 | 3.50E-02 | 0.26 | 6.92E-01 | 1.16 | 4.13E-04 | DJ1A | Protein DJ-1 homolog A |
| AT1G22070 | -0.93 | 2.77E-02 | 0.13 | 7.70E-01 | 1.04 | 9.39E-03 | At1g22070 | Uncharacterized protein |
| AT5G28830 | -0.93 | 2.61E-02 | 0.11 | 8.77E-01 | 1.03 | 1.25E-02 | At5g28830 | Calcium-binding EF hand family protein |
| AT1G63860 | -0.93 | 1.38E-02 | -0.16 | 6.90E-01 | 0.76 | 3.12E-02 | At1g63860 | Uncharacterized protein |

| | | | | | | | | |
|---|---|---|---|---|---|---|---|---|
| AT5G66070 | -0.93 | 2.40E-02 | 0.42 | 4.97E-01 | 1.34 | 2.49E-04 | ATARRE | RING/U-box superfamily protein |
| AT3G23530 | -0.93 | 1.54E-03 | -0.87 | 1.81E-09 | 0.05 | 9.24E-01 | At3g23530 | Amine oxidase domain-containing protein |
| AT1G66920 | -0.93 | 4.51E-03 | 0.87 | 1.18E-04 | 1.79 | 5.29E-09 | At1g66920 | Protein kinase superfamily protein |
| AT4G19990 | -0.93 | 8.88E-03 | -0.26 | 5.80E-01 | 0.66 | 7.52E-02 | FRS1 | Protein FAR1-RELATED SEQUENCE |
| AT5G24980 | -0.93 | 4.15E-02 | -0.56 | 1.79E-01 | 0.36 | 4.82E-01 | At5g24980 | |
| AT2G37940 | -0.93 | 4.46E-03 | -0.31 | 1.23E-01 | 0.61 | 8.09E-02 | IPCS2 | Phosphatidylinositol:ceramide inositolphosphotransferase 2 |
| AT1G34130 | -0.93 | 3.47E-02 | -0.45 | 2.41E-02 | 0.47 | 3.38E-01 | STT3B | Dolichyl-diphosphooligosaccharide--protein glycosyltransferase subunit STT3B |
| AT2G14720 | -0.93 | 4.23E-02 | -0.03 | 9.72E-01 | 0.89 | 4.74E-02 | VSR4 | Vacuolar-sorting receptor 4 |
| AT4G35310 | -0.94 | 2.70E-02 | -0.30 | 9.04E-02 | 0.62 | 1.47E-01 | CPK5 | Calcium-dependent protein kinase 5 |
| AT5G66210 | -0.94 | 2.51E-02 | 0.19 | 5.89E-01 | 1.11 | 1.57E-03 | At5g66210 | AT5G66210 protein |
| AT1G20780 | -0.94 | 1.17E-02 | -0.39 | 2.37E-01 | 0.53 | 1.60E-01 | PUB44 | U-box domain-containing protein 44 |
| AT4G11530 | -0.94 | 4.30E-02 | -0.62 | 9.41E-02 | 0.31 | 5.72E-01 | CRK34 | Cysteine-rich RLK |
| AT4G18950 | -0.94 | 6.45E-03 | -0.61 | 4.62E-02 | 0.32 | 3.49E-01 | BHP | AT4g18950/F13C5_120 |
| AT5G58220 | -0.94 | 1.98E-02 | -0.13 | 8.72E-01 | 0.80 | 5.72E-02 | TTL | Uric acid degradation bifunctional protein TTL |
| AT5G49760 | -0.94 | 1.46E-02 | -0.34 | 3.71E-02 | 0.59 | 1.37E-01 | HPCA1 | Leucine-rich repeat receptor protein kinase HPCA1 |
| AT1G35580 | -0.94 | 1.07E-02 | -0.04 | 9.45E-01 | 0.88 | 1.18E-02 | CINV1 | Alkaline/neutral invertase |
| AT1G22890 | -0.94 | 2.24E-03 | -0.02 | 9.94E-01 | 0.91 | 2.67E-03 | PROSCOOP14 | Serine rich endogenous peptide 14 |
| AT5G10610 | -0.94 | 3.16E-02 | -0.20 | 5.45E-01 | 0.72 | 9.94E-02 | CYP81K1 | Cytochrome P450, family 81, subfamily K, polypeptide 1 |
| AT4G31500 | -0.94 | 4.68E-02 | 0.65 | 8.27E-06 | 1.58 | 3.31E-05 | CYP83B1 | Cytochrome P450 83B1 |
| AT4G00500 | -0.95 | 6.65E-03 | -0.11 | 9.23E-01 | 0.82 | 2.94E-02 | At4g00500 | |
| AT4G08480 | -0.95 | 6.03E-03 | -0.42 | 8.28E-02 | 0.52 | 1.83E-01 | MEKK2 | Mitogen-activated protein kinase kinase kinase 9 |
| AT2G45500 | -0.95 | 1.38E-02 | -0.01 | 1.00E+00 | 0.93 | 9.86E-03 | At2g45500 | AAA-type ATPase family protein |
| AT1G27100 | -0.95 | 3.05E-03 | -0.01 | 1.00E+00 | 0.92 | 5.46E-04 | At1g27100 | T7N9.16 |
| AT1G74310 | -0.95 | 1.90E-02 | 0.12 | 9.33E-01 | 1.05 | 2.95E-02 | HSP101 | Heat shock protein 101 |
| AT2G17840 | -0.95 | 5.85E-03 | 0.62 | 2.92E-03 | 1.55 | 1.20E-07 | ERD7 | Senescence/dehydration-associated protein-like protein |
| AT4G25640 | -0.95 | 1.60E-02 | -0.23 | 3.64E-01 | 0.71 | 8.27E-02 | DTX35 | Protein DETOXIFICATION |
| AT1G52160 | -0.95 | 1.44E-02 | -0.52 | 1.27E-01 | 0.42 | 3.71E-01 | TRZ3 | tRNase Z TRZ3, mitochondrial |
| AT1G75510 | -0.95 | 1.86E-02 | -0.27 | 5.83E-01 | 0.67 | 1.59E-01 | At1g75510 | TFIIF beta subunit HTH domain-containing protein |
| AT3G60800 | -0.95 | 4.17E-02 | -0.09 | 8.90E-01 | 0.85 | 5.94E-02 | PAT14 | Probable protein S-acyltransferase 14 |
| AT5G35200 | -0.96 | 3.85E-02 | -0.17 | 5.14E-01 | 0.77 | 8.91E-02 | At5g35200 | Putative clathrin assembly protein At5g35200 |
| AT1G55820 | -0.96 | 4.59E-02 | -0.46 | 2.59E-01 | 0.47 | 2.90E-01 | GIP1L | GBF-interacting protein 1-like |
| AT5G47420 | -0.96 | 2.25E-02 | -0.11 | 9.07E-01 | 0.83 | 8.18E-02 | At5g47420 | Altered inheritance of mitochondria protein 24, mitochondrial |
| AT4G08390 | -0.96 | 4.03E-02 | -0.83 | 2.12E-07 | 0.11 | 8.61E-01 | SAPX | L-ascorbate peroxidase |
| AT1G17890 | -0.96 | 2.73E-02 | -0.17 | 7.72E-01 | 0.77 | 5.25E-02 | GER2 | GDP-L-fucose synthase |
| AT4G02405 | -0.97 | 4.61E-02 | -0.44 | 2.41E-01 | 0.51 | 3.61E-01 | At4g02405 | S-adenosyl-L-methionine-dependent methyltransferases superfamily protein |
| AT3G15355 | -0.97 | 5.27E-03 | 0.00 | 1.00E+00 | 0.95 | 3.71E-03 | UBC25 | Probable ubiquitin-conjugating enzyme E2 25 |
| AT2G31740 | -0.97 | 6.95E-03 | -0.32 | 2.69E-01 | 0.64 | 1.02E-01 | At2g31740 | Methyltransferase type 11 domain-containing protein |
| AT3G60190 | -0.97 | 4.01E-02 | -0.02 | 9.84E-01 | 0.94 | 3.86E-02 | At3g60190 | UMP-CMP kinase |
| AT1G17430 | -0.97 | 1.43E-02 | -0.82 | 7.40E-02 | 0.14 | 8.28E-01 | F28G4.9 | F28G4.9 protein |
| AT3G54850 | -0.97 | 4.99E-03 | -0.07 | 9.51E-01 | 0.88 | 9.22E-03 | At3g54850 | AT3G54850 protein |
| AT5G03290 | -0.97 | 4.61E-02 | -0.13 | 7.36E-01 | 0.83 | 8.95E-02 | IDH5 | Isocitrate dehydrogenase [NAD] catalytic subunit 5, mitochondrial |
| AT4G34480 | -0.97 | 2.71E-02 | -0.21 | 4.06E-01 | 0.75 | 8.09E-02 | At4g34480 | glucan endo-1,3-beta-D-glucosidase |
| AT4G01370 | -0.97 | 2.67E-02 | -0.47 | 4.73E-03 | 0.49 | 3.16E-01 | MPK4 | Mitogen-activated protein kinase 4 |
| AT3G47820 | -0.98 | 1.63E-02 | 0.01 | 1.00E+00 | 0.97 | 1.26E-02 | PUB39 | U-box domain-containing protein 39 |
| AT4G25515 | -0.98 | 2.35E-02 | 0.17 | 8.42E-01 | 1.13 | 1.12E-03 | SLK3 | Probable transcriptional regulator SLK3 |
| AT4G13040 | -0.98 | 4.20E-03 | -0.52 | 4.84E-02 | 0.45 | 2.04E-01 | APD1 | Integrase-type DNA-binding superfamily protein |
| AT4G17830 | -0.98 | 3.53E-02 | -0.17 | 7.43E-01 | 0.80 | 6.20E-02 | ATNAOD | Peptidase M20/M25/M40 family protein |
| AT5G25790 | -0.99 | 1.59E-02 | -0.16 | 9.06E-01 | 0.81 | 4.54E-02 | At5g25790 | Tesmin/TSO1-like CXC domain-containing protein |
| AT3G24180 | -0.99 | 7.48E-04 | -0.25 | 2.35E-01 | 0.73 | 1.55E-02 | At3g24180 | Non-lysosomal glucosylceramidase |
| AT4G21534 | -0.99 | 3.97E-02 | -0.41 | 3.38E-01 | 0.56 | 2.82E-01 | SPHK2 | Sphingosine kinase 2 |
| AT5G54910 | -0.99 | 2.65E-02 | -0.80 | 1.23E-01 | 0.17 | 8.02E-01 | RH32 | DEAD-box ATP-dependent RNA helicase 32 |
| AT1G32700 | -0.99 | 8.16E-03 | -0.15 | 6.78E-01 | 0.82 | 2.10E-02 | At1g32700 | F6N18.8 |
| AT4G39820 | -0.99 | 2.09E-02 | -0.12 | 8.77E-01 | 0.86 | 6.45E-02 | TRAPPC12 | At4g39820 |
| AT2G41010 | -0.99 | 2.37E-02 | -0.04 | 9.71E-01 | 0.94 | 2.24E-02 | CAMBP25 | Calmodulin-binding protein 25 |
| AT1G56540 | -1.00 | 2.99E-03 | -0.11 | 8.68E-01 | 0.87 | 9.79E-03 | At1g56540 | Disease resistance protein |
| AT5G15730 | -1.00 | 2.09E-02 | -0.18 | 7.25E-01 | 0.80 | 8.44E-02 | At5g15730 | Protein kinase domain-containing protein |
| AT1G51270 | -1.00 | 1.27E-02 | 0.49 | 2.56E-01 | 1.48 | 3.87E-04 | At1g51270 | Vesicle-associated protein 1-4 |
| AT4G29050 | -1.00 | 2.72E-02 | -0.23 | 5.22E-01 | 0.75 | 1.11E-01 | LECRK-V9 | non-specific serine/threonine protein kinase |
| AT2G35940 | -1.00 | 9.89E-03 | 0.12 | 7.87E-01 | 1.10 | 1.25E-03 | At2g35940 | Homeobox domain-containing protein |
| AT5G55070 | -1.00 | 4.66E-02 | -0.13 | 7.74E-01 | 0.86 | 8.27E-02 | At5g55070 | Dihydrolipoyllysine-residue succinyltransferase component<br>of 2-oxoglutarate dehydrogenase complex 1, mitochondrial |
| AT3G42790 | -1.00 | 3.14E-02 | -0.33 | 3.00E-01 | 0.65 | 1.63E-01 | AL3 | PHD finger protein ALFIN-LIKE 3 |
| AT5G27600 | -1.00 | 4.86E-02 | 0.44 | 2.32E-01 | 1.44 | 1.30E-03 | LACS7 | Long chain acyl-CoA synthetase 7, peroxisomal |
| AT3G04120 | -1.01 | 2.70E-02 | -0.25 | 1.73E-01 | 0.74 | 1.14E-01 | GAPC1 | Glyceraldehyde-3-phosphate dehydrogenase GAPC1, cytosolic |
| AT1G14170 | -1.01 | 1.09E-02 | -0.08 | 9.66E-01 | 0.90 | 4.61E-02 | At1g14170 | AT1G14170 protein |
| AT3G16700 | -1.01 | 4.71E-02 | -0.66 | 1.91E-01 | 0.33 | 5.75E-01 | At3g16700 | Fumarylacetoacetate |
| AT5G27920 | -1.01 | 5.46E-03 | -0.19 | 7.75E-01 | 0.80 | 2.18E-02 | At5g27920 | At5g27920 |
| AT4G23650 | -1.01 | 1.37E-02 | -0.11 | 8.15E-01 | 0.88 | 1.98E-02 | CPK3 | Calcium-dependent protein kinase 3 |
| AT2G23450 | -1.01 | 2.70E-02 | -0.17 | 7.21E-01 | 0.82 | 4.06E-02 | WAKL14 | Wall-associated receptor kinase-like 14 |
| AT1G71697 | -1.01 | 8.79E-03 | -0.24 | 6.14E-01 | 0.75 | 7.37E-02 | CK1 | Probable choline kinase 1 |
| AT4G02410 | -1.01 | 1.65E-03 | -0.33 | 9.46E-02 | 0.67 | 5.11E-02 | LECRK43 | L-type lectin-domain containing receptor kinase IV.3 |
| AT4G03820 | -1.01 | 4.01E-02 | -0.14 | 8.22E-01 | 0.85 | 6.75E-02 | At4g03820 | Transmembrane protein, putative |
| AT4G17615 | -1.01 | 1.67E-02 | -0.12 | 9.32E-01 | 0.88 | 5.65E-03 | CBL1 | Calcineurin B-like protein |
| AT3G07570 | -1.01 | 1.12E-02 | -0.13 | 8.23E-01 | 0.87 | 1.74E-02 | At3g07570 | Cytochrome b561 and DOMON domain-containing protein At3g07570 |
| AT3G07780 | -1.01 | 1.96E-02 | -0.14 | 7.29E-01 | 0.86 | 4.11E-02 | OBE1 | Protein OBERON |
| AT3G05970 | -1.02 | 1.25E-02 | -0.04 | 9.63E-01 | 0.97 | 1.42E-02 | LACS6 | Long chain acyl-CoA synthetase 6, peroxisomal |
| AT4G18010 | -1.02 | 6.57E-04 | 0.22 | 6.00E-01 | 1.23 | 1.36E-05 | IP5PII | Myo-inositol polyphosphate 5-phosphatase 2 |
| AT1G30270 | -1.02 | 4.66E-02 | 0.01 | 1.00E+00 | 1.02 | 4.35E-02 | CIPK23 | non-specific serine/threonine protein kinase |
| AT2G15042 | -1.03 | 1.04E-02 | -0.25 | 3.66E-01 | 0.77 | 4.27E-02 | RLP18 | Receptor-like protein 18 |

| | | | | | | | | |
|---|---|---|---|---|---|---|---|---|
| AT2G01180 | -1.03 | 6.11E-03 | -0.38 | 1.79E-01 | 0.64 | 1.05E-01 | PAP1 | Phosphatidic acid phosphatase 1 |
| AT1G16970 | -1.03 | 1.63E-02 | -0.51 | 1.13E-01 | 0.51 | 2.22E-01 | At1g16970 | KU70 |
| AT1G17080 | -1.03 | 1.04E-02 | -0.05 | 9.64E-01 | 0.97 | 1.60E-02 | At1g17080 | 60S ribosomal protein L18a-like protein |
| AT4G36210 | -1.04 | 2.63E-03 | -0.01 | 1.00E+00 | 1.01 | 2.63E-03 | At4g36210 | Transmembrane/coiled-coil protein |
| AT2G02800 | -1.04 | 5.65E-03 | -0.36 | 1.40E-01 | 0.66 | 9.12E-02 | PBL3 | Probable serine/threonine-protein kinase PBL3 |
| AT1G03400 | -1.04 | 1.33E-04 | -0.40 | 4.23E-02 | 0.62 | 2.48E-02 | At1g03400 | 2-oxoglutarate |
| AT2G12550 | -1.04 | 4.58E-02 | 0.26 | 8.26E-01 | 1.29 | 6.09E-03 | At2g12550 | Uncharacterized protein At2g12550 |
| AT5G65200 | -1.05 | 2.60E-02 | -0.63 | 6.24E-02 | 0.40 | 4.20E-01 | PUB38 | U-box domain-containing protein 38 |
| AT5G47730 | -1.05 | 4.56E-03 | -0.32 | 5.31E-01 | 0.71 | 7.44E-02 | MCA23.5 | AT5G47730 protein |
| AT2G17220 | -1.05 | 1.09E-02 | -0.34 | 8.25E-02 | 0.70 | 1.08E-01 | PIX13 | Probable serine/threonine-protein kinase PIX13 |
| AT5G47200 | -1.05 | 3.42E-02 | -0.17 | 7.57E-01 | 0.87 | 5.38E-02 | RABD2B | Ras-related protein RABD2b |
| AT5G16880 | -1.05 | 6.42E-03 | -0.17 | 5.17E-01 | 0.87 | 2.28E-02 | At5g16880 | Target of Myb protein 1 |
| AT1G64280 | -1.06 | 5.56E-04 | -0.42 | 1.15E-02 | 0.62 | 6.03E-02 | NPR1 | Regulatory protein NPR1 |
| AT1G19270 | -1.06 | 4.77E-03 | 0.05 | 9.58E-01 | 1.09 | 3.10E-03 | DA1 | Protein DA1 |
| AT1G63460 | -1.06 | 1.10E-02 | 0.00 | 1.00E+00 | 1.04 | 3.03E-03 | At1g63460 | Glutathione peroxidase |
| AT2G17790 | -1.06 | 1.83E-02 | -0.25 | 3.69E-01 | 0.79 | 9.47E-02 | VPS35A | Vacuolar protein sorting-associated protein 35A |
| AT3G14050 | -1.06 | 2.33E-02 | 0.17 | 6.33E-01 | 1.21 | 4.55E-03 | At3g14050 | Putative GTP pyrophosphokinase |
| AT1G71040 | -1.06 | 1.59E-02 | -0.14 | 5.93E-01 | 0.91 | 3.52E-02 | LPR2 | Multicopper oxidase LPR2 |
| AT2G40000 | -1.06 | 3.70E-02 | 0.48 | 3.25E-01 | 1.53 | 6.78E-09 | HSPRO2 | Nematode resistance protein-like HSPRO2 |
| AT2G16530 | -1.06 | 4.86E-02 | 0.35 | 6.11E-01 | 1.40 | 5.79E-03 | PPRD2 | Polyprenol reductase |
| AT5G04860 | -1.06 | 4.86E-02 | -0.42 | 5.84E-01 | 0.63 | 1.70E-01 | MUK11.18 | Splicing factor 3A subunit |
| AT4G23250 | -1.06 | 5.91E-03 | -0.56 | 1.73E-03 | 0.49 | 2.62E-01 | EMB1290 | Cysteine-rich receptor-like protein kinase 17 |
| AT4G19660 | -1.07 | 1.35E-03 | -0.32 | 8.72E-02 | 0.73 | 3.91E-02 | NPR4 | NPR1-like protein 4 |
| AT4G01920 | -1.07 | 3.80E-02 | -0.16 | 8.72E-01 | 0.88 | 2.04E-02 | At4g01920 | AT4g01920/T7B11_18 |
| AT2G28890 | -1.07 | 1.04E-02 | -0.31 | 2.42E-01 | 0.73 | 6.70E-02 | PLL4 | Probable protein phosphatase 2C 23 |
| AT5G05190 | -1.07 | 2.48E-02 | -0.16 | 7.77E-01 | 0.90 | 4.07E-02 | EDR4 | Protein ENHANCED DISEASE RESISTANCE 4 |
| AT5G19550 | -1.07 | 2.67E-03 | -0.01 | 1.00E+00 | 1.05 | 2.95E-03 | At5g19550 | Aspartate aminotransferase |
| AT3G47570 | -1.07 | 2.26E-03 | -0.27 | 4.81E-01 | 0.79 | 3.16E-02 | At3g47570 | Probable LRR receptor-like serine/threonine-protein kinase At3g47570 |
| AT1G02170 | -1.08 | 1.23E-02 | -0.26 | 4.11E-01 | 0.80 | 5.78E-02 | AMC1 | Metacaspase-1 |
| AT5G61390 | -1.08 | 2.32E-02 | -0.11 | 9.53E-01 | 0.95 | 4.79E-02 | NEN2 | Protein NEN2 |
| AT3G59660 | -1.08 | 1.38E-02 | 0.34 | 1.69E-01 | 1.40 | 4.56E-04 | BAGP1 | C2 domain-containing protein / GRAM domain-containing protein |
| AT2G17290 | -1.08 | 1.67E-02 | -0.14 | 8.19E-01 | 0.93 | 1.77E-02 | CPK6 | Calcium-dependent protein kinase 6 |
| AT2G39660 | -1.08 | 1.79E-02 | -0.02 | 1.00E+00 | 1.06 | 1.67E-02 | BIK1 | Serine/threonine-protein kinase BIK1 |
| AT1G54320 | -1.08 | 1.90E-02 | -0.05 | 9.46E-01 | 1.02 | 2.34E-02 | ALIS3 | ALA-interacting subunit 3 |
| AT5G19250 | -1.09 | 1.53E-02 | -0.46 | 2.66E-04 | 0.61 | 2.08E-01 | At5g19250 | Uncharacterized GPI-anchored protein At5g19250 |
| AT5G66850 | -1.09 | 4.80E-02 | -0.24 | 3.24E-01 | 0.83 | 1.33E-01 | MAPKKK5 | Mitogen-activated protein kinase kinase kinase 5 |
| AT3G57640 | -1.09 | 3.60E-02 | -0.72 | 2.37E-01 | 0.36 | 5.50E-01 | ZRK15 | Non-functional pseudokinase ZRK15 |
| AT5G56350 | -1.09 | 3.17E-02 | -0.15 | 6.76E-01 | 0.92 | 5.93E-02 | MCD7.8 | Pyruvate kinase |
| AT1G49750 | -1.09 | 6.82E-03 | -0.62 | 2.27E-07 | 0.46 | 3.23E-01 | At1g49750 | AT1G49750 protein |
| AT3G05660 | -1.09 | 2.34E-02 | 0.08 | 9.26E-01 | 1.16 | 9.86E-03 | At3g05660 | RLP33 |
| AT1G75750 | -1.10 | 2.85E-02 | -0.24 | 5.51E-01 | 0.84 | 5.36E-02 | GASA1 | GAST1 protein homolog 1 |
| AT4G01720 | -1.10 | 3.92E-02 | 0.33 | 9.04E-02 | 1.41 | 3.45E-03 | WRKY47 | Probable WRKY transcription factor 47 |
| AT5G61380 | -1.10 | 4.98E-03 | -0.13 | 8.75E-01 | 0.95 | 1.60E-02 | APRR1 | Two-component response regulator-like APRR1 |
| AT2G22480 | -1.10 | 3.64E-03 | -0.24 | 4.48E-01 | 0.85 | 2.34E-02 | PFK5 | ATP-dependent 6-phosphofructokinase 5, chloroplastic |
| AT1G69250 | -1.10 | 1.81E-02 | -0.41 | 4.41E-01 | 0.67 | 1.42E-01 | F23O10.17 | At1g69250 |
| AT2G07180 | -1.10 | 1.42E-03 | -0.01 | 1.00E+00 | 1.08 | 1.46E-03 | PBL17 | Probable serine/threonine-protein kinase PBL17 |
| AT2G11520 | -1.10 | 1.50E-02 | -0.25 | 3.64E-01 | 0.83 | 7.92E-02 | CRCK3 | Calmodulin-binding receptor-like cytoplasmic kinase 3 |
| AT5G08300 | -1.10 | 3.02E-02 | -0.02 | 9.87E-01 | 1.06 | 2.25E-02 | At5g08300 | Succinate--CoA ligase [ADP-forming] subunit alpha-1, mitochondrial |
| AT1G74280 | -1.10 | 7.39E-03 | -0.10 | 9.50E-01 | 0.98 | 1.53E-02 | At1g74280 | Alpha/beta-Hydrolases superfamily protein |
| AT3G01720 | -1.10 | 1.96E-02 | -0.12 | 7.87E-01 | 0.96 | 4.79E-02 | SERGT1 | Peptidyl serine alpha-galactosyltransferase |
| AT1G55450 | -1.10 | 1.67E-02 | -0.46 | 6.36E-02 | 0.63 | 1.36E-01 | At1g55450 | S-adenosyl-L-methionine-dependent methyltransferases superfamily protein |
| AT1G67470 | -1.10 | 8.92E-03 | -0.40 | 4.37E-01 | 0.68 | 7.02E-02 | ZRK12 | Inactive serine/threonine-protein kinase ZRK12 |
| AT3G47010 | -1.10 | 7.42E-03 | -0.25 | 7.42E-01 | 0.84 | 5.80E-02 | F13I12.60 | beta-glucosidase |
| AT4G26780 | -1.11 | 1.30E-02 | -0.68 | 2.47E-01 | 0.41 | 4.33E-01 | Mge2 | GrpE protein homolog 2, mitochondrial |
| AT3G55450 | -1.11 | 3.11E-02 | 0.34 | 3.81E-01 | 1.43 | 2.60E-03 | PBL1 | PBS1-like 1 |
| AT5G18490 | -1.11 | 1.52E-02 | 0.07 | 9.59E-01 | 1.16 | 3.55E-03 | At5g18490 | At5g18490 |
| AT2G27080 | -1.11 | 4.53E-03 | 0.96 | 1.20E-02 | 2.06 | 7.69E-13 | NHL13 | NDR1/HIN1-like protein 13 |
| AT5G07360 | -1.11 | 4.82E-03 | -0.12 | 8.32E-01 | 0.99 | 1.07E-02 | At5g07360 | Amidase family protein |
| AT5G39040 | -1.11 | 2.35E-02 | -0.28 | 2.40E-01 | 0.82 | 1.03E-01 | ABCB27 | Transporter associated with antigen processing protein 2 |
| AT2G05940 | -1.12 | 2.64E-03 | -0.12 | 7.65E-01 | 0.99 | 7.90E-03 | RIPK | Serine/threonine-protein kinase RIPK |
| AT4G23180 | -1.12 | 1.04E-02 | -0.56 | 1.55E-05 | 0.54 | 2.64E-01 | At4g23180 | Uncharacterized protein |
| AT2G39740 | -1.12 | 4.98E-04 | -0.58 | 1.98E-03 | 0.52 | 1.58E-01 | At2g39740 | Uncharacterized protein |
| AT4G05050 | -1.12 | 2.13E-02 | -0.09 | 8.38E-01 | 1.02 | 2.73E-02 | UBQ11 | Polyubiquitin |
| AT3G57330 | -1.12 | 5.94E-04 | -0.47 | 8.81E-04 | 0.64 | 7.75E-02 | ACA11 | Calcium-transporting ATPase |
| AT4G15520 | -1.13 | 2.76E-02 | 0.26 | 8.15E-01 | 1.38 | 1.22E-02 | dl3800w | OBP33pep like protein |
| AT3G49350 | -1.13 | 1.97E-03 | -0.39 | 2.98E-01 | 0.73 | 9.00E-02 | At3g49350 | Rab-GAP TBC domain-containing protein |
| AT4G22670 | -1.13 | 9.08E-03 | -0.24 | 4.11E-01 | 0.88 | 4.70E-02 | At4g22670 | TPR11 |
| AT5G64370 | -1.13 | 2.71E-03 | -0.28 | 4.62E-01 | 0.84 | 3.25E-02 | PYD3 | Beta-ureidopropionase |
| AT1G50740 | -1.13 | 1.32E-02 | -0.13 | 8.64E-01 | 0.98 | 1.02E-02 | FAX5 | Protein FATTY ACID EXPORT 5 |
| AT2G32160 | -1.14 | 9.97E-05 | -0.49 | 2.79E-03 | 0.63 | 2.64E-02 | At2g32160 | S-adenosyl-L-methionine-dependent methyltransferases superfamily protein |
| AT5G47140 | -1.14 | 1.25E-02 | -0.15 | 7.79E-01 | 0.98 | 2.87E-02 | At5g47140 | GATA27 |
| AT1G17500 | -1.14 | 2.66E-02 | 0.06 | 9.19E-01 | 1.18 | 1.47E-02 | ALA4 | Probable phospholipid-transporting ATPase 4 |
| AT4G37520 | -1.14 | 1.14E-02 | -0.05 | 9.43E-01 | 1.07 | 1.49E-02 | At4g37520 | Peroxidase |
| AT4G21390 | -1.14 | 1.88E-02 | 0.26 | 8.24E-01 | 1.39 | 1.39E-03 | At4g21390 | Receptor-like serine/threonine-protein kinase |
| AT1G50520 | -1.15 | 4.49E-02 | -1.11 | 7.11E-02 | 0.02 | 9.91E-01 | CYP705A27 | Cytochrome P450, family 705, subfamily A, polypeptide 27 |
| AT2G05320 | -1.15 | 4.42E-02 | 0.12 | 9.32E-01 | 1.25 | 1.81E-02 | GNT2 | Alpha-1,6-mannosyl-glycoprotein 2-beta-N-acetylglucosaminyltransferase |
| AT1G66590 | -1.15 | 3.82E-02 | -0.55 | 3.91E-01 | 0.57 | 3.22E-01 | COX19-1 | Cytochrome c oxidase 19-1 |
| AT3G53670 | -1.15 | 6.61E-04 | -0.26 | 4.83E-01 | 0.87 | 1.23E-02 | At3g53670 | |
| AT3G03560 | -1.15 | 1.04E-03 | -0.16 | 7.38E-01 | 0.97 | 6.19E-03 | At3g03560 | Uncharacterized protein |

| | | | | | | | | |
|---|---|---|---|---|---|---|---|---|
| AT3G16030 | -1.15 | 1.67E-02 | -0.02 | 1.00E+00 | 1.12 | 4.59E-03 | At3g16030 | Receptor-like serine/threonine-protein kinase |
| AT1G61570 | -1.15 | 3.21E-02 | -0.67 | 2.01E-01 | 0.46 | 4.11E-01 | TIM13 | Mitochondrial import inner membrane translocase subunit TIM13 |
| AT1G34380 | -1.15 | 3.46E-02 | -0.74 | 1.84E-01 | 0.40 | 5.03E-01 | OEX2 | 5'-3' exonuclease family protein |
| AT3G21630 | -1.15 | 9.58E-04 | -0.14 | 7.03E-01 | 1.00 | 4.46E-03 | At3g21630 | Protein kinase domain-containing protein |
| AT2G15480 | -1.15 | 3.76E-02 | 0.24 | 8.67E-01 | 1.38 | 1.16E-02 | UGT73B5 | Glycosyltransferase |
| AT5G53370 | -1.15 | 1.22E-02 | 0.08 | 8.93E-01 | 1.21 | 6.35E-03 | PMEPCRF | Pectinesterase |
| AT3G46930 | -1.16 | 2.52E-02 | 0.40 | 6.58E-01 | 1.54 | 8.95E-04 | RAF43 | Protein kinase superfamily protein |
| AT1G32870 | -1.16 | 4.16E-03 | -0.39 | 5.64E-01 | 0.75 | 7.49E-02 | NAC13 | NAC domain protein 13 |
| AT5G39020 | -1.17 | 5.85E-04 | -0.21 | 4.93E-01 | 0.94 | 6.58E-03 | At5g39020 | Probable receptor-like protein kinase At5g39020 |
| AT5G22630 | -1.17 | 2.95E-02 | -0.12 | 8.48E-01 | 1.03 | 4.68E-02 | ADT5 | Arogenate dehydratase 5, chloroplastic |
| AT1G47480 | -1.17 | 2.10E-03 | 0.04 | 9.87E-01 | 1.20 | 1.16E-03 | CXE2 | Probable carboxylesterase 2 |
| AT3G11040 | -1.17 | 2.09E-02 | 0.11 | 9.55E-01 | 1.27 | 1.09E-02 | ENGASE85B | mannosyl-glycoprotein endo-beta-N-acetylglucosaminidase |
| AT5G17060 | -1.17 | 2.63E-02 | -0.35 | 6.30E-01 | 0.81 | 1.07E-01 | At5g17060 | ADP-ribosylation factor |
| AT1G27060 | -1.17 | 4.80E-02 | -0.18 | 9.62E-01 | 0.97 | 2.66E-01 | | |
| AT1G25580 | -1.17 | 2.61E-02 | -0.35 | 6.58E-01 | 0.82 | 7.79E-02 | At1g25580 | NAC domain-containing protein |
| AT5G56030 | -1.17 | 1.85E-02 | -0.57 | 1.39E-04 | 0.60 | 2.80E-01 | HSP81-2 | Heat shock protein 81-2 |
| AT5G02290 | -1.18 | 2.17E-02 | -0.26 | 5.80E-01 | 0.91 | 2.92E-02 | PBL11 | Probable serine/threonine-protein kinase PBL11 |
| AT3G17700 | -1.18 | 4.83E-04 | -0.24 | 4.16E-01 | 0.93 | 6.89E-03 | CNBT1 | Cyclic nucleotide-binding transporter 1 |
| AT3G49420 | -1.18 | 4.04E-02 | -0.54 | 7.71E-01 | 0.61 | 3.60E-01 | | |
| AT1G32127 | -1.19 | 1.90E-02 | -0.67 | 2.47E-01 | 0.50 | 4.38E-01 | F3C3.9 | Uncharacterized protein F3C3.9 |
| AT4G23010 | -1.19 | 2.65E-02 | -0.06 | 9.49E-01 | 1.11 | 2.60E-02 | At4g23010 | Uncharacterized protein |
| AT4G14220 | -1.19 | 6.57E-03 | -0.27 | 2.38E-01 | 0.90 | 4.44E-02 | At4g14220 | AT4g14220/dl3150w |
| AT4G39740 | -1.19 | 3.81E-02 | -0.22 | 8.60E-01 | 0.95 | 1.12E-01 | At4g39740 | Uncharacterized protein At4g39740/T19P19_130 |
| AT5G49520 | -1.19 | 1.14E-02 | 0.18 | 7.95E-01 | 1.36 | 1.51E-03 | At5g49520 | WRKY domain-containing protein |
| AT2G25520 | -1.19 | 1.14E-02 | -0.26 | 1.78E-01 | 0.92 | 5.72E-02 | At2g25520 | Probable sugar phosphate/phosphate translocator At2g25520 |
| AT3G23100 | -1.20 | 1.93E-02 | -0.66 | 3.26E-01 | 0.53 | 3.58E-01 | At3g23100 | Carbamoyl-phosphate synthetase large subunit-like ATP-binding domain-containing protein |
| AT1G64170 | -1.20 | 3.62E-02 | 1.07 | 1.06E-01 | 2.26 | 9.55E-04 | CHX16 | Cation/H |
| AT5G15820 | -1.20 | 3.42E-02 | -0.69 | 6.69E-01 | 0.48 | 5.05E-01 | | |
| AT2G17760 | -1.20 | 1.76E-02 | -0.18 | 6.42E-01 | 1.01 | 4.17E-02 | APF1 | Aspartyl protease family protein 1 |
| AT3G21810 | -1.20 | 1.42E-02 | -0.80 | 6.40E-02 | 0.39 | 4.56E-01 | At3g21810 | Zinc finger C-x8-C-x5-C-x3-H type family protein |
| AT4G16960 | -1.20 | 2.33E-03 | -0.08 | 9.67E-01 | 1.11 | 9.16E-03 | SIKIC3 | Disease resistance protein |
| AT3G45040 | -1.20 | 2.19E-02 | 0.06 | 9.68E-01 | 1.23 | 1.05E-02 | DOK1 | dolichol kinase |
| AT5G26030 | -1.20 | 2.12E-03 | 0.18 | 7.36E-01 | 1.36 | 5.48E-04 | FC1 | Ferrochelatase-1, chloroplastic/mitochondrial |
| AT3G04770 | -1.20 | 2.29E-02 | -1.14 | 6.46E-02 | 0.04 | 9.74E-01 | RPSAB | Small ribosomal subunit protein uS2 |
| AT3G13330 | -1.20 | 1.86E-03 | -0.19 | 3.93E-01 | 1.00 | 8.86E-03 | PA200 | Proteasome activator subunit 4 |
| AT1G55610 | -1.20 | 2.26E-02 | -0.40 | 6.00E-01 | 0.78 | 1.01E-01 | BRL1 | Serine/threonine-protein kinase BRI1-like 1 |
| AT3G20250 | -1.20 | 2.69E-03 | 0.03 | 9.75E-01 | 1.22 | 1.16E-03 | PUM5 | Pumilio 5 |
| AT2G26170 | -1.21 | 1.94E-02 | -0.39 | 2.35E-01 | 0.79 | 1.59E-01 | CYP711A1 | Cytochrome P450 711A1 |
| AT2G04400 | -1.21 | 3.24E-02 | 0.10 | 8.77E-01 | 1.30 | 1.43E-02 | IGPS | Indole-3-glycerol phosphate synthase, chloroplastic |
| AT3G58600 | -1.22 | 1.25E-03 | -0.23 | 6.48E-01 | 0.97 | 7.71E-03 | At3g58600 | NECAP PHear domain-containing protein |
| AT3G16860 | -1.22 | 2.79E-03 | 0.40 | 5.17E-01 | 1.61 | 1.47E-06 | COBL8 | COBRA-like protein 8 |
| AT5G19750 | -1.22 | 1.15E-02 | -1.15 | 1.95E-03 | 0.06 | 9.46E-01 | At5g19750 | Peroxisomal membrane 22 kDa |
| AT1G32920 | -1.22 | 2.22E-03 | 0.56 | 1.78E-01 | 1.77 | 1.39E-06 | At1g32920 | At1g32920/F9L11_25 |
| AT3G47530 | -1.22 | 8.37E-03 | -0.30 | 7.68E-01 | 0.91 | 6.65E-02 | PCMP-H76 | Pentatricopeptide repeat-containing protein At3g47530 |
| AT3G02520 | -1.22 | 2.71E-02 | -0.28 | 2.77E-01 | 0.93 | 1.07E-01 | GRF7 | 14-3-3-like protein GF14 nu |
| AT5G57035 | -1.22 | 3.07E-03 | -0.37 | 5.21E-02 | 0.84 | 5.47E-02 | At5g57035 | U-box domain-containing protein kinase family protein |
| AT5G56230 | -1.22 | 2.98E-02 | -0.12 | 9.50E-01 | 1.09 | 6.86E-02 | At5g56230 | PRA1 family protein |
| AT1G30040 | -1.23 | 2.17E-02 | 0.01 | 1.00E+00 | 1.21 | 1.21E-01 | At1g30040 | GA2OX2 |
| AT1G59870 | -1.23 | 8.33E-03 | 0.09 | 7.81E-01 | 1.30 | 2.79E-03 | ABCG36 | ABC transporter G family member 36 |
| AT1G05010 | -1.23 | 4.89E-02 | -0.37 | 2.33E-01 | 0.85 | 1.61E-01 | ACO4 | 1-aminocyclopropane-1-carboxylate oxidase 4 |
| AT1G31540 | -1.23 | 2.59E-03 | -0.30 | 3.27E-01 | 0.91 | 8.74E-03 | T8E3.20 | ADP-ribosyl cyclase/cyclic ADP-ribose hydrolase |
| AT1G21270 | -1.23 | 2.20E-03 | -0.46 | 4.77E-03 | 0.76 | 6.86E-02 | WAK2 | Wall-associated receptor kinase 2 |
| AT5G47580 | -1.23 | 3.15E-02 | 0.00 | 1.00E+00 | 1.21 | 2.93E-02 | At5g47580 | ASG7 |
| AT2G17130 | -1.24 | 2.19E-02 | -0.11 | 9.59E-01 | 1.11 | 2.40E-02 | At2g17130 | |
| AT1G33610 | -1.24 | 4.85E-04 | -1.08 | 8.83E-06 | 0.14 | 8.17E-01 | At1g33610 | Leucine-rich repeat-containing N-terminal plant-type domain-containing protein |
| AT1G22280 | -1.24 | 2.15E-03 | -0.21 | 4.97E-01 | 1.02 | 1.02E-02 | PAPP2C | Phytochrome-associated protein phosphatase type 2C |
| AT1G74300 | -1.24 | 1.75E-02 | -0.29 | 8.02E-01 | 0.93 | 1.26E-01 | At1g74300 | Alpha/beta-Hydrolases superfamily protein |
| AT2G39270 | -1.24 | 2.12E-02 | 0.07 | 9.99E-01 | 1.34 | 4.68E-02 | | P-loop containing nucleoside triphosphate hydrolases superfamily protein |
| AT1G30420 | -1.24 | 2.69E-04 | -0.20 | 7.36E-01 | 1.03 | 2.67E-03 | ABCC11 | Multidrug resistance-associated protein 12 |
| AT4G38260 | -1.24 | 3.74E-02 | 0.06 | 9.94E-01 | 1.29 | 4.13E-02 | At4g38260 | Uncharacterized protein At4g38260 |
| AT3G56800 | -1.25 | 3.98E-02 | -0.19 | 5.88E-01 | 1.05 | 6.89E-02 | CAM3 | Calmodulin-3 |
| AT4G22780 | -1.25 | 3.75E-04 | -0.06 | 9.55E-01 | 1.18 | 4.21E-04 | ACR7 | ACT domain-containing protein ACR7 |
| AT5G18480 | -1.26 | 5.28E-03 | -0.06 | 9.31E-01 | 1.17 | 1.07E-02 | IPUT1 | Inositol phosphorylceramide glucuronosyltransferase 1 |
| AT5G61010 | -1.26 | 2.42E-04 | -0.45 | 2.23E-02 | 0.79 | 1.65E-02 | EXO70E2 | Exocyst complex component EXO70E2 |
| AT2G41440 | -1.26 | 3.72E-02 | 0.22 | 9.47E-01 | 1.45 | 2.52E-02 | | |
| AT1G03370 | -1.26 | 3.06E-04 | -0.48 | 5.85E-03 | 0.76 | 3.58E-02 | At1g03370 | C2 calcium/lipid-binding and GRAM domain containing protein |
| AT4G14746 | -1.26 | 1.70E-03 | -0.09 | 9.04E-01 | 1.15 | 2.60E-03 | At4g14746 | Neurogenic locus notch-like protein |
| AT4G36710 | -1.26 | 2.71E-02 | 0.27 | 8.63E-01 | 1.51 | 3.47E-03 | SCL15 | Scarecrow-like protein 15 |
| AT3G17240 | -1.26 | 3.83E-03 | -0.20 | 6.49E-01 | 1.05 | 1.60E-02 | At3g17240 | Dihydrolipoyl dehydrogenase |
| AT2G40270 | -1.26 | 1.53E-03 | -0.29 | 1.09E-01 | 0.95 | 2.02E-02 | At2g40270 | Protein kinase family protein |
| AT2G03120 | -1.26 | 6.41E-03 | -0.15 | 6.49E-01 | 1.09 | 2.07E-02 | SPP | Signal peptide peptidase |
| AT5G20830 | -1.26 | 1.66E-02 | -0.20 | 8.36E-01 | 1.05 | 1.43E-02 | SUS1 | Sucrose synthase 1 |
| AT1G23710 | -1.26 | 9.31E-03 | -0.28 | 8.11E-01 | 0.97 | 1.04E-01 | At1g23710 | At1g23710 |
| AT3G10010 | -1.27 | 1.83E-04 | -0.47 | 5.13E-02 | 0.79 | 2.22E-02 | DML2 | DEMETER-like protein 2 |
| AT5G52870 | -1.27 | 4.66E-03 | -0.30 | 6.17E-01 | 0.95 | 3.41E-02 | MAKR5 | Membrane-associated kinase regulator |
| AT3G53810 | -1.27 | 9.64E-03 | -0.28 | 4.43E-01 | 0.97 | 4.46E-02 | LECRK42 | L-type lectin-domain containing receptor kinase IV.2 |
| AT5G36930 | -1.27 | 6.97E-05 | -0.52 | 2.61E-04 | 0.73 | 3.78E-02 | MLF18.50 | Disease resistance protein |
| AT2G23200 | -1.27 | 2.15E-04 | -0.60 | 5.42E-04 | 0.66 | 7.05E-02 | At2g23200 | Probable receptor-like protein kinase At2g23200 |

| | | | | | | | | |
|---|---|---|---|---|---|---|---|---|
| AT5G63680 | -1.27 | 4.59E-02 | -0.11 | 9.99E-01 | 1.15 | 1.66E-01 | MBK5.16 | Pyruvate kinase |
| AT2G23320 | -1.28 | 1.32E-04 | -0.25 | 3.58E-01 | 1.02 | 2.43E-03 | WRKY15 | WRKY DNA-binding protein 15 |
| AT4G38150 | -1.28 | 4.03E-02 | -0.69 | 6.40E-01 | 0.60 | 4.03E-01 | | |
| AT3G59880 | -1.28 | 1.42E-02 | -0.83 | 4.69E-02 | 0.43 | 4.94E-01 | F24G16.150 | Uncharacterized protein F24G16.150 |
| AT2G05630 | -1.28 | 1.60E-03 | -0.11 | 9.63E-01 | 1.16 | 6.97E-03 | ATG8D | Autophagy-related protein |
| AT4G08230 | -1.28 | 3.00E-02 | -0.17 | 6.84E-01 | 1.10 | 6.45E-02 | At4g08230 | Glycine-rich protein |
| AT1G22410 | -1.28 | 2.60E-02 | -0.24 | 4.03E-01 | 1.03 | 7.09E-02 | At1g22410 | Phospho-2-dehydro-3-deoxyheptonate aldolase |
| AT2G34250 | -1.29 | 1.95E-02 | -0.21 | 3.54E-01 | 1.05 | 5.03E-02 | At2g34250 | AT2G34250 protein |
| AT3G53780 | -1.29 | 1.66E-02 | -0.36 | 4.20E-01 | 0.90 | 1.06E-01 | RBL4 | RHOMBOID-like protein |
| AT2G47000 | -1.29 | 3.69E-03 | 0.14 | 8.94E-01 | 1.41 | 1.03E-03 | ABCB4 | ATP binding cassette subfamily B4 |
| AT3G21220 | -1.29 | 3.90E-03 | -0.14 | 7.56E-01 | 1.14 | 9.16E-03 | MKK5 | Mitogen-activated protein kinase kinase 5 |
| AT1G50745 | -1.29 | 1.59E-02 | -0.26 | 7.36E-01 | 1.01 | 9.16E-02 | At1g50745 | Plant mobile domain protein |
| AT4G01750 | -1.29 | 6.75E-03 | 0.12 | 8.73E-01 | 1.40 | 3.73E-03 | RGXT2 | UDP-D-xylose:L-fucose alpha-1,3-D-xylosyltransferase |
| AT4G00026 | -1.29 | 1.70E-02 | 0.20 | 8.89E-01 | 1.47 | 5.09E-03 | TIM21 | Probable mitochondrial import inner membrane translocase subunit TIM21 |
| AT5G11650 | -1.29 | 2.73E-03 | -0.46 | 4.37E-01 | 0.81 | 8.17E-02 | MAGL13 | Alpha/beta-Hydrolases superfamily protein |
| AT5G58120 | -1.29 | 1.13E-03 | -0.10 | 9.04E-01 | 1.18 | 5.17E-04 | At5g58120 | ADP-ribosyl cyclase/cyclic ADP-ribose hydrolase |
| AT3G13235 | -1.29 | 3.60E-03 | 0.08 | 9.40E-01 | 1.35 | 2.67E-03 | DDI1 | Ubiquitin family protein |
| AT1G72930 | -1.29 | 6.18E-04 | -0.38 | 1.29E-01 | 0.90 | 7.17E-03 | TIR | Toll/interleukin-1 receptor-like protein |
| AT5G11000 | -1.29 | 4.82E-02 | -0.18 | 5.56E-01 | 1.10 | 9.05E-02 | At5g11000 | DUF868 family protein |
| AT5G44070 | -1.29 | 1.51E-03 | -0.53 | 3.30E-03 | 0.75 | 7.53E-02 | CAD1 | glutathione gamma-glutamylcysteinyltransferase |
| AT5G18310 | -1.29 | 2.49E-02 | -0.04 | 9.83E-01 | 1.24 | 3.35E-02 | At5g18310 | Ubiquitin hydrolase |
| AT4G11840 | -1.29 | 8.42E-04 | -0.50 | 5.97E-02 | 0.78 | 3.63E-02 | PLDGAMMA | Phospholipase D |
| AT3G16720 | -1.29 | 1.99E-03 | 0.00 | 1.00E+00 | 1.28 | 2.27E-04 | ATL2 | RING-H2 finger protein ATL2 |
| AT1G69730 | -1.30 | 1.57E-03 | -0.32 | 3.35E-01 | 0.97 | 5.73E-03 | WAKL9 | Wall-associated receptor kinase-like 9 |
| AT4G28400 | -1.30 | 1.45E-02 | 0.04 | 9.75E-01 | 1.32 | 1.21E-02 | At4g28400 | Protein phosphatase 2C family protein |
| AT3G16510 | -1.30 | 5.31E-03 | -0.38 | 3.94E-01 | 0.91 | 2.34E-02 | At3g16510 | |
| AT5G39785 | -1.31 | 5.69E-03 | -0.03 | 1.00E+00 | 1.26 | 1.21E-02 | At5g39780 | Gb\|AAF22924.1 |
| AT5G19980 | -1.31 | 6.75E-03 | -0.47 | 4.73E-03 | 0.82 | 1.16E-01 | GFT1 | GDP-fucose transporter 1 |
| AT5G38990 | -1.31 | 2.23E-03 | -0.58 | 1.11E-05 | 0.71 | 1.39E-01 | At5g38990 | Probable receptor-like protein kinase At5g38990 |
| AT1G62730 | -1.31 | 4.55E-02 | -0.44 | 6.37E-01 | 0.87 | 1.76E-01 | At1g62730 | Terpenoid synthases superfamily protein |
| AT2G30250 | -1.31 | 4.77E-03 | -0.20 | 5.83E-01 | 1.11 | 1.22E-02 | WRKY25 | Probable WRKY transcription factor 25 |
| AT1G52200 | -1.31 | 6.57E-04 | -0.22 | 6.36E-01 | 1.08 | 2.28E-03 | At1g52200 | PLAC8 family protein |
| AT2G45010 | -1.32 | 1.20E-02 | -0.42 | 1.93E-01 | 0.88 | 1.01E-01 | At2g45010 | PLAC8 family protein |
| AT2G28940 | -1.32 | 1.66E-03 | 0.24 | 8.42E-01 | 1.55 | 1.91E-03 | PBL37 | At2g28940 |
| AT1G62305 | -1.32 | 1.52E-02 | -0.79 | 9.46E-02 | 0.52 | 4.21E-01 | At1g62305 | At1g62305 |
| AT3G26090 | -1.32 | 3.46E-03 | -0.40 | 2.58E-01 | 0.91 | 5.29E-02 | RGS1 | REGULATOR OF G-PROTEIN SIGNALING 1 |
| AT2G16595 | -1.32 | 1.67E-02 | -0.27 | 7.81E-01 | 1.03 | 6.99E-02 | At2g16595 | Translocon-associated protein subunit alpha |
| AT4G33300 | -1.32 | 3.78E-04 | -0.13 | 6.92E-01 | 1.18 | 1.47E-03 | At4g33300 | Probable disease resistance protein At4g33300 |
| AT3G04480 | -1.33 | 2.52E-05 | -0.70 | 3.63E-06 | 0.61 | 1.05E-01 | At3g04480 | Diphthine--ammonia ligase |
| AT5G49570 | -1.33 | 9.48E-04 | -0.27 | 4.25E-01 | 1.04 | 1.41E-02 | PNG1 | Peptide-N |
| AT4G35110 | -1.33 | 3.23E-02 | -0.15 | 8.95E-01 | 1.17 | 1.07E-01 | PEARLI | Phospholipase-like protein |
| AT5G39950 | -1.33 | 5.39E-03 | -0.16 | 8.28E-01 | 1.15 | 1.84E-02 | TRX2 | Thioredoxin H2 |
| AT1G76600 | -1.33 | 1.35E-02 | 0.72 | 1.45E-01 | 2.04 | 3.61E-05 | At1g76600 | |
| AT3G12740 | -1.33 | 8.49E-03 | -0.03 | 9.74E-01 | 1.29 | 9.09E-03 | ALIS1 | ALA-interacting subunit 1 |
| AT5G64550 | -1.33 | 4.52E-03 | -0.18 | 8.52E-01 | 1.13 | 1.03E-02 | At5g64550 | Uncharacterized protein At5g64550 |
| AT4G01740 | -1.33 | 9.16E-03 | -0.74 | 1.83E-01 | 0.58 | 3.33E-01 | At4g01740 | Cysteine/Histidine-rich C1 domain family protein |
| AT1G07630 | -1.33 | 1.40E-03 | -0.25 | 3.31E-01 | 1.07 | 1.15E-02 | PLL5 | Probable protein phosphatase 2C 4 |
| AT1G62422 | -1.33 | 3.60E-02 | -0.38 | 6.24E-01 | 0.94 | 1.67E-01 | At1g62422 | Uncharacterized protein |
| AT1G72700 | -1.33 | 2.29E-02 | -0.23 | 4.88E-01 | 1.10 | 6.61E-02 | ALA5 | Probable phospholipid-transporting ATPase 5 |
| AT4G08470 | -1.34 | 2.02E-05 | -0.37 | 1.86E-01 | 0.95 | 1.95E-04 | MEKK3 | MAPK/ERK kinase kinase 3 |
| AT3G52420 | -1.34 | 2.69E-02 | 0.39 | 9.09E-01 | 1.80 | 5.11E-03 | | |
| AT3G47580 | -1.34 | 4.97E-02 | -0.91 | 6.03E-01 | 0.45 | 5.43E-01 | | |
| AT2G40520 | -1.34 | 5.04E-03 | -0.14 | 8.43E-01 | 1.19 | 5.86E-03 | At2g40520 | Polymerase nucleotidyl transferase domain-containing protein |
| AT1G02360 | -1.34 | 2.77E-03 | -0.36 | 7.03E-01 | 0.97 | 3.08E-02 | At1g02360 | At1g02360 |
| AT1G25220 | -1.34 | 1.53E-02 | 0.04 | 9.94E-01 | 1.36 | 1.08E-02 | ASB1 | Anthranilate synthase beta subunit 1 |
| AT4G23540 | -1.35 | 4.16E-05 | -0.44 | 5.38E-02 | 0.89 | 9.64E-03 | At4g23540 | ARM repeat superfamily protein |
| AT5G26860 | -1.35 | 6.07E-03 | -0.62 | 4.74E-02 | 0.72 | 1.59E-01 | At5g26860 | AT5g26860/F2P16_120 |
| AT1G33560 | -1.35 | 9.74E-03 | -0.38 | 2.57E-01 | 0.96 | 9.06E-02 | ADR1 | Disease resistance protein ADR1 |
| AT5G14930 | -1.35 | 2.70E-03 | -0.12 | 7.72E-01 | 1.21 | 7.40E-03 | SAG101 | Senescence-associated gene 101 |
| AT3G51980 | -1.35 | 4.50E-02 | -0.10 | 9.28E-01 | 1.24 | 5.84E-02 | At3g51980 | Nucleotide exchange factor Fes1 domain-containing protein |
| AT3G27010 | -1.35 | 6.95E-03 | -0.81 | 3.65E-01 | 0.53 | 5.16E-01 | TCP20 | Transcription factor TCP20 |
| AT5G52540 | -1.35 | 1.87E-04 | -0.26 | 4.49E-01 | 1.08 | 4.25E-03 | At5g52540 | |
| AT5G59550 | -1.35 | 8.17E-05 | -0.21 | 7.96E-01 | 1.13 | 7.01E-03 | RDUF2 | RING-type E3 ubiquitin transferase |
| AT5G54100 | -1.36 | 2.36E-03 | -0.73 | 2.16E-01 | 0.61 | 2.21E-01 | SLP2 | At5g54100 |
| AT1G09070 | -1.36 | 3.64E-03 | 0.19 | 6.47E-01 | 1.53 | 3.41E-04 | SRC2 | Protein SRC2 homolog |
| AT3G02230 | -1.36 | 3.70E-02 | -0.55 | 6.07E-02 | 0.79 | 2.46E-01 | RGP1 | UDP-arabinopyranose mutase 1 |
| AT5G39450 | -1.36 | 9.16E-05 | -0.67 | 1.24E-01 | 0.67 | 1.47E-01 | At5g39450 | |
| AT4G26910 | -1.36 | 2.01E-03 | -0.30 | 2.41E-01 | 1.05 | 2.56E-02 | At4g26910 | Dihydrolipoyllysine-residue succinyltransferase component of 2-oxoglutarate dehydrogenase complex 2, mitochondrial |
| AT3G24503 | -1.37 | 4.17E-02 | -0.04 | 9.55E-01 | 1.31 | 4.49E-02 | ALDH2C4 | Aldehyde dehydrogenase family 2 member C4 |
| AT5G53120 | -1.37 | 4.09E-02 | 0.02 | 1.00E+00 | 1.36 | 3.26E-02 | SPDS3 | Spermidine synthase 3 |
| AT1G65800 | -1.37 | 6.82E-04 | -0.25 | 4.11E-01 | 1.10 | 2.13E-03 | RK2 | Receptor kinase 2 |
| AT1G11310 | -1.37 | 1.18E-03 | -0.39 | 6.57E-03 | 0.96 | 2.71E-02 | At1g11310 | At1g11310/T28P6_23 |
| AT4G00240 | -1.37 | 1.67E-02 | 0.27 | 7.98E-01 | 1.63 | 3.90E-03 | PLDBETA2 | Phospholipase D |
| AT3G23560 | -1.37 | 9.73E-03 | -0.47 | 5.39E-01 | 0.87 | 1.51E-01 | ALF5 | MATE efflux family protein |
| AT1G75000 | -1.37 | 1.82E-03 | -0.28 | 5.28E-01 | 1.07 | 1.25E-02 | ELO3 | F25A4.4 |
| AT3G09350 | -1.37 | 3.10E-03 | -0.33 | 5.21E-01 | 1.03 | 4.03E-02 | FES1A | Fes1A |
| AT4G25390 | -1.37 | 3.72E-02 | 0.33 | 6.98E-01 | 1.69 | 5.58E-03 | At4g25390 | Receptor-like serine/threonine-protein kinase At4g25390 |
| AT3G49370 | -1.38 | 2.45E-03 | -0.18 | 9.05E-01 | 1.19 | 1.02E-02 | CRK6 | CDPK-related kinase 6 |

| | | | | | | | | |
|---|---|---|---|---|---|---|---|---|
| AT4G01910 | -1.38 | 2.13E-03 | 0.04 | 9.96E-01 | 1.40 | 1.54E-03 | At4g01910 | Cysteine/Histidine-rich C1 domain family protein |
| AT2G17720 | -1.38 | 1.50E-02 | 0.09 | 9.28E-01 | 1.46 | 8.48E-03 | P4H5 | Prolyl 4-hydroxylase 5 |
| AT4G09570 | -1.38 | 3.45E-03 | -0.11 | 9.18E-01 | 1.26 | 5.83E-03 | CPK4 | Calcium-dependent protein kinase 4 |
| AT3G52100 | -1.38 | 3.64E-03 | -0.79 | 1.17E-03 | 0.57 | 3.06E-01 | At3g52100 | RING/FYVE/PHD-type zinc finger family protein |
| AT1G16260 | -1.38 | 4.85E-05 | -0.51 | 5.01E-03 | 0.86 | 9.48E-03 | At1g16260 | Wall-associated receptor kinase-like |
| AT2G37110 | -1.38 | 3.53E-03 | -0.09 | 9.42E-01 | 1.28 | 1.74E-02 | At2g37110 | |
| AT2G33530 | -1.38 | 4.48E-04 | -0.61 | 6.27E-04 | 0.75 | 8.56E-02 | SCPL46 | Serine carboxypeptidase-like 46 |
| AT4G00960 | -1.39 | 1.86E-02 | -0.69 | 4.85E-01 | 0.69 | 2.95E-01 | A_TM018A10 | Protein kinase superfamily protein |
| AT1G50180 | -1.39 | 4.86E-02 | -0.01 | 9.97E-01 | 1.35 | 2.41E-02 | | |
| AT5G54840 | -1.39 | 1.06E-03 | -0.29 | 6.38E-01 | 1.08 | 1.41E-02 | SGP1 | At5g54840 |
| AT2G31990 | -1.39 | 1.83E-02 | -0.03 | 1.00E+00 | 1.35 | 1.90E-02 | At2g31990 | Exostosin family protein |
| AT1G52800 | -1.39 | 9.73E-03 | 0.81 | 5.98E-01 | 2.20 | 7.01E-05 | At1g52800 | hypothetical protein |
| AT5G15860 | -1.40 | 1.25E-05 | 0.03 | 9.84E-01 | 1.41 | 1.18E-05 | ICME | Isoprenylcysteine alpha-carbonyl methylesterase ICME |
| AT4G01010 | -1.40 | 1.26E-02 | 0.15 | 7.76E-01 | 1.54 | 3.01E-03 | CNGC13 | Putative cyclic nucleotide-gated ion channel 13 |
| AT3G22060 | -1.40 | 3.48E-08 | -0.82 | 1.74E-07 | 0.56 | 3.44E-02 | CRRSP38 | Cysteine-rich repeat secretory protein 38 |
| AT3G03855 | -1.40 | 1.16E-04 | -0.49 | 2.07E-01 | 0.90 | 1.66E-02 | | |
| AT3G11650 | -1.40 | 1.14E-02 | 0.04 | 9.87E-01 | 1.43 | 8.78E-03 | NHL2 | NDR1/HIN1-like protein 2 |
| AT1G69840 | -1.40 | 9.40E-03 | 0.16 | 8.03E-01 | 1.55 | 1.09E-03 | HIR2 | Hypersensitive-induced response protein 2 |
| AT5G45800 | -1.41 | 1.31E-02 | -0.47 | 5.45E-02 | 0.92 | 9.91E-02 | CAMRLK | Calmodulin-binding receptor kinase CaMRLK |
| AT3G05830 | -1.41 | 2.48E-02 | -0.52 | 5.35E-01 | 0.88 | 9.97E-02 | ATNEAP1 | Spindle pole body component-like protein |
| AT4G23570 | -1.41 | 7.25E-04 | 0.01 | 9.99E-01 | 1.41 | 2.52E-04 | SGT1A | Protein SGT1 homolog A |
| AT5G45110 | -1.41 | 4.99E-04 | 0.04 | 9.65E-01 | 1.44 | 4.00E-05 | NPR3 | NPR1-like protein 3 |
| AT2G03450 | -1.41 | 2.77E-02 | -0.08 | 9.63E-01 | 1.30 | 1.73E-02 | PAP9 | Probable inactive purple acid phosphatase 9 |
| AT3G03440 | -1.41 | 3.13E-03 | -0.33 | 4.26E-01 | 1.07 | 3.38E-02 | At3g03440 | ARM repeat superfamily protein |
| AT2G30020 | -1.41 | 9.89E-07 | 0.59 | 2.15E-02 | 1.99 | 2.35E-13 | At2g30020 | Probable protein phosphatase 2C 25 |
| AT1G71210 | -1.41 | 1.66E-03 | -0.52 | 2.54E-01 | 0.87 | 8.88E-02 | At1g71210 | Pentatricopeptide repeat-containing protein At1g71210, mitochondrial |
| AT4G00955 | -1.42 | 6.05E-06 | -0.60 | 5.36E-02 | 0.81 | 8.45E-03 | At4g00955 | Uncharacterized protein At4g00955 |
| AT3G45640 | -1.42 | 8.39E-04 | -0.02 | 9.88E-01 | 1.38 | 3.52E-04 | MPK3 | Mitogen-activated protein kinase 3 |
| AT1G50750 | -1.42 | 1.15E-02 | -0.33 | 8.87E-01 | 1.08 | 8.01E-02 | At1g50750 | Aminotransferase-like, mobile domain protein |
| AT3G60520 | -1.42 | 3.93E-02 | -0.09 | 9.37E-01 | 1.31 | 6.08E-02 | At3g60520 | Zinc ion-binding protein |
| AT5G66320 | -1.42 | 3.06E-03 | 0.12 | 8.90E-01 | 1.52 | 5.55E-04 | GATA5 | GATA transcription factor 5 |
| AT2G01650 | -1.42 | 2.93E-03 | -0.24 | 5.70E-01 | 1.17 | 1.25E-02 | PUX2 | Plant UBX domain-containing protein 2 |
| AT4G18890 | -1.42 | 1.16E-02 | -0.05 | 9.93E-01 | 1.37 | 2.10E-02 | BEH3 | Protein BZR1 homolog |
| AT1G13210 | -1.42 | 8.29E-04 | -0.21 | 4.62E-01 | 1.20 | 2.71E-03 | ALA11 | Probable phospholipid-transporting ATPase 11 |
| AT2G28500 | -1.43 | 1.52E-02 | -0.59 | 7.03E-01 | 0.84 | 1.98E-01 | | |
| AT4G12120 | -1.43 | 4.49E-04 | -0.35 | 4.52E-01 | 1.07 | 1.65E-02 | SEC1B | Protein transport Sec1b |
| AT4G17070 | -1.43 | 2.16E-04 | -0.44 | 3.72E-01 | 0.98 | 1.66E-02 | At4g17070 | PPIase cyclophilin-type domain-containing protein |
| AT3G19580 | -1.43 | 5.20E-04 | 0.04 | 9.76E-01 | 1.45 | 1.36E-05 | AZF2 | Zinc finger protein AZF2 |
| AT2G43850 | -1.43 | 1.11E-05 | -0.06 | 9.62E-01 | 1.36 | 6.45E-05 | ILK1 | Integrin-linked protein kinase family |
| AT3G20960 | -1.44 | 3.76E-02 | 0.29 | 7.88E-01 | 1.70 | 5.94E-03 | CYP705A33 | Cytochrome P450, family 705, subfamily A, polypeptide 33 |
| AT1G80610 | -1.44 | 3.08E-04 | -0.11 | 8.73E-01 | 1.31 | 8.08E-04 | At1g80610 | At1g80610/T21F11_6 |
| AT1G74200 | -1.44 | 5.98E-03 | 0.39 | 8.75E-01 | 1.75 | 4.17E-04 | RLP16 | Putative receptor-like protein 16 |
| AT2G02390 | -1.44 | 2.93E-03 | -0.02 | 9.94E-01 | 1.41 | 4.16E-03 | GSTZ1 | Glutathione S-transferase zeta 1 |
| AT3G55980 | -1.44 | 2.40E-03 | 0.26 | 6.48E-01 | 1.69 | 3.35E-06 | SZF1 | Salt-inducible zinc finger 1 |
| AT1G27350 | -1.44 | 1.37E-02 | -0.18 | 7.58E-01 | 1.26 | 2.57E-02 | At1g27330 | At1g27330 |
| AT1G03290 | -1.44 | 1.30E-03 | -0.22 | 5.96E-01 | 1.21 | 4.68E-03 | At1g03290 | ELKS/Rab6-interacting/CAST family protein |
| AT5G01610 | -1.45 | 9.82E-03 | -0.27 | 8.30E-01 | 1.17 | 2.32E-02 | At5g01610 | Uncharacterized protein At5g01610 |
| AT4G33420 | -1.45 | 2.35E-03 | 0.62 | 7.50E-01 | 2.05 | 5.48E-04 | At4g33420 | Peroxidase |
| AT1G29310 | -1.45 | 2.91E-02 | -0.30 | 5.59E-01 | 1.13 | 1.14E-01 | SEC | SecY protein transport family protein |
| AT5G42440 | -1.45 | 1.65E-03 | -0.15 | 8.60E-01 | 1.29 | 3.87E-03 | At5g42440 | Protein kinase domain-containing protein |
| AT4G17720 | -1.45 | 2.61E-02 | 0.00 | 1.00E+00 | 1.43 | 2.71E-02 | BPL1 | Putative RRM-containing protein |
| AT1G67850 | -1.45 | 9.38E-04 | 0.12 | 9.12E-01 | 1.56 | 1.34E-04 | At1g67850 | |
| AT1G23140 | -1.45 | 1.79E-02 | -0.69 | 7.22E-01 | 0.74 | 4.25E-01 | At1g23140 | C2 domain-containing protein |
| AT5G23850 | -1.45 | 1.49E-02 | -0.09 | 8.94E-01 | 1.34 | 2.85E-02 | MRO11.11 | At5g23850 |
| AT5G45500 | -1.45 | 4.00E-04 | -0.26 | 2.95E-01 | 1.18 | 5.58E-03 | MFC19.17 | RNI-like superfamily protein |
| AT2G40950 | -1.46 | 1.45E-02 | -0.23 | 5.70E-01 | 1.21 | 5.00E-02 | BZIP17 | bZIP transcription factor 17 |
| AT1G18890 | -1.46 | 1.02E-02 | -0.25 | 4.82E-01 | 1.20 | 4.03E-02 | CPK10 | Calcium-dependent protein kinase 10 |
| AT5G06750 | -1.46 | 1.29E-02 | -0.50 | 4.47E-01 | 0.94 | 1.24E-01 | APD8 | Protein phosphatase 2C family protein |
| AT3G55470 | -1.47 | 3.35E-03 | 0.16 | 9.22E-01 | 1.61 | 7.50E-03 | At3g55470 | Calcium-dependent lipid-binding |
| AT3G15760 | -1.47 | 1.26E-03 | -0.62 | 3.28E-01 | 0.83 | 9.16E-02 | At3g15760 | Transmembrane protein |
| AT5G09590 | -1.47 | 1.99E-02 | -0.20 | 7.25E-01 | 1.25 | 5.35E-02 | HSP70-10 | Heat shock 70 kDa protein 10, mitochondrial |
| AT2G19710 | -1.47 | 5.11E-04 | -0.22 | 6.14E-01 | 1.24 | 3.84E-03 | At2g19710 | Uncharacterized protein At2g19710 |
| AT1G13990 | -1.47 | 2.26E-02 | 0.09 | 8.94E-01 | 1.55 | 6.01E-03 | At1g13990 | Plant/protein |
| AT2G38830 | -1.47 | 9.89E-03 | -0.07 | 9.76E-01 | 1.38 | 3.59E-02 | At2g38830 | At2g38830 |
| AT3G26230 | -1.47 | 1.20E-03 | -0.43 | 6.66E-02 | 1.03 | 2.39E-02 | At3g26230 | Uncharacterized protein |
| AT5G19930 | -1.47 | 2.15E-03 | -0.85 | 1.28E-02 | 0.60 | 2.96E-01 | PGR | Protein PGR |
| AT1G18570 | -1.48 | 3.48E-03 | 0.18 | 6.23E-01 | 1.64 | 8.17E-04 | MYB51 | Transcription factor MYB51 |
| AT5G05320 | -1.48 | 4.32E-02 | | | 2.06 | 8.66E-03 | | |
| AT1G07220 | -1.49 | 3.72E-03 | 0.42 | 5.31E-01 | 1.90 | 1.43E-04 | At1g07220 | F10K1.7 protein |
| AT1G52780 | -1.49 | 3.47E-04 | -0.32 | 9.12E-02 | 1.16 | 6.45E-03 | At1g52780 | PII, uridylyltransferase |
| AT5G54720 | -1.49 | 4.30E-04 | 0.02 | 9.79E-01 | 1.50 | 2.42E-04 | At5g54720 | Uncharacterized protein |
| AT1G67800 | -1.49 | 2.01E-03 | -0.13 | 7.56E-01 | 1.35 | 5.31E-03 | RGLG5 | Copine |
| AT3G19010 | -1.49 | 4.49E-04 | -0.26 | 2.81E-01 | 1.21 | 3.41E-03 | 2-OXOGLUT/ | 2-oxoglutarate |
| AT5G21090 | -1.49 | 4.81E-02 | -0.01 | 9.97E-01 | 1.47 | 4.11E-02 | LRR1 | Leucine-rich repeat protein 1 |
| AT5G08380 | -1.49 | 1.62E-02 | 0.07 | 9.50E-01 | 1.55 | 1.25E-02 | AGAL1 | Alpha-galactosidase |
| AT4G02660 | -1.50 | 2.68E-03 | -0.17 | 8.77E-01 | 1.31 | 1.26E-02 | BCHA2 | Beige/BEACH and WD40 domain-containing protein |
| AT2G44180 | -1.50 | 8.30E-05 | -0.31 | 2.82E-01 | 1.17 | 3.27E-03 | At2g44180 | Methionine aminopeptidase 2 |
| AT2G16870 | -1.50 | 9.75E-05 | -0.57 | 1.73E-01 | 0.91 | 2.50E-02 | At2g16870 | Disease resistance protein |
| AT3G59570 | -1.50 | 6.01E-04 | -0.89 | 2.11E-02 | 0.60 | 2.26E-01 | At3g59570 | Uncharacterized protein At3g59570/T16L24_120 |

| | | | | | | | | |
|---|---|---|---|---|---|---|---|---|
| AT5G61210 | -1.50 | 4.49E-04 | -0.20 | 4.27E-01 | 1.29 | 2.77E-03 | SNAP33 | SNAP25 homologous protein SNAP33 |
| AT3G05650 | -1.50 | 1.54E-03 | 0.10 | 8.43E-01 | 1.59 | 3.85E-04 | RLP32 | Receptor-like protein 32 |
| AT1G19180 | -1.51 | 2.71E-02 | 0.93 | 8.44E-02 | 2.43 | 1.34E-08 | JAZ1 | Protein TIFY |
| AT1G09970 | -1.51 | 6.45E-03 | -0.06 | 9.38E-01 | 1.44 | 4.80E-03 | At1g09970 | Protein kinase domain-containing protein |
| AT1G60960 | -1.51 | 9.15E-06 | -0.08 | 8.93E-01 | 1.41 | 4.02E-06 | IRT3 | Iron regulated transporter 3 |
| AT3G09810 | -1.51 | 3.23E-03 | -0.26 | 5.97E-01 | 1.23 | 2.08E-02 | F11F8_40 | Putative dehydrogenase |
| AT1G13750 | -1.51 | 6.77E-03 | -0.40 | 3.93E-01 | 1.10 | 1.59E-02 | PAP1 | Probable inactive purple acid phosphatase 1 |
| AT2G23810 | -1.51 | 8.92E-03 | 0.08 | 9.20E-01 | 1.58 | 1.01E-03 | TET8 | Tetraspanin-8 |
| AT1G66970 | -1.52 | 2.14E-04 | -0.75 | 1.76E-09 | 0.76 | 9.91E-02 | SVL2 | glycerophosphodiester phosphodiesterase |
| AT4G11300 | -1.52 | 3.16E-04 | -0.63 | 4.11E-01 | 0.87 | 1.05E-01 | At4g11300 | ROH1, |
| AT5G27830 | -1.53 | 9.25E-03 | -0.23 | 7.42E-01 | 1.29 | 2.14E-02 | At5g27830 | Folate receptor family protein |
| AT1G16670 | -1.54 | 3.75E-04 | -0.18 | 8.01E-01 | 1.35 | 5.37E-04 | CRPK1 | Cold-responsive protein kinase 1 |
| AT3G46620 | -1.54 | 2.13E-03 | -0.03 | 1.00E+00 | 1.49 | 1.36E-03 | RDUF1 | E3 ubiquitin-protein ligase RDUF1 |
| AT4G39890 | -1.54 | 6.36E-03 | 0.24 | 8.83E-01 | 1.76 | 1.01E-04 | RABH1C | Ras-related protein RABH1c |
| AT1G61250 | -1.54 | 2.03E-04 | -0.15 | 7.46E-01 | 1.37 | 1.46E-03 | SC3 | Secretory carrier-associated membrane protein |
| AT3G10500 | -1.55 | 1.63E-03 | -0.14 | 8.60E-01 | 1.39 | 4.61E-03 | NAC053 | NAC domain containing protein 53 |
| AT4G29810 | -1.55 | 9.05E-04 | -0.36 | 1.64E-01 | 1.18 | 1.53E-02 | At4g29810 | AT4G29810 protein |
| AT1G21370 | -1.55 | 2.34E-03 | 0.01 | 1.00E+00 | 1.53 | 3.05E-03 | At1g21370 | |
| AT5G48570 | -1.55 | 1.58E-02 | -0.05 | 9.94E-01 | 1.50 | 2.43E-02 | FKBP65 | Peptidyl-prolyl cis-trans isomerase FKBP65 |
| AT1G16000 | -1.55 | 2.65E-02 | -1.48 | 1.51E-02 | 0.06 | 9.66E-01 | At1g16000 | GAG1At protein |
| AT1G27770 | -1.55 | 1.69E-04 | -0.27 | 4.62E-01 | 1.27 | 2.82E-04 | ACA1 | Calcium-transporting ATPase |
| AT4G04570 | -1.56 | 1.06E-02 | -0.40 | 4.66E-02 | 1.14 | 6.30E-02 | CRK40 | Cysteine-rich RLK |
| AT5G13190 | -1.56 | 4.39E-03 | 0.24 | 5.99E-01 | 1.78 | 4.70E-04 | GILP | GSH-induced LITAF domain protein |
| AT4G20830 | -1.56 | 1.88E-02 | -0.24 | 9.38E-01 | 1.28 | 6.92E-02 | At4g20830 | Berberine bridge enzyme-like 19 |
| AT1G69610 | -1.56 | 3.02E-02 | -0.78 | 1.22E-01 | 0.78 | 3.01E-01 | T6C23.19 | Uncharacterized protein F24J1.24 |
| AT1G19960 | -1.56 | 1.66E-03 | -0.77 | 9.93E-03 | 0.78 | 1.53E-01 | At1g19960 | Uncharacterized protein |
| AT1G16500 | -1.57 | 4.74E-02 | -0.45 | 4.97E-01 | 1.10 | 1.90E-01 | At1g16500 | Uncharacterized protein |
| AT3G03470 | -1.57 | 4.26E-02 | 0.05 | 9.99E-01 | 1.60 | 8.75E-02 | CYP89A9 | Cytochrome P450 89A9 |
| AT3G50470 | -1.57 | 4.09E-04 | -0.12 | 9.55E-01 | 1.44 | 1.01E-03 | HR3 | RPW8-like protein 3 |
| AT5G04720 | -1.57 | 5.01E-04 | -0.27 | 3.42E-01 | 1.29 | 4.13E-03 | At5g04720 | Probable disease resistance protein At5g04720 |
| AT5G01850 | -1.57 | 1.73E-03 | -0.04 | 9.88E-01 | 1.51 | 3.29E-03 | At5g01850 | Protein kinase ATN1-like protein |
| AT3G09830 | -1.57 | 4.32E-04 | -0.42 | 1.99E-01 | 1.15 | 8.05E-03 | PCRK1 | Serine/threonine-protein kinase PCRK1 |
| AT4G36150 | -1.57 | 1.22E-05 | -0.86 | 1.11E-05 | 0.69 | 9.01E-02 | At4g36150 | ADP-ribosyl cyclase/cyclic ADP-ribose hydrolase |
| AT4G08950 | -1.57 | 1.17E-02 | -0.33 | 6.94E-01 | 1.22 | 1.03E-05 | At4g08950 | EXO |
| AT3G16565 | -1.57 | 1.48E-03 | 0.37 | 6.18E-01 | 1.93 | 1.43E-04 | At3g16565 | Threonyl and alanyl tRNA synthetase second additional domain-containing protein |
| AT2G25000 | -1.57 | 3.24E-02 | 1.48 | 1.78E-02 | 3.05 | 1.10E-06 | WRKY60 | WRKY DNA-binding protein 60 |
| AT4G18880 | -1.58 | 2.15E-04 | -0.17 | 7.12E-01 | 1.38 | 1.26E-03 | HSFA4A | Heat stress transcription factor A-4a |
| AT5G24810 | -1.58 | 7.50E-05 | -0.39 | 2.50E-02 | 1.18 | 4.36E-03 | At5g24810 | ABC1 family protein |
| AT4G35985 | -1.58 | 3.47E-03 | 0.22 | 8.66E-01 | 1.79 | 1.46E-03 | At4g35985 | Senescence/dehydration-associated protein-like protein |
| AT5G04930 | -1.58 | 2.17E-04 | -0.28 | 9.32E-02 | 1.29 | 3.03E-03 | ALA1 | Phospholipid-transporting ATPase 1 |
| AT4G26060 | -1.58 | 4.44E-03 | -0.24 | 8.36E-01 | 1.34 | 5.92E-03 | At4g26060 | At4g26060 |
| AT3G19250 | -1.58 | 1.43E-02 | -0.91 | 5.85E-02 | 0.65 | 3.87E-01 | At3g19250 | UPF0496 protein At3g19250 |
| AT3G19970 | -1.59 | 2.74E-02 | -0.03 | 1.00E+00 | 1.56 | 2.99E-02 | At3g19970 | |
| AT2G37630 | -1.59 | 1.38E-02 | -1.54 | 1.55E-02 | 0.03 | 9.97E-01 | AS1 | Transcription factor AS1 |
| AT4G14420 | -1.59 | 3.24E-02 | -0.21 | 8.10E-01 | 1.36 | 8.51E-02 | At4g14420 | |
| AT4G26070 | -1.59 | 7.27E-05 | -0.77 | 1.29E-03 | 0.82 | 5.11E-02 | MEK1 | MAP kinase/ ERK kinase 1 |
| AT2G39710 | -1.59 | 2.20E-02 | -0.54 | 1.55E-01 | 1.04 | 1.20E-01 | At2g39710 | Eukaryotic aspartyl protease family protein |
| AT1G52855 | -1.59 | 1.45E-02 | -0.98 | 7.11E-02 | 0.58 | 4.58E-01 | At1g52855 | Uncharacterized protein |
| AT2G31020 | -1.60 | 9.35E-04 | -0.36 | 5.09E-01 | 1.23 | 4.70E-03 | ORP1A | OSBP |
| AT4G11350 | -1.60 | 2.19E-02 | 0.28 | 9.14E-01 | 1.85 | 6.66E-03 | At4g11350 | Transferring glycosyl group transferase |
| AT1G65040 | -1.60 | 1.75E-02 | -0.19 | 7.60E-01 | 1.41 | 2.65E-02 | HRD1B | RING/U-box superfamily protein |
| AT1G64610 | -1.60 | 2.73E-04 | 0.06 | 9.68E-01 | 1.64 | 2.82E-04 | At1g64610 | Transducin/WD40 repeat-like superfamily protein |
| AT1G30620 | -1.60 | 3.18E-03 | 0.33 | 7.38E-01 | 1.92 | 1.30E-03 | MUR4 | NAD |
| AT3G13560 | -1.61 | 2.64E-02 | -0.47 | 1.10E-01 | 1.11 | 1.39E-01 | At3g13560 | glucan endo-1,3-beta-D-glucosidase |
| AT3G07720 | -1.61 | 1.05E-02 | -0.11 | 9.50E-01 | 1.47 | 2.30E-02 | F17A17.6 | AT3g07720/F17A17_6 |
| AT1G01010 | -1.61 | 1.07E-02 | 0.17 | 9.38E-01 | 1.77 | 1.27E-02 | NAC001 | NAC domain-containing protein 1 |
| AT1G25400 | -1.61 | 2.54E-02 | 0.72 | 5.46E-01 | 2.32 | 1.48E-04 | At1g25400 | Transmembrane protein |
| AT4G35600 | -1.61 | 3.62E-04 | -0.40 | 1.04E-01 | 1.19 | 1.42E-02 | CST | Protein kinase superfamily protein |
| AT4G23470 | -1.62 | 2.63E-04 | -0.35 | 2.62E-02 | 1.25 | 6.70E-03 | At4g23470 | PLAC8 family protein |
| AT5G54710 | -1.62 | 7.25E-04 | -0.07 | 8.82E-01 | 1.53 | 9.65E-04 | At5g54710 | Similarity to ankyrin-like protein |
| AT1G60730 | -1.62 | 5.53E-03 | 0.02 | 1.00E+00 | 1.62 | 2.50E-02 | At1g60730 | NAD |
| AT4G23200 | -1.62 | 4.57E-03 | -0.43 | 2.96E-01 | 1.18 | 4.02E-02 | CRK12 | Cysteine-rich RLK |
| AT4G14610 | -1.62 | 2.81E-04 | -0.52 | 2.68E-01 | 1.09 | 3.70E-03 | At4g14610 | Probable disease resistance protein At4g14610 |
| AT5G54490 | -1.62 | 2.67E-03 | 0.23 | 9.25E-01 | 1.86 | 7.45E-04 | PBP1 | Calcium-binding protein PBP1 |
| AT4G22980 | -1.63 | 2.37E-03 | -0.39 | 5.61E-01 | 1.22 | 4.18E-02 | At4g22980 | Molybdenum cofactor sulfurase-like protein |
| AT5G48540 | -1.63 | 1.03E-04 | -0.62 | 1.05E-02 | 1.00 | 3.21E-02 | CRRSP55 | Cysteine-rich repeat secretory protein 55 |
| AT5G47070 | -1.63 | 2.03E-05 | -0.29 | 4.30E-01 | 1.32 | 8.05E-04 | PBL19 | Probable serine/threonine-protein kinase PBL19 |
| AT4G23130 | -1.63 | 2.83E-06 | -0.60 | 4.49E-02 | 1.03 | 1.87E-04 | At4g23130 | Uncharacterized protein |
| AT3G15350 | -1.63 | 2.81E-02 | -0.35 | 2.14E-01 | 1.27 | 9.28E-02 | At3g15350 | |
| AT5G66630 | -1.64 | 6.18E-03 | 0.24 | 7.09E-01 | 1.85 | 6.56E-04 | DAR5 | Protein DA1-related 5 |
| AT4G01540 | -1.64 | 1.90E-02 | -0.04 | 9.93E-01 | 1.60 | 1.82E-02 | NTM1 | NAC with transmembrane motif1 |
| AT3G45290 | -1.64 | 1.29E-04 | -0.23 | 6.63E-01 | 1.39 | 2.64E-03 | MLO3 | MLO-like protein 3 |
| AT2G33020 | -1.64 | 2.17E-02 | -0.66 | 4.87E-01 | 0.97 | 1.84E-01 | RLP24 | Receptor like protein 24 |
| AT5G45510 | -1.64 | 1.68E-04 | -0.53 | 1.51E-04 | 1.10 | 1.56E-02 | At5g45510 | Probable disease resistance protein At5g45510 |
| AT2G03510 | -1.64 | 1.88E-03 | -0.60 | 1.26E-03 | 1.02 | 8.51E-02 | At2g03510 | Expressed protein |
| AT4G33360 | -1.64 | 1.46E-03 | -0.65 | 2.17E-01 | 0.98 | 2.90E-02 | FLDH | NAD |
| AT3G50950 | -1.64 | 6.83E-05 | -0.24 | 4.80E-01 | 1.39 | 1.70E-04 | RPP13L4 | Disease resistance RPP13-like protein 4 |
| AT4G25900 | -1.64 | 1.44E-02 | 0.03 | 9.73E-01 | 1.66 | 1.07E-02 | At4g25900 | glucose-6-phosphate 1-epimerase |
| AT1G14370 | -1.64 | 4.35E-03 | 0.13 | 9.31E-01 | 1.76 | 7.84E-05 | PBL2 | Probable serine/threonine-protein kinase PBL2 |

| | | | | | | | | |
|---|---|---|---|---|---|---|---|---|
| AT2G21188 | -1.65 | 9.74E-03 | 0.08 | 9.80E-01 | 1.73 | 2.12E-03 | | |
| AT3G00720 | -1.65 | 1.86E-02 | -0.26 | 9.39E-01 | 1.38 | 1.08E-01 | | |
| AT4G24570 | -1.65 | 1.29E-02 | -0.23 | 6.80E-01 | 1.41 | 2.48E-02 | PUMP4 | Mitochondrial uncoupling protein 4 |
| AT3G25780 | -1.66 | 2.63E-02 | 1.06 | 2.10E-01 | 2.71 | 6.98E-05 | AOC3 | Allene oxide cyclase 3, chloroplastic |
| AT1G71100 | -1.66 | 1.99E-03 | -0.21 | 7.85E-01 | 1.43 | 1.37E-02 | RPI1 | Probable ribose-5-phosphate isomerase 1 |
| AT3G11840 | -1.66 | 6.90E-05 | -0.35 | 4.27E-01 | 1.29 | 2.15E-03 | At3g11840 | U-box domain-containing protein |
| AT5G58620 | -1.66 | 2.50E-02 | 0.44 | 5.62E-01 | 2.09 | 2.95E-03 | At5g58620 | Zinc finger CCCH domain-containing protein 66 |
| AT5G64660 | -1.67 | 2.98E-03 | -0.01 | 9.97E-01 | 1.70 | 9.38E-04 | PUB27 | U-box domain-containing protein 27 |
| AT4G34131 | -1.67 | 2.48E-02 | -0.01 | 9.97E-01 | 1.63 | 1.37E-02 | UGT73B3 | UDP-glycosyltransferase 73B3 |
| AT3G14470 | -1.67 | 5.69E-04 | -0.66 | 2.28E-02 | 1.01 | 3.84E-02 | RPPL1 | Putative disease resistance RPP13-like protein 1 |
| AT1G14010 | -1.67 | 1.29E-02 | -0.33 | 5.90E-01 | 1.31 | 6.01E-02 | At1g14010 | Transmembrane emp24 domain-containing protein p24delta7 |
| AT1G03210 | -1.67 | 2.16E-03 | 0.33 | 7.25E-01 | 1.99 | 2.39E-04 | At1g03210 | Uncharacterized protein |
| AT1G42990 | -1.67 | 1.14E-03 | -0.47 | 1.39E-01 | 1.20 | 2.11E-02 | BZIP60 | bZIP transcription factor 60 |
| AT3G16990 | -1.68 | 1.50E-02 | 0.05 | 9.93E-01 | 1.68 | 2.69E-02 | TENA_E | Bifunctional TENA-E protein |
| AT1G61640 | -1.68 | 1.03E-02 | -0.89 | 1.80E-01 | 0.78 | 2.50E-01 | T25B24.1 | T25B24.1 protein |
| AT4G02420 | -1.68 | 1.09E-04 | -1.07 | 2.62E-14 | 0.59 | 2.65E-01 | LECRK44 | L-type lectin-domain containing receptor kinase IV.4 |
| AT5G50460 | -1.68 | 3.48E-03 | -0.36 | 2.24E-01 | 1.30 | 2.74E-02 | At5g50460 | Protein transport protein Sec61 subunit gamma |
| AT3G44400 | -1.68 | 1.53E-05 | -0.27 | 3.94E-01 | 1.40 | 3.09E-04 | T22K7_80 | ADP-ribosyl cyclase/cyclic ADP-ribose hydrolase |
| AT4G24970 | -1.68 | 1.63E-04 | -1.10 | 9.93E-03 | 0.58 | 2.00E-01 | MORC7 | Protein MICRORCHIDIA 7 |
| AT1G80820 | -1.68 | 3.42E-03 | 0.28 | 9.10E-01 | 1.95 | 2.00E-03 | CCR2 | Cinnamoyl coa reductase |
| AT1G68710 | -1.69 | 2.67E-04 | -0.28 | 4.93E-01 | 1.39 | 2.95E-03 | At1g68710 | Phospholipid-transporting ATPase |
| AT1G71910 | -1.69 | 2.49E-03 | -0.01 | 1.00E+00 | 1.67 | 1.10E-02 | At1g71910 | |
| AT1G02220 | -1.69 | 8.55E-03 | -0.08 | 9.72E-01 | 1.58 | 9.71E-03 | NAC003 | NAC domain-containing protein 3 |
| AT1G69720 | -1.69 | 6.79E-04 | 0.08 | 9.71E-01 | 1.76 | 1.60E-04 | HO3 | Heme oxygenase 3, chloroplastic |
| AT3G16050 | -1.69 | 4.53E-02 | -0.63 | 7.24E-01 | 1.03 | 2.61E-01 | | |
| AT1G17745 | -1.69 | 3.47E-02 | 0.16 | 9.99E-01 | 1.85 | 6.82E-02 | PGDH2 | D-3-phosphoglycerate dehydrogenase 2, chloroplastic |
| AT3G49120 | -1.69 | 8.21E-03 | -0.03 | 9.78E-01 | 1.65 | 4.96E-03 | PER34 | Peroxidase 34 |
| AT3G13437 | -1.69 | 1.50E-04 | -0.85 | 9.89E-02 | 0.83 | 5.86E-02 | EWR1 | Transmembrane protein |
| AT3G63390 | -1.69 | 1.42E-03 | 0.21 | 8.38E-01 | 1.90 | 3.52E-04 | MAA21_20 | Uncharacterized protein MAA21_20 |
| AT5G44820 | -1.69 | 3.35E-06 | -0.74 | 7.34E-03 | 0.94 | 2.70E-02 | At5g44820 | Nucleotide-diphospho-sugar transferase domain-containing protein |
| AT3G26220 | -1.69 | 4.19E-05 | 0.03 | 9.89E-01 | 1.71 | 9.56E-06 | At3g26220 | At3g26230 |
| AT4G23190 | -1.69 | 5.52E-04 | -0.78 | 1.22E-02 | 0.91 | 5.64E-02 | At4g23190 | Uncharacterized protein |
| AT5G41160 | -1.70 | 8.54E-04 | -0.49 | 7.76E-01 | 1.20 | 6.27E-03 | PUP12 | Probable purine permease |
| AT5G42050 | -1.70 | 1.36E-02 | -0.12 | 7.29E-01 | 1.56 | 1.84E-02 | NRP | DCD domain-containing protein NRP |
| AT5G56870 | -1.70 | 2.15E-04 | 0.00 | 1.00E+00 | 1.68 | 1.72E-04 | BGAL4 | Beta-galactosidase 4 |
| AT2G25110 | -1.70 | 7.81E-03 | -0.21 | 4.65E-01 | 1.48 | 2.11E-02 | SDF2 | Stromal cell-derived factor 2-like protein |
| AT1G49050 | -1.71 | 7.73E-05 | -0.39 | 9.14E-02 | 1.32 | 2.21E-03 | APCB1 | Eukaryotic aspartyl protease family protein |
| AT2G45510 | -1.72 | 4.61E-04 | 0.48 | 3.95E-01 | 2.19 | 4.87E-05 | CYP704A2 | At2g45510 |
| AT4G34150 | -1.72 | 9.02E-04 | -0.17 | 5.64E-01 | 1.54 | 2.05E-03 | At4g34150 | |
| AT2G16700 | -1.73 | 3.35E-03 | -0.31 | 6.51E-01 | 1.41 | 2.59E-02 | ADF5 | Actin depolymerizing factor 5 |
| AT5G44390 | -1.73 | 9.99E-03 | -0.03 | 1.00E+00 | 1.69 | 3.03E-02 | At5g44390 | Berberine bridge enzyme-like 25 |
| AT1G05570 | -1.73 | 3.36E-05 | 0.04 | 9.38E-01 | 1.76 | 1.71E-05 | CALS1 | 1,3-beta-glucan synthase |
| AT2G19130 | -1.73 | 7.27E-03 | 0.01 | 1.00E+00 | 1.73 | 4.13E-03 | At2g19130 | G-type lectin S-receptor-like serine/threonine-protein kinase At2g19130 |
| AT3G44720 | -1.73 | 2.15E-04 | 0.04 | 9.65E-01 | 1.76 | 1.27E-04 | ADT4 | Arogenate dehydratase 4, chloroplastic |
| AT5G46230 | -1.74 | 1.57E-02 | -0.21 | 8.55E-01 | 1.53 | 3.15E-02 | At5g46230 | |
| AT5G19875 | -1.74 | 1.02E-02 | 0.05 | 9.91E-01 | 1.77 | 9.05E-03 | At5g19875 | |
| AT3G56030 | -1.74 | 1.11E-02 | -0.79 | 1.80E-01 | 0.93 | 2.17E-01 | At3g56030 | Pentatricopeptide repeat-containing protein At3g56030, mitochondrial |
| AT3G47540 | -1.75 | 5.91E-03 | -0.39 | 6.00E-01 | 1.34 | 1.33E-02 | At3g47540 | Putative endochitinase |
| AT5G62180 | -1.75 | 4.77E-03 | -0.35 | 8.91E-01 | 1.38 | 3.80E-02 | CXE20 | Probable carboxylesterase 120 |
| AT1G17600 | -1.75 | 2.35E-06 | -0.86 | 2.14E-02 | 0.88 | 5.86E-02 | SOC3 | Disease resistance protein |
| AT3G55950 | -1.75 | 1.25E-03 | -0.41 | 2.85E-01 | 1.32 | 1.31E-02 | CCR3 | Putative serine/threonine-protein kinase-like protein CCR3 |
| AT1G30755 | -1.76 | 2.94E-03 | -0.24 | 6.90E-01 | 1.52 | 8.21E-03 | T5I8.20 | T5I8.20 protein |
| AT1G23840 | -1.76 | 7.86E-04 | -0.31 | 3.64E-01 | 1.43 | 6.17E-03 | At1g23840 | Transmembrane protein |
| AT3G03640 | -1.76 | 3.38E-03 | -0.77 | 8.89E-04 | 0.98 | 1.37E-01 | BGLU25 | Probable inactive beta-glucosidase 25 |
| AT5G12930 | -1.76 | 1.53E-02 | -0.55 | 5.61E-01 | 1.19 | 1.84E-01 | At5g12930 | Inactive rhomboid protein |
| AT3G57630 | -1.76 | 5.18E-04 | -0.75 | 9.40E-03 | 1.01 | 6.74E-02 | EXAD | At3g57630 |
| AT2G26190 | -1.76 | 1.54E-05 | -0.67 | 2.49E-05 | 1.08 | 1.08E-02 | IQM4 | Calmodulin-binding family protein |
| AT5G46520 | -1.76 | 5.09E-04 | 0.17 | 9.20E-01 | 1.91 | 1.23E-04 | VICTR | ADP-ribosyl cyclase/cyclic ADP-ribose hydrolase |
| AT5G16170 | -1.77 | 2.01E-06 | -0.28 | 5.17E-01 | 1.48 | 8.91E-05 | T21H19_90 | Uncharacterized protein T21H19_90 |
| AT1G21390 | -1.77 | 8.16E-03 | 0.32 | 8.28E-01 | 2.08 | 2.05E-03 | At1g21390 | Transmembrane protein |
| AT1G51760 | -1.77 | 3.17E-02 | 0.51 | 5.82E-02 | 2.26 | 2.47E-03 | ILL4 | IAA-amino acid hydrolase ILR1-like 4 |
| AT5G05460 | -1.77 | 1.14E-06 | -0.55 | 5.82E-02 | 1.21 | 4.44E-04 | ENGASE1 | Cytosolic endo-beta-N-acetylglucosaminidase 1 |
| AT5G61250 | -1.77 | 6.64E-04 | -0.10 | 9.65E-01 | 1.66 | 1.21E-03 | At5g61250 | Heparanase-like protein 2 |
| AT3G13910 | -1.78 | 6.31E-03 | 0.17 | 9.56E-01 | 1.92 | 2.53E-02 | At3g13910 | |
| AT1G72520 | -1.78 | 4.44E-05 | 0.16 | 8.46E-01 | 1.93 | 9.39E-13 | LOX4 | Lipoxygenase 4, chloroplastic |
| AT3G49780 | -1.78 | 1.47E-02 | -0.16 | 8.03E-01 | 1.61 | 1.85E-02 | PSK3 | Phytosulfokines 3 |
| AT3G26170 | -1.78 | 1.48E-05 | -0.19 | 7.09E-01 | 1.58 | 1.42E-05 | CYP71B19 | Cytochrome P450 71B19 |
| AT3G14620 | -1.78 | 8.35E-04 | 0.26 | 6.35E-01 | 2.03 | 1.10E-05 | CYP72A8 | AT3g14620/MIE1_12 |
| AT5G20960 | -1.79 | 1.47E-03 | 0.44 | 4.09E-01 | 2.21 | 3.87E-05 | AAO1 | Indole-3-acetaldehyde oxidase |
| AT2G46620 | -1.79 | 2.15E-03 | -0.01 | 1.00E+00 | 1.76 | 2.59E-03 | At2g46620 | AAA-ATPase At2g46620 |
| AT1G77500 | -1.79 | 6.31E-04 | -0.33 | 2.90E-01 | 1.44 | 7.46E-03 | At1g77500 | Uncharacterized protein |
| AT5G07340 | -1.79 | 2.88E-02 | -0.09 | 9.34E-01 | 1.69 | 3.09E-02 | At5g07340 | Calreticulin family protein |
| AT2G24600 | -1.79 | 2.86E-05 | -0.24 | 6.38E-01 | 1.54 | 1.16E-05 | At2g24600 | Ankyrin repeat family protein |
| AT5G40720 | -1.79 | 1.62E-02 | -0.17 | 8.46E-01 | 1.60 | 1.54E-02 | MNF13.28 | Glycosyltransferase family 92 protein |
| AT1G21900 | -1.79 | 1.70E-02 | -0.14 | 8.95E-01 | 1.64 | 1.66E-02 | At1g21900 | Transmembrane emp24 domain-containing protein p24delta5 |
| AT3G60580 | -1.80 | 1.85E-02 | -1.14 | 4.66E-01 | 0.71 | 3.97E-01 | | |
| AT3G26600 | -1.80 | 2.30E-05 | -0.42 | 2.45E-01 | 1.37 | 2.59E-03 | At3g26600 | |
| AT5G24210 | -1.80 | 6.81E-05 | -0.26 | 4.82E-01 | 1.53 | 1.16E-04 | MOP9.2 | Alpha/beta-Hydrolases superfamily protein |
| AT2G28400 | -1.80 | 1.15E-02 | | | 2.08 | 1.01E-02 | S40-4 | Protein S40-4 |

| | | | | | | | | |
|---|---|---|---|---|---|---|---|---|
| AT1G61260 | -1.81 | 2.91E-03 | 1.15 | 2.67E-01 | 2.96 | 2.77E-03 | F11P17.2 | F11P17.2 protein |
| AT3G11080 | -1.81 | 2.70E-03 | -0.06 | 9.42E-01 | 1.73 | 4.61E-03 | RLP35 | Receptor-like protein 35 |
| AT2G41100 | -1.82 | 1.14E-07 | -0.60 | 1.44E-02 | 1.20 | 8.91E-05 | TCH3 | Calcium-binding EF hand family protein |
| AT3G20600 | -1.82 | 1.93E-05 | -0.67 | 4.83E-03 | 1.14 | 9.14E-03 | NDR1 | Protein NDR1 |
| AT3G14280 | -1.82 | 1.32E-02 | 0.48 | 4.70E-01 | 2.29 | 8.36E-04 | At3g14280 | AT3G14280 protein |
| AT3G09520 | -1.83 | 1.49E-02 | -1.14 | 1.41E-01 | 0.68 | 4.64E-01 | At3g09520 | Exocyst subunit Exo70 family protein |
| AT1G53100 | -1.83 | 3.70E-02 | -0.13 | 9.70E-01 | 1.66 | 6.03E-02 | At1g53100 | Core-2/I-branching beta-1,6-N-acetylglucosaminyltransferase family protein |
| AT1G18210 | -1.84 | 2.26E-04 | -0.14 | 8.60E-01 | 1.68 | 1.16E-03 | CML27 | Probable calcium-binding protein CML27 |
| AT1G31580 | -1.84 | 5.73E-05 | -0.54 | 5.27E-03 | 1.29 | 3.03E-03 | ECS1 | Protein ECS1 |
| AT3G26840 | -1.85 | 2.30E-02 | 0.49 | 5.89E-01 | 2.32 | 1.27E-03 | PES2 | Phytyl ester synthase 2, chloroplastic |
| AT1G67360 | -1.85 | 8.86E-03 | 0.17 | 8.47E-01 | 2.00 | 3.43E-03 | At1g67360 | REF/SRPP-like protein At1g67360 |
| AT1G55910 | -1.85 | 1.67E-04 | -0.68 | 1.98E-03 | 1.14 | 3.84E-02 | ZIP11 | Zinc transporter 11 |
| AT3G51430 | -1.85 | 1.37E-03 | -0.28 | 4.87E-01 | 1.56 | 4.35E-03 | YLS2 | Calcium-dependent phosphotriesterase superfamily protein |
| AT5G26340 | -1.85 | 4.64E-04 | 0.06 | 9.09E-01 | 1.89 | 2.95E-04 | STP13 | Sugar transport protein 13 |
| AT4G28390 | -1.85 | 3.06E-03 | -0.11 | 8.89E-01 | 1.73 | 7.39E-03 | AAC3 | ADP,ATP carrier protein 3, mitochondrial |
| AT5G07760 | -1.85 | 6.61E-04 | -0.71 | 2.95E-01 | 1.13 | 3.61E-02 | MBK20.22 | Formin-like protein |
| AT1G65845 | -1.85 | 4.56E-03 | -0.14 | 8.15E-01 | 1.71 | 8.93E-03 | At1g65845 | At1g65844 |
| AT1G72950 | -1.86 | 1.43E-03 | | | 1.78 | 1.87E-02 | At1g72950 | TIR domain-containing protein |
| AT2G36580 | -1.86 | 9.89E-04 | -0.43 | 7.92E-02 | 1.42 | 1.66E-02 | At2g36580 | Pyruvate kinase |
| AT2G13790 | -1.86 | 8.61E-07 | -0.61 | 3.28E-05 | 1.23 | 1.64E-03 | SERK4 | Somatic embryogenesis receptor kinase 4 |
| AT2G29990 | -1.86 | 6.17E-04 | 0.02 | 1.00E+00 | 1.87 | 6.55E-04 | NDA2 | Internal alternative NAD |
| AT4G36090 | -1.86 | 3.19E-05 | -0.42 | 4.83E-01 | 1.43 | 4.12E-03 | ALKBH9C | Oxidoreductase, 2OG-Fe |
| AT5G20400 | -1.87 | 1.53E-02 | 0.38 | 8.31E-01 | 2.24 | 2.35E-03 | At5g20400 | hypothetical protein |
| AT3G60450 | -1.87 | 2.93E-03 | 0.27 | 8.40E-01 | 2.13 | 2.10E-04 | At3g60450 | |
| AT3G29240 | -1.87 | 6.57E-04 | -0.22 | 4.64E-01 | 1.63 | 3.25E-03 | At3g29240 | |
| AT4G24920 | -1.87 | 1.38E-02 | -0.31 | 2.75E-01 | 1.55 | 3.98E-02 | At4g24920 | SecE/sec61-gamma protein transport protein |
| AT1G09240 | -1.87 | 4.97E-02 | -1.27 | 1.28E-02 | 0.59 | 5.60E-01 | NAS3 | Nicotianamine synthase 3 |
| AT1G68690 | -1.88 | 2.90E-02 | 0.09 | 9.25E-01 | 1.94 | 2.00E-02 | PERK9 | non-specific serine/threonine protein kinase |
| AT5G66910 | -1.88 | 2.86E-06 | -0.64 | 3.67E-02 | 1.22 | 3.41E-03 | At5g66910 | Probable disease resistance protein At5g66910 |
| AT5G07770 | -1.88 | 2.64E-08 | -0.29 | 3.34E-01 | 1.58 | 3.04E-06 | FH16 | Formin-like protein |
| AT5G46500 | -1.88 | 2.11E-05 | -0.22 | 7.57E-01 | 1.65 | 2.15E-04 | At5g46500 | Protein VARIATION IN COMPOUND TRIGGERED ROOT growth protein |
| AT3G17420 | -1.89 | 1.21E-02 | -0.09 | 9.60E-01 | 1.78 | 2.03E-02 | At3g17420 | Serine/threonine protein kinase |
| AT3G08870 | -1.89 | 9.88E-07 | -0.94 | 4.46E-05 | 0.94 | 1.50E-02 | LECRK-VI1 | Concanavalin A-like lectin protein kinase family protein |
| AT2G41090 | -1.89 | 3.20E-05 | -0.96 | 6.71E-04 | 0.92 | 6.87E-03 | At2g41090 | AT2G41090 protein |
| AT2G35910 | -1.89 | 9.53E-03 | -0.36 | 8.32E-01 | 1.51 | 3.38E-02 | At2g35910 | |
| AT3G51890 | -1.89 | 9.69E-05 | -0.12 | 9.14E-01 | 1.75 | 4.66E-04 | At3g51890 | Clathrin light chain |
| AT4G37640 | -1.90 | 6.26E-05 | -0.26 | 3.85E-01 | 1.62 | 8.47E-04 | ACA2 | Calcium-transporting ATPase 2, plasma membrane-type |
| AT2G41835 | -1.90 | 2.65E-04 | -0.44 | 3.78E-01 | 1.44 | 1.06E-02 | SAP11 | Zinc finger AN1 and C2H2 domain-containing stress-associated protein 11 |
| AT1G61340 | -1.90 | 2.70E-03 | | | 2.42 | 1.11E-02 | At1g61340 | FBS1 |
| AT5G39090 | -1.90 | 9.12E-04 | 0.84 | 2.20E-01 | 2.72 | 2.78E-06 | At5g39090 | Uncharacterized protein |
| AT4G08850 | -1.90 | 1.63E-04 | -0.58 | 1.04E-04 | 1.31 | 1.13E-02 | MIK2 | MDIS1-interacting receptor like kinase 2 |
| AT3G09440 | -1.90 | 4.05E-03 | -0.57 | 2.18E-04 | 1.32 | 5.66E-02 | HSP70-3 | Heat shock 70 kDa protein 3 |
| AT3G11820 | -1.90 | 4.05E-05 | -0.50 | 2.23E-02 | 1.38 | 5.53E-03 | SYP121 | Syntaxin-121 |
| AT3G59700 | -1.91 | 9.69E-05 | -0.13 | 9.04E-01 | 1.75 | 2.74E-04 | LECRK55 | L-type lectin-domain containing receptor kinase V.5 |
| AT3G47090 | -1.91 | 2.09E-06 | -0.26 | 5.64E-01 | 1.64 | 1.56E-04 | F13I12.140 | Leucine-rich repeat protein kinase family protein |
| AT5G18780 | -1.91 | 8.28E-06 | 0.04 | 9.95E-01 | 1.94 | 1.49E-05 | At5g18780 | F-box/RNI-like superfamily protein |
| AT3G24090 | -1.91 | 3.48E-03 | -0.62 | 2.70E-01 | 1.28 | 8.04E-02 | GFAT1 | Glutamine--fructose-6-phosphate aminotransferase [isomerizing] 1 |
| AT1G34180 | -1.91 | 5.75E-03 | 0.44 | 8.25E-01 | 2.34 | 1.34E-04 | NAC016 | NAC domain containing protein 16 |
| AT1G74440 | -1.92 | 1.62E-06 | -1.00 | 2.34E-04 | 0.90 | 5.21E-02 | At1g74440 | ER membrane protein, putative |
| AT2G46440 | -1.92 | 8.77E-08 | -0.75 | 5.09E-07 | 1.15 | 2.20E-03 | CNGC11 | Cyclic nucleotide-gated channels |
| AT4G23320 | -1.92 | 2.58E-05 | 0.10 | 9.72E-01 | 2.01 | 2.39E-05 | CRK24 | Cysteine-rich RLK |
| AT2G47470 | -1.93 | 4.15E-03 | -0.18 | 5.80E-01 | 1.73 | 1.15E-02 | UNE5 | Thioredoxin family protein |
| AT1G20350 | -1.94 | 1.10E-02 | | | 2.05 | 6.18E-03 | TIM17-1 | Mitochondrial import inner membrane translocase subunit TIM17-1 |
| AT2G41190 | -1.94 | 4.90E-03 | -0.50 | 4.60E-01 | 1.42 | 3.11E-02 | At2g41190 | Transmembrane amino acid transporter family protein |
| AT1G22400 | -1.94 | 4.45E-04 | -0.22 | 7.41E-01 | 1.71 | 9.31E-04 | UGT85A1 | UDP-glycosyltransferase 85A1 |
| AT1G61370 | -1.95 | 2.15E-02 | -1.24 | 8.62E-02 | 0.68 | 4.95E-01 | At1g61370 | S-locus lectin protein kinase family protein |
| AT4G37530 | -1.95 | 1.49E-02 | -0.03 | 9.82E-01 | 1.90 | 1.40E-02 | At4g37530 | Peroxidase |
| AT2G31060 | -1.95 | 2.42E-03 | -0.46 | 1.71E-01 | 1.47 | 2.86E-02 | At2g31060 | Putative GTP-binding protein |
| AT3G28480 | -1.95 | 5.61E-04 | -0.49 | 3.07E-01 | 1.44 | 1.27E-02 | At3g28480 | procollagen-proline 4-dioxygenase |
| AT1G72680 | -1.95 | 7.82E-04 | -0.33 | 6.28E-01 | 1.60 | 6.80E-03 | CAD1 | Probable cinnamyl alcohol dehydrogenase 1 |
| AT4G02220 | -1.96 | 1.81E-05 | -0.69 | 2.03E-02 | 1.25 | 1.35E-02 | T10M13.22 | Putative zinc finger protein |
| AT5G50350 | -1.96 | 3.21E-02 | -0.38 | 7.74E-01 | 1.58 | 8.38E-02 | MXI22.6 | Uncharacterized protein At5g50350 |
| AT2G04430 | -1.96 | 3.24E-04 | -0.80 | 1.68E-01 | 1.15 | 5.84E-02 | NUDT5 | Nudix hydrolase |
| AT5G53550 | -1.97 | 8.80E-06 | -0.31 | 3.82E-01 | 1.64 | 2.42E-04 | YSL3 | YELLOW STRIPE like 3 |
| AT3G51340 | -1.97 | 1.28E-04 | -1.24 | 2.54E-02 | 0.72 | 2.24E-01 | F24M12.380 | Uncharacterized protein F24M12.380 |
| AT1G11340 | -1.97 | 2.38E-07 | -0.68 | 4.09E-01 | 1.28 | 4.10E-03 | At1g11340 | Receptor-like serine/threonine-protein kinase |
| AT1G10040 | -1.98 | 4.86E-02 | | | 2.64 | 1.04E-02 | At1g10040 | Alpha/beta-Hydrolases superfamily protein |
| AT1G59590 | -1.98 | 1.39E-05 | -0.37 | 4.51E-01 | 1.60 | 5.58E-04 | ZCF37 | At1g59590 |
| AT4G22305 | -1.98 | 1.16E-05 | -1.26 | 1.32E-08 | 0.72 | 1.80E-01 | SOBER1 | Alpha/beta-Hydrolases superfamily protein |
| AT5G22060 | -1.98 | 5.96E-05 | -0.45 | 6.66E-02 | 1.52 | 3.29E-03 | ATJ2 | Chaperone protein dnaJ 2 |
| AT2G20110 | -1.98 | 1.34E-03 | -0.09 | 9.76E-01 | 1.89 | 2.17E-03 | At2g20110 | Tesmin/TSO1-like CXC domain-containing protein |
| AT5G15870 | -1.99 | 4.99E-05 | -0.36 | 1.51E-01 | 1.61 | 1.49E-03 | At5g15870 | glucan endo-1,3-beta-D-glucosidase |
| AT4G34390 | -1.99 | 4.30E-05 | -0.32 | 1.26E-01 | 1.65 | 9.14E-04 | XLG2 | Extra-large GTP-binding protein 2 |
| AT4G26990 | -1.99 | 5.97E-03 | -0.04 | 9.84E-01 | 1.93 | 7.50E-03 | At4g26990 | Ataxin 2 SM domain-containing protein |
| AT4G37690 | -1.99 | 4.93E-03 | -0.23 | 9.44E-01 | 1.74 | 7.38E-03 | GT6 | Glycosyltransferase 6 |
| AT3G07580 | -2.00 | 1.53E-02 | -0.53 | 4.92E-01 | 1.45 | 9.00E-02 | At3g07580 | At3g07580 |
| AT2G38290 | -2.01 | 6.04E-06 | -0.18 | 7.90E-01 | 1.81 | 7.88E-05 | AMT2 | Ammonium transporter 2 |
| AT1G28370 | -2.01 | 2.38E-02 | 0.32 | 8.82E-01 | 2.31 | 2.59E-08 | ERF11 | ERF domain protein 11 |
| AT3G09490 | -2.01 | 1.26E-03 | -0.81 | 8.02E-02 | 1.19 | 7.18E-02 | At3g09490 | Uncharacterized protein |

| | | | | | | | | |
|---|---|---|---|---|---|---|---|---|
| AT5G42530 | -2.01 | 6.18E-06 | -1.01 | 1.29E-04 | 0.99 | 3.29E-03 | At5g42530 | |
| AT1G26690 | -2.02 | 9.12E-04 | -0.29 | 5.28E-01 | 1.72 | 5.81E-03 | At1g26690 | |
| AT5G01640 | -2.02 | 4.16E-03 | 0.18 | 9.58E-01 | 2.19 | 4.70E-03 | PRA1B5 | PRA1 family protein B5 |
| AT4G22530 | -2.02 | 5.92E-04 | -0.38 | 7.77E-01 | 1.64 | 8.50E-04 | At4g22530 | |
| AT4G12290 | -2.03 | 5.80E-05 | -0.10 | 8.97E-01 | 1.91 | 1.16E-04 | CUAO | Amine oxidase |
| AT1G74810 | -2.03 | 5.76E-03 | -1.96 | 1.01E-03 | 0.05 | 9.76E-01 | BOR5 | HCO3-transporter family |
| AT4G34135 | -2.03 | 6.39E-03 | -0.32 | 5.20E-01 | 1.70 | 2.35E-02 | UGT73B2 | UDP-glucosyltransferase 73B2 |
| AT2G23680 | -2.03 | 3.85E-04 | -0.21 | 8.34E-01 | 1.81 | 9.38E-04 | At2g23680 | Uncharacterized protein |
| AT5G47120 | -2.04 | 9.80E-04 | -0.24 | 4.56E-01 | 1.79 | 3.90E-03 | BI-1 | Bax inhibitor 1 |
| AT4G11910 | -2.04 | 4.88E-02 | | | 3.05 | 6.63E-03 | SGR2 | Magnesium dechelatase SGR2, chloroplastic |
| AT1G61460 | -2.04 | 4.37E-03 | -0.75 | 6.82E-01 | 1.34 | 7.29E-02 | At1g61460 | G-type lectin S-receptor-like serine/threonine-protein kinase At1g61460 |
| AT1G53920 | -2.05 | 5.95E-04 | -1.67 | 2.01E-02 | 0.36 | 6.45E-01 | At1g53920 | GLIP5 |
| AT5G25440 | -2.05 | 4.20E-08 | -0.97 | 9.60E-07 | 1.06 | 5.19E-03 | SZE1 | Protein kinase superfamily protein |
| AT1G18910 | -2.05 | 1.19E-04 | -0.71 | 7.40E-02 | 1.32 | 1.41E-02 | At1g18910 | Zinc finger protein BRUTUS-like At1g18910 |
| AT3G20510 | -2.05 | 1.67E-03 | -0.86 | 1.99E-02 | 1.16 | 9.82E-02 | FAX6 | Protein FATTY ACID EXPORT 6 |
| AT4G29780 | -2.05 | 3.29E-04 | 0.75 | 1.13E-01 | 2.78 | 1.58E-07 | F27B13.20 | Uncharacterized protein AT4g29780 |
| AT1G66090 | -2.05 | 1.15E-05 | -0.02 | 1.00E+00 | 2.02 | 4.70E-06 | At1g66090 | Disease resistance protein |
| AT5G40780 | -2.05 | 2.94E-04 | -0.24 | 2.71E-01 | 1.80 | 1.81E-03 | LHT1 | Lysine histidine transporter 1 |
| AT1G56340 | -2.06 | 8.49E-03 | -0.29 | 1.30E-01 | 1.75 | 2.54E-02 | CRT1A | Calreticulin |
| AT1G29240 | -2.06 | 2.39E-04 | -0.52 | 7.27E-02 | 1.53 | 9.44E-03 | At1g29240 | |
| AT1G34750 | -2.07 | 9.58E-07 | -0.50 | 3.04E-01 | 1.55 | 3.40E-04 | CIPP1 | Protein phosphatase 2C family protein |
| AT4G19370 | -2.08 | 5.57E-04 | -1.12 | 3.40E-01 | 0.98 | 9.26E-02 | At4g19370 | Transmembrane protein |
| AT1G45145 | -2.08 | 1.28E-03 | 0.96 | 2.17E-02 | 3.03 | 5.53E-06 | TRX5 | Thioredoxin H5 |
| AT4G38830 | -2.08 | 1.78E-02 | -1.29 | 3.30E-01 | 0.83 | 3.47E-01 | CRK26 | Cysteine-rich receptor-like protein kinase 26 |
| AT3G45620 | -2.08 | 3.41E-04 | -0.57 | 1.00E-01 | 1.50 | 8.34E-03 | At3g45620 | Transducin/WD40 repeat-like superfamily protein |
| AT4G31800 | -2.09 | 4.48E-03 | 0.89 | 1.02E-01 | 2.97 | 2.14E-07 | WRKY18 | WRKY DNA-binding protein 18 |
| AT2G24160 | -2.09 | 1.52E-05 | -0.87 | 7.46E-06 | 1.21 | 1.46E-02 | | |
| AT4G11000 | -2.09 | 6.08E-07 | -1.10 | 8.34E-06 | 0.98 | 3.73E-02 | F25I24.210 | Uncharacterized protein AT4g11000 |
| AT4G13180 | -2.09 | 8.42E-04 | -0.79 | 2.41E-01 | 1.28 | 5.69E-02 | At4g13180 | Uncharacterized protein |
| AT3G26500 | -2.09 | 1.04E-03 | 0.09 | 9.58E-01 | 2.18 | 4.44E-04 | PIRL2 | Plant intracellular Ras-group-related LRR protein 2 |
| AT1G15670 | -2.10 | 1.06E-02 | 0.00 | 1.00E+00 | 2.09 | 1.45E-06 | At1g15670 | F-box/kelch-repeat protein At1g15670 |
| AT5G66640 | -2.11 | 2.74E-05 | -0.33 | 6.73E-01 | 1.77 | 6.86E-04 | DAR3 | DA1-related protein 3 |
| AT1G15790 | -2.11 | 2.77E-05 | -0.29 | 7.07E-01 | 1.80 | 4.66E-04 | At1g15790 | Mediator complex subunit 15 KIX domain-containing protein |
| AT5G06320 | -2.12 | 8.22E-05 | -0.58 | 3.54E-03 | 1.52 | 5.05E-03 | NHL3 | NDR1/HIN1-like protein 3 |
| AT4G22590 | -2.12 | 1.52E-04 | 0.19 | 8.76E-01 | 2.30 | 1.43E-04 | TPPG | Probable trehalose-phosphate phosphatase G |
| AT3G48520 | -2.13 | 3.08E-02 | 0.31 | 8.74E-01 | 2.43 | 5.77E-04 | CYP94B3 | Cytochrome P450 94B3 |
| AT1G33720 | -2.15 | 3.26E-13 | -0.19 | 7.09E-01 | 1.95 | 1.12E-12 | CYP76C6 | Cytochrome P450, family 76, subfamily C, polypeptide 6 |
| AT1G35210 | -2.15 | 2.96E-06 | 0.16 | 8.85E-01 | 2.30 | 5.06E-07 | At1g35210 | At1g35210 |
| AT2G29120 | -2.16 | 1.13E-03 | -0.01 | 1.00E+00 | 2.14 | 1.00E-03 | GLR2.7 | Glutamate receptor 2.7 |
| AT1G09210 | -2.16 | 6.16E-03 | -0.48 | 9.42E-04 | 1.66 | 4.18E-02 | CRT2 | Calreticulin-2 |
| AT5G41740 | -2.16 | 3.92E-08 | -0.18 | 6.89E-01 | 1.97 | 3.45E-08 | MUF8.2 | Disease resistance protein |
| AT1G09560 | -2.16 | 5.50E-04 | -0.42 | 1.20E-02 | 1.73 | 7.37E-03 | GLP4 | Germin-like protein subfamily 2 member 1 |
| AT4G15233 | -2.16 | 1.70E-07 | -0.69 | 2.81E-02 | 1.46 | 6.31E-05 | ABCG42 | ABC-2 and Plant PDR ABC-type transporter family protein |
| AT5G62620 | -2.16 | 3.33E-03 | -0.09 | 9.51E-01 | 2.07 | 4.21E-03 | GALT6 | Galactosyltransferase family protein |
| AT1G51660 | -2.16 | 2.15E-04 | -0.69 | 5.58E-02 | 1.47 | 1.05E-02 | MKK4 | Mitogen-activated protein kinase kinase 4 |
| AT1G14870 | -2.16 | 3.61E-03 | -0.50 | 5.91E-01 | 1.64 | 4.89E-02 | PCR2 | PLANT CADMIUM RESISTANCE 2 |
| AT1G05300 | -2.17 | 1.39E-08 | 0.73 | 1.87E-02 | 2.88 | 2.07E-19 | ZIP5 | Zinc transporter 5 |
| AT4G28490 | -2.17 | 1.45E-05 | -0.80 | 2.53E-06 | 1.36 | 9.78E-03 | RLK5 | Receptor-like protein kinase 5 |
| AT1G10970 | -2.17 | 2.11E-07 | 0.40 | 3.92E-01 | 2.54 | 1.73E-10 | At1g10970 | |
| AT3G29034 | -2.17 | 2.54E-03 | | | 2.86 | 6.89E-04 | At3g29034 | Transmembrane protein |
| AT1G14360 | -2.17 | 9.05E-03 | -0.36 | 7.06E-01 | 1.78 | 2.56E-02 | UTR3 | UDP-galactose/UDP-glucose transporter 3 |
| AT3G22930 | -2.18 | 1.89E-03 | | | 1.91 | 6.99E-02 | | |
| AT5G59662 | -2.18 | 1.07E-02 | | | 1.33 | 6.36E-02 | | |
| AT1G64065 | -2.19 | 1.67E-02 | 0.12 | 9.76E-01 | 2.30 | 3.01E-02 | At1g64065 | Late embryogenesis abundant protein At1g64065 |
| AT5G48380 | -2.20 | 1.63E-04 | -0.19 | 5.02E-01 | 2.00 | 8.12E-04 | At5g48380 | Probably inactive leucine-rich repeat receptor-like protein kinase At5g48380 |
| AT5G39050 | -2.20 | 1.49E-04 | -0.11 | 8.59E-01 | 2.07 | 2.32E-04 | PMAT1 | Phenolic glucoside malonyltransferase 1 |
| AT1G62300 | -2.20 | 5.84E-05 | -0.46 | 4.94E-01 | 1.71 | 2.91E-03 | At1g62300 | Truncated WRKY6 protein |
| AT3G52400 | -2.20 | 1.21E-05 | 0.11 | 8.78E-01 | 2.30 | 2.79E-06 | SYP122 | Syntaxin-122 |
| AT3G50800 | -2.21 | 2.67E-03 | | | 2.05 | 6.03E-02 | | |
| AT4G13820 | -2.21 | 4.21E-03 | -0.15 | 9.51E-01 | 2.03 | 3.41E-04 | At4g13820 | Disease resistance like protein |
| AT4G36990 | -2.22 | 1.03E-03 | 0.25 | 7.42E-01 | 2.45 | 7.12E-04 | At4g36990 | Truncated heat shock factor B1 |
| AT2G40095 | -2.22 | 1.81E-05 | -0.74 | 1.12E-02 | 1.45 | 6.93E-03 | At2g40095 | Alpha/beta hydrolase related protein |
| AT2G37710 | -2.23 | 3.30E-05 | -0.75 | 1.17E-08 | 1.46 | 9.65E-03 | LECRK41 | L-type lectin-domain containing receptor kinase IV.1 |
| AT4G16660 | -2.23 | 8.03E-04 | -0.36 | 6.53E-02 | 1.86 | 5.77E-03 | HSP70 | Heat shock protein 70 |
| AT3G28340 | -2.23 | 1.09E-06 | -0.76 | 7.11E-02 | 1.45 | 5.18E-04 | GATL10 | Probable galacturonosyltransferase-like 10 |
| AT5G10380 | -2.24 | 2.17E-04 | -0.65 | 6.96E-02 | 1.58 | 1.75E-03 | ATL55 | E3 ubiquitin-protein ligase RING1 |
| AT1G76980 | -2.24 | 9.84E-03 | -0.38 | 7.41E-01 | 1.87 | 1.96E-02 | At1g76980 | Patatin-like phospholipase domain protein |
| AT1G21520 | -2.24 | 5.17E-05 | -0.18 | 8.92E-01 | 2.06 | 4.40E-04 | At1g21520 | |
| AT1G56660 | -2.24 | 2.58E-05 | 0.70 | 3.88E-01 | 2.92 | 6.42E-07 | At1g56660 | Uncharacterized protein |
| AT1G65500 | -2.25 | 1.66E-02 | -0.20 | 9.57E-01 | 2.03 | 3.55E-03 | STMP6 | Secreted transmembrane peptide 6 |
| AT4G12720 | -2.25 | 7.21E-05 | -0.05 | 9.64E-01 | 2.19 | 5.02E-05 | NUDT7 | Nudix hydrolase |
| AT3G45860 | -2.26 | 3.68E-06 | -0.29 | 5.45E-01 | 1.96 | 4.19E-07 | CRK4 | Cysteine-rich receptor-like protein kinase 4 |
| AT1G28190 | -2.27 | 4.49E-02 | 0.37 | 7.57E-01 | 2.63 | 7.48E-03 | At1g28190 | |
| AT1G27330 | -2.27 | 1.59E-04 | -0.56 | 5.91E-02 | 1.70 | 7.96E-03 | At1g27330 | F17L21.12 |
| AT4G11850 | -2.27 | 4.51E-06 | -0.62 | 8.07E-03 | 1.65 | 1.04E-03 | PLDGAMMA | Phospholipase D gamma 1 |
| AT4G21380 | -2.27 | 2.55E-06 | -1.11 | 3.70E-05 | 1.15 | 5.34E-03 | RK3 | Receptor kinase 3 |
| AT1G24147 | -2.28 | 5.46E-04 | -0.69 | 2.94E-01 | 1.57 | 1.13E-04 | At1g24147 | Transmembrane protein |
| AT1G67520 | -2.28 | 1.25E-02 | -0.51 | 8.16E-01 | 1.78 | 8.70E-02 | At1g67520 | Receptor-like serine/threonine-protein kinase |
| AT2G31865 | -2.28 | 5.21E-05 | -0.18 | 7.75E-01 | 2.10 | 1.79E-04 | PARG2 | poly |

| | | | | | | | | |
|---|---|---|---|---|---|---|---|---|
| AT1G24150 | -2.28 | 4.99E-05 | -0.96 | 1.10E-08 | 1.31 | 2.86E-02 | FH4 | Formin-like protein |
| AT3G23010 | -2.29 | 9.49E-05 | -0.86 | 2.37E-02 | 1.42 | 5.10E-03 | RLP36 | Receptor like protein |
| AT2G38470 | -2.29 | 6.15E-07 | -0.38 | 8.51E-02 | 1.89 | 1.03E-05 | At2g38470 | Truncated WRKY33 protein |
| AT5G08760 | -2.29 | 1.15E-04 | -0.53 | 3.59E-01 | 1.75 | 2.60E-05 | PROSCOOP15 | Serine rich endogenous peptide 15 |
| AT5G10740 | -2.29 | 1.06E-03 | -0.39 | 8.65E-01 | 1.91 | 8.86E-03 | At5g10740 | Probable protein phosphatase 2C 69 |
| AT5G03160 | -2.29 | 1.94E-03 | -0.75 | 1.51E-02 | 1.54 | 4.15E-02 | P58IPK | DnaJ protein P58IPK homolog |
| AT5G54140 | -2.30 | 4.93E-04 | -1.07 | 8.53E-02 | 1.20 | 7.60E-02 | ILL3 | IAA-amino acid hydrolase ILR1-like 3 |
| AT5G37600 | -2.31 | 1.95E-05 | -0.01 | 1.00E+00 | 2.28 | 1.96E-05 | GLN1-1 | Glutamine synthetase cytosolic isozyme 1-1 |
| AT5G19240 | -2.31 | 2.11E-05 | -0.99 | 1.25E-08 | 1.30 | 1.90E-02 | At5g19240 | Uncharacterized GPI-anchored protein At5g19240 |
| AT1G67000 | -2.31 | 1.71E-04 | 0.45 | 6.40E-01 | 2.76 | 1.10E-06 | LRK10L-2.8 | LEAF RUST 10 DISEASE-RESISTANCE LOCUS RECEPTOR-LIKE PROTEIN KINASE-like 2.8 |
| AT4G26270 | -2.31 | 7.40E-05 | -0.24 | 8.26E-01 | 2.07 | 6.43E-04 | PFK3 | ATP-dependent 6-phosphofructokinase 3 |
| AT3G25610 | -2.32 | 1.15E-04 | 0.05 | 9.65E-01 | 2.36 | 3.35E-05 | ALA10 | Phospholipid-transporting ATPase 10 |
| AT1G32940 | -2.33 | 8.04E-03 | -0.08 | 9.50E-01 | 2.24 | 1.10E-02 | SBT35 | Subtilase family protein |
| AT5G63790 | -2.33 | 1.22E-05 | -1.49 | 6.40E-03 | 0.83 | 9.71E-02 | At5g63790 | |
| AT2G25510 | -2.33 | 6.97E-05 | -0.55 | 1.12E-01 | 1.77 | 3.28E-04 | At2g25510 | Transmembrane protein |
| AT1G13830 | -2.34 | 3.71E-02 | 1.31 | 2.02E-01 | 3.61 | 2.12E-03 | At1g13830 | Carbohydrate-binding X8 domain superfamily protein |
| AT2G01540 | -2.34 | 1.01E-04 | -1.32 | 9.80E-03 | 1.00 | 1.74E-01 | CAR10 | Protein C2-DOMAIN ABA-RELATED 10 |
| AT1G79680 | -2.34 | 4.38E-02 | -0.06 | 1.00E+00 | 2.27 | 2.42E-02 | WAKL10 | Wall-associated receptor kinase-like 10 |
| AT1G65486 | -2.34 | 1.55E-05 | -0.37 | 5.91E-01 | 1.97 | 6.04E-04 | STMP4 | Transmembrane protein |
| AT5G35735 | -2.35 | 6.05E-04 | 0.19 | 6.22E-01 | 2.52 | 1.74E-04 | At5g35735 | Cytochrome b561 and DOMON domain-containing protein At5g35735 |
| AT2G41640 | -2.37 | 2.88E-05 | 0.28 | 8.70E-01 | 2.64 | 8.73E-14 | At2g41640 | Glycosyltransferase family 61 protein |
| AT2G28110 | -2.37 | 2.70E-02 | 0.12 | 9.70E-01 | 2.46 | 1.57E-02 | IRX7 | Probable glucuronoxylan glucuronosyltransferase IRX7 |
| AT5G56050 | -2.37 | 1.91E-03 | -0.40 | 8.87E-01 | 1.95 | 1.01E-02 | At5g56050 | Late embryogenesis abundant protein LEA-2 subgroup domain-containing protein |
| AT5G52640 | -2.37 | 5.80E-04 | -0.30 | 6.27E-01 | 2.07 | 1.83E-03 | HSP90-1 | Heat shock protein 90-1 |
| AT3G08720 | -2.37 | 8.28E-06 | -0.25 | 7.48E-01 | 2.10 | 1.21E-04 | ATPK2 | Serine/threonine-protein kinase AtPK2/AtPK19 |
| AT2G38860 | -2.37 | 1.70E-04 | -0.32 | 6.29E-01 | 2.03 | 1.79E-03 | YLS5 | Class I glutamine amidotransferase-like superfamily protein |
| AT2G29720 | -2.38 | 2.38E-05 | -0.77 | 3.17E-01 | 1.58 | 1.23E-02 | CTF2B | FAD/NAD |
| AT4G23170 | -2.38 | 9.21E-06 | -0.83 | 5.25E-08 | 1.53 | 5.85E-03 | CRK9 | Putative cysteine-rich receptor-like protein kinase 9 |
| AT5G38710 | -2.38 | 8.93E-03 | 0.23 | 8.24E-01 | 2.59 | 3.42E-03 | PDH2 | Proline dehydrogenase |
| AT5G08790 | -2.38 | 1.21E-08 | -0.66 | 7.08E-03 | 1.71 | 3.45E-05 | NAC081 | Protein ATAF2 |
| AT5G25930 | -2.38 | 3.33E-05 | -0.85 | 2.82E-04 | 1.52 | 1.01E-02 | T1N24.22 | Kinase family with leucine-rich repeat domain-containing protein |
| AT1G11300 | -2.38 | 1.04E-02 | 0.17 | 9.32E-01 | 2.54 | 1.02E-02 | EGM1 | Receptor-like serine/threonine-protein kinase |
| AT4G04695 | -2.39 | 5.84E-05 | -1.57 | 1.60E-03 | 0.81 | 2.13E-01 | CPK31 | Calcium-dependent protein kinase 31 |
| AT3G22400 | -2.39 | 6.50E-03 | -0.16 | 8.92E-01 | 2.20 | 9.71E-03 | LOX5 | Linoleate 9S-lipoxygenase 5 |
| AT3G09010 | -2.39 | 2.11E-05 | -0.53 | 4.70E-01 | 1.85 | 1.43E-04 | MZB10.4 | Putative receptor ser/thr protein kinase |
| AT3G01290 | -2.41 | 1.51E-04 | -0.51 | 2.19E-01 | 1.89 | 3.85E-04 | HIR3 | Hypersensitive-induced response protein 3 |
| AT2G46430 | -2.42 | 5.06E-08 | -0.99 | 1.87E-09 | 1.42 | 1.67E-03 | CNGC3 | Probable cyclic nucleotide-gated ion channel 3 |
| AT3G48090 | -2.42 | 4.11E-06 | -0.86 | 4.94E-08 | 1.54 | 5.92E-03 | EDS1 | Protein EDS1 |
| AT1G72900 | -2.42 | 4.14E-05 | -0.39 | 5.89E-01 | 2.03 | 5.48E-04 | At1g72900 | Similar to part of disease resistance protein |
| AT3G06890 | -2.44 | 2.12E-03 | | | 2.79 | 4.41E-04 | At3g06890 | hypothetical protein |
| AT2G24165 | -2.45 | 6.61E-04 | -1.18 | 5.85E-02 | 1.26 | 5.14E-02 | | |
| AT3G07520 | -2.45 | 9.32E-05 | -0.74 | 2.14E-02 | 1.68 | 8.63E-03 | At3g07520 | Glutamate receptor |
| AT5G55170 | -2.45 | 2.56E-03 | | | 3.11 | 9.11E-05 | | |
| AT5G50530 | -2.46 | 1.53E-02 | -0.07 | 9.87E-01 | 2.40 | 1.51E-02 | CBSCBSPB4 | CBS domain-containing protein CBSCBSPB4 |
| AT1G07000 | -2.47 | 1.38E-09 | -0.55 | 2.01E-01 | 1.91 | 8.78E-06 | EXO70B2 | Exocyst complex component EXO70B2 |
| AT1G15010 | -2.48 | 9.75E-05 | | | 2.62 | 1.50E-04 | T15D22.5 | T15D22.5 protein |
| AT2G21850 | -2.48 | 1.47E-09 | -0.79 | 1.09E-01 | 1.67 | 2.60E-05 | At2g21850 | Cysteine/Histidine-rich C1 domain family protein |
| AT5G44568 | -2.49 | 1.39E-05 | -0.95 | 1.73E-10 | 1.53 | 1.10E-02 | PROSCOOP4 | Serine rich endogenous peptide 4 |
| AT2G17120 | -2.51 | 2.12E-04 | -0.88 | 2.25E-09 | 1.62 | 2.33E-02 | LYM2 | LysM domain-containing GPI-anchored protein 2 |
| AT3G54150 | -2.51 | 2.07E-04 | 0.62 | 6.81E-01 | 3.13 | 1.92E-06 | EFD | Embryonic abundant protein-like |
| AT3G13850 | -2.52 | 2.77E-02 | -0.46 | 8.87E-01 | 2.03 | 3.59E-02 | LBD22 | LOB domain-containing protein 22 |
| AT1G70690 | -2.52 | 2.55E-03 | -0.13 | 9.02E-01 | 2.38 | 3.54E-03 | PDLP5 | Plasmodesmata-located protein 5 |
| AT1G72910 | -2.52 | 4.88E-06 | -0.42 | 5.18E-01 | 2.09 | 2.82E-06 | At1g72910 | Similar to part of disease resistance protein |
| AT3G52430 | -2.52 | 4.70E-07 | -0.32 | 3.13E-01 | 2.19 | 3.42E-05 | PAD4 | Lipase-like PAD4 |
| AT5G25820 | -2.52 | 1.05E-05 | -0.03 | 1.00E+00 | 2.49 | 7.46E-05 | At5g25820 | Exostosin family protein |
| AT4G23220 | -2.53 | 2.74E-06 | -0.34 | 3.64E-01 | 2.18 | 2.03E-05 | At4g23220 | Uncharacterized protein |
| AT1G66725 | -2.53 | 1.29E-03 | -1.45 | 5.78E-02 | 1.08 | 7.29E-02 | | |
| AT1G21750 | -2.53 | 4.44E-03 | -0.37 | 7.62E-02 | 2.14 | 1.59E-02 | PDIL1-1 | Protein disulfide-isomerase |
| AT5G58940 | -2.53 | 2.07E-04 | -0.67 | 2.16E-01 | 1.83 | 1.24E-02 | CRCK1 | Calmodulin-binding receptor-like cytoplasmic kinase 1 |
| AT3G13080 | -2.54 | 1.42E-05 | -0.24 | 5.00E-01 | 2.28 | 7.11E-05 | ABCC3 | Multidrug resistance-associated protein 3 |
| AT5G49680 | -2.54 | 9.89E-07 | -0.44 | 4.39E-01 | 2.10 | 2.03E-05 | KIP | Protein KINKY POLLEN |
| AT1G55210 | -2.55 | 1.53E-02 | -0.01 | 1.00E+00 | 2.52 | 1.52E-02 | DIR20 | Dirigent protein 20 |
| AT3G30120 | -2.56 | 3.74E-03 | -0.43 | 8.73E-01 | 2.10 | 1.18E-02 | | |
| AT1G52770 | -2.57 | 1.85E-05 | -2.63 | 2.92E-06 | -0.08 | 9.40E-01 | At1g52770 | At1g52770 |
| AT1G79450 | -2.57 | 4.13E-02 | -0.07 | 1.00E+00 | 2.50 | 8.91E-03 | ALIS5 | ALA-interacting subunit 5 |
| AT1G48370 | -2.57 | 1.49E-02 | -0.92 | 6.12E-01 | 1.64 | 1.26E-01 | YSL8 | YELLOW STRIPE like 8 |
| AT4G23230 | -2.57 | 2.06E-05 | -0.91 | 1.48E-02 | 1.66 | 1.51E-03 | CRK15 | Cysteine-rich receptor-like protein kinase 15 |
| AT2G15390 | -2.57 | 3.35E-05 | 0.25 | 6.48E-01 | 2.82 | 2.22E-06 | FUT4 | Probable fucosyltransferase 4 |
| AT3G61280 | -2.58 | 2.41E-06 | -0.35 | 3.78E-01 | 2.22 | 3.32E-06 | T20K12.180 | Uncharacterized protein T20K12.180 |
| AT1G34420 | -2.58 | 4.29E-05 | -0.36 | 5.41E-01 | 2.21 | 3.92E-04 | At1g34420 | Protein kinase domain-containing protein |
| AT4G17670 | -2.59 | 4.50E-03 | -0.02 | 1.00E+00 | 2.55 | 7.26E-04 | FLZ2 | FCS-Like Zinc finger 2 |
| AT1G01340 | -2.59 | 5.95E-04 | -0.35 | 6.20E-01 | 2.24 | 3.13E-03 | CNGC10 | Probable cyclic nucleotide-gated ion channel 10 |
| AT3G13790 | -2.59 | 1.65E-05 | -1.69 | 6.16E-17 | 0.88 | 2.28E-01 | ATBFRUCT1 | Glycosyl hydrolases family 32 protein |
| AT5G37540 | -2.59 | 2.93E-02 | 0.12 | 8.78E-01 | 2.70 | 1.91E-02 | At5g37540 | Peptidase A1 domain-containing protein |
| AT1G74360 | -2.59 | 4.15E-05 | -0.30 | 5.83E-01 | 2.29 | 3.60E-04 | At1g74360 | Probable LRR receptor-like serine/threonine-protein kinase At1g74360 |
| AT5G46350 | -2.59 | 1.38E-02 | -0.45 | 6.90E-01 | 2.15 | 2.59E-02 | WRKY8 | WRKY transcription factor 8 |
| AT3G01080 | -2.59 | 2.15E-07 | -0.74 | 1.38E-01 | 1.83 | 2.87E-04 | WRKY58 | WRKY DNA-binding protein 58 |
| AT1G59865 | -2.60 | 2.39E-04 | 0.14 | 9.80E-01 | 2.74 | 2.11E-02 | At1g59865 | Transmembrane protein |

| | | | | | | | | |
|---|---|---|---|---|---|---|---|---|
| AT3G52748 | -2.60 | 5.93E-03 | -0.83 | 6.34E-01 | 1.77 | 6.32E-02 | | |
| AT1G76970 | -2.61 | 6.05E-06 | -0.46 | 3.01E-01 | 2.14 | 3.73E-04 | At1g76970 | Target of Myb protein 1 |
| AT4G15417 | -2.62 | 2.17E-02 | | | 3.21 | 2.24E-02 | | |
| AT2G30550 | -2.62 | 1.74E-03 | -0.17 | 7.24E-01 | 2.44 | 3.87E-03 | DALL3 | Alpha/beta-Hydrolases superfamily protein |
| AT4G23160 | -2.62 | 9.94E-07 | -0.19 | 9.14E-01 | 2.42 | 1.01E-05 | At4g23160 | Uncharacterized protein |
| AT5G45380 | -2.63 | 1.08E-06 | -0.66 | 1.58E-01 | 1.96 | 7.05E-07 | DUR3 | Urea-proton symporter DUR3 |
| AT1G19020 | -2.63 | 5.78E-03 | -0.86 | 1.83E-01 | 1.76 | 4.43E-02 | At1g19020 | CDP-diacylglycerol-glycerol-3-phosphate 3-phosphatidyltransferase |
| AT1G27730 | -2.63 | 4.29E-05 | 0.56 | 5.03E-01 | 3.18 | 1.82E-08 | ZAT10 | Zinc finger protein ZAT10 |
| AT4G21903 | -2.63 | 1.28E-03 | 0.09 | 9.80E-01 | 2.71 | 3.77E-04 | At4g21903 | Protein DETOXIFICATION |
| AT5G63130 | -2.64 | 7.04E-06 | -0.74 | 6.15E-01 | 1.85 | 2.04E-04 | At5g63130 | hypothetical protein |
| AT5G64310 | -2.66 | 7.65E-09 | -0.08 | 9.74E-01 | 2.57 | 9.98E-09 | AGP1 | Classical arabinogalactan protein 1 |
| AT1G12420 | -2.66 | 1.00E-06 | -0.79 | 1.53E-01 | 1.84 | 1.58E-03 | ACR8 | ACT domain-containing protein ACR |
| AT5G54860 | -2.67 | 6.04E-06 | -0.92 | 6.10E-04 | 1.74 | 6.15E-03 | At5g54860 | Probable folate-biopterin transporter 4 |
| AT1G67970 | -2.67 | 6.26E-05 | -0.38 | 5.95E-01 | 2.28 | 8.16E-04 | HSFA8 | Heat stress transcription factor A-8 |
| AT1G58390 | -2.67 | 2.17E-04 | | | 2.93 | 3.28E-04 | At1g58390 | Probable disease resistance protein At1g58390 |
| AT4G24190 | -2.67 | 1.89E-03 | -0.50 | 3.54E-03 | 2.16 | 1.30E-02 | SHD | Chaperone protein htpG family protein |
| AT3G25020 | -2.67 | 1.59E-06 | -0.50 | 4.78E-03 | 2.15 | 1.08E-04 | RLP42 | Receptor-like protein 42 |
| AT1G68390 | -2.69 | 2.01E-03 | -0.95 | 5.64E-01 | 1.72 | 4.54E-02 | At1g68390 | Core-2/I-branching beta-1,6-N-acetylglucosaminyltransferase family protein |
| AT3G10930 | -2.70 | 4.30E-04 | | | 2.30 | 4.10E-03 | IDL7 | Uncharacterized protein At3g10930 |
| AT1G48210 | -2.70 | 1.74E-08 | -0.51 | 3.38E-01 | 2.18 | 7.36E-06 | At1g48210 | Protein kinase domain-containing protein |
| AT1G09715 | -2.70 | 1.54E-03 | -0.45 | 8.87E-01 | 2.21 | 1.17E-02 | | |
| AT5G01550 | -2.70 | 1.79E-02 | 1.22 | 4.70E-01 | 3.93 | 1.93E-04 | LECRK-VI3 | Lectin receptor kinase a4.1 |
| AT3G22160 | -2.71 | 1.02E-05 | -0.27 | 7.25E-01 | 2.43 | 1.21E-04 | VQ22 | VQ motif-containing protein 22 |
| AT1G76650 | -2.71 | 1.87E-02 | | | 3.84 | 1.12E-12 | CML38 | Calcium-binding protein CML38 |
| AT4G14400 | -2.72 | 5.37E-10 | -1.32 | 7.34E-19 | 1.38 | 1.16E-03 | ACD6 | Protein ACCELERATED CELL DEATH 6 |
| AT1G66465 | -2.72 | 8.10E-04 | -0.81 | 2.31E-01 | 1.88 | 3.02E-02 | At1g66465 | Transmembrane protein |
| AT3G12830 | -2.72 | 1.82E-04 | -0.75 | 2.78E-01 | 1.96 | 1.10E-02 | SAUR72 | Auxin-responsive protein SAUR72 |
| AT1G21120 | -2.72 | 3.44E-03 | | | 2.10 | 3.95E-02 | IGMT2 | O-methyltransferase family protein |
| AT3G21150 | -2.73 | 2.15E-04 | | | 2.10 | 2.92E-02 | BBX32 | B-box zinc finger protein 32 |
| AT5G41750 | -2.74 | 4.45E-07 | -0.08 | 9.32E-01 | 2.65 | 1.96E-08 | MUF8.3 | Disease resistance protein |
| AT3G05500 | -2.75 | 1.85E-05 | -0.45 | 3.35E-01 | 2.27 | 9.67E-04 | LDAP3 | Rubber elongation factor protein |
| AT3G11402 | -2.75 | 2.79E-07 | -1.15 | 8.54E-05 | 1.58 | 6.35E-03 | At3g11402 | Cysteine/Histidine-rich C1 domain family protein |
| AT4G37370 | -2.77 | 1.51E-04 | -0.67 | 1.58E-01 | 2.09 | 4.03E-03 | CYP81D8 | Cytochrome P450, family 81, subfamily D, polypeptide 8 |
| AT4G33050 | -2.77 | 1.24E-09 | -0.50 | 1.96E-02 | 2.26 | 8.17E-07 | EDA39 | Calmodulin-binding family protein |
| AT2G39210 | -2.77 | 9.24E-07 | -0.80 | 4.02E-04 | 1.97 | 4.85E-04 | PIC30 | At2g39210/T16B24.15 |
| AT2G26560 | -2.78 | 2.81E-05 | 0.47 | 5.40E-01 | 3.23 | 1.12E-12 | PLP2 | Patatin-like protein 2 |
| AT5G40230 | -2.78 | 5.15E-08 | -0.52 | 8.27E-01 | 2.34 | 6.55E-06 | At5g40230 | WAT1-related protein |
| AT3G23510 | -2.79 | 1.51E-05 | -1.72 | 2.30E-03 | 1.07 | 1.20E-01 | At3g23510 | Amine oxidase domain-containing protein |
| AT5G13490 | -2.80 | 2.58E-03 | -0.70 | 6.00E-01 | 2.08 | 3.02E-02 | AAC2 | ADP,ATP carrier protein 2, mitochondrial |
| AT3G26440 | -2.81 | 3.84E-03 | 0.28 | 8.39E-01 | 3.08 | 1.71E-04 | At3g26440 | Transmembrane protein, putative |
| AT5G07780 | -2.81 | 1.31E-07 | 0.51 | 7.71E-01 | 3.32 | 1.96E-08 | FH19 | Formin-like protein 19 |
| AT1G04600 | -2.81 | 1.54E-03 | -0.43 | 7.77E-01 | 2.36 | 3.09E-03 | At1g04600 | Uncharacterized protein |
| AT4G24000 | -2.81 | 2.93E-02 | | | 3.90 | 9.79E-03 | CSLG2 | Cellulose synthase like G2 |
| AT2G33580 | -2.82 | 4.76E-07 | -0.68 | 2.29E-05 | 2.13 | 1.71E-04 | LYK5 | Protein LYK5 |
| AT4G20110 | -2.82 | 1.96E-05 | -0.23 | 7.68E-01 | 2.57 | 3.35E-05 | VSR7 | VACUOLAR SORTING RECEPTOR 7 |
| AT4G01870 | -2.83 | 5.84E-05 | -0.26 | 6.90E-01 | 2.53 | 3.76E-04 | At4g01870 | TolB protein-like protein |
| AT2G30140 | -2.83 | 3.26E-06 | -0.61 | 9.52E-02 | 2.22 | 2.15E-04 | UGT87A2 | UDP-glycosyltransferase 87A2 |
| AT5G24206 | -2.83 | 2.34E-04 | | | 2.46 | 5.67E-04 | | |
| AT3G62600 | -2.84 | 7.05E-04 | -0.60 | 5.10E-03 | 2.22 | 9.83E-03 | ERDJ3B | DnaJ protein ERDJ3B |
| AT4G34380 | -2.84 | 4.00E-02 | | | 2.61 | 1.38E-02 | | |
| AT1G67920 | -2.86 | 1.94E-02 | -0.32 | 9.27E-01 | 2.45 | 3.30E-02 | At1g67920 | Uncharacterized protein |
| AT2G22500 | -2.86 | 4.34E-06 | -0.90 | 2.96E-03 | 1.94 | 1.08E-03 | PUMP5 | Mitochondrial uncoupling protein 5 |
| AT5G26170 | -2.87 | 2.02E-05 | -1.16 | 5.59E-02 | 1.68 | 7.01E-03 | At5g26170 | WRKY50 |
| AT1G29290 | -2.87 | 2.08E-02 | -1.24 | 9.56E-01 | 2.00 | 3.56E-03 | CEP14 | C-terminally encoded peptide 14 |
| AT5G18270 | -2.87 | 9.74E-03 | | | 4.18 | 3.71E-04 | ANAC087 | NAC domain containing protein 87 |
| AT3G62780 | -2.87 | 9.47E-06 | -1.28 | 6.86E-02 | 1.57 | 1.22E-02 | F26K9_210 | At3g62780 |
| AT2G40140 | -2.88 | 2.24E-07 | -0.31 | 7.30E-01 | 2.55 | 2.28E-07 | At2g40140 | Zinc finger CCCH domain-containing protein 29 |
| AT3G51350 | -2.89 | 3.48E-03 | | | 2.38 | 2.04E-03 | | |
| AT1G07620 | -2.89 | 9.64E-06 | -0.47 | 5.23E-01 | 2.42 | 1.16E-04 | ATOBGM | GTP-binding protein Obg/CgtA |
| AT4G29520 | -2.90 | 9.74E-03 | -0.47 | 5.48E-01 | 2.42 | 2.35E-02 | SES1 | Nucleophosmin |
| AT5G60950 | -2.91 | 5.17E-05 | -1.33 | 9.22E-04 | 1.57 | 1.66E-02 | COBL5 | COBRA-like protein 5 |
| AT2G17040 | -2.92 | 5.63E-05 | -0.80 | 2.61E-02 | 2.10 | 1.08E-03 | At2g17040 | At2g17040 |
| AT3G50480 | -2.93 | 3.50E-12 | -0.79 | 2.34E-08 | 2.13 | 4.42E-07 | HR4 | Homolog of RPW8 4 |
| AT1G76960 | -2.93 | 3.67E-04 | -0.68 | 2.64E-01 | 2.24 | 3.52E-04 | At1g76960 | F22K20.6 protein |
| AT4G05020 | -2.93 | 6.25E-05 | -0.21 | 8.35E-01 | 2.72 | 3.43E-04 | NDB2 | NADH:ubiquinone reductase |
| AT3G50280 | -2.94 | 1.63E-02 | | | 4.62 | 8.48E-04 | | |
| AT5G54610 | -2.95 | 7.97E-11 | -1.15 | 2.61E-08 | 1.79 | 8.64E-06 | BAD1 | Ankyrin repeat-containing protein BDA1 |
| AT5G55460 | -2.97 | 1.70E-04 | -0.45 | 8.22E-01 | 2.47 | 4.33E-04 | ATLTP45 | At5g55460 |
| AT2G22470 | -2.97 | 5.16E-03 | -0.36 | 8.60E-01 | 2.61 | 8.48E-03 | AGP2 | Classical arabinogalactan protein 2 |
| AT5G12880 | -2.98 | 6.95E-03 | | | 2.84 | 3.79E-02 | | |
| AT2G26440 | -2.98 | 1.14E-07 | -0.75 | 1.11E-05 | 2.21 | 5.40E-05 | PME12 | Probable pectinesterase/pectinesterase inhibitor 12 |
| AT5G01100 | -2.98 | 3.08E-08 | -0.41 | 5.34E-01 | 2.55 | 5.54E-06 | FRB1 | Protein FRIABLE 1 |
| AT1G61420 | -2.98 | 6.74E-05 | -0.38 | 8.74E-01 | 2.55 | 7.82E-04 | At1g61420 | S-locus lectin protein kinase family protein |
| AT1G72000 | -2.99 | 3.45E-03 | | | 2.32 | 2.96E-02 | | |
| AT5G61790 | -2.99 | 5.68E-04 | -0.56 | 9.96E-04 | 2.42 | 6.15E-03 | CNX1 | Calnexin homolog 1 |
| AT2G44290 | -2.99 | 4.29E-04 | -0.78 | 3.39E-06 | 2.20 | 1.16E-02 | LTPG13 | Non-specific lipid transfer protein GPI-anchored 13 |
| AT3G63380 | -2.99 | 3.25E-03 | -0.51 | 3.53E-01 | 2.48 | 1.61E-02 | ACA12 | Calcium-transporting ATPase 12, plasma membrane-type |
| AT5G57480 | -3.00 | 1.07E-03 | -1.19 | 3.89E-01 | 1.86 | 6.56E-02 | At5g57480 | AAA-ATPase At5g57480 |
| AT3G51440 | -3.01 | 5.08E-04 | 0.16 | 9.02E-01 | 3.17 | 3.77E-04 | SSL6 | Protein STRICTOSIDINE SYNTHASE-LIKE 6 |

| | | | | | | | | |
|---|---|---|---|---|---|---|---|---|
| AT2G46270 | -3.01 | 7.20E-07 | -0.83 | 2.70E-01 | 2.18 | 3.77E-04 | At2g46270 | BZIP domain-containing protein |
| AT5G59680 | -3.01 | 6.54E-09 | -1.19 | 5.28E-03 | 1.82 | 1.94E-06 | At5g59680 | Probable LRR receptor-like serine/threonine-protein kinase At5g59680 |
| AT1G05340 | -3.01 | 1.58E-02 | 0.04 | 9.93E-01 | 3.08 | 2.42E-02 | CYSTM1 | Protein CYSTEINE-RICH TRANSMEMBRANE MODULE 1 |
| AT1G80840 | -3.02 | 1.59E-06 | 0.97 | 3.70E-01 | 3.97 | 3.00E-20 | WRKY40 | Probable WRKY transcription factor 40 |
| AT1G63720 | -3.02 | 2.40E-06 | -0.60 | 2.66E-01 | 2.40 | 3.02E-04 | F24D7.9 | Uncharacterized protein F24D7.9 |
| AT3G12220 | -3.02 | 7.30E-06 | -1.40 | 2.50E-02 | 1.62 | 1.21E-04 | SCPL16 | Serine carboxypeptidase-like 16 |
| AT1G08450 | -3.04 | 6.39E-07 | -0.74 | 1.75E-05 | 2.29 | 1.48E-04 | At1g08450 | Calreticulin like protein |
| AT5G26920 | -3.04 | 1.37E-06 | -0.38 | 2.69E-01 | 2.65 | 2.28E-05 | CBP60G | Cam-binding protein 60-like G |
| AT5G60900 | -3.05 | 1.48E-09 | -1.41 | 2.78E-11 | 1.63 | 8.56E-04 | RLK1 | G-type lectin S-receptor-like serine/threonine-protein kinase RLK1 |
| AT3G56710 | -3.05 | 1.38E-08 | -0.72 | 1.67E-02 | 2.31 | 4.92E-06 | SIB1 | Sigma factor binding protein 1, chloroplastic |
| AT5G25250 | -3.07 | 9.39E-06 | -0.17 | 7.51E-01 | 2.88 | 3.14E-05 | At5g25250 | Flotillin-like |
| AT5G24205 | -3.07 | 1.14E-07 | -0.30 | 9.14E-01 | 2.73 | 9.12E-07 | | |
| AT1G74710 | -3.08 | 9.50E-06 | -0.29 | 3.71E-01 | 2.78 | 9.30E-05 | ICS1 | Isochorismate synthase 1, chloroplastic |
| AT1G56550 | -3.10 | 7.71E-07 | -0.99 | 1.27E-01 | 2.10 | 4.66E-04 | At1g56550 | At1g56550 |
| AT3G07195 | -3.10 | 1.41E-06 | -1.05 | 5.23E-02 | 2.03 | 1.48E-03 | At3g07195 | RPM1-interacting protein 4 |
| AT3G26210 | -3.10 | 3.21E-06 | -0.42 | 1.22E-01 | 2.66 | 3.42E-05 | CYP71B23 | Cytochrome P450 71B23 |
| AT5G43910 | -3.11 | 4.91E-06 | -0.86 | 5.41E-02 | 2.25 | 1.44E-03 | At5g43910 | PfkB-like carbohydrate kinase family protein |
| AT1G14260 | -3.11 | 6.44E-03 | -0.98 | 5.85E-01 | 2.17 | 2.61E-02 | At1g14260 | At1g14260 |
| AT4G14370 | -3.12 | 3.68E-06 | -1.04 | 7.15E-02 | 2.08 | 7.90E-04 | DL3225C | Disease resistance protein |
| AT4G13900 | -3.12 | 4.34E-06 | -0.85 | 5.46E-04 | 2.26 | 4.86E-04 | RLP49 | Receptor-like protein 49 |
| AT1G24140 | -3.13 | 4.80E-06 | -0.73 | 2.48E-01 | 2.40 | 1.49E-05 | 3MMP | Metalloendoproteinase 3-MMP |
| AT3G25510 | -3.13 | 3.18E-04 | -0.34 | 8.88E-01 | 2.78 | 4.33E-09 | At3g25510 | ADP-ribosyl cyclase/cyclic ADP-ribose hydrolase |
| AT5G27420 | -3.13 | 1.67E-04 | 0.39 | 8.24E-01 | 3.51 | 1.85E-05 | ATL31 | E3 ubiquitin-protein ligase ATL31 |
| AT1G65790 | -3.14 | 1.10E-07 | -1.11 | 2.90E-05 | 2.01 | 1.17E-04 | RK1 | Receptor kinase 1 |
| AT2G18680 | -3.15 | 1.46E-04 | | | 3.12 | 1.27E-05 | At2g18680 | Uncharacterized protein At2g18680 |
| AT3G04220 | -3.15 | 1.42E-02 | 0.04 | 9.93E-01 | 3.19 | 5.08E-03 | At3g04220 | Disease resistance protein family |
| AT4G15610 | -3.16 | 2.41E-02 | 0.18 | 9.60E-01 | 3.35 | 1.35E-02 | CASPL1D1 | CASP-like protein |
| AT5G28540 | -3.16 | 3.12E-04 | -0.55 | 3.47E-03 | 2.60 | 3.29E-03 | BIP1 | Heat shock 70 kDa protein BIP1 |
| AT4G23810 | -3.17 | 2.83E-08 | -1.23 | 6.17E-09 | 1.92 | 1.04E-03 | WRKY53 | Probable WRKY transcription factor 53 |
| AT1G51790 | -3.17 | 1.15E-06 | -1.41 | 3.04E-03 | 1.75 | 3.46E-05 | At1g51790 | Leucine-rich repeat protein kinase family protein |
| AT3G48650 | -3.18 | 2.74E-10 | -0.95 | 6.67E-03 | 2.22 | 3.63E-06 | At3g48650 | UPF0496 protein At3g48650 |
| AT3G09940 | -3.18 | 3.36E-03 | 0.06 | 9.91E-01 | 3.22 | 2.20E-03 | MDAR3 | Monodehydroascorbate reductase |
| AT3G48080 | -3.18 | 6.73E-07 | -1.27 | 8.90E-03 | 1.91 | 5.09E-04 | EDS1B | Protein EDS1B |
| AT3G13100 | -3.18 | 2.19E-06 | -0.12 | 9.42E-01 | 3.05 | 6.39E-06 | ABCC7 | ABC-type xenobiotic transporter |
| AT1G66880 | -3.18 | 8.60E-06 | -0.77 | 8.35E-08 | 2.40 | 8.81E-04 | At1g66880 | Protein kinase superfamily protein |
| AT1G72240 | -3.20 | 3.98E-04 | -1.33 | 4.46E-01 | 1.91 | 3.55E-02 | At1g72240 | Uncharacterized protein |
| AT5G21900 | -3.21 | 1.97E-06 | | | 0.78 | 2.66E-01 | RAD7C | RNI-like superfamily protein |
| AT5G52050 | -3.22 | 9.67E-10 | -0.72 | 6.20E-01 | 2.43 | 3.71E-07 | DTX50 | Protein DETOXIFICATION 50 |
| AT2G40750 | -3.22 | 8.64E-11 | -1.28 | 4.12E-07 | 1.94 | 4.59E-06 | WRKY54 | Probable WRKY transcription factor 54 |
| AT4G26120 | -3.22 | 9.22E-04 | -0.77 | 7.78E-01 | 2.53 | 1.96E-05 | At4g26120 | Ankyrin repeat family protein / BTB/POZ domain-containing protein |
| AT2G47800 | -3.22 | 1.03E-07 | -0.99 | 1.03E-06 | 2.22 | 3.00E-04 | ABCC4 | ABC transporter C family member 4 |
| AT5G60800 | -3.23 | 3.97E-05 | -1.16 | 1.28E-02 | 2.05 | 1.10E-02 | HIPP3 | Heavy metal transport/detoxification superfamily protein |
| AT5G62520 | -3.23 | 2.63E-07 | | | 4.22 | 6.17E-09 | SRO5 | Probable inactive poly [ADP-ribose] polymerase SRO5 |
| AT5G59670 | -3.24 | 3.10E-07 | -1.66 | 3.88E-05 | 1.56 | 9.77E-05 | At5g59670 | Receptor-like protein kinase At5g59670 |
| AT2G31880 | -3.24 | 4.82E-06 | -0.77 | 5.58E-05 | 2.46 | 4.21E-04 | SOBIR1 | Leucine-rich repeat receptor-like serine/threonine/tyrosine-protein kinase SOBIR1 |
| AT4G39030 | -3.26 | 8.01E-09 | -0.12 | 9.51E-01 | 3.13 | 1.96E-08 | DTX47 | Protein DETOXIFICATION 47, chloroplastic |
| AT5G25910 | -3.27 | 1.55E-05 | -0.91 | 1.92E-01 | 2.33 | 2.87E-03 | RLP52 | Receptor-like protein 52 |
| AT5G66620 | -3.27 | 2.16E-04 | -1.73 | 6.96E-03 | 1.49 | 6.55E-02 | DAR6 | DA1-related protein 6 |
| AT3G22231 | -3.27 | 2.36E-07 | -2.23 | 3.63E-15 | 1.04 | 7.48E-02 | PCC1 | Cysteine-rich and transmembrane domain-containing protein PCC1 |
| AT2G20142 | -3.28 | 7.10E-06 | -0.57 | 4.92E-01 | 2.71 | 6.02E-05 | At2g20142 | Toll-Interleukin-Resistance |
| AT5G67340 | -3.28 | 4.00E-06 | -1.03 | 1.41E-05 | 2.24 | 1.99E-03 | PUB2 | U-box domain-containing protein 2 |
| AT2G02810 | -3.28 | 1.17E-05 | -0.97 | 1.33E-02 | 2.29 | 4.66E-03 | UTR1 | UDP-galactose/UDP-glucose transporter 1 |
| AT1G76040 | -3.28 | 3.48E-04 | -0.49 | 3.69E-02 | 2.79 | 2.87E-03 | CPK29 | Calcium-dependent protein kinase 29 |
| AT1G77810 | -3.29 | 3.10E-03 | 0.02 | 1.00E+00 | 3.31 | 3.11E-03 | At1g77810 | Hexosyltransferase |
| AT1G72280 | -3.29 | 4.44E-05 | -0.92 | 7.27E-02 | 2.35 | 5.05E-03 | At1g72280 | |
| AT3G54960 | -3.29 | 1.68E-04 | -0.76 | 6.93E-03 | 2.53 | 4.97E-03 | PDIL1-3 | Protein disulfide isomerase-like 1-3 |
| AT5G08240 | -3.30 | 1.57E-05 | -0.07 | 9.94E-01 | 3.21 | 8.64E-06 | At5g08240 | Transmembrane protein |
| AT3G22235 | -3.30 | 5.06E-08 | -2.01 | 4.95E-13 | 1.27 | 1.32E-02 | CYSTM8 | Protein CYSTEINE-RICH TRANSMEMBRANE MODULE 8 |
| AT5G27060 | -3.30 | 2.00E-05 | 0.10 | 9.85E-01 | 3.38 | 1.34E-04 | RLP53 | Receptor-like protein 53 |
| AT5G35580 | -3.31 | 1.90E-03 | | | 1.63 | 1.75E-01 | | |
| AT5G43420 | -3.31 | 1.06E-03 | | | 2.21 | 3.54E-02 | ATL16 | RING-H2 finger protein ATL16 |
| AT3G08970 | -3.32 | 3.58E-04 | -0.96 | 2.36E-01 | 2.35 | 1.56E-02 | ERDJ3A | DnaJ protein ERDJ3A |
| AT3G13380 | -3.32 | 7.40E-05 | -0.55 | 3.11E-01 | 2.76 | 1.08E-03 | BRL3 | Receptor-like protein kinase BRI1-like 3 |
| AT3G47780 | -3.35 | 4.05E-05 | -0.75 | 4.49E-01 | 2.58 | 1.09E-02 | ABCA7 | ABC transporter A family member 7 |
| AT5G20230 | -3.36 | 4.06E-05 | -0.03 | 9.95E-01 | 3.32 | 2.90E-05 | BCB | Blue copper protein |
| AT5G62770 | -3.36 | 1.15E-04 | -0.05 | 9.99E-01 | 3.30 | 5.94E-05 | At5g62770 | Uncharacterized protein |
| AT5G59820 | -3.37 | 3.69E-06 | | | 4.11 | 3.10E-06 | ZAT12 | Zinc finger protein ZAT12 |
| AT5G52750 | -3.38 | 1.47E-07 | -0.85 | 1.24E-02 | 2.52 | 3.44E-05 | At5g52750 | Heavy metal transport/detoxification superfamily protein |
| AT5G03350 | -3.40 | 1.62E-06 | -2.08 | 1.26E-13 | 1.30 | 4.94E-02 | LLP | Lectin-like protein |
| AT5G50200 | -3.41 | 9.87E-08 | -0.72 | 1.08E-01 | 2.67 | 1.80E-05 | NRT3.1 | High-affinity nitrate transporter 3.1 |
| AT5G24530 | -3.42 | 3.50E-12 | -1.63 | 6.71E-09 | 1.78 | 3.45E-08 | DMR6 | Protein DOWNY MILDEW RESISTANCE 6 |
| AT3G61190 | -3.44 | 2.65E-06 | | | 4.50 | 6.77E-06 | At3g61190 | C2 domain-containing protein |
| AT1G33760 | -3.44 | 1.38E-04 | | | 5.01 | 2.94E-05 | | |
| AT5G64510 | -3.44 | 2.09E-05 | -0.71 | 2.82E-01 | 2.71 | 1.20E-03 | At5g64510 | Tunicamycin induced protein |
| AT3G47480 | -3.44 | 4.97E-04 | -0.89 | 9.46E-02 | 2.55 | 2.62E-03 | CML47 | Probable calcium-binding protein CML47 |
| AT5G52390 | -3.45 | 3.23E-03 | | | 2.56 | 2.20E-02 | | |
| AT3G56400 | -3.46 | 1.24E-09 | -0.99 | 1.49E-07 | 2.46 | 4.71E-06 | WRKY70 | Probable WRKY transcription factor 70 |
| AT1G21250 | -3.47 | 4.15E-07 | -1.30 | 1.32E-08 | 2.16 | 5.98E-04 | WAK1 | Wall-associated receptor kinase 1 |
| AT4G18253 | -3.48 | 1.35E-06 | -1.26 | 2.39E-07 | 2.22 | 2.98E-03 | At4g18253 | Receptor Serine/Threonine kinase-like protein |

| | | | | | | | | |
|---|---|---|---|---|---|---|---|---|
| AT2G04070 | -3.49 | 9.76E-04 | 0.05 | 9.93E-01 | 3.44 | 4.34E-04 | DTX4 | Protein DETOXIFICATION 4 |
| AT3G60415 | -3.49 | 4.61E-09 | -0.77 | 1.94E-02 | 2.70 | 2.82E-06 | At3g60415 | Phosphoglycerate mutase family protein |
| AT1G02920 | -3.49 | 1.37E-06 | -0.58 | 7.75E-02 | 2.89 | 1.64E-05 | GSTF7 | Glutathione S-transferase F7 |
| AT1G56120 | -3.49 | 9.74E-12 | -0.89 | 1.45E-03 | 2.59 | 7.23E-07 | At1g56120 | Leucine-rich repeat transmembrane protein kinase |
| AT2G44383 | -3.50 | 1.17E-05 | | | 2.27 | 2.05E-03 | | |
| AT5G45630 | -3.52 | 8.63E-08 | | | 2.28 | 3.08E-03 | | |
| AT3G17690 | -3.53 | 1.38E-04 | | | 2.60 | 7.48E-03 | | |
| AT2G32140 | -3.54 | 1.55E-08 | -0.37 | 9.02E-01 | 3.18 | 3.88E-10 | At2g32140 | Transmembrane receptor |
| AT4G21850 | -3.54 | 1.12E-04 | 0.19 | 9.59E-01 | 3.68 | 6.31E-04 | MSRB9 | Peptide methionine sulfoxide reductase B9-S-oxide reductase |
| AT3G28540 | -3.57 | 2.85E-06 | -0.06 | 9.85E-01 | 3.50 | 4.19E-06 | At3g28540 | AAA+ ATPase domain-containing protein |
| AT1G77510 | -3.60 | 3.16E-04 | -0.58 | 1.61E-02 | 3.01 | 2.46E-03 | PDIL1-2 | Protein disulfide isomerase-like 1-2 |
| AT5G05340 | -3.62 | 2.15E-04 | | | 2.70 | 2.46E-03 | PER52 | Peroxidase 52 |
| AT3G02705 | -3.65 | 1.68E-02 | | | 4.52 | 1.34E-04 | | |
| AT4G18250 | -3.66 | 2.67E-07 | -0.28 | 8.50E-01 | 3.35 | 2.70E-08 | At4g18250 | Receptor Serine/Threonine kinase-like protein |
| AT5G48657 | -3.66 | 1.38E-09 | -0.63 | 2.52E-01 | 3.01 | 3.51E-07 | At5g48657 | Defense protein-like protein |
| AT1G73810 | -3.68 | 2.80E-06 | 0.13 | 9.63E-01 | 3.80 | 7.25E-06 | At1g73810 | Core-2/I-branching beta-1,6-N-acetylglucosaminyltransferase family protein |
| AT2G44400 | -3.69 | 5.13E-08 | -1.78 | 9.94E-02 | 1.99 | 1.80E-03 | At2g44400 | Cysteine/Histidine-rich C1 domain family protein |
| AT2G44380 | -3.69 | 5.95E-08 | -0.93 | 6.73E-01 | 2.68 | 6.17E-09 | At2g44380 | hypothetical protein |
| AT3G51330 | -3.69 | 4.90E-07 | -0.90 | 2.01E-02 | 2.78 | 1.09E-04 | At3g51330 | Eukaryotic aspartyl protease family protein |
| AT1G51620 | -3.70 | 1.17E-02 | | | 2.90 | 1.11E-01 | | |
| AT1G35710 | -3.70 | 8.98E-09 | -1.35 | 6.56E-08 | 2.34 | 2.83E-05 | At1g35710 | Probable leucine-rich repeat receptor-like protein kinase At1g35710 |
| AT1G26420 | -3.72 | 2.15E-04 | 0.59 | 7.37E-01 | 4.27 | 1.84E-05 | FOX5 | Berberine bridge enzyme-like 7 |
| AT4G03450 | -3.72 | 1.56E-09 | -0.95 | 2.41E-02 | 2.77 | 4.72E-07 | At4g03450 | Ankyrin repeat family protein |
| AT4G21120 | -3.73 | 6.12E-06 | -1.06 | 5.03E-01 | 2.70 | 3.19E-03 | AAT1 | Amino acid transporter 1 |
| AT1G57560 | -3.73 | 4.85E-04 | -1.06 | 5.58E-01 | 2.75 | 9.92E-04 | MYB50 | At1g57560 |
| AT4G30640 | -3.73 | 1.59E-06 | | | 1.95 | 1.10E-02 | | |
| AT3G25882 | -3.76 | 8.06E-03 | | | 2.73 | 8.68E-02 | | |
| AT5G56960 | -3.76 | 1.76E-03 | | | 2.08 | 3.27E-02 | | |
| AT3G23250 | -3.77 | 6.85E-11 | | | 4.08 | 3.52E-06 | MYB15 | Myb domain protein 15 |
| AT5G67450 | -3.78 | 2.55E-05 | -0.91 | 3.81E-01 | 2.86 | 7.74E-04 | AZF1 | Zinc finger protein AZF1 |
| AT1G78410 | -3.79 | 3.10E-03 | -1.68 | 2.95E-01 | 2.06 | 1.96E-02 | VQ10 | VQ motif-containing protein 10 |
| AT5G38250 | -3.80 | 1.10E-04 | -1.14 | 1.35E-01 | 2.62 | 8.50E-03 | At5g38250 | Protein kinase domain-containing protein |
| AT5G44567 | -3.80 | 3.48E-04 | | | 3.18 | 4.10E-03 | | |
| AT4G25110 | -3.83 | 1.37E-13 | -0.60 | 2.62E-01 | 3.21 | 1.31E-11 | AMC2 | Metacaspase-2 |
| AT1G24145 | -3.83 | 2.17E-05 | -0.78 | 1.96E-01 | 3.05 | 2.90E-04 | At1g24145 | At1g24145 |
| AT5G46050 | -3.84 | 8.80E-06 | -0.71 | 5.44E-01 | 3.10 | 1.88E-05 | NPF52 | Peptide transporter 3 |
| AT1G03850 | -3.86 | 3.58E-06 | -0.77 | 4.43E-01 | 3.09 | 1.16E-05 | GRXS13 | Glutaredoxin family protein |
| AT1G02230 | -3.89 | 9.58E-04 | -2.93 | 2.15E-02 | 1.28 | 1.80E-01 | NAC004 | NAC domain-containing protein 4 |
| AT2G43140 | -3.90 | 3.74E-06 | | | 1.97 | 2.95E-02 | BHLH129 | Basic helix-loop-helix DNA-binding superfamily protein |
| AT4G23210 | -3.90 | 8.77E-08 | -0.78 | 4.09E-01 | 3.12 | 2.60E-05 | CRK13 | Cysteine-rich RLK |
| AT5G52810 | -3.92 | 1.76E-11 | -1.64 | 9.51E-14 | 2.28 | 5.26E-05 | At5g52810 | AT5G52810 protein |
| AT5G42020 | -3.94 | 2.37E-05 | -0.94 | 1.97E-07 | 2.99 | 1.33E-03 | BIP2 | Heat shock protein 70 |
| AT2G27660 | -3.96 | 3.68E-06 | -0.77 | 5.39E-01 | 3.20 | 4.83E-07 | At2g27660 | Cysteine/Histidine-rich C1 domain family protein |
| AT2G43000 | -3.97 | 3.44E-07 | -1.12 | 8.28E-02 | 2.82 | 3.01E-04 | NAC042 | NAC domain containing protein 42 |
| AT1G58225 | -3.97 | 1.81E-05 | -0.87 | 8.20E-02 | 3.09 | 7.90E-04 | At1g58225 | Transmembrane protein |
| AT1G51800 | -4.03 | 1.30E-11 | -0.86 | 5.91E-02 | 3.14 | 3.14E-08 | IOS1 | LRR receptor-like serine/threonine-protein kinase IOS1 |
| AT4G23700 | -4.03 | 8.46E-04 | | | 4.58 | 3.52E-04 | CHX17 | Cation/Hantiporter 17 |
| AT2G34940 | -4.04 | 5.86E-06 | -2.48 | 4.40E-05 | 1.52 | 1.77E-02 | VSR5 | Vacuolar-sorting receptor 5 |
| AT5G17760 | -4.04 | 2.35E-06 | -1.00 | 6.63E-02 | 3.03 | 5.98E-04 | At5g17760 | AAA-ATPase At5g17760 |
| AT3G57460 | -4.06 | 9.41E-13 | -2.21 | 8.24E-08 | 1.84 | 1.88E-04 | T8H10.60 | Uncharacterized protein T8H10.60 |
| AT5G52740 | -4.08 | 4.37E-10 | | | 2.83 | 5.09E-06 | At5g52740 | Copper transport protein family |
| AT4G14365 | -4.08 | 2.54E-08 | -1.08 | 3.89E-08 | 2.98 | 3.20E-05 | XBAT34 | Putative E3 ubiquitin-protein ligase XBAT34 |
| AT3G50930 | -4.09 | 2.68E-09 | -0.30 | 5.69E-01 | 3.78 | 5.33E-08 | At3g50930 | AAA+ ATPase domain-containing protein |
| AT1G08050 | -4.11 | 3.92E-07 | | | 2.77 | 2.76E-03 | At1g08050 | Zinc finger family protein |
| AT1G43910 | -4.11 | 8.75E-04 | -1.82 | 5.46E-02 | 2.24 | 2.15E-04 | At1g43910 | AAA-ATPase At1g43910 |
| AT1G32960 | -4.11 | 4.14E-05 | | | 4.97 | 3.52E-06 | SBT3.3 | Subtilisin-like protease SBT3.3 |
| AT1G03660 | -4.12 | 1.59E-06 | | | 4.09 | 2.79E-06 | At1g03660 | Ankyrin-repeat containing protein |
| AT3G23120 | -4.12 | 3.58E-13 | -1.78 | 5.49E-23 | 2.33 | 2.03E-05 | RLP38 | Receptor-like protein 38 |
| AT3G28580 | -4.14 | 9.41E-05 | | | 4.01 | 2.83E-04 | At3g28580 | AAA+ ATPase domain-containing protein |
| AT2G18690 | -4.14 | 7.94E-06 | -0.31 | 2.54E-01 | 3.82 | 4.09E-05 | At2g18690 | At2g18690/MSF3.7 |
| AT5G10760 | -4.14 | 4.27E-08 | -1.34 | 8.84E-11 | 2.79 | 7.52E-05 | AED1 | Aspartyl protease AED1 |
| AT3G11010 | -4.15 | 5.57E-09 | -1.48 | 3.51E-20 | 2.66 | 1.09E-04 | RLP34 | Receptor like protein 34 |
| AT5G18470 | -4.15 | 4.32E-04 | | | 3.41 | 3.80E-03 | At5g18470 | Bulb-type lectin domain-containing protein |
| AT4G39830 | -4.15 | 5.86E-06 | -1.48 | 3.42E-05 | 2.65 | 4.22E-03 | AO | L-ascorbate oxidase |
| AT5G22520 | -4.17 | 1.64E-03 | -0.95 | 6.23E-01 | 3.23 | 4.16E-03 | MQJ16.6 | At5g22520 |
| AT2G29110 | -4.18 | 1.60E-06 | -0.05 | 9.89E-01 | 4.11 | 4.54E-06 | GLR28 | Glutamate receptor |
| AT5G13320 | -4.18 | 1.81E-07 | -0.45 | 2.35E-01 | 3.72 | 3.52E-06 | PBS3 | Auxin-responsive GH3 family protein |
| AT3G00850 | -4.19 | 2.57E-08 | -1.50 | 1.37E-01 | 2.67 | 1.84E-05 | | |
| AT5G05300 | -4.19 | 1.53E-05 | | | 2.34 | 1.92E-02 | | |
| AT5G53110 | -4.22 | 1.24E-05 | -0.43 | 8.55E-01 | 3.78 | 7.88E-05 | At5g53110 | RING-type E3 ubiquitin transferase |
| AT4G11655 | -4.24 | 1.68E-02 | | | 3.57 | 2.20E-04 | | |
| AT2G14560 | -4.24 | 3.39E-15 | -1.84 | 1.18E-33 | 2.39 | 2.19E-06 | LURP1 | LURP-one-like protein |
| AT3G60540 | -4.26 | 4.42E-05 | | | 2.86 | 1.79E-03 | T8B10_200 | Protein transport protein Sec61 subunit beta |
| AT2G20150 | -4.27 | 3.20E-05 | | | 2.76 | 1.74E-02 | At2g20150 | Uncharacterized protein |
| AT2G04495 | -4.27 | 2.06E-06 | -1.14 | 4.75E-01 | 3.12 | 3.14E-04 | At2g04495 | At2g04495 |
| AT4G25000 | -4.30 | 1.92E-05 | -1.41 | 3.01E-02 | 2.85 | 2.49E-03 | AMY1 | Alpha-amylase 1 |
| AT4G23280 | -4.31 | 1.24E-04 | | | 2.70 | 9.45E-02 | | |
| AT3G29000 | -4.31 | 2.03E-05 | | | 3.43 | 3.68E-04 | CML45 | Probable calcium-binding protein CML45 |
| AT2G42350 | -4.33 | 4.15E-05 | | | 4.21 | 2.42E-04 | | |

| | | | | | | | | |
|---|---|---|---|---|---|---|---|---|
| AT1G73805 | -4.35 | 1.76E-11 | -1.15 | 3.47E-03 | 3.20 | 3.66E-09 | SARD1 | Protein SAR DEFICIENT 1 |
| AT1G13340 | -4.36 | 5.68E-06 | -0.94 | 4.88E-01 | 3.42 | 3.18E-04 | ISTL6 | Regulator of Vps4 activity in the MVB pathway protein |
| AT1G51850 | -4.36 | 1.78E-07 | -1.73 | 7.30E-03 | 2.60 | 2.39E-03 | SIF2 | Leucine-rich repeat protein kinase family protein |
| AT1G30900 | -4.45 | 1.73E-06 | -1.46 | 3.62E-06 | 2.99 | 8.45E-04 | VSR6 | Vacuolar-sorting receptor 6 |
| AT2G32680 | -4.45 | 1.48E-09 | -1.71 | 4.36E-14 | 2.73 | 3.87E-05 | RLP23 | Receptor like protein 23 |
| AT2G46400 | -4.50 | 1.39E-08 | -0.87 | 1.85E-01 | 3.61 | 3.45E-08 | WRKY46 | Probable WRKY transcription factor 46 |
| AT5G42830 | -4.51 | 8.52E-10 | -2.17 | 9.60E-07 | 2.34 | 1.92E-04 | MBD2.2 | HXXXD-type acyl-transferase family protein |
| AT5G44570 | -4.53 | 5.99E-05 | | | 4.10 | 3.59E-04 | | |
| AT5G22545 | -4.53 | 2.96E-04 | | | 8.75 | 3.49E-06 | | |
| AT4G37010 | -4.53 | 6.86E-04 | | | 3.57 | 6.76E-03 | CEN2 | Centrin 2 |
| AT1G51860 | -4.54 | 1.27E-04 | | | 4.20 | 5.76E-03 | At1g51860 | Leucine-rich repeat protein kinase family protein |
| AT4G23140 | -4.57 | 4.95E-09 | -1.59 | 4.71E-38 | 2.96 | 1.23E-04 | CRK6 | Cysteine-rich receptor-like protein kinase 6 |
| AT4G38560 | -4.58 | 2.36E-09 | -0.96 | 9.69E-02 | 3.59 | 8.45E-07 | At4g38560 | |
| AT4G02380 | -4.58 | 6.84E-05 | -1.17 | 5.77E-01 | 3.31 | 4.46E-04 | SAG21 | Senescence-associated gene 21 |
| AT1G09932 | -4.59 | 4.14E-07 | -0.97 | 1.13E-01 | 3.61 | 3.73E-06 | At1g09932 | Phosphoglycerate mutase family protein |
| AT1G13470 | -4.67 | 6.88E-13 | -1.53 | 5.43E-08 | 3.14 | 1.34E-08 | At1g13470 | Uncharacterized protein At1g13470 |
| AT2G29350 | -4.67 | 2.37E-05 | -0.50 | 4.62E-01 | 4.16 | 1.16E-04 | SAG13 | Senescence-associated protein 13 |
| AT3G02840 | -4.68 | 8.86E-07 | | | 5.09 | 2.27E-06 | At3g02840 | U-box domain-containing protein |
| AT3G25010 | -4.70 | 5.63E-07 | -1.48 | 4.53E-11 | 3.21 | 3.51E-04 | RLP41 | Receptor like protein 41 |
| AT1G02930 | -4.70 | 6.76E-08 | -1.58 | 8.33E-08 | 3.12 | 5.20E-05 | GSTF6 | Glutathione S-transferase F6 |
| AT1G65690 | -4.70 | 4.63E-07 | -1.35 | 1.96E-02 | 3.32 | 2.46E-04 | NHL6 | NDR1/HIN1-like protein 6 |
| AT2G04515 | -4.73 | 1.12E-04 | | | 3.40 | 9.61E-03 | | |
| AT2G24850 | -4.73 | 2.23E-06 | -0.98 | 2.18E-01 | 3.74 | 4.22E-06 | TAT3 | Probable aminotransferase TAT3 |
| AT2G47130 | -4.73 | 8.53E-11 | -1.95 | 1.25E-07 | 2.77 | 7.24E-05 | SDR3a | Short-chain dehydrogenase reductase 3a |
| AT4G04510 | -4.77 | 1.66E-05 | | | 4.84 | 2.87E-05 | CRK38 | Cysteine-rich RLK 38 |
| AT3G23109 | -4.78 | 1.59E-06 | | | 2.08 | 2.71E-02 | | |
| AT3G28890 | -4.80 | 2.71E-15 | -1.31 | 4.83E-03 | 3.46 | 1.52E-09 | RLP43 | Receptor-like protein 43 |
| AT1G10340 | -4.80 | 1.05E-09 | -1.16 | 2.28E-02 | 3.64 | 4.19E-07 | At1g10340 | Ankyrin repeat family protein |
| AT3G09270 | -4.82 | 2.23E-03 | | | 3.23 | 8.04E-03 | GSTU8 | Glutathione S-transferase U8 |
| AT5G48400 | -4.82 | 1.07E-04 | | | 6.15 | 5.98E-06 | GLR1.2 | Glutamate receptor 1.2 |
| AT4G11890 | -4.82 | 2.12E-10 | -1.86 | 5.52E-06 | 2.96 | 1.46E-05 | ARCK1 | Protein kinase superfamily protein |
| AT3G24900 | -4.84 | 1.00E-06 | -1.46 | 5.63E-05 | 3.37 | 3.32E-04 | RLP39 | Receptor like protein 39 |
| AT3G60470 | -4.85 | 4.34E-06 | | | 2.69 | 2.15E-04 | T8B10_130 | Transmembrane protein, putative |
| AT4G25070 | -4.86 | 3.57E-08 | -1.28 | 2.47E-01 | 3.57 | 4.49E-05 | F13M23.200 | Uncharacterized protein AT4g25060 |
| AT4G04490 | -4.86 | 6.28E-12 | -1.56 | 8.28E-05 | 3.29 | 3.10E-06 | CRK36 | Cysteine-rich receptor-like protein kinase 36 |
| AT4G12490 | -4.91 | 1.42E-02 | | | 8.37 | 8.15E-04 | | |
| AT5G02490 | -4.92 | 1.57E-06 | -0.73 | 1.64E-01 | 4.19 | 2.03E-05 | HSP70-2 | Heat shock 70 kDa protein 2 |
| AT4G11170 | -4.93 | 9.67E-10 | | | 3.84 | 3.87E-04 | At4g11170 | Putative disease resistance protein At4g11170 |
| AT4G17660 | -4.95 | 2.07E-04 | | | 2.87 | 3.27E-02 | | |
| AT5G01900 | -4.96 | 8.72E-04 | | | 4.14 | 9.33E-04 | | |
| AT5G25260 | -4.98 | 1.14E-06 | -1.09 | 4.93E-01 | 3.90 | 2.51E-04 | FLOT2 | Flotillin-like protein 2 |
| AT2G25440 | -4.99 | 6.04E-06 | | | 4.71 | 1.27E-09 | RLP20 | Receptor-like protein 20 |
| AT5G09290 | -5.07 | 2.01E-05 | -2.63 | 6.09E-02 | 2.55 | 5.10E-03 | SAL4 | Probable SAL4 phosphatase |
| AT5G11210 | -5.12 | 5.46E-04 | | | 4.03 | 1.58E-03 | | |
| AT5G64810 | -5.18 | 8.17E-05 | | | 3.21 | 3.90E-03 | | |
| AT4G23310 | -5.18 | 1.60E-06 | -1.81 | 4.14E-03 | 3.35 | 1.72E-05 | CRK23 | Cysteine-rich RLK |
| AT4G23215 | -5.21 | 5.85E-08 | | | 3.71 | 1.94E-04 | | |
| AT1G01560 | -5.24 | 2.74E-08 | -2.45 | 8.02E-06 | 2.81 | 9.02E-05 | MPK11 | Mitogen-activated protein kinase |
| AT1G04980 | -5.27 | 3.54E-05 | -2.03 | 4.78E-03 | 3.15 | 3.52E-04 | At1g04980 | PDIL2-2 |
| AT1G02450 | -5.27 | 1.40E-04 | | | 2.56 | 2.61E-02 | | |
| AT3G61198 | -5.30 | 2.37E-07 | | | 2.66 | 1.71E-02 | | |
| AT2G33080 | -5.30 | 1.62E-06 | -1.34 | 3.96E-01 | 3.99 | 9.68E-06 | RLP28 | Receptor like protein 28 |
| AT3G57240 | -5.31 | 1.55E-11 | -3.29 | 2.55E-07 | 2.02 | 2.31E-04 | At3g57240 | glucan endo-1,3-beta-D-glucosidase |
| AT1G09080 | -5.32 | 7.32E-06 | -1.92 | 1.14E-03 | 3.36 | 1.93E-03 | At1g09080 | |
| AT5G22570 | -5.32 | 9.99E-09 | -2.27 | 3.15E-07 | 3.01 | 4.08E-04 | WRKY38 | Probable WRKY transcription factor 38 |
| AT5G47850 | -5.34 | 4.55E-09 | | | 4.59 | 1.37E-07 | CCR4 | Serine/threonine-protein kinase-like protein CCR4 |
| AT4G14390 | -5.37 | 1.87E-06 | -3.24 | 1.51E-05 | 2.11 | 1.18E-02 | DL3235W | Ankyrin repeat family protein |
| AT1G57650 | -5.37 | 1.25E-05 | | | 4.20 | 3.25E-05 | At1g57650 | ATP binding protein |
| AT5G52760 | -5.39 | 7.52E-08 | -1.46 | 3.09E-04 | 3.92 | 4.39E-05 | At5g52760 | Copper transport protein family |
| AT1G57640 | -5.41 | 2.57E-06 | | | 3.04 | 1.26E-03 | | |
| AT2G19190 | -5.42 | 1.38E-09 | -0.95 | 4.54E-01 | 4.47 | 2.13E-07 | SIRK | Senescence-induced receptor-like serine/threonine-protein kinase |
| AT5G26690 | -5.43 | 3.60E-05 | -1.39 | 5.16E-01 | 4.16 | 3.61E-06 | HIPP02 | Heavy metal-associated isoprenylated plant protein 2 |
| AT1G20310 | -5.43 | 2.55E-09 | | | 3.19 | 1.36E-05 | | |
| AT3G11000 | -5.46 | 1.82E-05 | | | 2.16 | 3.34E-02 | At3g11000 | DCD (Development and Cell Death) domain protein |
| AT2G18660 | -5.50 | 1.93E-14 | -3.29 | 3.88E-18 | 2.17 | 1.10E-03 | EGC2 | EG45-like domain containing protein 2 |
| AT3G15536 | -5.51 | 1.32E-07 | -1.62 | 5.35E-05 | 3.90 | 5.25E-05 | | |
| AT1G58420 | -5.52 | 8.94E-03 | -2.95 | 4.96E-02 | 2.67 | 5.33E-02 | | |
| AT1G36622 | -5.53 | 9.39E-06 | | | 3.92 | 3.03E-03 | | |
| AT1G35230 | -5.54 | 6.84E-05 | -1.15 | 1.55E-03 | 4.39 | 1.53E-03 | At1g35230 | |
| AT3G22910 | -5.57 | 4.76E-06 | -1.60 | 6.39E-04 | 3.97 | 8.20E-04 | ACA13 | Putative calcium-transporting ATPase 13, plasma membrane-type |
| AT1G51890 | -5.58 | 3.51E-08 | -0.74 | 5.78E-01 | 4.82 | 3.10E-06 | At1g51890 | Leucine-rich repeat protein kinase family protein |
| AT5G11920 | -5.58 | 2.34E-06 | | | 3.95 | 3.57E-05 | CWINV6 | 6-&1-fructan exohydrolase |
| AT1G66960 | -5.61 | 6.57E-12 | -2.89 | 2.77E-11 | 2.67 | 4.89E-05 | LUP5 | Terpene cyclase/mutase family member |
| AT1G01680 | -5.61 | 5.84E-11 | -2.14 | 9.96E-04 | 3.48 | 1.46E-05 | PUB54 | U-box domain-containing protein 54 |
| AT1G07900 | -5.68 | 4.16E-05 | | | 3.58 | 2.71E-03 | | |
| AT3G57260 | -5.70 | 3.10E-04 | -1.47 | 8.73E-03 | 4.22 | 2.09E-03 | PR2 | glucan endo-1,3-beta-D-glucosidase |
| AT1G21240 | -5.76 | 1.91E-03 | -1.60 | 1.87E-01 | 4.15 | 1.10E-05 | WAK3 | Wall-associated receptor kinase-like |
| AT4G00700 | -5.77 | 1.24E-09 | -2.05 | 5.75E-11 | 3.69 | 3.52E-05 | MCTP9 | Multiple C2 domain and transmembrane region protein 9 |
| AT2G00570 | -5.78 | 1.93E-05 | | | 2.92 | 9.58E-03 | | |

| | | | | | | | | |
|---|---|---|---|---|---|---|---|---|
| AT3G61390 | -5.78 | 1.71E-06 | | | 8.44 | 1.99E-06 | | |
| AT2G04450 | -5.82 | 4.07E-17 | -2.47 | 5.16E-13 | 3.36 | 5.33E-08 | NUDT6 | Nudix hydrolase 6 |
| AT4G23150 | -5.83 | 2.67E-07 | -1.98 | 8.62E-04 | 3.86 | 7.25E-06 | CRK7 | Cysteine-rich receptor-like protein kinase 7 |
| AT5G45000 | -5.84 | 1.48E-09 | | | 2.40 | 2.47E-03 | At5g45000 | Disease resistance protein family |
| AT3G48640 | -5.84 | 4.76E-07 | -3.45 | 2.30E-03 | 2.59 | 4.68E-03 | T8P19.150 | Transmembrane protein |
| AT1G68620 | -5.85 | 4.36E-04 | | | 4.13 | 1.00E-03 | | |
| AT1G57630 | -5.88 | 2.85E-05 | | | 4.09 | 2.10E-04 | At1g57630 | Disease resistance protein RPP1-WsB, putative domain family protein |
| AT4G14630 | -5.89 | 6.84E-05 | | | 4.15 | 3.17E-02 | | |
| AT2G45760 | -5.99 | 2.12E-04 | | | 8.81 | 2.28E-05 | | |
| AT3G44350 | -6.00 | 9.41E-05 | | | 6.30 | 4.46E-04 | NAC061 | NAC domain containing protein 61 |
| AT4G01360 | -6.09 | 1.31E-12 | | | 4.62 | 3.84E-10 | A_IG002N01. | A_IG002N01.26 protein |
| AT3G13950 | -6.10 | 2.42E-04 | | | 4.75 | 1.48E-04 | At3g13950 | At3g13950 |
| AT2G25297 | -6.12 | 2.03E-06 | | | 3.73 | 1.08E-03 | At2g25297 | Transmembrane protein |
| AT5G22530 | -6.13 | 3.94E-05 | | | 3.25 | 1.46E-02 | | |
| AT1G15520 | -6.14 | 5.57E-09 | -0.85 | 1.81E-01 | 5.27 | 1.66E-07 | ABCG40 | Pleiotropic drug resistance 12 |
| AT1G33950 | -6.15 | 4.07E-17 | | | 3.65 | 6.90E-06 | IAN7 | Immune-associated nucleotide-binding protein 7 |
| AT4G04500 | -6.18 | 9.47E-08 | -2.21 | 1.26E-05 | 3.99 | 1.25E-04 | CRK37 | Cysteine-rich RLK |
| AT5G22380 | -6.19 | 7.56E-09 | | | 3.85 | 5.43E-05 | NAC090 | NAC domain-containing protein 90 |
| AT1G65610 | -6.22 | 2.97E-04 | | | 6.45 | 5.33E-05 | KOR2 | Endoglucanase 7 |
| AT4G39670 | -6.25 | 4.87E-06 | | | 6.69 | 2.45E-07 | At4g39670 | ACD11 homolog protein |
| AT1G26380 | -6.25 | 6.49E-05 | | | 4.11 | 2.04E-03 | | |
| AT5G24110 | -6.28 | 7.96E-05 | -2.98 | 1.73E-02 | 3.37 | 3.38E-03 | WRKY30 | Probable WRKY transcription factor 30 |
| AT5G38900 | -6.32 | 4.95E-06 | | | 4.25 | 7.52E-05 | PDI | Thioredoxin superfamily protein |
| AT3G46090 | -6.35 | 1.05E-09 | | | 5.21 | 4.39E-05 | ZAT7 | Zinc finger protein ZAT7 |
| AT1G75040 | -6.38 | 8.82E-05 | -2.25 | 6.13E-10 | 4.12 | 5.77E-03 | At1g75040 | Pathogenesis-related protein 5 |
| AT1G47890 | -6.40 | 5.08E-08 | -2.04 | 1.56E-01 | 4.48 | 6.07E-06 | RLP7 | Receptor-like protein 7 |
| AT4G21840 | -6.47 | 2.86E-05 | | | 4.82 | 5.03E-04 | MSRB8 | Peptide methionine sulfoxide reductase B8-S-oxide reductase |
| AT1G14880 | -6.52 | 2.19E-11 | -3.04 | 1.00E-28 | 3.48 | 1.58E-05 | PCR1 | Protein PLANT CADMIUM RESISTANCE 1 |
| AT5G24200 | -6.59 | 2.07E-12 | -2.81 | 4.39E-03 | 3.82 | 4.06E-07 | At5g24200 | Alpha/beta-Hydrolases superfamily protein |
| AT2G40740 | -6.62 | 6.36E-06 | | | 3.30 | 2.28E-03 | WRKY55 | WRKY DNA-binding protein 55 |
| AT3G22600 | -6.65 | 8.81E-07 | -2.30 | 3.17E-02 | 4.46 | 1.94E-04 | LTPG5 | Non-specific lipid transfer protein GPI-anchored 5 |
| AT5G40010 | -6.68 | 1.38E-09 | | | 3.70 | 2.17E-04 | At5g40010 | AAA+ ATPase domain-containing protein |
| AT1G61800 | -6.76 | 1.53E-08 | | | 6.07 | 2.64E-08 | GPT2 | Glucose-6-phosphate/phosphate translocator 2 |
| AT1G44130 | -6.77 | 1.92E-04 | | | 5.11 | 5.03E-04 | At1g44130 | Peptidase A1 domain-containing protein |
| AT5G57010 | -6.86 | 9.32E-05 | | | 4.10 | 1.87E-02 | | |
| AT3G13433 | -6.87 | 9.09E-04 | | | 10.33 | 3.65E-05 | | |
| AT1G33960 | -6.93 | 8.14E-18 | -3.13 | 5.04E-21 | 3.77 | 2.14E-07 | AIG1 | P-loop containing nucleoside triphosphate hydrolases superfamily protein |
| AT3G60966 | -6.96 | 1.09E-06 | | | 3.87 | 9.65E-03 | At3g60966 | RING-type E3 ubiquitin transferase |
| AT2G29460 | -7.04 | 5.64E-07 | | | 4.14 | 9.65E-04 | GSTU4 | Glutathione S-transferase U4 |
| AT4G04540 | -7.06 | 1.40E-08 | | | 4.45 | 1.34E-04 | CRK39 | Cysteine-rich RLK 39 |
| AT1G65483 | -7.38 | 8.85E-07 | | | 4.53 | 2.89E-04 | At1g65483 | KH type-2 domain-containing protein |
| AT4G35180 | -7.50 | 2.55E-09 | | | 3.87 | 5.51E-05 | LHT7 | LYS/HIS transporter 7 |
| AT2G43570 | -7.53 | 9.68E-07 | -2.52 | 3.89E-08 | 5.04 | 2.13E-04 | CHI | Endochitinase CHI |
| AT1G56060 | -7.58 | 1.39E-06 | | | 4.25 | 1.25E-04 | ATHCYSTM3 | Cysteine-rich/transmembrane domain protein B |
| AT1G32350 | -7.60 | 1.38E-05 | | | 8.02 | 6.79E-06 | AOX1D | Ubiquinol oxidase |
| AT4G10500 | -7.64 | 1.48E-09 | -3.31 | 7.15E-31 | 4.36 | 7.52E-05 | DLO1 | Protein DMR6-LIKE OXYGENASE 1 |
| AT5G42380 | -7.79 | 8.28E-11 | | | 6.27 | 1.71E-08 | CML37 | Calcium-binding protein CML37 |
| AT3G26830 | -7.87 | 2.54E-08 | | | 5.64 | 3.51E-07 | At3g26830 | PAD3 |
| AT2G45220 | -7.99 | 1.60E-06 | | | 5.82 | 3.48E-05 | PME17 | Pectinesterase inhibitor 17] |
| AT5G51465 | -8.19 | 9.21E-06 | | | 4.85 | 5.26E-04 | PROSCOOP1 | Serine rich endogenous peptide 18 |
| AT2G34500 | -8.22 | 1.05E-09 | | | 6.64 | 1.48E-07 | CYP710A1 | Cytochrome P450 710A1 |
| AT5G66890 | -8.32 | 7.20E-08 | | | 8.10 | 6.11E-07 | NRG1C | N REQUIREMENT GENE 1.3 |
| AT1G75945 | -8.95 | 2.92E-03 | | | 4.37 | 1.08E-01 | | hypothetical protein |
| AT5G44990 | -9.00 | 9.16E-09 | | | 2.78 | 3.90E-03 | At5g44990 | Glutathione S-transferase family protein |
| AT2G35980 | -9.03 | 2.96E-07 | | | 8.29 | 7.61E-06 | NHL10 | NDR1/HIN1-like protein 10 |
| AT1G19250 | -9.12 | 1.59E-13 | | | 5.55 | 1.14E-08 | FMO1 | Probable flavin-containing monooxygenase 1 |
| AT1G05880 | -9.12 | 3.50E-12 | | | 8.90 | 2.91E-10 | ARI12 | RING/U-box superfamily protein |
| AT3G46080 | -9.33 | 1.02E-08 | | | 9.11 | 4.42E-07 | ZAT8 | Zinc finger protein ZAT8 |
| AT2G30770 | -9.41 | 6.22E-07 | | | 6.21 | 8.93E-05 | CYP71A13 | Indoleacetaldoxime dehydratase |
| AT2G14610 | -9.45 | 1.48E-09 | -4.56 | 1.20E-47 | 4.86 | 8.59E-05 | At2g14610 | Pathogenesis-related protein 1 |
| AT3G11340 | -9.49 | 5.60E-28 | -5.49 | 1.60E-13 | 3.88 | 1.52E-12 | UGT76B1 | UDP-glycosyltransferase 76B1 |
| AT1G26390 | -9.66 | 5.37E-10 | | | 6.11 | 1.09E-06 | FOX2 | Berberine bridge enzyme-like 4 |
| AT3G45330 | -9.70 | 5.26E-06 | | | 4.62 | 7.74E-03 | LECRK-I.1 | L-type lectin receptor kinase I.1 |
| AT2G39530 | -9.80 | 1.14E-06 | | | 4.66 | 1.84E-02 | CASPL4D1 | CASP-LIKE PROTEIN 4D1 |
| AT3G28510 | -9.90 | 6.84E-09 | | | 5.54 | 5.54E-06 | At3g28510 | P-loop containing nucleoside triphosphate hydrolases superfamily protein |
| AT3G21520 | -10.13 | 8.77E-08 | | | 5.67 | 9.83E-05 | DUF679 | DOMAIN MEMBRANE PROTEIN 1 |
| AT2G13810 | -10.14 | 7.17E-27 | -5.85 | 7.38E-13 | 4.21 | 1.73E-10 | ALD1 | Aminotransferase ALD1, chloroplastic |
| AT4G13890 | -10.27 | 1.09E-06 | | | 5.26 | 2.98E-04 | EDA36 | Serine hydroxymethyltransferase |
| AT3G53150 | -10.40 | 4.38E-07 | | | 5.47 | 4.55E-04 | UGT73D1 | UDP-glycosyltransferase 73D1 |
| AT3G48850 | -11.28 | 8.52E-10 | | | 4.67 | 3.53E-05 | MPT2 | Mitochondrial phosphate carrier protein 2, mitochondrial |
| AT2G14620 | -11.47 | 4.43E-11 | | | 3.76 | 4.06E-04 | XTH10 | Xyloglucan endotransglucosylase/hydrolase |
| AT1G71390 | -11.50 | 1.48E-09 | | | 3.63 | 2.85E-03 | RLP11 | Receptor like protein 11 |
| AT3G13610 | -12.07 | 6.62E-15 | | | 4.95 | 2.13E-07 | F6'H1 | Feruloyl CoA ortho-hydroxylase 1 |
| AT2G26400 | -13.06 | 3.00E-18 | | | 5.86 | 2.02E-08 | ARD3 | Acireductone dioxygenase |

Indian Farmers Fertilizer Cooperative (IFFCO)'s liquid nano urea formulation (NUF) was applied as a foliar spray to one-month-old *Arabidopsis thaliana* plants, and the leaf transcriptome was compared to that of equimolar bulk urea treatment. Mock treatment refers to a spray with distilled water. Comparisons between NUF and urea, NUF and mock, and urea and mock treatments are shown with $\log_2$(fold-change) in expression levels and Benjamini-Hochberg *false discovery rate* (*FDR*)-adjusted *P*-values. $\text{Log}_2\text{FC} > 0.5$ or $< -0.5$ are highlighted in shades of red and blue, respectively. *FDR* $< 0.05$ are indicated in red font.

**Table S2. Oligonucleotide sequences used for real time q-PCR**

| AGI code | Primer name | Sequence (5'--> 3') |
|---|---|---|
| AT5G59320 | LTP3_RT_Fw | AAAAGGTGGGTTGGTGCCAC |
| | LTP3_RT_Rv | CTTGTTGGCGGTCTGGTGTG |
| AT3G47340 | ASN1_RT_Fw | CGTGGCTTGTTCGACTGCAA |
| | ASN1_RT_Rv | ATGCCACGTTCTTGCCATCG |
| AT4G27440 | PORB_RT_Fw | GGTGAGTGATCCAAGCTTGACG |
| | PORB_RT_Rv | GCCCACGAGCTTCTCACTGA |
| AT1G02205 | CER1_RT_Fw | GCGCTTATTCACACCGGACC |
| | CER1_RT_Rv | GGTGGGAATGGTAGCGGGAA |
| AT5G50200 | NRT3.1_RT_Fw | GCGTTCAGGCTATCAGTGGC |
| | NRT3.1_RT_Rv | AGACGACAAGAGCCACGACG |
| AT3G52590 | UBQ1 _RT_Fw | TCGTAAGTACAATCAGGATAAGA |
| | UBQ1 _RT_Rv | CACTGAAACAAGAAAAACAAACC |

**Table S3.** ***Arabidopsis thaliana*** **genes of major functional categories down-regulated under IFFCO nano urea versus equimolar bulk urea**

| AGI code | Gene Symbol | Short Description | NUF vs Urea | | NUF vs Mock | | Urea vs Mock | |
|---|---|---|---|---|---|---|---|---|
| | | | $log_2FC$ | *FDR* | $log_2FC$ | *FDR* | $log_2FC$ | *FDR* |
| **Cell death** | | | | | | | | |
| AT3G13610 | F6'H1 | Feruloyl CoA ortho-hydroxylase 1 | -12.07 | **6.62E-15** | | | 4.95 | **2.13E-07** |
| AT5G11650 | MAGL13 | Alpha/beta-Hydrolases superfamily protein | -1.29 | **2.73E-03** | -0.46 | 4.37E-01 | 0.81 | 8.17E-02 |
| **Abscission** | | | | | | | | |
| AT1G74710 | ICS1 | Isochorismate synthase 1, chloroplastic | -3.08 | **9.50E-06** | -0.29 | 3.71E-01 | 2.78 | **9.30E-05** |
| AT4G28490 | RLK5 | Receptor-like protein kinase 5 | -2.17 | **1.45E-05** | -0.80 | **2.53E-06** | 1.36 | **9.78E-03** |
| AT1G51660 | MKK4 | Mitogen-activated protein kinase kinase 4 | -2.16 | **2.15E-04** | -0.69 | 5.58E-02 | 1.47 | **1.05E-02** |
| AT4G35600 | CST | Protein kinase superfamily protein | -1.61 | **3.62E-04** | -0.40 | 1.04E-01 | 1.19 | **1.42E-02** |
| AT5G14930 | SAG101 | Senescence-associated gene 101 | -1.35 | **2.70E-03** | -0.12 | 7.72E-01 | 1.21 | **7.40E-03** |
| AT3G21220 | MKK5 | Mitogen-activated protein kinase kinase 5 | -1.29 | **3.90E-03** | -0.14 | 7.56E-01 | 1.14 | **9.16E-03** |
| **Senescence** | | | | | | | | |
| AT2G13810 | ALD1 | Aminotransferase ALD1, chloroplastic | -10.14 | **7.17E-27** | -5.85 | **7.38E-13** | 4.21 | **1.73E-10** |
| AT3G11340 | UGT76B1 | UDP-glycosyltransferase 76B1 | -9.49 | **5.60E-28** | -5.49 | **1.60E-13** | 3.88 | **1.52E-12** |
| AT2G35980 | NHL10 | NDR1/HIN1-like protein 10 | -9.03 | **2.96E-07** | | | 8.29 | **7.61E-06** |
| AT4G10500 | DLO1 | Protein DMR6-LIKE OXYGENASE 1 | -7.64 | **1.48E-09** | -3.31 | **7.15E-31** | 4.36 | **7.52E-05** |
| AT4G21840 | MSRB8 | Peptide methionine sulfoxide reductase B8-S-oxide reductase | -6.47 | **2.86E-05** | | | 4.82 | **5.03E-04** |
| AT5G24110 | WRKY30 | Probable WRKY transcription factor 30 | -6.28 | **7.96E-05** | -2.98 | **1.73E-02** | 3.37 | **3.38E-03** |
| AT1G21240 | WAK3 | Wall-associated receptor kinase-like | -5.76 | **1.91E-03** | -1.60 | 1.87E-01 | 4.15 | **1.10E-05** |
| AT1G01680 | PUB54 | U-box domain-containing protein 54 | -5.61 | **5.84E-11** | -2.14 | **9.96E-04** | 3.48 | **1.46E-05** |
| AT3G22910 | ACA13 | Putative calcium-transporting ATPase 13, plasma membrane-type | -5.57 | **4.76E-06** | -1.60 | **6.39E-04** | 3.97 | **8.20E-04** |
| AT3G25010 | RLP41 | Receptor like protein 41 | -4.70 | **5.63E-07** | -1.48 | **4.53E-11** | 3.21 | **3.51E-04** |
| AT4G02380 | SAG21 | Senescence-associated gene 21 | -4.58 | **6.84E-05** | -1.17 | 5.77E-01 | 3.31 | **4.46E-04** |
| AT2G18690 | At2g18690 | At2g18690/MSF3.7 | -4.14 | **7.94E-06** | -0.31 | 2.54E-01 | 3.82 | **4.09E-05** |
| AT1G08050 | At1g08050 | Zinc finger family protein | -4.11 | **3.92E-07** | | | 2.77 | **2.76E-03** |
| AT4G14365 | XBAT34 | Putative E3 ubiquitin-protein ligase XBAT34 | -4.08 | **2.54E-08** | -1.08 | **3.89E-08** | 2.98 | **3.20E-05** |
| AT2G43000 | NAC042 | NAC domain containing protein 42 | -3.97 | **3.44E-07** | -1.12 | 8.28E-02 | 2.82 | **3.01E-04** |
| AT1G02230 | NAC004 | NAC domain-containing protein 4 | -3.89 | **9.58E-04** | -2.93 | **2.15E-02** | 1.28 | 1.80E-01 |
| AT2G32140 | At2g32140 | Transmembrane receptor | -3.54 | **1.55E-08** | -0.37 | 9.02E-01 | 3.18 | **3.88E-10** |
| AT3G60415 | At3g60415 | Phosphoglycerate mutase family protein | -3.49 | **4.61E-09** | -0.77 | **1.94E-02** | 2.70 | **2.82E-06** |
| AT3G56400 | WRKY70 | Probable WRKY transcription factor 70 | -3.46 | **1.24E-09** | -0.99 | **1.49E-07** | 2.46 | **4.71E-06** |
| AT5G24530 | DMR6 | Protein DOWNY MILDEW RESISTANCE 6 | -3.42 | **3.50E-12** | -1.63 | **6.71E-09** | 1.78 | **3.45E-08** |
| AT2G31880 | SOBIR1 | Leucine-rich repeat receptor-like serine/threonine/tyrosine-protein kinase SOBIR1 | -3.24 | **4.82E-06** | -0.77 | **5.58E-05** | 2.46 | **4.21E-04** |
| AT2G40750 | WRKY54 | Probable WRKY transcription factor 54 | -3.22 | **8.64E-11** | -1.28 | **4.12E-07** | 1.94 | **4.59E-06** |
| AT1G66880 | At1g66880 | Protein kinase superfamily protein | -3.18 | **8.60E-06** | -0.77 | **8.35E-08** | 2.40 | **8.81E-04** |
| AT3G48080 | EDS1B | Protein EDS1B | -3.18 | **6.73E-07** | -1.27 | **8.90E-03** | 1.91 | **5.09E-04** |
| AT4G23810 | WRKY53 | Probable WRKY transcription factor 53 | -3.17 | **2.83E-08** | -1.23 | **6.17E-09** | 1.92 | **1.04E-03** |
| AT4G15610 | CASPL1D1 | CASP-like protein | -3.16 | **2.41E-02** | 0.18 | 9.60E-01 | 3.35 | **1.35E-02** |
| AT4G14370 | DL3225C | Disease resistance protein | -3.12 | **3.68E-06** | -1.04 | 7.15E-02 | 2.08 | **7.90E-04** |
| AT3G26210 | CYP71B23 | Cytochrome P450 71B23 | -3.10 | **3.21E-06** | -0.42 | 1.22E-01 | 2.66 | **3.42E-05** |
| AT1G05340 | CYSTM1 | Protein CYSTEINE-RICH TRANSMEMBRANE MODULE 1 | -3.01 | **1.58E-02** | 0.04 | 9.93E-01 | 3.08 | **2.42E-02** |
| AT2G22470 | AGP2 | Classical arabinogalactan protein 2 | -2.97 | **5.16E-03** | -0.36 | 8.60E-01 | 2.61 | **8.48E-03** |
| AT4G01870 | At4g01870 | TolB protein-like protein | -2.83 | **5.84E-05** | -0.26 | 6.90E-01 | 2.53 | **3.76E-04** |
| AT5G63130 | At5g63130 | hypothetical protein | -2.64 | **7.04E-06** | -0.74 | 6.15E-01 | 1.85 | **2.04E-04** |
| AT2G30550 | DALL3 | Alpha/beta-Hydrolases superfamily protein | -2.62 | **1.74E-03** | -0.17 | 7.24E-01 | 2.44 | **3.87E-03** |
| AT1G34420 | At1g34420 | Protein kinase domain-containing protein | -2.58 | **4.29E-05** | -0.36 | 5.41E-01 | 2.21 | **3.92E-04** |
| AT3G52430 | PAD4 | Lipase-like PAD4 | -2.52 | **4.70E-07** | -0.32 | 3.13E-01 | 2.19 | **3.42E-05** |
| AT3G48090 | EDS1 | Protein EDS1 | -2.42 | **4.11E-06** | -0.86 | **4.94E-08** | 1.54 | **5.92E-03** |
| AT5G25930 | T1N24.22 | Kinase family with leucine-rich repeat domain-containing protein | -2.38 | **3.33E-05** | -0.85 | **2.82E-04** | 1.52 | **1.01E-02** |
| AT5G37600 | GLN1-1 | Glutamine synthetase cytosolic isozyme 1-1 | -2.31 | **1.95E-05** | -0.01 | 1.00E+00 | 2.28 | **1.96E-05** |
| AT1G76980 | At1g76980 | Patatin-like phospholipase domain protein | -2.24 | **9.84E-03** | -0.38 | 7.41E-01 | 1.87 | **1.96E-02** |
| AT5G39050 | PMAT1 | Phenolic glucoside malonyltransferase 1 | -2.20 | **1.49E-04** | -0.11 | 8.59E-01 | 2.07 | **2.32E-04** |
| AT1G34180 | NAC016 | NAC domain containing protein 16 | -1.91 | **5.75E-03** | 0.44 | 8.25E-01 | 2.34 | **1.34E-04** |
| AT2G13790 | SERK4 | Somatic embryogenesis receptor kinase 4 | -1.86 | **8.61E-07** | -0.61 | **3.28E-05** | 1.23 | **1.64E-03** |
| AT3G26840 | PES2 | Phytyl ester synthase 2, chloroplastic | -1.85 | **2.30E-02** | 0.49 | 5.89E-01 | 2.32 | **1.27E-03** |
| AT1G18210 | CML27 | Probable calcium-binding protein CML27 | -1.84 | **2.26E-04** | -0.14 | 8.60E-01 | 1.68 | **1.16E-03** |
| AT5G56870 | BGAL4 | Beta-galactosidase 4 | -1.70 | **2.15E-04** | 0.00 | 1.00E+00 | 1.68 | **1.72E-04** |
| AT1G02220 | NAC003 | NAC domain-containing protein 3 | -1.69 | **8.55E-03** | -0.08 | 9.72E-01 | 1.58 | **9.71E-03** |
| AT3G10500 | NAC053 | NAC domain containing protein 53 | -1.55 | **1.63E-03** | -0.14 | 8.60E-01 | 1.39 | **4.61E-03** |
| AT1G13990 | At1g13990 | Plant/protein | -1.47 | **2.26E-02** | 0.09 | 8.94E-01 | 1.55 | **6.01E-03** |
| AT3G15760 | At3g15760 | Transmembrane protein | -1.47 | **1.26E-03** | -0.62 | 3.28E-01 | 0.83 | 9.16E-02 |
| AT3G27010 | TCP20 | Transcription factor TCP20 | -1.35 | **6.95E-03** | -0.81 | 3.65E-01 | 0.53 | 5.16E-01 |
| AT2G17760 | APF1 | Aspartyl protease family protein 1 | -1.20 | **1.76E-02** | -0.18 | 6.42E-01 | 1.01 | **4.17E-02** |
| AT1G19270 | DA1 | Protein DA1 | -1.06 | **4.77E-03** | 0.05 | 9.58E-01 | 1.09 | **3.10E-03** |
| AT3G60800 | PAT14 | Probable protein S-acyltransferase 14 | -0.95 | **4.17E-02** | -0.09 | 8.90E-01 | 0.85 | 5.94E-02 |
| AT1G20780 | PUB44 | U-box domain-containing protein 44 | -0.94 | **1.17E-02** | -0.39 | 2.37E-01 | 0.53 | 1.60E-01 |
| AT3G19190 | ATG2 | Autophagy-related protein 2 | -0.80 | **1.53E-02** | -0.02 | 9.76E-01 | 0.76 | **1.35E-02** |
| AT1G54115 | CCX4 | Cation/calcium exchanger 4 | -0.76 | **1.94E-02** | 0.09 | 9.00E-01 | 0.84 | **4.04E-03** |
| **Cellular catabolic process** | | | | | | | | |
| AT4G15610 | CASPL1D1 | CASP-like protein | -3.16 | **2.41E-02** | 0.18 | 9.60E-01 | 3.35 | **1.35E-02** |
| AT1G05340 | CYSTM1 | Protein CYSTEINE-RICH TRANSMEMBRANE MODULE 1 | -3.01 | **1.58E-02** | 0.04 | 9.93E-01 | 3.08 | **2.42E-02** |
| AT3G63380 | ACA12 | Calcium-transporting ATPase 12, plasma membrane-type | -2.99 | **3.25E-03** | -0.51 | 3.53E-01 | 2.48 | **1.61E-02** |
| AT2G22470 | AGP2 | Classical arabinogalactan protein 2 | -2.97 | **5.16E-03** | -0.36 | 8.60E-01 | 2.61 | **8.48E-03** |
| AT4G01870 | At4g01870 | TolB protein-like protein | -2.83 | **5.84E-05** | -0.26 | 6.90E-01 | 2.53 | **3.76E-04** |
| AT5G35735 | At5g35735 | Cytochrome b561 and DOMON domain-containing protein At5g35735 | -2.35 | **6.05E-04** | 0.19 | 6.22E-01 | 2.52 | **1.74E-04** |
| AT1G13830 | At1g13830 | Carbohydrate-binding X8 domain superfamily protein | -2.34 | **3.71E-02** | 1.31 | 2.02E-01 | 3.61 | **2.12E-03** |
| AT3G49780 | PSK3 | Phytosulfokines 3 | -1.78 | **1.47E-02** | -0.16 | 8.03E-01 | 1.61 | **1.85E-02** |
| AT5G56870 | BGAL4 | Beta-galactosidase 4 | -1.70 | **2.15E-04** | 0.00 | 1.00E+00 | 1.68 | **1.72E-04** |
| AT3G26220 | At3g26220 | At3g26230 | -1.69 | **4.19E-05** | 0.03 | 9.89E-01 | 1.71 | **9.56E-06** |
| AT1G02220 | NAC003 | NAC domain-containing protein 3 | -1.69 | **8.55E-03** | -0.08 | 9.72E-01 | 1.58 | **9.71E-03** |
| AT1G27350 | At1g27330 | At1g27330 | -1.44 | **1.37E-02** | -0.18 | 7.58E-01 | 1.26 | **2.57E-02** |
| AT3G51980 | At3g51980 | Nucleotide exchange factor Fes1 domain-containing protein | -1.35 | **4.50E-02** | -0.10 | 9.28E-01 | 1.24 | 5.84E-02 |
| AT5G64550 | At5g64550 | Uncharacterized protein At5g64550 | -1.33 | **4.52E-03** | -0.18 | 8.52E-01 | 1.13 | **1.03E-02** |
| AT1G62305 | At1g62305 | At1g62305 | -1.32 | **1.52E-02** | -0.79 | 9.46E-02 | 0.52 | 4.21E-01 |
| AT4G08230 | At4g08230 | Glycine-rich protein | -1.28 | **3.00E-02** | -0.17 | 6.84E-01 | 1.10 | 6.45E-02 |
| AT2G25520 | At2g25520 | Probable sugar phosphate/phosphate translocator At2g25520 | -1.19 | **1.14E-02** | -0.26 | 1.78E-01 | 0.92 | 5.72E-02 |

| | | | | | | | | |
|---|---|---|---|---|---|---|---|---|
| AT1G54320 | ALIS3 | ALA-interacting subunit 3 | -1.08 | **1.90E-02** | -0.05 | 9.46E-01 | 1.02 | **2.34E-02** |
| AT4G34480 | At4g34480 | glucan endo-1,3-beta-D-glucosidase | -0.97 | **2.71E-02** | -0.21 | 4.06E-01 | 0.75 | 8.09E-02 |
| AT2G45500 | At2g45500 | AAA-type ATPase family protein | -0.95 | **1.38E-02** | -0.01 | 1.00E+00 | 0.93 | **9.86E-03** |
| AT3G23530 | At3g23530 | Amine oxidase domain-containing protein | -0.93 | **1.54E-03** | -0.87 | **1.81E-09** | 0.05 | 9.24E-01 |
| AT1G56460 | At1g56460 | HIT zinc finger and PAPA-1-like domain-containing protein | -0.83 | **3.17E-02** | 0.01 | 9.97E-01 | 0.82 | **2.21E-02** |
| AT5G61530 | At5g61530 | Small G protein family protein / RhoGAP family protein | -0.81 | **1.90E-02** | 0.13 | 7.73E-01 | 0.93 | **7.46E-03** |
| **Signal transduction** | | | | | | | | |
| AT1G71390 | RLP11 | Receptor like protein 11 | -11.50 | **1.48E-09** | | | 3.63 | **2.85E-03** |
| AT3G11340 | UGT76B1 | UDP-glycosyltransferase 76B1 | -9.49 | **5.60E-28** | -5.49 | **1.60E-13** | 3.88 | **1.52E-12** |
| AT3G22600 | LTPG5 | Non-specific lipid transfer protein GPI-anchored 5 | -6.65 | **8.81E-07** | -2.30 | **3.17E-02** | 4.46 | **1.94E-04** |
| AT4G21840 | MSRB8 | Peptide methionine sulfoxide reductase B8-S-oxide reductase | -6.47 | **2.86E-05** | | | 4.82 | **5.03E-04** |
| AT1G47890 | RLP7 | Receptor-like protein 7 | -6.40 | **5.08E-08** | -2.04 | 1.56E-01 | 4.48 | **6.07E-06** |
| AT1G15520 | ABCG40 | Pleiotropic drug resistance 12 | -6.14 | **5.57E-09** | -0.85 | 1.81E-01 | 5.27 | **1.66E-07** |
| AT3G13950 | At3g13950 | At3g13950 | -6.10 | **2.42E-04** | | | 4.75 | **1.48E-04** |
| AT1G57630 | At1g57630 | Disease resistance protein RPP1-WsB, putative domain family protein | -5.88 | **2.85E-05** | | | 4.09 | **2.10E-04** |
| AT3G48640 | T8P19.150 | Transmembrane protein | -5.84 | **4.76E-07** | -3.45 | **2.30E-03** | 2.59 | **4.68E-03** |
| AT5G45000 | At5g45000 | Disease resistance protein family | -5.84 | **1.48E-09** | | | 2.40 | **2.47E-03** |
| AT2G04450 | NUDT6 | Nudix hydrolase 6 | -5.82 | **4.07E-17** | -2.47 | **5.16E-13** | 3.36 | **5.33E-08** |
| AT4G00700 | MCTP9 | Multiple C2 domain and transmembrane region protein 9 | -5.77 | **1.24E-09** | -2.05 | **5.75E-11** | 3.69 | **3.52E-05** |
| AT1G21240 | WAK3 | Wall-associated receptor kinase-like | -5.76 | **1.91E-03** | -1.60 | 1.87E-01 | 4.15 | **1.10E-05** |
| AT1G01680 | PUB54 | U-box domain-containing protein 54 | -5.61 | **5.84E-11** | -2.14 | **9.96E-04** | 3.48 | **1.46E-05** |
| AT3G22910 | ACA13 | Putative calcium-transporting ATPase 13, plasma membrane-type | -5.57 | **4.76E-06** | -1.60 | **6.39E-04** | 3.97 | **8.20E-04** |
| AT1G35230 | At1g35230 | | -5.54 | **6.84E-05** | -1.15 | **1.55E-03** | 4.39 | **1.53E-03** |
| AT5G26690 | HIPP02 | Heavy metal-associated isoprenylated plant protein 2 | -5.43 | **3.60E-05** | -1.39 | 5.16E-01 | 4.16 | **3.61E-06** |
| AT4G14390 | DL3235W | Ankyrin repeat family protein | -5.37 | **1.87E-06** | -3.24 | **1.51E-05** | 2.11 | **1.18E-02** |
| AT5G22570 | WRKY38 | Probable WRKY transcription factor 38 | -5.32 | **9.99E-09** | -2.27 | **3.15E-07** | 3.01 | **4.08E-04** |
| AT2G33080 | RLP28 | Receptor like protein 28 | -5.30 | **1.62E-06** | -1.34 | 3.96E-01 | 3.99 | **9.68E-06** |
| AT1G01560 | MPK11 | Mitogen-activated protein kinase | -5.24 | **2.74E-08** | -2.45 | **8.02E-06** | 2.81 | **9.02E-05** |
| AT2G25440 | RLP20 | Receptor-like protein 20 | -4.99 | **6.04E-06** | | | 4.71 | **1.27E-09** |
| AT4G11170 | At4g11170 | Putative disease resistance protein At4g11170 | -4.93 | **9.67E-10** | | | 3.84 | **3.87E-04** |
| AT3G24900 | RLP39 | Receptor like protein 39 | -4.84 | **1.00E-06** | -1.46 | **5.63E-05** | 3.37 | **3.32E-04** |
| AT4G11890 | ARCK1 | Protein kinase superfamily protein | -4.82 | **2.12E-10** | -1.86 | **5.52E-06** | 2.96 | **1.46E-05** |
| AT5G48400 | GLR1.2 | Glutamate receptor 1.2 | -4.82 | **1.07E-04** | | | 6.15 | **5.98E-06** |
| AT1G10340 | At1g10340 | Ankyrin repeat family protein | -4.80 | **1.05E-09** | -1.16 | **2.28E-02** | 3.64 | **4.19E-07** |
| AT3G28890 | RLP43 | Receptor-like protein 43 | -4.80 | **2.71E-15** | -1.31 | **4.83E-03** | 3.46 | **1.52E-09** |
| AT3G25010 | RLP41 | Receptor like protein 41 | -4.70 | **5.63E-07** | -1.48 | **4.53E-11** | 3.21 | **3.51E-04** |
| AT1G13470 | At1g13470 | Uncharacterized protein At1g13470 | -4.67 | **6.88E-13** | -1.53 | **5.43E-08** | 3.14 | **1.34E-08** |
| AT2G46400 | WRKY46 | Probable WRKY transcription factor 46 | -4.50 | **1.39E-08** | -0.87 | 1.85E-01 | 3.61 | **3.45E-08** |
| AT2G32680 | RLP23 | Receptor like protein 23 | -4.45 | **1.48E-09** | -1.71 | **4.36E-14** | 2.73 | **3.87E-05** |
| AT3G29000 | CML45 | Probable calcium-binding protein CML45 | -4.31 | **2.03E-05** | | | 3.43 | **3.68E-04** |
| AT2G04495 | At2g04495 | At2g04495 | -4.27 | **2.06E-06** | -1.14 | 4.75E-01 | 3.12 | **3.14E-04** |
| AT3G60540 | T8B10_200 | Protein transport protein Sec61 subunit beta | -4.26 | **4.42E-05** | | | 2.86 | **1.79E-03** |
| AT2G29110 | GLR28 | Glutamate receptor | -4.18 | **1.60E-06** | -0.05 | 9.89E-01 | 4.11 | **4.54E-06** |
| AT3G11010 | RLP34 | Receptor like protein 34 | -4.15 | **5.57E-09** | -1.48 | **3.51E-20** | 2.66 | **1.09E-04** |
| AT2G18690 | At2g18690 | At2g18690/MSF3.7 | -4.14 | **7.94E-06** | -0.31 | 2.54E-01 | 3.82 | **4.09E-05** |
| AT3G23120 | RLP38 | Receptor-like protein 38 | -4.12 | **3.58E-13** | -1.78 | **5.49E-23** | 2.33 | **2.03E-05** |
| AT1G03660 | At1g03660 | Ankyrin-repeat containing protein | -4.12 | **1.59E-06** | | | 4.09 | **2.79E-06** |
| AT1G08050 | At1g08050 | Zinc finger family protein | -4.11 | **3.92E-07** | | | 2.77 | **2.76E-03** |
| AT3G50930 | At3g50930 | AAA+ ATPase domain-containing protein | -4.09 | **2.68E-09** | -0.30 | 5.69E-01 | 3.78 | **5.33E-08** |
| AT4G14365 | XBAT34 | Putative E3 ubiquitin-protein ligase XBAT34 | -4.08 | **2.54E-08** | -1.08 | **3.89E-08** | 2.98 | **3.20E-05** |
| AT1G51800 | IOS1 | LRR receptor-like serine/threonine-protein kinase IOS1 | -4.03 | **1.30E-11** | -0.86 | 5.91E-02 | 3.14 | **3.14E-08** |
| AT2G27660 | At2g27660 | Cysteine/Histidine-rich C1 domain family protein | -3.96 | **3.68E-06** | -0.77 | 5.39E-01 | 3.20 | **4.83E-07** |
| AT1G24145 | At1g24145 | At1g24145 | -3.83 | **2.17E-05** | -0.78 | 1.96E-01 | 3.05 | **2.90E-04** |
| AT4G03450 | At4g03450 | Ankyrin repeat family protein | -3.72 | **1.56E-09** | -0.95 | **2.41E-02** | 2.77 | **4.72E-07** |
| AT1G26420 | FOX5 | Berberine bridge enzyme-like 7 | -3.72 | **2.15E-04** | 0.59 | 7.37E-01 | 4.27 | **1.84E-05** |
| AT1G35710 | At1g35710 | Probable leucine-rich repeat receptor-like protein kinase At1g35710 | -3.70 | **8.98E-09** | -1.35 | **6.56E-08** | 2.34 | **2.83E-05** |
| AT3G51330 | At3g51330 | Eukaryotic aspartyl protease family protein | -3.69 | **4.90E-07** | -0.90 | **2.01E-02** | 2.78 | **1.09E-04** |
| AT5G48657 | At5g48657 | Defense protein-like protein | -3.66 | **1.38E-09** | -0.63 | 2.52E-01 | 3.01 | **3.51E-07** |
| AT4G21850 | MSRB9 | Peptide methionine sulfoxide reductase B9-S-oxide reductase | -3.54 | **1.12E-04** | 0.19 | 9.59E-01 | 3.68 | **6.31E-04** |
| AT2G32140 | At2g32140 | Transmembrane receptor | -3.54 | **1.55E-08** | -0.37 | 9.02E-01 | 3.18 | **3.88E-10** |
| AT3G60415 | At3g60415 | Phosphoglycerate mutase family protein | -3.49 | **4.61E-09** | -0.77 | **1.94E-02** | 2.70 | **2.82E-06** |
| AT1G21250 | WAK1 | Wall-associated receptor kinase 1 | -3.47 | **4.15E-07** | -1.30 | **1.32E-08** | 2.16 | **5.98E-04** |
| AT3G56400 | WRKY70 | Probable WRKY transcription factor 70 | -3.46 | **1.24E-09** | -0.99 | **1.49E-07** | 2.46 | **4.71E-06** |
| AT5G62770 | At5g62770 | Uncharacterized protein | -3.36 | **1.15E-04** | -0.05 | 9.99E-01 | 3.30 | **5.94E-05** |
| AT3G13380 | BRL3 | Receptor-like protein kinase BRI1-like 3 | -3.32 | **7.40E-05** | -0.55 | 3.11E-01 | 2.76 | **1.08E-03** |
| AT5G43420 | ATL16 | RING-H2 finger protein ATL16 | -3.31 | **1.06E-03** | | | 2.21 | **3.54E-02** |
| AT5G27060 | RLP53 | Receptor-like protein 53 | -3.30 | **2.00E-05** | 0.10 | 9.85E-01 | 3.38 | **1.34E-04** |
| AT5G08240 | At5g08240 | Transmembrane protein | -3.30 | **1.57E-05** | -0.07 | 9.94E-01 | 3.21 | **8.64E-06** |
| AT1G76040 | CPK29 | Calcium-dependent protein kinase 29 | -3.28 | **3.48E-04** | -0.49 | **3.69E-02** | 2.79 | **2.87E-03** |
| AT5G67340 | PUB2 | U-box domain-containing protein 2 | -3.28 | **4.00E-06** | -1.03 | **1.41E-05** | 2.24 | **1.99E-03** |
| AT2G20142 | At2g20142 | Toll-Interleukin-Resistance | -3.28 | **7.10E-06** | -0.57 | 4.92E-01 | 2.71 | **6.02E-05** |
| AT3G22231 | PCC1 | Cysteine-rich and transmembrane domain-containing protein PCC1 | -3.27 | **2.36E-07** | -2.23 | **3.63E-15** | 1.04 | 7.48E-02 |
| AT5G25910 | RLP52 | Receptor-like protein 52 | -3.27 | **1.55E-05** | -0.91 | 1.92E-01 | 2.33 | **2.87E-03** |
| AT2G40750 | WRKY54 | Probable WRKY transcription factor 54 | -3.22 | **8.64E-11** | -1.28 | **4.12E-07** | 1.94 | **4.59E-06** |
| AT1G72240 | At1g72240 | Uncharacterized protein | -3.20 | **3.98E-04** | -1.33 | 4.46E-01 | 1.91 | **3.55E-02** |
| AT1G66880 | At1g66880 | Protein kinase superfamily protein | -3.18 | **8.60E-06** | -0.77 | **8.35E-08** | 2.40 | **8.81E-04** |
| AT3G48080 | EDS1B | Protein EDS1B | -3.18 | **6.73E-07** | -1.27 | **8.90E-03** | 1.91 | **5.09E-04** |
| AT1G51790 | At1g51790 | Leucine-rich repeat protein kinase family protein | -3.17 | **1.15E-06** | -1.41 | **3.04E-03** | 1.75 | **3.46E-05** |
| AT4G23810 | WRKY53 | Probable WRKY transcription factor 53 | -3.17 | **2.83E-08** | -1.23 | **6.17E-09** | 1.92 | **1.04E-03** |
| AT4G15610 | CASPL1D1 | CASP-like protein | -3.16 | **2.41E-02** | 0.18 | 9.60E-01 | 3.35 | **1.35E-02** |
| AT3G04220 | At3g04220 | Disease resistance protein family | -3.15 | **1.42E-02** | 0.04 | 9.93E-01 | 3.19 | **5.08E-03** |
| AT4G14370 | DL3225C | Disease resistance protein | -3.12 | **3.68E-06** | -1.04 | 7.15E-02 | 2.08 | **7.90E-04** |
| AT3G26210 | CYP71B23 | Cytochrome P450 71B23 | -3.10 | **3.21E-06** | -0.42 | 1.22E-01 | 2.66 | **3.42E-05** |
| AT3G07195 | At3g07195 | RPM1-interacting protein 4 | -3.10 | **1.41E-06** | -1.05 | 5.23E-02 | 2.03 | **1.48E-03** |
| AT5G60900 | RLK1 | G-type lectin S-receptor-like serine/threonine-protein kinase RLK1 | -3.05 | **1.48E-09** | -1.41 | **2.78E-11** | 1.63 | **8.56E-04** |
| AT5G26920 | CBP60G | Cam-binding protein 60-like G | -3.04 | **1.37E-06** | -0.38 | 2.69E-01 | 2.65 | **2.28E-05** |
| AT1G63720 | F24D7.9 | Uncharacterized protein F24D7.9 | -3.02 | **2.40E-06** | -0.60 | 2.66E-01 | 2.40 | **3.02E-04** |
| AT1G05340 | CYSTM1 | Protein CYSTEINE-RICH TRANSMEMBRANE MODULE 1 | -3.01 | **1.58E-02** | 0.04 | 9.93E-01 | 3.08 | **2.42E-02** |
| AT3G63380 | ACA12 | Calcium-transporting ATPase 12, plasma membrane-type | -2.99 | **3.25E-03** | -0.51 | 3.53E-01 | 2.48 | **1.61E-02** |

| | | | | | | | | |
|---|---|---|---|---|---|---|---|---|
| AT5G18270 | ANAC087 | NAC domain containing protein 87 | -2.87 | **9.74E-03** | | | 4.18 | **3.71E-04** |
| AT5G26170 | At5g26170 | WRKY50 | -2.87 | **2.02E-05** | -1.16 | 5.59E-02 | 1.68 | **7.01E-03** |
| AT1G67920 | At1g67920 | Uncharacterized protein | -2.86 | **1.94E-02** | -0.32 | 9.27E-01 | 2.45 | **3.30E-02** |
| AT3G62600 | ERDJ3B | DnaJ protein ERDJ3B | -2.84 | **7.05E-04** | -0.60 | **5.10E-03** | 2.22 | **9.83E-03** |
| AT4G01870 | At4g01870 | TolB protein-like protein | -2.83 | **5.84E-05** | -0.26 | 6.90E-01 | 2.53 | **3.76E-04** |
| AT3G11402 | At3g11402 | Cysteine/Histidine-rich C1 domain family protein | -2.75 | **2.79E-07** | -1.15 | **8.54E-05** | 1.58 | **6.35E-03** |
| AT5G41750 | MUF8.3 | Disease resistance protein | -2.74 | **4.45E-07** | -0.08 | 9.32E-01 | 2.65 | **1.96E-08** |
| AT3G21150 | BBX32 | B-box zinc finger protein 32 | -2.73 | **2.15E-04** | | | 2.10 | **2.92E-02** |
| AT4G14400 | ACD6 | Protein ACCELERATED CELL DEATH 6 | -2.72 | **5.37E-10** | -1.32 | **7.34E-19** | 1.38 | **1.16E-03** |
| AT3G22160 | VQ22 | VQ motif-containing protein 22 | -2.71 | **1.02E-05** | -0.27 | 7.25E-01 | 2.43 | **1.21E-04** |
| AT5G01550 | LECRK-VI3 | Lectin receptor kinase a4.1 | -2.70 | **1.79E-02** | 1.22 | 4.70E-01 | 3.93 | **1.93E-04** |
| AT3G25020 | RLP42 | Receptor-like protein 42 | -2.67 | **1.59E-06** | -0.50 | **4.78E-03** | 2.15 | **1.08E-04** |
| AT5G64310 | AGP1 | Classical arabinogalactan protein 1 | -2.66 | **7.65E-09** | -0.08 | 9.74E-01 | 2.57 | **9.98E-09** |
| AT5G63130 | At5g63130 | hypothetical protein | -2.64 | **7.04E-06** | -0.74 | 6.15E-01 | 1.85 | **2.04E-04** |
| AT2G30550 | DALL3 | Alpha/beta-Hydrolases superfamily protein | -2.62 | **1.74E-03** | -0.17 | 7.24E-01 | 2.44 | **3.87E-03** |
| AT1G76970 | At1g76970 | Target of Myb protein 1 | -2.61 | **6.05E-06** | -0.46 | 3.01E-01 | 2.14 | **3.73E-04** |
| AT1G34420 | At1g34420 | Protein kinase domain-containing protein | -2.58 | **4.29E-05** | -0.36 | 5.41E-01 | 2.21 | **3.92E-04** |
| AT3G61280 | T20K12.180 | Uncharacterized protein T20K12.180 | -2.58 | **2.41E-06** | -0.35 | 3.78E-01 | 2.22 | **3.32E-06** |
| AT4G23220 | At4g23220 | Uncharacterized protein | -2.53 | **2.74E-06** | -0.34 | 3.64E-01 | 2.18 | **2.03E-05** |
| AT3G52430 | PAD4 | Lipase-like PAD4 | -2.52 | **4.70E-07** | -0.32 | 3.13E-01 | 2.19 | **3.42E-05** |
| AT1G72910 | At1g72910 | Similar to part of disease resistance protein | -2.52 | **4.88E-06** | -0.42 | 5.18E-01 | 2.09 | **2.82E-06** |
| AT2G17120 | LYM2 | LysM domain-containing GPI-anchored protein 2 | -2.51 | **2.12E-04** | -0.88 | **2.25E-09** | 1.62 | **2.33E-02** |
| AT3G07520 | At3g07520 | Glutamate receptor | -2.45 | **9.32E-05** | -0.74 | **2.14E-02** | 1.68 | **8.63E-03** |
| AT3G06890 | At3g06890 | hypothetical protein | -2.44 | **2.12E-03** | | | 2.79 | **4.41E-04** |
| AT1G72900 | At1g72900 | Similar to part of disease resistance protein | -2.42 | **4.14E-05** | -0.39 | 5.89E-01 | 2.03 | **5.48E-04** |
| AT2G46430 | CNGC3 | Probable cyclic nucleotide-gated ion channel 3 | -2.42 | **5.06E-08** | -0.99 | **1.87E-09** | 1.42 | **1.67E-03** |
| AT3G09010 | MZB10.4 | Putative receptor ser/thr protein kinase | -2.39 | **2.11E-05** | -0.53 | 4.70E-01 | 1.85 | **1.43E-04** |
| AT4G04695 | CPK31 | Calcium-dependent protein kinase 31 | -2.39 | **5.84E-05** | -1.57 | **1.60E-03** | 0.81 | 2.13E-01 |
| AT5G25930 | T1N24.22 | Kinase family with leucine-rich repeat domain-containing protein | -2.38 | **3.33E-05** | -0.85 | **2.82E-04** | 1.52 | **1.01E-02** |
| AT2G29720 | CTF2B | FAD/NAD | -2.38 | **2.38E-05** | -0.77 | 3.17E-01 | 1.58 | **1.23E-02** |
| AT2G41640 | At2g41640 | Glycosyltransferase family 61 protein | -2.37 | **2.88E-05** | 0.28 | 8.70E-01 | 2.64 | **8.73E-14** |
| AT5G35735 | At5g35735 | Cytochrome b561 and DOMON domain-containing protein At5g35735 | -2.35 | **6.05E-04** | 0.19 | 6.22E-01 | 2.52 | **1.74E-04** |
| AT1G79680 | WAKL10 | Wall-associated receptor kinase-like 10 | -2.34 | **4.38E-02** | -0.06 | 1.00E+00 | 2.27 | **2.42E-02** |
| AT2G01540 | CAR10 | Protein C2-DOMAIN ABA-RELATED 10 | -2.34 | **1.01E-04** | -1.32 | **9.80E-03** | 1.00 | 1.74E-01 |
| AT2G25510 | At2g25510 | Transmembrane protein | -2.33 | **6.97E-05** | -0.55 | 1.12E-01 | 1.77 | **3.28E-04** |
| AT1G32940 | SBT35 | Subtilase family protein | -2.33 | **8.04E-03** | -0.08 | 9.50E-01 | 2.24 | **1.10E-02** |
| AT1G67000 | LRK10L-2.8 | LEAF RUST 10 DISEASE-RESISTANCE LOCUS RECEPTOR-LIKE PROTEIN KINASE-like 2.8 | -2.31 | **1.71E-04** | 0.45 | 6.40E-01 | 2.76 | **1.10E-06** |
| AT5G19240 | At5g19240 | Uncharacterized GPI-anchored protein At5g19240 | -2.31 | **2.11E-05** | -0.99 | **1.25E-08** | 1.30 | **1.90E-02** |
| AT3G23010 | RLP36 | Receptor like protein | -2.29 | **9.49E-05** | -0.86 | **2.37E-02** | 1.42 | **5.10E-03** |
| AT1G56660 | At1g56660 | Uncharacterized protein | -2.24 | **2.58E-05** | 0.70 | 3.88E-01 | 2.92 | **6.42E-07** |
| AT4G16660 | HSP70 | Heat shock protein 70 | -2.23 | **8.03E-04** | -0.36 | 6.53E-02 | 1.86 | **5.77E-03** |
| AT2G40095 | At2g40095 | Alpha/beta hydrolase related protein | -2.22 | **1.81E-05** | -0.74 | **1.12E-02** | 1.45 | **6.93E-03** |
| AT4G13820 | At4g13820 | Disease resistance like protein | -2.21 | **4.21E-03** | -0.15 | 9.51E-01 | 2.03 | **3.41E-04** |
| AT3G52400 | SYP122 | Syntaxin-122 | -2.20 | **1.21E-05** | 0.11 | 8.78E-01 | 2.30 | **2.79E-06** |
| AT5G39050 | PMAT1 | Phenolic glucoside malonyltransferase 1 | -2.20 | **1.49E-04** | -0.11 | 8.59E-01 | 2.07 | **2.32E-04** |
| AT1G64065 | At1g64065 | Late embryogenesis abundant protein At1g64065 | -2.19 | **1.67E-02** | 0.12 | 9.76E-01 | 2.30 | **3.01E-02** |
| AT4G15233 | ABCG42 | ABC-2 and Plant PDR ABC-type transporter family protein | -2.16 | **1.70E-07** | -0.69 | **2.81E-02** | 1.46 | **6.31E-05** |
| AT5G41740 | MUF8.2 | Disease resistance protein | -2.16 | **3.92E-08** | -0.18 | 6.89E-01 | 1.97 | **3.45E-08** |
| AT2G29120 | GLR2.7 | Glutamate receptor 2.7 | -2.16 | **1.13E-03** | -0.01 | 1.00E+00 | 2.14 | **1.00E-03** |
| AT1G15670 | At1g15670 | F-box/kelch-repeat protein At1g15670 | -2.10 | **1.06E-02** | 0.00 | 1.00E+00 | 2.09 | **1.45E-06** |
| AT4G11000 | F25I24.210 | Uncharacterized protein AT4g11000 | -2.09 | **6.08E-07** | -1.10 | **8.34E-06** | 0.98 | **3.73E-02** |
| AT3G45620 | At3g45620 | Transducin/WD40 repeat-like superfamily protein | -2.08 | **3.41E-04** | -0.57 | 1.00E-01 | 1.50 | **8.34E-03** |
| AT1G66090 | At1g66090 | Disease resistance protein | -2.05 | **1.15E-05** | -0.02 | 1.00E+00 | 2.02 | **4.70E-06** |
| AT4G29780 | F27B13.20 | Uncharacterized protein AT4g29780 | -2.05 | **3.29E-04** | 0.75 | 1.13E-01 | 2.78 | **1.58E-07** |
| AT2G23680 | At2g23680 | Uncharacterized protein | -2.03 | **3.85E-04** | -0.21 | 8.34E-01 | 1.81 | **9.38E-04** |
| AT5G42530 | At5g42530 | | -2.01 | **6.18E-06** | -1.01 | **1.29E-04** | 0.99 | **3.29E-03** |
| AT1G28370 | ERF11 | ERF domain protein 11 | -2.01 | **2.38E-02** | 0.32 | 8.82E-01 | 2.31 | **2.59E-08** |
| AT3G07580 | At3g07580 | At3g07580 | -2.00 | **1.53E-02** | -0.53 | 4.92E-01 | 1.45 | 9.00E-02 |
| AT4G34390 | XLG2 | Extra-large GTP-binding protein 2 | -1.99 | **4.30E-05** | -0.32 | 1.26E-01 | 1.65 | **9.14E-04** |
| AT5G15870 | At5g15870 | glucan endo-1,3-beta-D-glucosidase | -1.99 | **4.99E-05** | -0.36 | 1.51E-01 | 1.61 | **1.49E-03** |
| AT1G59590 | ZCF37 | At1g59590 | -1.98 | **1.39E-05** | -0.37 | 4.51E-01 | 1.60 | **5.58E-04** |
| AT1G10040 | At1g10040 | Alpha/beta-Hydrolases superfamily protein | -1.98 | **4.86E-02** | | | 2.64 | **1.04E-02** |
| AT1G22400 | UGT85A1 | UDP-glycosyltransferase 85A1 | -1.94 | **4.45E-04** | -0.22 | 7.41E-01 | 1.71 | **9.31E-04** |
| AT1G74440 | At1g74440 | ER membrane protein, putative | -1.92 | **1.62E-06** | -1.00 | **2.34E-04** | 0.90 | 5.21E-02 |
| AT5G18780 | At5g18780 | F-box/RNI-like superfamily protein | -1.91 | **8.28E-06** | 0.04 | 9.95E-01 | 1.94 | **1.49E-05** |
| AT3G47090 | F13I12.140 | Leucine-rich repeat protein kinase family protein | -1.91 | **2.09E-06** | -0.26 | 5.64E-01 | 1.64 | **1.56E-04** |
| AT3G11820 | SYP121 | Syntaxin-121 | -1.90 | **4.05E-05** | -0.50 | **2.23E-02** | 1.38 | **5.53E-03** |
| AT4G08850 | MIK2 | MDIS1-interacting receptor like kinase 2 | -1.90 | **1.63E-04** | -0.58 | **1.04E-04** | 1.31 | **1.13E-02** |
| AT5G39090 | At5g39090 | Uncharacterized protein | -1.90 | **9.12E-04** | 0.84 | 2.20E-01 | 2.72 | **2.78E-06** |
| AT2G41835 | SAP11 | Zinc finger AN1 and C2H2 domain-containing stress-associated protein 11 | -1.90 | **2.65E-04** | -0.44 | 3.78E-01 | 1.44 | **1.06E-02** |
| AT5G07770 | FH16 | Formin-like protein | -1.88 | **2.64E-08** | -0.29 | 3.34E-01 | 1.58 | **3.04E-06** |
| AT5G66910 | At5g66910 | Probable disease resistance protein At5g66910 | -1.88 | **2.86E-06** | -0.64 | **3.67E-02** | 1.22 | **3.41E-03** |
| AT1G68690 | PERK9 | non-specific serine/threonine protein kinase | -1.88 | **2.90E-02** | 0.09 | 9.25E-01 | 1.94 | **2.00E-02** |
| AT5G20400 | At5g20400 | hypothetical protein | -1.87 | **1.53E-02** | 0.38 | 8.31E-01 | 2.24 | **2.35E-03** |
| AT4G36090 | ALKBH9C | Oxidoreductase, 2OG-Fe | -1.86 | **3.19E-05** | -0.42 | 4.83E-01 | 1.43 | **4.12E-03** |
| AT2G29990 | NDA2 | Internal alternative NAD | -1.86 | **6.17E-04** | 0.02 | 1.00E+00 | 1.87 | **6.55E-04** |
| AT2G13790 | SERK4 | Somatic embryogenesis receptor kinase 4 | -1.86 | **8.61E-07** | -0.61 | **3.28E-05** | 1.23 | **1.64E-03** |
| AT1G72950 | At1g72950 | TIR domain-containing protein | -1.86 | **1.43E-03** | | | 1.78 | **1.87E-02** |
| AT1G18210 | CML27 | Probable calcium-binding protein CML27 | -1.84 | **2.26E-04** | -0.14 | 8.60E-01 | 1.68 | **1.16E-03** |
| AT3G11080 | RLP35 | Receptor-like protein 35 | -1.81 | **2.70E-03** | -0.06 | 9.42E-01 | 1.73 | **4.61E-03** |
| AT1G61260 | F11P17.2 | F11P17.2 protein | -1.81 | **2.91E-03** | 1.15 | 2.67E-01 | 2.96 | **2.77E-03** |
| AT2G28400 | S40-4 | Protein S40-4 | -1.80 | **1.15E-02** | | | 2.08 | **1.01E-02** |
| AT3G26600 | At3g26600 | | -1.80 | **2.30E-05** | -0.42 | 2.45E-01 | 1.37 | **2.59E-03** |
| AT2G24600 | At2g24600 | Ankyrin repeat family protein | -1.79 | **2.86E-05** | -0.24 | 6.38E-01 | 1.54 | **1.16E-05** |
| AT1G77500 | At1g77500 | Uncharacterized protein | -1.79 | **6.31E-04** | -0.33 | 2.90E-01 | 1.44 | **7.46E-03** |
| AT2G46620 | At2g46620 | AAA-ATPase At2g46620 | -1.79 | **2.15E-03** | -0.01 | 1.00E+00 | 1.76 | **2.59E-03** |
| AT3G26170 | CYP71B19 | Cytochrome P450 71B19 | -1.78 | **1.48E-05** | -0.19 | 7.09E-01 | 1.58 | **1.42E-05** |
| AT3G49780 | PSK3 | Phytosulfokines 3 | -1.78 | **1.47E-02** | -0.16 | 8.03E-01 | 1.61 | **1.85E-02** |
| AT5G61250 | At5g61250 | Heparanase-like protein 2 | -1.77 | **6.64E-04** | -0.10 | 9.65E-01 | 1.66 | **1.21E-03** |
| AT5G46520 | VICTR | ADP-ribosyl cyclase/cyclic ADP-ribose hydrolase | -1.76 | **5.09E-04** | 0.17 | 9.20E-01 | 1.91 | **1.23E-04** |

| | | | | | | | | |
|---|---|---|---|---|---|---|---|---|
| AT5G12930 | At5g12930 | Inactive rhomboid protein | -1.76 | **1.53E-02** | -0.55 | 5.61E-01 | 1.19 | 1.84E-01 |
| AT3G55950 | CCR3 | Putative serine/threonine-protein kinase-like protein CCR3 | -1.75 | **1.25E-03** | -0.41 | 2.85E-01 | 1.32 | **1.31E-02** |
| AT1G17600 | SOC3 | Disease resistance protein | -1.75 | **2.35E-06** | -0.86 | **2.14E-02** | 0.88 | 5.86E-02 |
| AT5G46230 | At5g46230 | | -1.74 | **1.57E-02** | -0.21 | 8.55E-01 | 1.53 | **3.15E-02** |
| AT2G19130 | At2g19130 | G-type lectin S-receptor-like serine/threonine-protein kinase At2g19130 | -1.73 | **7.27E-03** | 0.01 | 1.00E+00 | 1.73 | **4.13E-03** |
| AT1G05570 | CALS1 | 1,3-beta-glucan synthase | -1.73 | **3.36E-05** | 0.04 | 9.38E-01 | 1.76 | **1.71E-05** |
| AT2G25110 | SDF2 | Stromal cell-derived factor 2-like protein | -1.70 | **7.81E-03** | -0.21 | 4.65E-01 | 1.48 | **2.11E-02** |
| AT3G26220 | At3g26220 | At3g26230 | -1.69 | **4.19E-05** | 0.03 | 9.89E-01 | 1.71 | **9.56E-06** |
| AT5G44820 | At5g44820 | Nucleotide-diphospho-sugar transferase domain-containing protein | -1.69 | **3.35E-06** | -0.74 | **7.34E-03** | 0.94 | **2.70E-02** |
| AT3G63390 | MAA21_20 | Uncharacterized protein MAA21_20 | -1.69 | **1.42E-03** | 0.21 | 8.38E-01 | 1.90 | **3.52E-04** |
| AT3G49120 | PER34 | Peroxidase 34 | -1.69 | **8.21E-03** | -0.03 | 9.78E-01 | 1.65 | **4.96E-03** |
| AT1G69720 | HO3 | Heme oxygenase 3, chloroplastic | -1.69 | **6.79E-04** | 0.08 | 9.71E-01 | 1.76 | **1.60E-04** |
| AT1G02220 | NAC003 | NAC domain-containing protein 3 | -1.69 | **8.55E-03** | -0.08 | 9.72E-01 | 1.58 | **9.71E-03** |
| AT3G44400 | T22K7_80 | ADP-ribosyl cyclase/cyclic ADP-ribose hydrolase | -1.68 | **1.53E-05** | -0.27 | 3.94E-01 | 1.40 | **3.09E-04** |
| AT4G33360 | FLDH | NAD | -1.64 | **1.46E-03** | -0.65 | 2.17E-01 | 0.98 | **2.90E-02** |
| AT2G03510 | At2g03510 | Expressed protein | -1.64 | **1.88E-03** | -0.60 | **1.26E-03** | 1.02 | 8.51E-02 |
| AT2G33020 | RLP24 | Receptor like protein 24 | -1.64 | **2.17E-02** | -0.66 | 4.87E-01 | 0.97 | 1.84E-01 |
| AT3G45290 | MLO3 | MLO-like protein 3 | -1.64 | **1.29E-04** | -0.23 | 6.63E-01 | 1.39 | **2.64E-03** |
| AT4G01540 | NTM1 | NAC with transmembrane motif1 | -1.64 | **1.90E-02** | -0.04 | 9.93E-01 | 1.60 | **1.82E-02** |
| AT4G22980 | At4g22980 | Molybdenum cofactor sulfurase-like protein | -1.63 | **2.37E-03** | -0.39 | 5.61E-01 | 1.22 | **4.18E-02** |
| AT5G54710 | At5g54710 | Similarity to ankyrin-like protein | -1.62 | **7.25E-04** | -0.07 | 8.82E-01 | 1.53 | **9.65E-04** |
| AT4G23470 | At4g23470 | PLAC8 family protein | -1.62 | **2.63E-04** | -0.35 | **2.62E-02** | 1.25 | **6.70E-03** |
| AT1G25400 | At1g25400 | Transmembrane protein | -1.61 | **2.54E-02** | 0.72 | 5.46E-01 | 2.32 | **1.48E-04** |
| AT1G64610 | At1g64610 | Transducin/WD40 repeat-like superfamily protein | -1.60 | **2.73E-04** | 0.06 | 9.68E-01 | 1.64 | **2.82E-04** |
| AT4G11350 | At4g11350 | Transferring glycosyl group transferase | -1.60 | **2.19E-02** | 0.28 | 9.14E-01 | 1.85 | **6.66E-03** |
| AT3G19970 | At3g19970 | | -1.59 | **2.74E-02** | -0.03 | 1.00E+00 | 1.56 | **2.99E-02** |
| AT4G26060 | At4g26060 | At4g26060 | -1.58 | **4.44E-03** | -0.24 | 8.36E-01 | 1.34 | **5.92E-03** |
| AT5G24810 | At5g24810 | ABC1 family protein | -1.58 | **7.50E-05** | -0.39 | **2.50E-02** | 1.18 | **4.36E-03** |
| AT3G16565 | At3g16565 | Threonyl and alanyl tRNA synthetase second additional domain-containing protein | -1.57 | **1.48E-03** | 0.37 | 6.18E-01 | 1.93 | **1.43E-04** |
| AT4G36150 | At4g36150 | ADP-ribosyl cyclase/cyclic ADP-ribose hydrolase | -1.57 | **1.22E-05** | -0.86 | **1.11E-05** | 0.69 | 9.01E-02 |
| AT3G09830 | PCRK1 | Serine/threonine-protein kinase PCRK1 | -1.57 | **4.32E-04** | -0.42 | 1.99E-01 | 1.15 | **8.05E-03** |
| AT5G01850 | At5g01850 | Protein kinase ATN1-like protein | -1.57 | **1.73E-03** | -0.04 | 9.88E-01 | 1.51 | **3.29E-03** |
| AT5G04720 | At5g04720 | Probable disease resistance protein At5g04720 | -1.57 | **5.01E-04** | -0.27 | 3.42E-01 | 1.29 | **4.13E-03** |
| AT1G19960 | At1g19960 | Uncharacterized protein | -1.56 | **1.66E-03** | -0.77 | **9.93E-03** | 0.78 | 1.53E-01 |
| AT1G69610 | T6C23.19 | Uncharacterized protein F24J1.24 | -1.56 | **3.02E-02** | -0.78 | 1.22E-01 | 0.78 | 3.01E-01 |
| AT1G27770 | ACA1 | Calcium-transporting ATPase | -1.55 | **1.69E-04** | -0.27 | 4.62E-01 | 1.27 | **2.82E-04** |
| AT1G21370 | At1g21370 | | -1.55 | **2.34E-03** | 0.01 | 1.00E+00 | 1.53 | **3.05E-03** |
| AT4G11300 | At4g11300 | ROH1, | -1.52 | **3.16E-04** | -0.63 | 4.11E-01 | 0.87 | 1.05E-01 |
| AT2G23810 | TET8 | Tetraspanin-8 | -1.51 | **8.92E-03** | 0.08 | 9.20E-01 | 1.58 | **1.01E-03** |
| AT1G13750 | PAP1 | Probable inactive purple acid phosphatase 1 | -1.51 | **6.77E-03** | -0.40 | 3.93E-01 | 1.10 | **1.59E-02** |
| AT1G19180 | JAZ1 | Protein TIFY | -1.51 | **2.71E-02** | 0.93 | 8.44E-02 | 2.43 | **1.34E-08** |
| AT3G05650 | RLP32 | Receptor-like protein 32 | -1.50 | **1.54E-03** | 0.10 | 8.43E-01 | 1.59 | **3.85E-04** |
| AT2G16870 | At2g16870 | Disease resistance protein | -1.50 | **9.75E-05** | -0.57 | 1.73E-01 | 0.91 | **2.50E-02** |
| AT5G08380 | AGAL1 | Alpha-galactosidase | -1.49 | **1.62E-02** | 0.07 | 9.50E-01 | 1.55 | **1.25E-02** |
| AT5G21090 | LRR1 | Leucine-rich repeat protein 1 | -1.49 | **4.81E-02** | -0.01 | 9.97E-01 | 1.47 | **4.11E-02** |
| AT3G19010 | 2-OXOGLUT | 2-oxoglutarate | -1.49 | **4.49E-04** | -0.26 | 2.81E-01 | 1.21 | **3.41E-03** |
| AT1G67800 | RGLG5 | Copine | -1.49 | **2.01E-03** | -0.13 | 7.56E-01 | 1.35 | **5.31E-03** |
| AT5G54720 | At5g54720 | Uncharacterized protein | -1.49 | **4.30E-04** | 0.02 | 9.79E-01 | 1.50 | **2.42E-04** |
| AT1G52780 | At1g52780 | PII, uridylyltransferase | -1.49 | **3.47E-04** | -0.32 | 9.12E-02 | 1.16 | **6.45E-03** |
| AT3G26230 | At3g26230 | Uncharacterized protein | -1.47 | **1.20E-03** | -0.43 | 6.66E-02 | 1.03 | **2.39E-02** |
| AT1G13990 | At1g13990 | Plant/protein | -1.47 | **2.26E-02** | 0.09 | 8.94E-01 | 1.55 | **6.01E-03** |
| AT2G19710 | At2g19710 | Uncharacterized protein At2g19710 | -1.47 | **5.11E-04** | -0.22 | 6.14E-01 | 1.24 | **3.84E-03** |
| AT3G15760 | At3g15760 | Transmembrane protein | -1.47 | **1.26E-03** | -0.62 | 3.28E-01 | 0.83 | 9.16E-02 |
| AT3G55470 | At3g55470 | Calcium-dependent lipid-binding | -1.47 | **3.35E-03** | 0.16 | 9.22E-01 | 1.61 | **7.50E-03** |
| AT1G18890 | CPK10 | Calcium-dependent protein kinase 10 | -1.46 | **1.02E-02** | -0.25 | 4.82E-01 | 1.20 | **4.03E-02** |
| AT5G45500 | MFC19.17 | RNI-like superfamily protein | -1.45 | **4.00E-04** | -0.26 | 2.95E-01 | 1.18 | **5.58E-03** |
| AT1G23140 | At1g23140 | C2 domain-containing protein | -1.45 | **1.79E-02** | -0.69 | 7.22E-01 | 0.74 | 4.25E-01 |
| AT5G42440 | At5g42440 | Protein kinase domain-containing protein | -1.45 | **1.65E-03** | -0.15 | 8.60E-01 | 1.29 | **3.87E-03** |
| AT1G74200 | RLP16 | Putative receptor-like protein 16 | -1.44 | **5.98E-03** | 0.39 | 8.75E-01 | 1.75 | **4.17E-04** |
| AT1G80610 | At1g80610 | At1g80610/T21F11_6 | -1.44 | **3.08E-04** | -0.11 | 8.73E-01 | 1.31 | **8.08E-04** |
| AT4G18890 | BEH3 | Protein BZR1 homolog | -1.42 | **1.16E-02** | -0.05 | 9.93E-01 | 1.37 | **2.10E-02** |
| AT1G50750 | At1g50750 | Aminotransferase-like, mobile domain protein | -1.42 | **1.15E-02** | -0.33 | 8.87E-01 | 1.08 | 8.01E-02 |
| AT3G45640 | MPK3 | Mitogen-activated protein kinase 3 | -1.42 | **8.39E-04** | -0.02 | 9.88E-01 | 1.38 | **3.52E-04** |
| AT4G00955 | At4g00955 | Uncharacterized protein At4g00955 | -1.42 | **6.05E-06** | -0.60 | 5.36E-02 | 0.81 | **8.45E-03** |
| AT2G30020 | At2g30020 | Probable protein phosphatase 2C 25 | -1.41 | **9.89E-07** | 0.59 | **2.15E-02** | 1.99 | **2.35E-13** |
| AT5G45110 | NPR3 | NPR1-like protein 3 | -1.41 | **4.99E-04** | 0.04 | 9.65E-01 | 1.44 | **4.00E-05** |
| AT4G01010 | CNGC13 | Putative cyclic nucleotide-gated ion channel 13 | -1.40 | **1.26E-02** | 0.15 | 7.76E-01 | 1.54 | **3.01E-03** |
| AT2G31990 | At2g31990 | Exostosin family protein | -1.39 | **1.83E-02** | -0.03 | 1.00E+00 | 1.35 | **1.90E-02** |
| AT1G16260 | At1g16260 | Wall-associated receptor kinase-like | -1.38 | **4.85E-05** | -0.51 | **5.01E-03** | 0.86 | **9.48E-03** |
| AT4G09570 | CPK4 | Calcium-dependent protein kinase 4 | -1.38 | **3.45E-03** | -0.11 | 9.18E-01 | 1.26 | **5.83E-03** |
| AT3G49370 | CRK6 | CDPK-related kinase 6 | -1.38 | **2.45E-03** | -0.18 | 9.05E-01 | 1.19 | **1.02E-02** |
| AT4G25390 | At4g25390 | Receptor-like serine/threonine-protein kinase At4g25390 | -1.37 | **3.72E-02** | 0.33 | 6.98E-01 | 1.69 | **5.58E-03** |
| AT1G65800 | RK2 | Receptor kinase 2 | -1.37 | **6.82E-04** | -0.25 | 4.11E-01 | 1.10 | **2.13E-03** |
| AT1G62422 | At1g62422 | Uncharacterized protein | -1.33 | **3.60E-02** | -0.38 | 6.24E-01 | 0.94 | 1.67E-01 |
| AT3G26090 | RGS1 | REGULATOR OF G-PROTEIN SIGNALING 1 | -1.32 | **3.46E-03** | -0.40 | 2.58E-01 | 0.91 | 5.29E-02 |
| AT2G28940 | PBL37 | At2g28940 | -1.32 | **1.66E-03** | 0.24 | 8.42E-01 | 1.55 | **1.91E-03** |
| AT5G39785 | At5g39780 | Gb\|AAF22924.1 | -1.31 | **5.69E-03** | -0.03 | 1.00E+00 | 1.26 | **1.21E-02** |
| AT3G16510 | At3g16510 | | -1.30 | **5.31E-03** | -0.38 | 3.94E-01 | 0.91 | **2.34E-02** |
| AT1G69730 | WAKL9 | Wall-associated receptor kinase-like 9 | -1.30 | **1.57E-03** | -0.32 | 3.35E-01 | 0.97 | **5.73E-03** |
| AT1G72930 | TIR | Toll/interleukin-1 receptor-like protein | -1.29 | **6.18E-04** | -0.38 | 1.29E-01 | 0.90 | **7.17E-03** |
| AT5G58120 | At5g58120 | ADP-ribosyl cyclase/cyclic ADP-ribose hydrolase | -1.29 | **1.13E-03** | -0.10 | 9.04E-01 | 1.18 | **5.17E-04** |
| AT5G11650 | MAGL13 | Alpha/beta-Hydrolases superfamily protein | -1.29 | **2.73E-03** | -0.46 | 4.37E-01 | 0.81 | 8.17E-02 |
| AT5G36930 | MLF18.50 | Disease resistance protein | -1.27 | **6.97E-05** | -0.52 | **2.61E-04** | 0.73 | **3.78E-02** |
| AT1G23710 | At1g23710 | At1g23710 | -1.26 | **9.31E-03** | -0.28 | 8.11E-01 | 0.97 | 1.04E-01 |
| AT2G03120 | SPP | Signal peptide peptidase | -1.26 | **6.41E-03** | -0.15 | 6.49E-01 | 1.09 | **2.07E-02** |
| AT2G40270 | At2g40270 | Protein kinase family protein | -1.26 | **1.53E-03** | -0.29 | 1.09E-01 | 0.95 | **2.02E-02** |
| AT1G03370 | At1g03370 | C2 calcium/lipid-binding and GRAM domain containing protein | -1.26 | **3.06E-04** | -0.48 | **5.85E-03** | 0.76 | **3.58E-02** |
| AT4G22780 | ACR7 | ACT domain-containing protein ACR7 | -1.25 | **3.75E-04** | -0.06 | 9.55E-01 | 1.18 | **4.21E-04** |
| AT3G56800 | CAM3 | Calmodulin-3 | -1.25 | **3.98E-02** | -0.19 | 5.88E-01 | 1.05 | 6.89E-02 |
| AT1G22280 | PAPP2C | Phytochrome-associated protein phosphatase type 2C | -1.24 | **2.15E-03** | -0.21 | 4.97E-01 | 1.02 | **1.02E-02** |

| | | | | | | | | |
|---|---|---|---|---|---|---|---|---|
| AT1G33610 | At1g33610 | Leucine-rich repeat-containing N-terminal plant-type domain-containing protein | -1.24 | **4.85E-04** | -1.08 | **8.83E-06** | 0.14 | 8.17E-01 |
| AT5G47580 | At5g47580 | ASG7 | -1.23 | **3.15E-02** | 0.00 | 1.00E+00 | 1.21 | **2.93E-02** |
| AT1G21270 | WAK2 | Wall-associated receptor kinase 2 | -1.23 | **2.20E-03** | -0.46 | **4.77E-03** | 0.76 | 6.86E-02 |
| AT1G31540 | T8E3.20 | ADP-ribosyl cyclase/cyclic ADP-ribose hydrolase | -1.23 | **2.59E-03** | -0.30 | 3.27E-01 | 0.91 | **8.74E-03** |
| AT1G59870 | ABCG36 | ABC transporter G family member 36 | -1.23 | **8.33E-03** | 0.09 | 7.81E-01 | 1.30 | **2.79E-03** |
| AT5G57035 | At5g57035 | U-box domain-containing protein kinase family protein | -1.22 | **3.07E-03** | -0.37 | 5.21E-02 | 0.84 | 5.47E-02 |
| AT3G02520 | GRF7 | 14-3-3-like protein GF14 nu | -1.22 | **2.71E-02** | -0.28 | 2.77E-01 | 0.93 | 1.07E-01 |
| AT3G16860 | COBL8 | COBRA-like protein 8 | -1.22 | **2.79E-03** | 0.40 | 5.17E-01 | 1.61 | **1.47E-06** |
| AT3G58600 | At3g58600 | NECAP PHear domain-containing protein | -1.22 | **1.25E-03** | -0.23 | 6.48E-01 | 0.97 | **7.71E-03** |
| AT1G55610 | BRL1 | Serine/threonine-protein kinase BRI1-like 1 | -1.20 | **2.26E-02** | -0.40 | 6.00E-01 | 0.78 | 1.01E-01 |
| AT3G13330 | PA200 | Proteasome activator subunit 4 | -1.20 | **1.86E-03** | -0.19 | 3.93E-01 | 1.00 | **8.86E-03** |
| AT4G16960 | SIKIC3 | Disease resistance protein | -1.20 | **2.33E-03** | -0.08 | 9.67E-01 | 1.11 | **9.16E-03** |
| AT3G21810 | At3g21810 | Zinc finger C-x8-C-x5-C-x3-H type family protein | -1.20 | **1.42E-02** | -0.80 | 6.40E-02 | 0.39 | 4.56E-01 |
| AT1G32127 | F3C3.9 | Uncharacterized protein F3C3.9 | -1.19 | **1.90E-02** | -0.67 | 2.47E-01 | 0.50 | 4.38E-01 |
| AT3G17700 | CNBT1 | Cyclic nucleotide-binding transporter 1 | -1.18 | **4.83E-04** | -0.24 | 4.16E-01 | 0.93 | **6.89E-03** |
| AT5G02290 | PBL11 | Probable serine/threonine-protein kinase PBL11 | -1.18 | **2.17E-02** | -0.26 | 5.80E-01 | 0.91 | **2.92E-02** |
| AT1G25580 | At1g25580 | NAC domain-containing protein | -1.17 | **2.61E-02** | -0.35 | 6.58E-01 | 0.82 | 7.79E-02 |
| AT1G32870 | NAC13 | NAC domain protein 13 | -1.16 | **4.16E-03** | -0.39 | 5.64E-01 | 0.75 | 7.49E-02 |
| AT3G46930 | RAF43 | Protein kinase superfamily protein | -1.16 | **2.52E-02** | 0.40 | 6.58E-01 | 1.54 | **8.95E-04** |
| AT3G21630 | At3g21630 | Protein kinase domain-containing protein | -1.15 | **9.58E-04** | -0.14 | 7.03E-01 | 1.00 | **4.46E-03** |
| AT3G03560 | At3g03560 | Uncharacterized protein | -1.15 | **1.04E-03** | -0.16 | 7.38E-01 | 0.97 | **6.19E-03** |
| AT5G18490 | At5g18490 | At5g18490 | -1.11 | **1.52E-02** | 0.07 | 9.59E-01 | 1.16 | **3.55E-03** |
| AT1G67470 | ZRK12 | Inactive serine/threonine-protein kinase ZRK12 | -1.10 | **8.92E-03** | -0.40 | 4.37E-01 | 0.68 | 7.02E-02 |
| AT1G55450 | At1g55450 | S-adenosyl-L-methionine-dependent methyltransferases superfamily protein | -1.10 | **1.67E-02** | -0.46 | 6.36E-02 | 0.63 | 1.36E-01 |
| AT1G74280 | At1g74280 | Alpha/beta-Hydrolases superfamily protein | -1.10 | **7.39E-03** | -0.10 | 9.50E-01 | 0.98 | **1.53E-02** |
| AT2G11520 | CRCK3 | Calmodulin-binding receptor-like cytoplasmic kinase 3 | -1.10 | **1.50E-02** | -0.25 | 3.64E-01 | 0.83 | 7.92E-02 |
| AT2G07180 | PBL17 | Probable serine/threonine-protein kinase PBL17 | -1.10 | **1.42E-03** | -0.01 | 1.00E+00 | 1.08 | **1.46E-03** |
| AT5G61380 | APRR1 | Two-component response regulator-like APRR1 | -1.10 | **4.98E-03** | -0.13 | 8.75E-01 | 0.95 | **1.60E-02** |
| AT4G01720 | WRKY47 | Probable WRKY transcription factor 47 | -1.10 | **3.92E-02** | 0.33 | 9.04E-02 | 1.41 | **3.45E-03** |
| AT3G05660 | At3g05660 | RLP33 | -1.09 | **2.34E-02** | 0.08 | 9.26E-01 | 1.16 | **9.86E-03** |
| AT5G66850 | MAPKKK5 | Mitogen-activated protein kinase kinase kinase 5 | -1.09 | **4.80E-02** | -0.24 | 3.24E-01 | 0.83 | 1.33E-01 |
| AT1G54320 | ALIS3 | ALA-interacting subunit 3 | -1.08 | **1.90E-02** | -0.05 | 9.46E-01 | 1.02 | **2.34E-02** |
| AT2G39660 | BIK1 | Serine/threonine-protein kinase BIK1 | -1.08 | **1.79E-02** | -0.02 | 1.00E+00 | 1.06 | **1.67E-02** |
| AT2G17290 | CPK6 | Calcium-dependent protein kinase 6 | -1.08 | **1.67E-02** | -0.14 | 8.19E-01 | 0.93 | **1.77E-02** |
| AT3G47570 | At3g47570 | Probable LRR receptor-like serine/threonine-protein kinase At3g47570 | -1.07 | **2.26E-03** | -0.27 | 4.81E-01 | 0.79 | **3.16E-02** |
| AT5G05190 | EDR4 | Protein ENHANCED DISEASE RESISTANCE 4 | -1.07 | **2.48E-02** | -0.16 | 7.77E-01 | 0.90 | **4.07E-02** |
| AT1G64280 | NPR1 | Regulatory protein NPR1 | -1.06 | **5.56E-04** | -0.42 | **1.15E-02** | 0.62 | 6.03E-02 |
| AT5G16880 | At5g16880 | Target of Myb protein 1 | -1.05 | **6.42E-03** | -0.17 | 5.17E-01 | 0.87 | **2.28E-02** |
| AT2G17220 | PIX13 | Probable serine/threonine-protein kinase PIX13 | -1.05 | **1.09E-02** | -0.34 | 8.25E-02 | 0.70 | 1.08E-01 |
| AT1G03400 | At1g03400 | 2-oxoglutarate | -1.04 | **1.33E-04** | -0.40 | **4.23E-02** | 0.62 | **2.48E-02** |
| AT2G02800 | PBL3 | Probable serine/threonine-protein kinase PBL3 | -1.04 | **5.65E-03** | -0.36 | 1.40E-01 | 0.66 | 9.12E-02 |
| AT4G36210 | At4g36210 | Transmembrane/coiled-coil protein | -1.04 | **2.63E-03** | -0.01 | 1.00E+00 | 1.01 | **2.63E-03** |
| AT2G15042 | RLP18 | Receptor-like protein 18 | -1.03 | **1.04E-02** | -0.25 | 3.66E-01 | 0.77 | **4.27E-02** |
| AT4G17615 | CBL1 | Calcineurin B-like protein | -1.01 | **1.67E-02** | -0.12 | 9.32E-01 | 0.88 | **5.65E-03** |
| AT4G03820 | At4g03820 | Transmembrane protein, putative | -1.01 | **4.01E-02** | -0.14 | 8.22E-01 | 0.85 | 6.75E-02 |
| AT2G23450 | WAKL14 | Wall-associated receptor kinase-like 14 | -1.01 | **2.70E-02** | -0.17 | 7.21E-01 | 0.82 | **4.06E-02** |
| AT4G23650 | CPK3 | Calcium-dependent protein kinase 3 | -1.01 | **1.37E-02** | -0.11 | 8.15E-01 | 0.88 | **1.98E-02** |
| AT3G42790 | AL3 | PHD finger protein ALFIN-LIKE 3 | -1.00 | **3.14E-02** | -0.33 | 3.00E-01 | 0.65 | 1.63E-01 |
| AT1G51270 | At1g51270 | Vesicle-associated protein 1-4 | -1.00 | **1.27E-02** | 0.49 | 2.56E-01 | 1.48 | **3.87E-04** |
| AT1G56540 | At1g56540 | Disease resistance protein | -1.00 | **2.99E-03** | -0.11 | 8.68E-01 | 0.87 | **9.79E-03** |
| AT4G39820 | TRAPPC12 | At4g39820 | -0.99 | **2.09E-02** | -0.12 | 8.77E-01 | 0.86 | 6.45E-02 |
| AT3G24180 | At3g24180 | Non-lysosomal glucosylceramidase | -0.99 | **7.48E-04** | -0.25 | 2.35E-01 | 0.73 | **1.55E-02** |
| AT3G47820 | PUB39 | U-box domain-containing protein 39 | -0.98 | **1.63E-02** | 0.01 | 1.00E+00 | 0.97 | **1.26E-02** |
| AT4G34480 | At4g34480 | glucan endo-1,3-beta-D-glucosidase | -0.97 | **2.71E-02** | -0.21 | 4.06E-01 | 0.75 | 8.09E-02 |
| AT1G27100 | At1g27100 | T7N9.16 | -0.95 | **3.05E-03** | -0.01 | 1.00E+00 | 0.92 | **5.46E-04** |
| AT5G49760 | HPCA1 | Leucine-rich repeat receptor protein kinase HPCA1 | -0.94 | **1.46E-02** | -0.34 | **3.71E-02** | 0.59 | 1.37E-01 |
| AT5G58220 | TTL | Uric acid degradation bifunctional protein TTL | -0.94 | **1.98E-02** | -0.13 | 8.72E-01 | 0.80 | 5.72E-02 |
| AT5G66210 | At5g66210 | AT5G66210 protein | -0.94 | **2.51E-02** | 0.19 | 5.89E-01 | 1.11 | **1.57E-03** |
| AT4G35310 | CPK5 | Calcium-dependent protein kinase 5 | -0.94 | **2.70E-02** | -0.30 | 9.04E-02 | 0.62 | 1.47E-01 |
| AT1G66920 | At1g66920 | Protein kinase superfamily protein | -0.93 | **4.51E-03** | 0.87 | **1.18E-04** | 1.79 | **5.29E-09** |
| AT5G66070 | ATARRE | RING/U-box superfamily protein | -0.93 | **2.40E-02** | 0.42 | 4.97E-01 | 1.34 | **2.49E-04** |
| AT1G63860 | At1g63860 | Uncharacterized protein | -0.93 | **1.38E-02** | -0.16 | 6.90E-01 | 0.76 | **3.12E-02** |
| AT3G20410 | CPK9 | Calcium-dependent protein kinase 9 | -0.93 | **4.40E-02** | 0.22 | 6.75E-01 | 1.13 | **4.79E-03** |
| AT5G42140 | MJC20.25 | Regulator of chromosome condensation | -0.92 | **1.72E-02** | -0.21 | 6.97E-01 | 0.70 | **4.61E-02** |
| AT1G25390 | LRK10L-1.1 | LEAF RUST 10 DISEASE-RESISTANCE LOCUS RECEPTOR-LIKE PROTEIN KINASE-like 1.1 | -0.92 | **1.04E-02** | -0.13 | 7.29E-01 | 0.77 | **2.99E-02** |
| AT5G20910 | AIP2 | E3 ubiquitin-protein ligase AIP2 | -0.92 | **1.57E-02** | 0.08 | 9.51E-01 | 0.97 | **7.04E-03** |
| AT1G76700 | ATJ10 | Chaperone protein dnaJ 10 | -0.90 | **2.66E-02** | -0.14 | 8.26E-01 | 0.75 | 7.03E-02 |
| AT1G56510 | ADR2 | Disease resistance protein ADR2 | -0.89 | **1.51E-02** | -0.20 | 6.02E-01 | 0.68 | **1.63E-02** |
| AT2G21500 | At2g21500 | Uncharacterized protein At2g21500 | -0.89 | **3.95E-02** | 0.23 | 5.49E-01 | 1.10 | **2.04E-03** |
| AT5G58350 | WNK4 | Probable serine/threonine-protein kinase WNK4 | -0.88 | **3.99E-02** | 0.11 | 8.57E-01 | 0.97 | **1.28E-02** |
| AT4G13810 | RLP47 | Receptor like protein | -0.88 | **2.77E-03** | -0.49 | **3.22E-03** | 0.37 | 2.66E-01 |
| AT5G43100 | MMG4.12 | Eukaryotic aspartyl protease family protein | -0.87 | **3.53E-02** | -0.04 | 9.55E-01 | 0.82 | **3.44E-02** |
| AT1G59620 | CW9 | Disease resistance protein | -0.87 | **1.43E-02** | -0.13 | 8.40E-01 | 0.73 | **1.65E-02** |
| AT2G40600 | At2g40600 | Expressed protein | -0.86 | **2.64E-02** | -0.45 | 8.15E-02 | 0.41 | 3.05E-01 |
| AT3G51920 | CML9 | Calmodulin-like protein 9 | -0.86 | **3.41E-03** | -0.28 | 3.14E-01 | 0.56 | 7.41E-02 |
| AT5G46490 | At5g46490 | Disease resistance protein | -0.86 | **1.85E-02** | -0.08 | 8.72E-01 | 0.76 | **1.84E-02** |
| AT2G24360 | STY13 | Serine/threonine-protein kinase STY13 | -0.86 | **3.73E-02** | -0.07 | 8.77E-01 | 0.78 | **4.97E-02** |
| AT4G03080 | BSL1 | Serine/threonine-protein phosphatase BSL1 | -0.85 | **5.47E-03** | -0.25 | 2.58E-01 | 0.59 | 5.69E-02 |
| AT5G46260 | MPL12.4 | ADP-ribosyl cyclase/cyclic ADP-ribose hydrolase | -0.85 | **2.65E-02** | -0.05 | 9.65E-01 | 0.78 | **1.65E-02** |
| AT5G23520 | At5g23520 | Smr domain-containing protein | -0.83 | **1.81E-02** | -0.60 | **1.30E-02** | 0.22 | 6.10E-01 |
| AT3G17410 | CARK1 | Receptor-like cytoplasmic kinase 1 | -0.83 | **2.73E-02** | -0.01 | 1.00E+00 | 0.81 | **1.80E-02** |
| AT2G40830 | RHC1A | Probable E3 ubiquitin-protein ligase RHC1A | -0.83 | **1.46E-02** | -0.44 | **4.33E-02** | 0.38 | 3.60E-01 |
| AT1G28380 | NSL1 | MACPF domain-containing protein NSL1 | -0.82 | **1.53E-02** | -0.14 | 6.86E-01 | 0.67 | 5.69E-02 |
| AT1G56520 | F13N6.13 | Disease resistance protein | -0.82 | **2.76E-02** | -0.17 | 6.21E-01 | 0.64 | 7.77E-02 |
| AT5G66675 | At5g66675 | Uncharacterized protein | -0.82 | **4.58E-02** | -0.03 | 9.89E-01 | 0.77 | **1.61E-02** |
| AT4G34460 | At4g34460 | At4g34460 | -0.81 | **2.86E-02** | -0.02 | 1.00E+00 | 0.77 | 5.35E-02 |
| AT5G61530 | At5g61530 | Small G protein family protein / RhoGAP family protein | -0.81 | **1.90E-02** | 0.13 | 7.73E-01 | 0.93 | **7.46E-03** |
| AT3G46510 | PUB13 | U-box domain-containing protein 13 | -0.80 | **4.18E-02** | 0.11 | 7.25E-01 | 0.90 | **9.74E-03** |
| AT1G05630 | 5PTASE13 | Endonuclease/exonuclease/phosphatase family protein | -0.80 | **2.72E-02** | -0.63 | **2.50E-04** | 0.16 | 7.18E-01 |
| AT3G15430 | At3g15430 | Regulator of chromosome condensation | -0.79 | **4.33E-02** | -0.14 | 7.87E-01 | 0.64 | 7.75E-02 |

| | | | | | | | | |
|---|---|---|---|---|---|---|---|---|
| AT2G40940 | ERS1 | Ethylene response sensor 1 | -0.79 | **3.50E-02** | -0.20 | 5.31E-01 | 0.57 | 1.23E-01 |
| AT1G12990 | F3F19.2 | F3F19.2 protein | -0.78 | **1.23E-02** | -0.15 | 6.41E-01 | 0.61 | **4.12E-02** |
| AT1G75020 | LPAT4 | Probable 1-acyl-sn-glycerol-3-phosphate acyltransferase 4 | -0.76 | **3.64E-02** | -0.21 | 5.96E-01 | 0.53 | 1.66E-01 |
| AT3G18370 | ATSYTF | AT3g18370/MYF24_8 | -0.75 | **2.13E-02** | -0.11 | 7.82E-01 | 0.62 | 5.68E-02 |
| AT1G78290 | SRK2C | Serine/threonine-protein kinase SRK2C | -0.74 | **4.40E-02** | -0.20 | 6.31E-01 | 0.52 | 1.69E-01 |
| AT3G59080 | F17J16_130 | AT3g59080/F17J16_130 | -0.72 | **4.10E-02** | -0.17 | 7.12E-01 | 0.54 | 1.31E-01 |
| AT2G44500 | At2g44500 | O-fucosyltransferase family protein | -0.72 | **4.86E-02** | 0.47 | **4.28E-02** | 1.18 | **5.23E-07** |
| AT1G79380 | RGLG4 | E3 ubiquitin-protein ligase RGLG4 | -0.70 | **3.27E-02** | -0.33 | **4.09E-02** | 0.36 | 2.98E-01 |
| AT2G39840 | TOPP4 | Serine/threonine-protein phosphatase PP1 isozyme 4 | -0.69 | **2.64E-02** | -0.02 | 1.00E+00 | 0.66 | **2.41E-02** |
| AT1G74170 | RLP13 | Receptor like protein 13 | -0.68 | **4.01E-02** | -0.46 | **3.64E-02** | 0.21 | 6.28E-01 |
| AT1G60490 | VPS34 | Phosphatidylinositol 3-kinase VPS34 | -0.65 | **3.78E-02** | -0.21 | 9.99E-01 | 0.44 | 3.88E-01 |
| AT4G12010 | DSC1 | Disease resistance-like protein DSC1 | -0.64 | **4.80E-02** | -0.43 | **3.61E-02** | 0.20 | 5.87E-01 |
| **Immune/ defense response/ response to biotic stress** | | | | | | | | |
| AT2G14620 | XTH10 | Xyloglucan endotransglucosylase/hydrolase | -11.47 | **4.43E-11** | | | 3.76 | **4.06E-04** |
| AT3G53150 | UGT73D1 | UDP-glycosyltransferase 73D1 | -10.40 | **4.38E-07** | | | 5.47 | **4.55E-04** |
| AT2G13810 | ALD1 | Aminotransferase ALD1, chloroplastic | -10.14 | **7.17E-27** | -5.85 | **7.38E-13** | 4.21 | **1.73E-10** |
| AT3G11340 | UGT76B1 | UDP-glycosyltransferase 76B1 | -9.49 | **5.60E-28** | -5.49 | **1.60E-13** | 3.88 | **1.52E-12** |
| AT2G30770 | CYP71A13 | Indoleacetaldoxime dehydratase | -9.41 | **6.22E-07** | | | 6.21 | **8.93E-05** |
| AT1G19250 | FMO1 | Probable flavin-containing monooxygenase 1 | -9.12 | **1.59E-13** | | | 5.55 | **1.14E-08** |
| AT2G35980 | NHL10 | NDR1/HIN1-like protein 10 | -9.03 | **2.96E-07** | | | 8.29 | **7.61E-06** |
| AT2G45220 | PME17 | Pectinesterase inhibitor 17] | -7.99 | **1.60E-06** | | | 5.82 | **3.48E-05** |
| AT3G26830 | At3g26830 | PAD3 | -7.87 | **2.54E-08** | | | 5.64 | **3.51E-07** |
| AT4G10500 | DLO1 | Protein DMR6-LIKE OXYGENASE 1 | -7.64 | **1.48E-09** | -3.31 | **7.15E-31** | 4.36 | **7.52E-05** |
| AT1G65483 | At1g65483 | KH type-2 domain-containing protein | -7.38 | **8.85E-07** | | | 4.53 | **2.89E-04** |
| AT2G29460 | GSTU4 | Glutathione S-transferase U4 | -7.04 | **5.64E-07** | | | 4.14 | **9.65E-04** |
| AT3G22600 | LTPG5 | Non-specific lipid transfer protein GPI-anchored 5 | -6.65 | **8.81E-07** | -2.30 | **3.17E-02** | 4.46 | **1.94E-04** |
| AT1G14880 | PCR1 | Protein PLANT CADMIUM RESISTANCE 1 | -6.52 | **2.19E-11** | -3.04 | **1.00E-28** | 3.48 | **1.58E-05** |
| AT1G75040 | At1g75040 | Pathogenesis-related protein 5 | -6.38 | **8.82E-05** | -2.25 | **6.13E-10** | 4.12 | **5.77E-03** |
| AT1G65610 | KOR2 | Endoglucanase 7 | -6.22 | **2.97E-04** | | | 6.45 | **5.33E-05** |
| AT1G15520 | ABCG40 | Pleiotropic drug resistance 12 | -6.14 | **5.57E-09** | -0.85 | 1.81E-01 | 5.27 | **1.66E-07** |
| AT2G25297 | At2g25297 | Transmembrane protein | -6.12 | **2.03E-06** | | | 3.73 | **1.08E-03** |
| AT3G13950 | At3g13950 | At3g13950 | -6.10 | **2.42E-04** | | | 4.75 | **1.48E-04** |
| AT3G48640 | T8P19.150 | Transmembrane protein | -5.84 | **4.76E-07** | -3.45 | **2.30E-03** | 2.59 | **4.68E-03** |
| AT4G23150 | CRK7 | Cysteine-rich receptor-like protein kinase 7 | -5.83 | **2.67E-07** | -1.98 | **8.62E-04** | 3.86 | **7.25E-06** |
| AT4G00700 | MCTP9 | Multiple C2 domain and transmembrane region protein 9 | -5.77 | **1.24E-09** | -2.05 | **5.75E-11** | 3.69 | **3.52E-05** |
| AT1G21240 | WAK3 | Wall-associated receptor kinase-like | -5.76 | **1.91E-03** | -1.60 | 1.87E-01 | 4.15 | **1.10E-05** |
| AT3G57260 | PR2 | glucan endo-1,3-beta-D-glucosidase | -5.70 | **3.10E-04** | -1.47 | **8.73E-03** | 4.22 | **2.09E-03** |
| AT1G51890 | At1g51890 | Leucine-rich repeat protein kinase family protein | -5.58 | **3.51E-08** | -0.74 | 5.78E-01 | 4.82 | **3.10E-06** |
| AT2G18660 | EGC2 | EG45-like domain containing protein 2 | -5.50 | **1.93E-14** | -3.29 | **3.88E-18** | 2.17 | **1.10E-03** |
| AT5G26690 | HIPP02 | Heavy metal-associated isoprenylated plant protein 2 | -5.43 | **3.60E-05** | -1.39 | 5.16E-01 | 4.16 | **3.61E-06** |
| AT2G19190 | SIRK | Senescence-induced receptor-like serine/threonine-protein kinase | -5.42 | **1.38E-09** | -0.95 | 4.54E-01 | 4.47 | **2.13E-07** |
| AT5G52760 | At5g52760 | Copper transport protein family | -5.39 | **7.52E-08** | -1.46 | **3.09E-04** | 3.92 | **4.39E-05** |
| AT1G57650 | At1g57650 | ATP binding protein | -5.37 | **1.25E-05** | | | 4.20 | **3.25E-05** |
| AT4G14390 | DL3235W | Ankyrin repeat family protein | -5.37 | **1.87E-06** | -3.24 | **1.51E-05** | 2.11 | **1.18E-02** |
| AT5G47850 | CCR4 | Serine/threonine-protein kinase-like protein CCR4 | -5.34 | **4.55E-09** | | | 4.59 | **1.37E-07** |
| AT5G22570 | WRKY38 | Probable WRKY transcription factor 38 | -5.32 | **9.99E-09** | -2.27 | **3.15E-07** | 3.01 | **4.08E-04** |
| AT4G23310 | CRK23 | Cysteine-rich RLK | -5.18 | **1.60E-06** | -1.81 | **4.14E-03** | 3.35 | **1.72E-05** |
| AT5G02490 | HSP70-2 | Heat shock 70 kDa protein 2 | -4.92 | **1.57E-06** | -0.73 | 1.64E-01 | 4.19 | **2.03E-05** |
| AT2G47130 | SDR3a | Short-chain dehydrogenase reductase 3a | -4.73 | **8.53E-11** | -1.95 | **1.25E-07** | 2.77 | **7.24E-05** |
| AT4G23140 | CRK6 | Cysteine-rich receptor-like protein kinase 6 | -4.57 | **4.95E-09** | -1.59 | **4.71E-38** | 2.96 | **1.23E-04** |
| AT1G51860 | At1g51860 | Leucine-rich repeat protein kinase family protein | -4.54 | **1.27E-04** | | | 4.20 | **5.76E-03** |
| AT5G42830 | MBD2.2 | HXXXD-type acyl-transferase family protein | -4.51 | **8.52E-10** | -2.17 | **9.60E-07** | 2.34 | **1.92E-04** |
| AT2G46400 | WRKY46 | Probable WRKY transcription factor 46 | -4.50 | **1.39E-08** | -0.87 | 1.85E-01 | 3.61 | **3.45E-08** |
| AT2G32680 | RLP23 | Receptor like protein 23 | -4.45 | **1.48E-09** | -1.71 | **4.36E-14** | 2.73 | **3.87E-05** |
| AT1G73805 | SARD1 | Protein SAR DEFICIENT 1 | -4.35 | **1.76E-11** | -1.15 | **3.47E-03** | 3.20 | **3.66E-09** |
| AT3G29000 | CML45 | Probable calcium-binding protein CML45 | -4.31 | **2.03E-05** | | | 3.43 | **3.68E-04** |
| AT2G14560 | LURP1 | LURP-one-like protein | -4.24 | **3.39E-15** | -1.84 | **1.18E-33** | 2.39 | **2.19E-06** |
| AT5G53110 | At5g53110 | RING-type E3 ubiquitin transferase | -4.22 | **1.24E-05** | -0.43 | 8.55E-01 | 3.78 | **7.88E-05** |
| AT5G13320 | PBS3 | Auxin-responsive GH3 family protein | -4.18 | **1.81E-07** | -0.45 | 2.35E-01 | 3.72 | **3.52E-06** |
| AT4G39830 | AO | L-ascorbate oxidase | -4.15 | **5.86E-06** | -1.48 | **3.42E-05** | 2.65 | **4.22E-03** |
| AT5G10760 | AED1 | Aspartyl protease AED1 | -4.14 | **4.27E-08** | -1.34 | **8.84E-11** | 2.79 | **7.52E-05** |
| AT2G18690 | At2g18690 | At2g18690/MSF3.7 | -4.14 | **7.94E-06** | -0.31 | 2.54E-01 | 3.82 | **4.09E-05** |
| AT1G32960 | SBT3.3 | Subtilisin-like protease SBT3.3 | -4.11 | **4.14E-05** | | | 4.97 | **3.52E-06** |
| AT1G08050 | At1g08050 | Zinc finger family protein | -4.11 | **3.92E-07** | | | 2.77 | **2.76E-03** |
| AT5G52740 | At5g52740 | Copper transport protein family | -4.08 | **4.37E-10** | | | 2.83 | **5.09E-06** |
| AT5G17760 | At5g17760 | AAA-ATPase At5g17760 | -4.04 | **2.35E-06** | -1.00 | 6.63E-02 | 3.03 | **5.98E-04** |
| AT1G51800 | IOS1 | LRR receptor-like serine/threonine-protein kinase IOS1 | -4.03 | **1.30E-11** | -0.86 | 5.91E-02 | 3.14 | **3.14E-08** |
| AT2G27660 | At2g27660 | Cysteine/Histidine-rich C1 domain family protein | -3.96 | **3.68E-06** | -0.77 | 5.39E-01 | 3.20 | **4.83E-07** |
| AT5G52810 | At5g52810 | AT5G52810 protein | -3.92 | **1.76E-11** | -1.64 | **9.51E-14** | 2.28 | **5.26E-05** |
| AT1G02230 | NAC004 | NAC domain-containing protein 4 | -3.89 | **9.58E-04** | -2.93 | **2.15E-02** | 1.28 | 1.80E-01 |
| AT1G03850 | GRXS13 | Glutaredoxin family protein | -3.86 | **3.58E-06** | -0.77 | 4.43E-01 | 3.09 | **1.16E-05** |
| AT5G46050 | NPF52 | Peptide transporter 3 | -3.84 | **8.80E-06** | -0.71 | 5.44E-01 | 3.10 | **1.88E-05** |
| AT1G24145 | At1g24145 | At1g24145 | -3.83 | **2.17E-05** | -0.78 | 1.96E-01 | 3.05 | **2.90E-04** |
| AT1G57560 | MYB50 | At1g57560 | -3.73 | **4.85E-04** | -1.06 | 5.58E-01 | 2.75 | **9.92E-04** |
| AT4G03450 | At4g03450 | Ankyrin repeat family protein | -3.72 | **1.56E-09** | -0.95 | **2.41E-02** | 2.77 | **4.72E-07** |
| AT1G26420 | FOX5 | Berberine bridge enzyme-like 7 | -3.72 | **2.15E-04** | 0.59 | 7.37E-01 | 4.27 | **1.84E-05** |
| AT3G51330 | At3g51330 | Eukaryotic aspartyl protease family protein | -3.69 | **4.90E-07** | -0.90 | **2.01E-02** | 2.78 | **1.09E-04** |
| AT2G44400 | At2g44400 | Cysteine/Histidine-rich C1 domain family protein | -3.69 | **5.13E-08** | -1.78 | 9.94E-02 | 1.99 | **1.80E-03** |
| AT5G48657 | At5g48657 | Defense protein-like protein | -3.66 | **1.38E-09** | -0.63 | 2.52E-01 | 3.01 | **3.51E-07** |
| AT4G21850 | MSRB9 | Peptide methionine sulfoxide reductase B9-S-oxide reductase | -3.54 | **1.12E-04** | 0.19 | 9.59E-01 | 3.68 | **6.31E-04** |
| AT2G32140 | At2g32140 | Transmembrane receptor | -3.54 | **1.55E-08** | -0.37 | 9.02E-01 | 3.18 | **3.88E-10** |
| AT1G56120 | At1g56120 | Leucine-rich repeat transmembrane protein kinase | -3.49 | **9.74E-12** | -0.89 | **1.45E-03** | 2.59 | **7.23E-07** |
| AT2G04070 | DTX4 | Protein DETOXIFICATION 4 | -3.49 | **9.76E-04** | 0.05 | 9.93E-01 | 3.44 | **4.34E-04** |
| AT4G18253 | At4g18253 | Receptor Serine/Threonine kinase-like protein | -3.48 | **1.35E-06** | -1.26 | **2.39E-07** | 2.22 | **2.98E-03** |
| AT1G21250 | WAK1 | Wall-associated receptor kinase 1 | -3.47 | **4.15E-07** | -1.30 | **1.32E-08** | 2.16 | **5.98E-04** |
| AT3G56400 | WRKY70 | Probable WRKY transcription factor 70 | -3.46 | **1.24E-09** | -0.99 | **1.49E-07** | 2.46 | **4.71E-06** |
| AT5G24530 | DMR6 | Protein DOWNY MILDEW RESISTANCE 6 | -3.42 | **3.50E-12** | -1.63 | **6.71E-09** | 1.78 | **3.45E-08** |
| AT5G03350 | LLP | Lectin-like protein | -3.40 | **1.62E-06** | -2.08 | **1.26E-13** | 1.30 | **4.94E-02** |
| AT5G62770 | At5g62770 | Uncharacterized protein | -3.36 | **1.15E-04** | -0.05 | 9.99E-01 | 3.30 | **5.94E-05** |

| | | | | | | | | |
|---|---|---|---|---|---|---|---|---|
| AT5G43420 | ATL16 | RING-H2 finger protein ATL16 | -3.31 | 1.06E-03 | | | 2.21 | 3.54E-02 |
| AT5G08240 | At5g08240 | Transmembrane protein | -3.30 | 1.57E-05 | -0.07 | 9.94E-01 | 3.21 | 8.64E-06 |
| AT5G67340 | PUB2 | U-box domain-containing protein 2 | -3.28 | 4.00E-06 | -1.03 | 1.41E-05 | 2.24 | 1.99E-03 |
| AT2G20142 | At2g20142 | Toll-Interleukin-Resistance | -3.28 | 7.10E-06 | -0.57 | 4.92E-01 | 2.71 | 6.02E-05 |
| AT3G22231 | PCC1 | Cysteine-rich and transmembrane domain-containing protein PCC1 | -3.27 | 2.36E-07 | -2.23 | 3.63E-15 | 1.04 | 7.48E-02 |
| AT5G25910 | RLP52 | Receptor-like protein 52 | -3.27 | 1.55E-05 | -0.91 | 1.92E-01 | 2.33 | 2.87E-03 |
| AT4G39030 | DTX47 | Protein DETOXIFICATION 47, chloroplastic | -3.26 | 8.01E-09 | -0.12 | 9.51E-01 | 3.13 | 1.96E-08 |
| AT2G31880 | SOBIR1 | Leucine-rich repeat receptor-like serine/threonine/tyrosine-protein kinase SOBIR1 | -3.24 | 4.82E-06 | -0.77 | 5.58E-05 | 2.46 | 4.21E-04 |
| AT5G59670 | At5g59670 | Receptor-like protein kinase At5g59670 | -3.24 | 3.10E-07 | -1.66 | 3.88E-05 | 1.56 | 9.77E-05 |
| AT4G26120 | At4g26120 | Ankyrin repeat family protein / BTB/POZ domain-containing protein | -3.22 | 9.22E-04 | -0.77 | 7.78E-01 | 2.53 | 1.96E-05 |
| AT2G40750 | WRKY54 | Probable WRKY transcription factor 54 | -3.22 | 8.64E-11 | -1.28 | 4.12E-07 | 1.94 | 4.59E-06 |
| AT1G72240 | At1g72240 | Uncharacterized protein | -3.20 | 3.98E-04 | -1.33 | 4.46E-01 | 1.91 | 3.55E-02 |
| AT4G15610 | CASPL1D1 | CASP-like protein | -3.16 | 2.41E-02 | 0.18 | 9.60E-01 | 3.35 | 1.35E-02 |
| AT3G04220 | At3g04220 | Disease resistance protein family | -3.15 | 1.42E-02 | 0.04 | 9.93E-01 | 3.19 | 5.08E-03 |
| AT5G27420 | ATL31 | E3 ubiquitin-protein ligase ATL31 | -3.13 | 1.67E-04 | 0.39 | 8.24E-01 | 3.51 | 1.85E-05 |
| AT1G24140 | 3MMP | Metalloendoproteinase 3-MMP | -3.13 | 4.80E-06 | -0.73 | 2.48E-01 | 2.40 | 1.96E-05 |
| AT4G14370 | DL3225C | Disease resistance protein | -3.12 | 3.68E-06 | -1.04 | 7.15E-02 | 2.08 | 7.90E-04 |
| AT5G43910 | At5g43910 | PfkB-like carbohydrate kinase family protein | -3.11 | 4.91E-06 | -0.86 | 5.41E-02 | 2.25 | 1.44E-03 |
| AT1G74710 | ICS1 | Isochorismate synthase 1, chloroplastic | -3.08 | 9.50E-06 | -0.29 | 3.71E-01 | 2.78 | 9.30E-05 |
| AT3G56710 | SIB1 | Sigma factor binding protein 1, chloroplastic | -3.05 | 1.38E-08 | -0.72 | 1.67E-02 | 2.31 | 4.92E-06 |
| AT5G26920 | CBP60G | Cam-binding protein 60-like G | -3.04 | 1.37E-06 | -0.38 | 2.69E-01 | 2.65 | 2.28E-05 |
| AT1G08450 | At1g08450 | Calreticulin like protein | -3.04 | 6.39E-07 | -0.74 | 1.75E-05 | 2.29 | 1.48E-04 |
| AT1G63720 | F24D7.9 | Uncharacterized protein F24D7.9 | -3.02 | 2.40E-06 | -0.60 | 2.66E-01 | 2.40 | 3.02E-04 |
| AT1G80840 | WRKY40 | Probable WRKY transcription factor 40 | -3.02 | 1.59E-06 | 0.97 | 3.70E-01 | 3.97 | 3.00E-20 |
| AT1G05340 | CYSTM1 | Protein CYSTEINE-RICH TRANSMEMBRANE MODULE 1 | -3.01 | 1.58E-02 | 0.04 | 9.93E-01 | 3.08 | 2.42E-02 |
| AT5G59680 | At5g59680 | Probable LRR receptor-like serine/threonine-protein kinase At5g59680 | -3.01 | 6.54E-09 | -1.19 | 5.28E-03 | 1.82 | 1.94E-06 |
| AT3G51440 | SSL6 | Protein STRICTOSIDINE SYNTHASE-LIKE 6 | -3.01 | 5.08E-04 | 0.16 | 9.02E-01 | 3.17 | 3.77E-04 |
| AT5G57480 | At5g57480 | AAA-ATPase At5g57480 | -3.00 | 1.07E-03 | -1.19 | 3.89E-01 | 1.86 | 6.56E-02 |
| AT3G63380 | ACA12 | Calcium-transporting ATPase 12, plasma membrane-type | -2.99 | 3.25E-03 | -0.51 | 3.53E-01 | 2.48 | 1.61E-02 |
| AT2G44290 | LTPG13 | Non-specific lipid transfer protein GPI-anchored 13 | -2.99 | 4.29E-04 | -0.78 | 3.39E-06 | 2.20 | 1.16E-02 |
| AT1G61420 | At1g61420 | S-locus lectin protein kinase family protein | -2.98 | 6.74E-05 | -0.38 | 8.74E-01 | 2.55 | 7.82E-04 |
| AT2G22470 | AGP2 | Classical arabinogalactan protein 2 | -2.97 | 5.16E-03 | -0.36 | 8.60E-01 | 2.61 | 8.48E-03 |
| AT5G55460 | ATLTP45 | At5g55460 | -2.97 | 1.70E-04 | -0.45 | 8.22E-01 | 2.47 | 4.33E-04 |
| AT5G54610 | BAD1 | Ankyrin repeat-containing protein BDA1 | -2.95 | 7.97E-11 | -1.15 | 2.61E-08 | 1.79 | 8.64E-06 |
| AT3G50480 | HR4 | Homolog of RPW8 4 | -2.93 | 3.50E-12 | -0.79 | 2.34E-08 | 2.13 | 4.42E-07 |
| AT2G40140 | At2g40140 | Zinc finger CCCH domain-containing protein 29 | -2.88 | 2.24E-07 | -0.31 | 7.30E-01 | 2.55 | 2.28E-07 |
| AT3G62780 | F26K9_210 | At3g62780 | -2.87 | 9.47E-06 | -1.28 | 6.86E-02 | 1.57 | 1.22E-02 |
| AT5G18270 | ANAC087 | NAC domain containing protein 87 | -2.87 | 9.74E-03 | | | 4.18 | 3.71E-04 |
| AT5G26170 | At5g26170 | WRKY50 | -2.87 | 2.02E-05 | -1.16 | 5.59E-02 | 1.68 | 7.01E-03 |
| AT1G67920 | At1g67920 | Uncharacterized protein | -2.86 | 1.94E-02 | -0.32 | 9.27E-01 | 2.45 | 3.30E-02 |
| AT4G01870 | At4g01870 | TolB protein-like protein | -2.83 | 5.84E-05 | -0.26 | 6.90E-01 | 2.53 | 3.76E-04 |
| AT2G33580 | LYK5 | Protein LYK5 | -2.82 | 4.76E-07 | -0.68 | 2.29E-05 | 2.13 | 1.71E-04 |
| AT3G26440 | At3g26440 | Transmembrane protein, putative | -2.81 | 3.84E-03 | 0.28 | 8.39E-01 | 3.08 | 1.71E-04 |
| AT2G26560 | PLP2 | Patatin-like protein 2 | -2.78 | 2.81E-05 | 0.47 | 5.40E-01 | 3.23 | 1.12E-12 |
| AT3G11402 | At3g11402 | Cysteine/Histidine-rich C1 domain family protein | -2.75 | 2.79E-07 | -1.15 | 8.54E-05 | 1.58 | 6.35E-03 |
| AT3G12830 | SAUR72 | Auxin-responsive protein SAUR72 | -2.72 | 1.82E-04 | -0.75 | 2.78E-01 | 1.96 | 1.10E-02 |
| AT1G66465 | At1g66465 | Transmembrane protein | -2.72 | 8.10E-04 | -0.81 | 2.31E-01 | 1.88 | 3.02E-02 |
| AT4G14400 | ACD6 | Protein ACCELERATED CELL DEATH 6 | -2.72 | 5.37E-10 | -1.32 | 7.34E-19 | 1.38 | 1.16E-03 |
| AT5G01550 | LECRK-VI3 | Lectin receptor kinase a4.1 | -2.70 | 1.79E-02 | 1.22 | 4.70E-01 | 3.93 | 1.93E-04 |
| AT3G10930 | IDL7 | Uncharacterized protein At3g10930 | -2.70 | 4.30E-04 | | | 2.30 | 4.10E-03 |
| AT1G68390 | At1g68390 | Core-2/I-branching beta-1,6-N-acetylglucosaminyltransferase family protein | -2.69 | 2.01E-03 | -0.95 | 5.64E-01 | 1.72 | 4.54E-02 |
| AT3G25020 | RLP42 | Receptor-like protein 42 | -2.67 | 1.59E-06 | -0.50 | 4.78E-03 | 2.15 | 1.08E-04 |
| AT5G54860 | At5g54860 | Probable folate-biopterin transporter 4 | -2.67 | 6.04E-06 | -0.92 | 6.10E-04 | 1.74 | 6.15E-03 |
| AT5G64310 | AGP1 | Classical arabinogalactan protein 1 | -2.66 | 7.65E-09 | -0.08 | 9.74E-01 | 2.57 | 9.98E-09 |
| AT5G63130 | At5g63130 | hypothetical protein | -2.64 | 7.04E-06 | -0.74 | 6.15E-01 | 1.85 | 2.04E-04 |
| AT4G21903 | At4g21903 | Protein DETOXIFICATION | -2.63 | 1.28E-03 | 0.09 | 9.80E-01 | 2.71 | 3.77E-04 |
| AT4G23160 | At4g23160 | Uncharacterized protein | -2.62 | 9.94E-07 | -0.19 | 9.14E-01 | 2.42 | 1.01E-05 |
| AT1G76970 | At1g76970 | Target of Myb protein 1 | -2.61 | 6.05E-06 | -0.46 | 3.01E-01 | 2.14 | 3.73E-04 |
| AT3G01080 | WRKY58 | WRKY DNA-binding protein 58 | -2.59 | 2.15E-07 | -0.74 | 1.38E-01 | 1.83 | 2.87E-04 |
| AT5G46350 | WRKY8 | WRKY transcription factor 8 | -2.59 | 1.38E-02 | -0.45 | 6.90E-01 | 2.15 | 2.59E-02 |
| AT1G74360 | At1g74360 | Probable LRR receptor-like serine/threonine-protein kinase At1g74360 | -2.59 | 4.15E-05 | -0.30 | 5.83E-01 | 2.29 | 3.60E-04 |
| AT3G13790 | ATBFRUCT1 | Glycosyl hydrolases family 32 protein | -2.59 | 1.65E-05 | -1.69 | 6.16E-17 | 0.88 | 2.28E-01 |
| AT3G61280 | T20K12.180 | Uncharacterized protein T20K12.180 | -2.58 | 2.41E-06 | -0.35 | 3.78E-01 | 2.22 | 3.32E-06 |
| AT4G23230 | CRK15 | Cysteine-rich receptor-like protein kinase 15 | -2.57 | 2.06E-05 | -0.91 | 1.48E-02 | 1.66 | 1.51E-03 |
| AT3G52430 | PAD4 | Lipase-like PAD4 | -2.52 | 4.70E-07 | -0.32 | 3.13E-01 | 2.19 | 3.42E-05 |
| AT1G70690 | PDLP5 | Plasmodesmata-located protein 5 | -2.52 | 2.55E-03 | -0.13 | 9.02E-01 | 2.38 | 3.54E-03 |
| AT2G17120 | LYM2 | LysM domain-containing GPI-anchored protein 2 | -2.51 | 2.12E-04 | -0.88 | 2.25E-09 | 1.62 | 2.33E-02 |
| AT5G44568 | PROSCOOP4 | Serine rich endogenous peptide 4 | -2.49 | 1.39E-05 | -0.95 | 1.73E-10 | 1.53 | 1.10E-02 |
| AT1G15010 | T15D22.5 | T15D22.5 protein | -2.48 | 9.75E-05 | | | 2.62 | 1.50E-04 |
| AT1G07000 | EXO70B2 | Exocyst complex component EXO70B2 | -2.47 | 1.38E-09 | -0.55 | 2.01E-01 | 1.91 | 8.78E-06 |
| AT3G06890 | At3g06890 | hypothetical protein | -2.44 | 2.12E-03 | | | 2.79 | 4.41E-04 |
| AT1G72900 | At1g72900 | Similar to part of disease resistance protein | -2.42 | 4.14E-05 | -0.39 | 5.89E-01 | 2.03 | 5.48E-04 |
| AT3G48090 | EDS1 | Protein EDS1 | -2.42 | 4.11E-06 | -0.86 | 4.94E-08 | 1.54 | 5.92E-03 |
| AT3G22400 | LOX5 | Linoleate 9S-lipoxygenase 5 | -2.39 | 6.50E-03 | -0.16 | 8.92E-01 | 2.20 | 9.71E-03 |
| AT5G25930 | T1N24.22 | Kinase family with leucine-rich repeat domain-containing protein | -2.38 | 3.33E-05 | -0.85 | 2.82E-04 | 1.52 | 1.01E-02 |
| AT4G23170 | CRK9 | Putative cysteine-rich receptor-like protein kinase 9 | -2.38 | 9.21E-06 | -0.83 | 5.25E-08 | 1.53 | 5.85E-03 |
| AT2G38860 | YLS5 | Class I glutamine amidotransferase-like superfamily protein | -2.37 | 1.70E-04 | -0.32 | 6.29E-01 | 2.03 | 1.79E-03 |
| AT5G52640 | HSP90-1 | Heat shock protein 90-1 | -2.37 | 5.80E-04 | -0.30 | 6.27E-01 | 2.07 | 1.83E-03 |
| AT2G41640 | At2g41640 | Glycosyltransferase family 61 protein | -2.37 | 2.88E-05 | 0.28 | 8.70E-01 | 2.64 | 8.73E-14 |
| AT1G79680 | WAKL10 | Wall-associated receptor kinase-like 10 | -2.34 | 4.38E-02 | -0.06 | 1.00E+00 | 2.27 | 2.42E-02 |
| AT1G67000 | LRK10L-2.8 | LEAF RUST 10 DISEASE-RESISTANCE LOCUS RECEPTOR-LIKE PROTEIN KINASE-like 2.8 | -2.31 | 1.71E-04 | 0.45 | 6.40E-01 | 2.76 | 1.10E-06 |
| AT5G19240 | At5g19240 | Uncharacterized GPI-anchored protein At5g19240 | -2.31 | 2.11E-05 | -0.99 | 1.25E-08 | 1.30 | 1.90E-02 |
| AT2G38470 | At2g38470 | Truncated WRKY33 protein | -2.29 | 6.15E-07 | -0.38 | 8.51E-02 | 1.89 | 1.03E-05 |
| AT1G24150 | FH4 | Formin-like protein | -2.28 | 4.99E-05 | -0.96 | 1.10E-08 | 1.31 | 2.86E-02 |
| AT2G31865 | PARG2 | poly | -2.28 | 5.21E-05 | -0.18 | 7.75E-01 | 2.10 | 1.79E-04 |
| AT1G67520 | At1g67520 | Receptor-like serine/threonine-protein kinase | -2.28 | 1.25E-02 | -0.51 | 8.16E-01 | 1.78 | 8.70E-02 |
| AT4G11850 | PLDGAMMA | Phospholipase D gamma 1 | -2.27 | 4.51E-06 | -0.62 | 8.07E-03 | 1.65 | 1.04E-03 |
| AT3G45860 | CRK4 | Cysteine-rich receptor-like protein kinase 4 | -2.26 | 3.68E-06 | -0.29 | 5.45E-01 | 1.96 | 4.19E-07 |
| AT4G12720 | NUDT7 | Nudix hydrolase | -2.25 | 7.21E-05 | -0.05 | 9.64E-01 | 2.19 | 5.02E-05 |
| AT1G76980 | At1g76980 | Patatin-like phospholipase domain protein | -2.24 | 9.84E-03 | -0.38 | 7.41E-01 | 1.87 | 1.96E-02 |

| | | | | | | | | |
|---|---|---|---|---|---|---|---|---|
| AT2G37710 | LECRK41 | L-type lectin-domain containing receptor kinase IV.1 | -2.23 | **3.30E-05** | -0.75 | **1.17E-08** | 1.46 | **9.65E-03** |
| AT2G40095 | At2g40095 | Alpha/beta hydrolase related protein | -2.22 | **1.81E-05** | -0.74 | **1.12E-02** | 1.45 | **6.93E-03** |
| AT3G52400 | SYP122 | Syntaxin-122 | -2.20 | **1.21E-05** | 0.11 | 8.78E-01 | 2.30 | **2.79E-06** |
| AT5G39050 | PMAT1 | Phenolic glucoside malonyltransferase 1 | -2.20 | **1.49E-04** | -0.11 | 8.59E-01 | 2.07 | **2.32E-04** |
| AT5G48380 | At5g48380 | Probably inactive leucine-rich repeat receptor-like protein kinase At5g48380 | -2.20 | **1.63E-04** | -0.19 | 5.02E-01 | 2.00 | **8.12E-04** |
| AT3G29034 | At3g29034 | Transmembrane protein | -2.17 | **2.54E-03** | | | 2.86 | **6.89E-04** |
| AT4G28490 | RLK5 | Receptor-like protein kinase 5 | -2.17 | **1.45E-05** | -0.80 | **2.53E-06** | 1.36 | **9.78E-03** |
| AT1G51660 | MKK4 | Mitogen-activated protein kinase kinase 4 | -2.16 | **2.15E-04** | -0.69 | 5.58E-02 | 1.47 | **1.05E-02** |
| AT4G15233 | ABCG42 | ABC-2 and Plant PDR ABC-type transporter family protein | -2.16 | **1.70E-07** | -0.69 | **2.81E-02** | 1.46 | **6.31E-05** |
| AT3G48520 | CYP94B3 | Cytochrome P450 94B3 | -2.13 | **3.08E-02** | 0.31 | 8.74E-01 | 2.43 | **5.77E-04** |
| AT5G06320 | NHL3 | NDR1/HIN1-like protein 3 | -2.12 | **8.22E-05** | -0.58 | **3.54E-03** | 1.52 | **5.05E-03** |
| AT4G31800 | WRKY18 | WRKY DNA-binding protein 18 | -2.09 | **4.48E-03** | 0.89 | 1.02E-01 | 2.97 | **2.14E-07** |
| AT3G45620 | At3g45620 | Transducin/WD40 repeat-like superfamily protein | -2.08 | **3.41E-04** | -0.57 | 1.00E-01 | 1.50 | **8.34E-03** |
| AT1G45145 | TRX5 | Thioredoxin H5 | -2.08 | **1.28E-03** | 0.96 | **2.17E-02** | 3.03 | **5.53E-06** |
| AT1G29240 | At1g29240 | | -2.06 | **2.39E-04** | -0.52 | 7.27E-02 | 1.53 | **9.44E-03** |
| AT5G40780 | LHT1 | Lysine histidine transporter 1 | -2.05 | **2.94E-04** | -0.24 | 2.71E-01 | 1.80 | **1.81E-03** |
| AT4G29780 | F27B13.20 | Uncharacterized protein AT4g29780 | -2.05 | **3.29E-04** | 0.75 | 1.13E-01 | 2.78 | **1.58E-07** |
| AT5G25440 | SZE1 | Protein kinase superfamily protein | -2.05 | **4.20E-08** | -0.97 | **9.60E-07** | 1.06 | **5.19E-03** |
| AT1G53920 | At1g53920 | GLIP5 | -2.05 | **5.95E-04** | -1.67 | **2.01E-02** | 0.36 | 6.45E-01 |
| AT1G61460 | At1g61460 | G-type lectin S-receptor-like serine/threonine-protein kinase At1g61460 | -2.04 | **4.37E-03** | -0.75 | 6.82E-01 | 1.34 | 7.29E-02 |
| AT2G23680 | At2g23680 | Uncharacterized protein | -2.03 | **3.85E-04** | -0.21 | 8.34E-01 | 1.81 | **9.38E-04** |
| AT4G22530 | At4g22530 | | -2.02 | **5.92E-04** | -0.38 | 7.77E-01 | 1.64 | **8.50E-04** |
| AT5G42530 | At5g42530 | | -2.01 | **6.18E-06** | -1.01 | **1.29E-04** | 0.99 | **3.29E-03** |
| AT3G07580 | At3g07580 | At3g07580 | -2.00 | **1.53E-02** | -0.53 | 4.92E-01 | 1.45 | 9.00E-02 |
| AT4G37690 | GT6 | Glycosyltransferase 6 | -1.99 | **4.93E-03** | -0.23 | 9.44E-01 | 1.74 | **7.38E-03** |
| AT4G34390 | XLG2 | Extra-large GTP-binding protein 2 | -1.99 | **4.30E-05** | -0.32 | 1.26E-01 | 1.65 | **9.14E-04** |
| AT5G15870 | At5g15870 | glucan endo-1,3-beta-D-glucosidase | -1.99 | **4.99E-05** | -0.36 | 1.51E-01 | 1.61 | **1.49E-03** |
| AT1G10040 | At1g10040 | Alpha/beta-Hydrolases superfamily protein | -1.98 | **4.86E-02** | | | 2.64 | **1.04E-02** |
| AT3G51340 | F24M12.380 | Uncharacterized protein F24M12.380 | -1.97 | **1.28E-04** | -1.24 | **2.54E-02** | 0.72 | 2.24E-01 |
| AT4G37530 | At4g37530 | Peroxidase | -1.95 | **1.49E-02** | -0.03 | 9.82E-01 | 1.90 | **1.40E-02** |
| AT1G61370 | At1g61370 | S-locus lectin protein kinase family protein | -1.95 | **2.15E-02** | -1.24 | 8.62E-02 | 0.68 | 4.95E-01 |
| AT1G22400 | UGT85A1 | UDP-glycosyltransferase 85A1 | -1.94 | **4.45E-04** | -0.22 | 7.41E-01 | 1.71 | **9.31E-04** |
| AT2G46440 | CNGC11 | Cyclic nucleotide-gated channels | -1.92 | **8.77E-08** | -0.75 | **5.09E-07** | 1.15 | **2.20E-03** |
| AT3G47090 | F13I12.140 | Leucine-rich repeat protein kinase family protein | -1.91 | **2.09E-06** | -0.26 | 5.64E-01 | 1.64 | **1.56E-04** |
| AT3G59700 | LECRK55 | L-type lectin-domain containing receptor kinase V.5 | -1.91 | **9.69E-05** | -0.13 | 9.04E-01 | 1.75 | **2.74E-04** |
| AT3G11820 | SYP121 | Syntaxin-121 | -1.90 | **4.05E-05** | -0.50 | **2.23E-02** | 1.38 | **5.53E-03** |
| AT3G09440 | HSP70-3 | Heat shock 70 kDa protein 3 | -1.90 | **4.05E-03** | -0.57 | **2.18E-04** | 1.32 | 5.66E-02 |
| AT2G41835 | SAP11 | Zinc finger AN1 and C2H2 domain-containing stress-associated protein 11 | -1.90 | **2.65E-04** | -0.44 | 3.78E-01 | 1.44 | **1.06E-02** |
| AT2G35910 | At2g35910 | | -1.89 | **9.53E-03** | -0.36 | 8.32E-01 | 1.51 | **3.38E-02** |
| AT3G08870 | LECRK-VI1 | Concanavalin A-like lectin protein kinase family protein | -1.89 | **9.88E-07** | -0.94 | **4.46E-05** | 0.94 | **1.50E-02** |
| AT3G17420 | At3g17420 | Serine/threonine protein kinase | -1.89 | **1.21E-02** | -0.09 | 9.60E-01 | 1.78 | **2.03E-02** |
| AT5G46500 | At5g46500 | Protein VARIATION IN COMPOUND TRIGGERED ROOT growth protein | -1.88 | **2.11E-05** | -0.22 | 7.57E-01 | 1.65 | **2.15E-04** |
| AT5G66910 | At5g66910 | Probable disease resistance protein At5g66910 | -1.88 | **2.86E-06** | -0.64 | **3.67E-02** | 1.22 | **3.41E-03** |
| AT3G60450 | At3g60450 | | -1.87 | **2.93E-03** | 0.27 | 8.40E-01 | 2.13 | **2.10E-04** |
| AT4G36090 | ALKBH9C | Oxidoreductase, 2OG-Fe | -1.86 | **3.19E-05** | -0.42 | 4.83E-01 | 1.43 | **4.12E-03** |
| AT2G29990 | NDA2 | Internal alternative NAD | -1.86 | **6.17E-04** | 0.02 | 1.00E+00 | 1.87 | **6.55E-04** |
| AT1G72950 | At1g72950 | TIR domain-containing protein | -1.86 | **1.43E-03** | | | 1.78 | **1.87E-02** |
| AT3G51430 | YLS2 | Calcium-dependent phosphotriesterase superfamily protein | -1.85 | **1.37E-03** | -0.28 | 4.87E-01 | 1.56 | **4.35E-03** |
| AT1G31580 | ECS1 | Protein ECS1 | -1.84 | **5.73E-05** | -0.54 | **5.27E-03** | 1.29 | **3.03E-03** |
| AT1G18210 | CML27 | Probable calcium-binding protein CML27 | -1.84 | **2.26E-04** | -0.14 | 8.60E-01 | 1.68 | **1.16E-03** |
| AT3G20600 | NDR1 | Protein NDR1 | -1.82 | **1.93E-05** | -0.67 | **4.83E-03** | 1.14 | **9.14E-03** |
| AT1G61260 | F11P17.2 | F11P17.2 protein | -1.81 | **2.91E-03** | 1.15 | 2.67E-01 | 2.96 | **2.77E-03** |
| AT2G28400 | S40-4 | Protein S40-4 | -1.80 | **1.15E-02** | | | 2.08 | **1.01E-02** |
| AT5G24210 | MOP9.2 | Alpha/beta-Hydrolases superfamily protein | -1.80 | **6.81E-05** | -0.26 | 4.82E-01 | 1.53 | **1.16E-04** |
| AT5G40720 | MNF13.28 | Glycosyltransferase family 92 protein | -1.79 | **1.62E-02** | -0.17 | 8.46E-01 | 1.60 | **1.54E-02** |
| AT1G77500 | At1g77500 | Uncharacterized protein | -1.79 | **6.31E-04** | -0.33 | 2.90E-01 | 1.44 | **7.46E-03** |
| AT2G46620 | At2g46620 | AAA-ATPase At2g46620 | -1.79 | **2.15E-03** | -0.01 | 1.00E+00 | 1.76 | **2.59E-03** |
| AT3G49780 | PSK3 | Phytosulfokines 3 | -1.78 | **1.47E-02** | -0.16 | 8.03E-01 | 1.61 | **1.85E-02** |
| AT3G13910 | At3g13910 | | -1.78 | **6.31E-03** | 0.17 | 9.56E-01 | 1.92 | **2.53E-02** |
| AT5G12930 | At5g12930 | Inactive rhomboid protein | -1.76 | **1.53E-02** | -0.55 | 5.61E-01 | 1.19 | 1.84E-01 |
| AT1G17600 | SOC3 | Disease resistance protein | -1.75 | **2.35E-06** | -0.86 | **2.14E-02** | 0.88 | 5.86E-02 |
| AT5G62180 | CXE20 | Probable carboxylesterase 120 | -1.75 | **4.77E-03** | -0.35 | 8.91E-01 | 1.38 | **3.80E-02** |
| AT3G47540 | At3g47540 | Putative endochitinase | -1.75 | **5.91E-03** | -0.39 | 6.00E-01 | 1.34 | **1.33E-02** |
| AT3G56030 | At3g56030 | Pentatricopeptide repeat-containing protein At3g56030, mitochondrial | -1.74 | **1.11E-02** | -0.79 | 1.80E-01 | 0.93 | 2.17E-01 |
| AT5G46230 | At5g46230 | | -1.74 | **1.57E-02** | -0.21 | 8.55E-01 | 1.53 | **3.15E-02** |
| AT3G44720 | ADT4 | Arogenate dehydratase 4, chloroplastic | -1.73 | **2.15E-04** | 0.04 | 9.65E-01 | 1.76 | **1.27E-04** |
| AT5G44390 | At5g44390 | Berberine bridge enzyme-like 25 | -1.73 | **9.99E-03** | -0.03 | 1.00E+00 | 1.69 | **3.03E-02** |
| AT1G49050 | APCB1 | Eukaryotic aspartyl protease family protein | -1.71 | **7.73E-05** | -0.39 | 9.14E-02 | 1.32 | **2.21E-03** |
| AT2G25110 | SDF2 | Stromal cell-derived factor 2-like protein | -1.70 | **7.81E-03** | -0.21 | 4.65E-01 | 1.48 | **2.11E-02** |
| AT5G56870 | BGAL4 | Beta-galactosidase 4 | -1.70 | **2.15E-04** | 0.00 | 1.00E+00 | 1.68 | **1.72E-04** |
| AT4G23190 | At4g23190 | Uncharacterized protein | -1.69 | **5.52E-04** | -0.78 | **1.22E-02** | 0.91 | 5.64E-02 |
| AT3G26220 | At3g26220 | At3g26230 | -1.69 | **4.19E-05** | 0.03 | 9.89E-01 | 1.71 | **9.56E-06** |
| AT5G44820 | At5g44820 | Nucleotide-diphospho-sugar transferase domain-containing protein | -1.69 | **3.35E-06** | -0.74 | **7.34E-03** | 0.94 | **2.70E-02** |
| AT3G13437 | EWR1 | Transmembrane protein | -1.69 | **1.50E-04** | -0.85 | 9.89E-02 | 0.83 | 5.86E-02 |
| AT3G49120 | PER34 | Peroxidase 34 | -1.69 | **8.21E-03** | -0.03 | 9.78E-01 | 1.65 | **4.96E-03** |
| AT1G02220 | NAC003 | NAC domain-containing protein 3 | -1.69 | **8.55E-03** | -0.08 | 9.72E-01 | 1.58 | **9.71E-03** |
| AT4G24970 | MORC7 | Protein MICRORCHIDIA 7 | -1.68 | **1.63E-04** | -1.10 | **9.93E-03** | 0.58 | 2.00E-01 |
| AT5G50460 | At5g50460 | Protein transport protein Sec61 subunit gamma | -1.68 | **3.48E-03** | -0.36 | 2.24E-01 | 1.30 | **2.74E-02** |
| AT4G02420 | LECRK44 | L-type lectin-domain containing receptor kinase IV.4 | -1.68 | **1.09E-04** | -1.07 | **2.62E-14** | 0.59 | 2.65E-01 |
| AT3G14470 | RPPL1 | Putative disease resistance RPP13-like protein 1 | -1.67 | **5.69E-04** | -0.66 | **2.28E-02** | 1.01 | **3.84E-02** |
| AT3G11840 | At3g11840 | U-box domain-containing protein | -1.66 | **6.90E-05** | -0.35 | 4.27E-01 | 1.29 | **2.15E-03** |
| AT1G14370 | PBL2 | Probable serine/threonine-protein kinase PBL2 | -1.64 | **4.35E-03** | 0.13 | 9.31E-01 | 1.76 | **7.84E-05** |
| AT4G25900 | At4g25900 | glucose-6-phosphate 1-epimerase | -1.64 | **1.44E-02** | 0.03 | 9.73E-01 | 1.66 | **1.07E-02** |
| AT3G50950 | RPP13L4 | Disease resistance RPP13-like protein 4 | -1.64 | **6.83E-05** | -0.24 | 4.80E-01 | 1.39 | **1.70E-04** |
| AT5G45510 | At5g45510 | Probable disease resistance protein At5g45510 | -1.64 | **1.68E-04** | -0.53 | **1.51E-04** | 1.10 | **1.56E-02** |
| AT3G15350 | At3g15350 | | -1.63 | **2.81E-02** | -0.35 | 2.14E-01 | 1.27 | 9.28E-02 |
| AT4G23130 | At4g23130 | Uncharacterized protein | -1.63 | **2.83E-06** | -0.60 | **4.49E-02** | 1.03 | **1.87E-04** |
| AT5G47070 | PBL19 | Probable serine/threonine-protein kinase PBL19 | -1.63 | **2.03E-05** | -0.29 | 4.30E-01 | 1.32 | **8.05E-04** |
| AT4G22980 | At4g22980 | Molybdenum cofactor sulfurase-like protein | -1.63 | **2.37E-03** | -0.39 | 5.61E-01 | 1.22 | **4.18E-02** |
| AT4G23200 | CRK12 | Cysteine-rich RLK | -1.62 | **4.57E-03** | -0.43 | 2.96E-01 | 1.18 | **4.02E-02** |

| | | | | | | | | |
|---|---|---|---|---|---|---|---|---|
| AT1G60730 | At1g60730 | NAD | -1.62 | **5.53E-03** | 0.02 | 1.00E+00 | 1.62 | **2.50E-02** |
| AT5G54710 | At5g54710 | Similarity to ankyrin-like protein | -1.62 | **7.25E-04** | -0.07 | 8.82E-01 | 1.53 | **9.65E-04** |
| AT4G23470 | At4g23470 | PLAC8 family protein | -1.62 | **2.63E-04** | -0.35 | **2.62E-02** | 1.25 | **6.70E-03** |
| AT1G01010 | NAC001 | NAC domain-containing protein 1 | -1.61 | **1.07E-02** | 0.17 | 9.38E-01 | 1.77 | **1.27E-02** |
| AT1G64610 | At1g64610 | Transducin/WD40 repeat-like superfamily protein | -1.60 | **2.73E-04** | 0.06 | 9.68E-01 | 1.64 | **2.82E-04** |
| AT4G11350 | At4g11350 | Transferring glycosyl group transferase | -1.60 | **2.19E-02** | 0.28 | 9.14E-01 | 1.85 | **6.66E-03** |
| AT3G19970 | At3g19970 | | -1.59 | **2.74E-02** | -0.03 | 1.00E+00 | 1.56 | **2.99E-02** |
| AT2G25000 | WRKY60 | WRKY DNA-binding protein 60 | -1.57 | **3.24E-02** | 1.48 | **1.78E-02** | 3.05 | **1.10E-06** |
| AT3G09830 | PCRK1 | Serine/threonine-protein kinase PCRK1 | -1.57 | **4.32E-04** | -0.42 | 1.99E-01 | 1.15 | **8.05E-03** |
| AT5G04720 | At5g04720 | Probable disease resistance protein At5g04720 | -1.57 | **5.01E-04** | -0.27 | 3.42E-01 | 1.29 | **4.13E-03** |
| AT3G50470 | HR3 | RPW8-like protein 3 | -1.57 | **4.09E-04** | -0.12 | 9.55E-01 | 1.44 | **1.01E-03** |
| AT4G20830 | At4g20830 | Berberine bridge enzyme-like 19 | -1.56 | **1.88E-02** | -0.24 | 9.38E-01 | 1.28 | 6.92E-02 |
| AT1G27770 | ACA1 | Calcium-transporting ATPase | -1.55 | **1.69E-04** | -0.27 | 4.62E-01 | 1.27 | **2.82E-04** |
| AT4G29810 | At4g29810 | AT4G29810 protein | -1.55 | **9.05E-04** | -0.36 | 1.64E-01 | 1.18 | **1.53E-02** |
| AT2G23810 | TET8 | Tetraspanin-8 | -1.51 | **8.92E-03** | 0.08 | 9.20E-01 | 1.58 | **1.01E-03** |
| AT1G09970 | At1g09970 | Protein kinase domain-containing protein | -1.51 | **6.45E-03** | -0.06 | 9.38E-01 | 1.44 | **4.80E-03** |
| AT1G19180 | JAZ1 | Protein TIFY | -1.51 | **2.71E-02** | 0.93 | 8.44E-02 | 2.43 | **1.34E-08** |
| AT2G16870 | At2g16870 | Disease resistance protein | -1.50 | **9.75E-05** | -0.57 | 1.73E-01 | 0.91 | **2.50E-02** |
| AT5G08380 | AGAL1 | Alpha-galactosidase | -1.49 | **1.62E-02** | 0.07 | 9.50E-01 | 1.55 | **1.25E-02** |
| AT1G18570 | MYB51 | Transcription factor MYB51 | -1.48 | **3.48E-03** | 0.18 | 6.23E-01 | 1.64 | **8.17E-04** |
| AT3G26230 | At3g26230 | Uncharacterized protein | -1.47 | **1.20E-03** | -0.43 | 6.66E-02 | 1.03 | **2.39E-02** |
| AT1G13990 | At1g13990 | Plant/protein | -1.47 | **2.26E-02** | 0.09 | 8.94E-01 | 1.55 | **6.01E-03** |
| AT5G09590 | HSP70-10 | Heat shock 70 kDa protein 10, mitochondrial | -1.47 | **1.99E-02** | -0.20 | 7.25E-01 | 1.25 | 5.35E-02 |
| AT3G15760 | At3g15760 | Transmembrane protein | -1.47 | **1.26E-03** | -0.62 | 3.28E-01 | 0.83 | 9.16E-02 |
| AT3G55470 | At3g55470 | Calcium-dependent lipid-binding | -1.47 | **3.35E-03** | 0.16 | 9.22E-01 | 1.61 | **7.50E-03** |
| AT5G45500 | MFC19.17 | RNI-like superfamily protein | -1.45 | **4.00E-04** | -0.26 | 2.95E-01 | 1.18 | **5.58E-03** |
| AT5G23850 | MRO11.11 | At5g23850 | -1.45 | **1.49E-02** | -0.09 | 8.94E-01 | 1.34 | **2.85E-02** |
| AT1G67850 | At1g67850 | | -1.45 | **9.38E-04** | 0.12 | 9.12E-01 | 1.56 | **1.34E-04** |
| AT1G80610 | At1g80610 | At1g80610/T21F11_6 | -1.44 | **3.08E-04** | -0.11 | 8.73E-01 | 1.31 | **8.08E-04** |
| AT2G43850 | ILK1 | Integrin-linked protein kinase family | -1.43 | **1.11E-05** | -0.06 | 9.62E-01 | 1.36 | **6.45E-05** |
| AT2G01650 | PUX2 | Plant UBX domain-containing protein 2 | -1.42 | **2.93E-03** | -0.24 | 5.70E-01 | 1.17 | **1.25E-02** |
| AT5G66320 | GATA5 | GATA transcription factor 5 | -1.42 | **3.06E-03** | 0.12 | 8.90E-01 | 1.52 | **5.55E-04** |
| AT4G00955 | At4g00955 | Uncharacterized protein At4g00955 | -1.42 | **6.05E-06** | -0.60 | 5.36E-02 | 0.81 | **8.45E-03** |
| AT2G30020 | At2g30020 | Probable protein phosphatase 2C 25 | -1.41 | **9.89E-07** | 0.59 | **2.15E-02** | 1.99 | **2.35E-13** |
| AT3G03440 | At3g03440 | ARM repeat superfamily protein | -1.41 | **3.13E-03** | -0.33 | 4.26E-01 | 1.07 | **3.38E-02** |
| AT5G45110 | NPR3 | NPR1-like protein 3 | -1.41 | **4.99E-04** | 0.04 | 9.65E-01 | 1.44 | **4.00E-05** |
| AT4G23570 | SGT1A | Protein SGT1 homolog A | -1.41 | **7.25E-04** | 0.01 | 9.99E-01 | 1.41 | **2.52E-04** |
| AT3G05830 | ATNEAP1 | Spindle pole body component-like protein | -1.41 | **2.48E-02** | -0.52 | 5.35E-01 | 0.88 | 9.97E-02 |
| AT1G69840 | HIR2 | Hypersensitive-induced response protein 2 | -1.40 | **9.40E-03** | 0.16 | 8.03E-01 | 1.55 | **1.09E-03** |
| AT3G11650 | NHL2 | NDR1/HIN1-like protein 2 | -1.40 | **1.14E-02** | 0.04 | 9.87E-01 | 1.43 | **8.78E-03** |
| AT2G31990 | At2g31990 | Exostosin family protein | -1.39 | **1.83E-02** | -0.03 | 1.00E+00 | 1.35 | **1.90E-02** |
| AT3G23560 | ALF5 | MATE efflux family protein | -1.37 | **9.73E-03** | -0.47 | 5.39E-01 | 0.87 | 1.51E-01 |
| AT1G11310 | At1g11310 | At1g11310/T28P6_23 | -1.37 | **1.18E-03** | -0.39 | **6.57E-03** | 0.96 | **2.71E-02** |
| AT5G53120 | SPDS3 | Spermidine synthase 3 | -1.37 | **4.09E-02** | 0.02 | 1.00E+00 | 1.36 | **3.26E-02** |
| AT1G09070 | SRC2 | Protein SRC2 homolog | -1.36 | **3.64E-03** | 0.19 | 6.47E-01 | 1.53 | **3.41E-04** |
| AT5G14930 | SAG101 | Senescence-associated gene 101 | -1.35 | **2.70E-03** | -0.12 | 7.72E-01 | 1.21 | **7.40E-03** |
| AT1G33560 | ADR1 | Disease resistance protein ADR1 | -1.35 | **9.74E-03** | -0.38 | 2.57E-01 | 0.96 | 9.06E-02 |
| AT1G62422 | At1g62422 | Uncharacterized protein | -1.33 | **3.60E-02** | -0.38 | 6.24E-01 | 0.94 | 1.67E-01 |
| AT4G01740 | At4g01740 | Cysteine/Histidine-rich C1 domain family protein | -1.33 | **9.16E-03** | -0.74 | 1.83E-01 | 0.58 | 3.33E-01 |
| AT5G64550 | At5g64550 | Uncharacterized protein At5g64550 | -1.33 | **4.52E-03** | -0.18 | 8.52E-01 | 1.13 | **1.03E-02** |
| AT4G33300 | At4g33300 | Probable disease resistance protein At4g33300 | -1.32 | **3.78E-04** | -0.13 | 6.92E-01 | 1.18 | **1.47E-03** |
| AT1G62305 | At1g62305 | At1g62305 | -1.32 | **1.52E-02** | -0.79 | 9.46E-02 | 0.52 | 4.21E-01 |
| AT2G28940 | PBL37 | At2g28940 | -1.32 | **1.66E-03** | 0.24 | 8.42E-01 | 1.55 | **1.91E-03** |
| AT5G39785 | At5g39780 | Gb\|AAF22924.1 | -1.31 | **5.69E-03** | -0.03 | 1.00E+00 | 1.26 | **1.21E-02** |
| AT3G16510 | At3g16510 | | -1.30 | **5.31E-03** | -0.38 | 3.94E-01 | 0.91 | **2.34E-02** |
| AT3G16720 | ATL2 | RING-H2 finger protein ATL2 | -1.29 | **1.99E-03** | 0.00 | 1.00E+00 | 1.28 | **2.27E-04** |
| AT5G18310 | At5g18310 | Ubiquitin hydrolase | -1.29 | **2.49E-02** | -0.04 | 9.83E-01 | 1.24 | **3.35E-02** |
| AT5G44070 | CAD1 | glutathione gamma-glutamylcysteinyltransferase | -1.29 | **1.51E-03** | -0.53 | **3.30E-03** | 0.75 | 7.53E-02 |
| AT5G11000 | At5g11000 | DUF868 family protein | -1.29 | **4.82E-02** | -0.18 | 5.56E-01 | 1.10 | 9.05E-02 |
| AT5G58120 | At5g58120 | ADP-ribosyl cyclase/cyclic ADP-ribose hydrolase | -1.29 | **1.13E-03** | -0.10 | 9.04E-01 | 1.18 | **5.17E-04** |
| AT5G11650 | MAGL13 | Alpha/beta-Hydrolases superfamily protein | -1.29 | **2.73E-03** | -0.46 | 4.37E-01 | 0.81 | 8.17E-02 |
| AT3G21220 | MKK5 | Mitogen-activated protein kinase kinase 5 | -1.29 | **3.90E-03** | -0.14 | 7.56E-01 | 1.14 | **9.16E-03** |
| AT4G08230 | At4g08230 | Glycine-rich protein | -1.28 | **3.00E-02** | -0.17 | 6.84E-01 | 1.10 | 6.45E-02 |
| AT3G59880 | F24G16.150 | Uncharacterized protein F24G16.150 | -1.28 | **1.42E-02** | -0.83 | **4.69E-02** | 0.43 | 4.94E-01 |
| AT2G23200 | At2g23200 | Probable receptor-like protein kinase At2g23200 | -1.27 | **2.15E-04** | -0.60 | **5.42E-04** | 0.66 | 7.05E-02 |
| AT3G53810 | LECRK42 | L-type lectin-domain containing receptor kinase IV.2 | -1.27 | **9.64E-03** | -0.28 | 4.43E-01 | 0.97 | **4.46E-02** |
| AT3G10010 | DML2 | DEMETER-like protein 2 | -1.27 | **1.83E-04** | -0.47 | 5.13E-02 | 0.79 | **2.22E-02** |
| AT1G23710 | At1g23710 | At1g23710 | -1.26 | **9.31E-03** | -0.28 | 8.11E-01 | 0.97 | 1.04E-01 |
| AT2G40270 | At2g40270 | Protein kinase family protein | -1.26 | **1.53E-03** | -0.29 | 1.09E-01 | 0.95 | **2.02E-02** |
| AT4G14746 | At4g14746 | Neurogenic locus notch-like protein | -1.26 | **1.70E-03** | -0.09 | 9.04E-01 | 1.15 | **2.60E-03** |
| AT5G61010 | EXO70E2 | Exocyst complex component EXO70E2 | -1.26 | **2.42E-04** | -0.45 | **2.23E-02** | 0.79 | **1.65E-02** |
| AT1G31540 | T8E3.20 | ADP-ribosyl cyclase/cyclic ADP-ribose hydrolase | -1.23 | **2.59E-03** | -0.30 | 3.27E-01 | 0.91 | **8.74E-03** |
| AT1G59870 | ABCG36 | ABC transporter G family member 36 | -1.23 | **8.33E-03** | 0.09 | 7.81E-01 | 1.30 | **2.79E-03** |
| AT5G19750 | At5g19750 | Peroxisomal membrane 22 kDa | -1.22 | **1.15E-02** | -1.15 | **1.95E-03** | 0.06 | 9.46E-01 |
| AT3G16860 | COBL8 | COBRA-like protein 8 | -1.22 | **2.79E-03** | 0.40 | 5.17E-01 | 1.61 | **1.47E-06** |
| AT3G20250 | PUM5 | Pumilio 5 | -1.20 | **2.69E-03** | 0.03 | 9.75E-01 | 1.22 | **1.16E-03** |
| AT3G13330 | PA200 | Proteasome activator subunit 4 | -1.20 | **1.86E-03** | -0.19 | 3.93E-01 | 1.00 | **8.86E-03** |
| AT4G16960 | SIKIC3 | Disease resistance protein | -1.20 | **2.33E-03** | -0.08 | 9.67E-01 | 1.11 | **9.16E-03** |
| AT2G17760 | APF1 | Aspartyl protease family protein 1 | -1.20 | **1.76E-02** | -0.18 | 6.42E-01 | 1.01 | **4.17E-02** |
| AT5G49520 | At5g49520 | WRKY domain-containing protein | -1.19 | **1.14E-02** | 0.18 | 7.95E-01 | 1.36 | **1.51E-03** |
| AT1G32127 | F3C3.9 | Uncharacterized protein F3C3.9 | -1.19 | **1.90E-02** | -0.67 | 2.47E-01 | 0.50 | 4.38E-01 |
| AT5G02290 | PBL11 | Probable serine/threonine-protein kinase PBL11 | -1.18 | **2.17E-02** | -0.26 | 5.80E-01 | 0.91 | **2.92E-02** |
| AT5G56030 | HSP81-2 | Heat shock protein 81-2 | -1.17 | **1.85E-02** | -0.57 | **1.39E-04** | 0.60 | 2.80E-01 |
| AT5G22630 | ADT5 | Arogenate dehydratase 5, chloroplastic | -1.17 | **2.95E-02** | -0.12 | 8.48E-01 | 1.03 | **4.68E-02** |
| AT3G21630 | At3g21630 | Protein kinase domain-containing protein | -1.15 | **9.58E-04** | -0.14 | 7.03E-01 | 1.00 | **4.46E-03** |
| AT3G16030 | At3g16030 | Receptor-like serine/threonine-protein kinase | -1.15 | **1.67E-02** | -0.02 | 1.00E+00 | 1.12 | **4.59E-03** |
| AT4G21390 | At4g21390 | Receptor-like serine/threonine-protein kinase | -1.14 | **1.88E-02** | 0.26 | 8.24E-01 | 1.39 | **1.39E-03** |
| AT2G32160 | At2g32160 | S-adenosyl-L-methionine-dependent methyltransferases superfamily protein | -1.14 | **9.97E-05** | -0.49 | **2.79E-03** | 0.63 | **2.64E-02** |
| AT3G57330 | ACA11 | Calcium-transporting ATPase | -1.12 | **5.94E-04** | -0.47 | **8.81E-04** | 0.64 | 7.75E-02 |
| AT4G23180 | At4g23180 | Uncharacterized protein | -1.12 | **1.04E-02** | -0.56 | **1.55E-05** | 0.54 | 2.64E-01 |

| | | | | | | | | |
|---|---|---|---|---|---|---|---|---|
| AT2G05940 | RIPK | Serine/threonine-protein kinase RIPK | -1.12 | **2.64E-03** | -0.12 | 7.65E-01 | 0.99 | **7.90E-03** |
| AT2G27080 | NHL13 | NDR1/HIN1-like protein 13 | -1.11 | **4.53E-03** | 0.96 | **1.20E-02** | 2.06 | **7.69E-13** |
| AT5G18490 | At5g18490 | At5g18490 | -1.11 | **1.52E-02** | 0.07 | 9.59E-01 | 1.16 | **3.55E-03** |
| AT3G55450 | PBL1 | PBS1-like 1 | -1.11 | **3.11E-02** | 0.34 | 3.81E-01 | 1.43 | **2.60E-03** |
| AT3G47010 | F13I12.60 | beta-glucosidase | -1.10 | **7.42E-03** | -0.25 | 7.42E-01 | 0.84 | 5.80E-02 |
| AT1G67470 | ZRK12 | Inactive serine/threonine-protein kinase ZRK12 | -1.10 | **8.92E-03** | -0.40 | 4.37E-01 | 0.68 | 7.02E-02 |
| AT2G11520 | CRCK3 | Calmodulin-binding receptor-like cytoplasmic kinase 3 | -1.10 | **1.50E-02** | -0.25 | 3.64E-01 | 0.83 | 7.92E-02 |
| AT2G07180 | PBL17 | Probable serine/threonine-protein kinase PBL17 | -1.10 | **1.42E-03** | -0.01 | 1.00E+00 | 1.08 | **1.46E-03** |
| AT4G01720 | WRKY47 | Probable WRKY transcription factor 47 | -1.10 | **3.92E-02** | 0.33 | 9.04E-02 | 1.41 | **3.45E-03** |
| AT5G66850 | MAPKKK5 | Mitogen-activated protein kinase kinase kinase 5 | -1.09 | **4.80E-02** | -0.24 | 3.24E-01 | 0.83 | 1.33E-01 |
| AT5G19250 | At5g19250 | Uncharacterized GPI-anchored protein At5g19250 | -1.09 | **1.53E-02** | -0.46 | **2.66E-04** | 0.61 | 2.08E-01 |
| AT1G54320 | ALIS3 | ALA-interacting subunit 3 | -1.08 | **1.90E-02** | -0.05 | 9.46E-01 | 1.02 | **2.34E-02** |
| AT2G39660 | BIK1 | Serine/threonine-protein kinase BIK1 | -1.08 | **1.79E-02** | -0.02 | 1.00E+00 | 1.06 | **1.67E-02** |
| AT3G59660 | BAGP1 | C2 domain-containing protein / GRAM domain-containing protein | -1.08 | **1.38E-02** | 0.34 | 1.69E-01 | 1.40 | **4.56E-04** |
| AT3G47570 | At3g47570 | Probable LRR receptor-like serine/threonine-protein kinase At3g47570 | -1.07 | **2.26E-03** | -0.27 | 4.81E-01 | 0.79 | **3.16E-02** |
| AT5G05190 | EDR4 | Protein ENHANCED DISEASE RESISTANCE 4 | -1.07 | **2.48E-02** | -0.16 | 7.77E-01 | 0.90 | **4.07E-02** |
| AT4G01920 | At4g01920 | AT4g01920/T7B11_18 | -1.07 | **3.80E-02** | -0.16 | 8.72E-01 | 0.88 | **2.04E-02** |
| AT4G19660 | NPR4 | NPR1-like protein 4 | -1.07 | **1.35E-03** | -0.32 | 8.72E-02 | 0.73 | **3.91E-02** |
| AT4G23250 | EMB1290 | Cysteine-rich receptor-like protein kinase 17 | -1.06 | **5.91E-03** | -0.56 | **1.73E-03** | 0.49 | 2.62E-01 |
| AT2G40000 | HSPRO2 | Nematode resistance protein-like HSPRO2 | -1.06 | **3.70E-02** | 0.48 | 3.25E-01 | 1.53 | **6.78E-09** |
| AT1G64280 | NPR1 | Regulatory protein NPR1 | -1.06 | **5.56E-04** | -0.42 | **1.15E-02** | 0.62 | 6.03E-02 |
| AT5G47730 | MCA23.5 | AT5G47730 protein | -1.05 | **4.56E-03** | -0.32 | 5.31E-01 | 0.71 | 7.44E-02 |
| AT1G03400 | At1g03400 | 2-oxoglutarate | -1.04 | **1.33E-04** | -0.40 | **4.23E-02** | 0.62 | **2.48E-02** |
| AT2G02800 | PBL3 | Probable serine/threonine-protein kinase PBL3 | -1.04 | **5.65E-03** | -0.36 | 1.40E-01 | 0.66 | 9.12E-02 |
| AT2G15042 | RLP18 | Receptor-like protein 18 | -1.03 | **1.04E-02** | -0.25 | 3.66E-01 | 0.77 | **4.27E-02** |
| AT3G07780 | OBE1 | Protein OBERON | -1.01 | **1.96E-02** | -0.14 | 7.29E-01 | 0.86 | **4.11E-02** |
| AT3G07570 | At3g07570 | Cytochrome b561 and DOMON domain-containing protein At3g07570 | -1.01 | **1.12E-02** | -0.13 | 8.23E-01 | 0.87 | **1.74E-02** |
| AT4G03820 | At4g03820 | Transmembrane protein, putative | -1.01 | **4.01E-02** | -0.14 | 8.22E-01 | 0.85 | 6.75E-02 |
| AT4G02410 | LECRK43 | L-type lectin-domain containing receptor kinase IV.3 | -1.01 | **1.65E-03** | -0.33 | 9.46E-02 | 0.67 | 5.11E-02 |
| AT2G23450 | WAKL14 | Wall-associated receptor kinase-like 14 | -1.01 | **2.70E-02** | -0.17 | 7.21E-01 | 0.82 | **4.06E-02** |
| AT3G42790 | AL3 | PHD finger protein ALFIN-LIKE 3 | -1.00 | **3.14E-02** | -0.33 | 3.00E-01 | 0.65 | 1.63E-01 |
| AT4G29050 | LECRK-V9 | non-specific serine/threonine protein kinase | -1.00 | **2.72E-02** | -0.23 | 5.22E-01 | 0.75 | 1.11E-01 |
| AT4G39820 | TRAPPC12 | At4g39820 | -0.99 | **2.09E-02** | -0.12 | 8.77E-01 | 0.86 | 6.45E-02 |
| AT4G13040 | APD1 | Integrase-type DNA-binding superfamily protein | -0.98 | **4.20E-03** | -0.52 | **4.84E-02** | 0.45 | 2.04E-01 |
| AT3G47820 | PUB39 | U-box domain-containing protein 39 | -0.98 | **1.63E-02** | 0.01 | 1.00E+00 | 0.97 | **1.26E-02** |
| AT4G01370 | MPK4 | Mitogen-activated protein kinase 4 | -0.97 | **2.67E-02** | -0.47 | **4.73E-03** | 0.49 | 3.16E-01 |
| AT4G34480 | At4g34480 | glucan endo-1,3-beta-D-glucosidase | -0.97 | **2.71E-02** | -0.21 | 4.06E-01 | 0.75 | 8.09E-02 |
| AT3G54850 | At3g54850 | AT3G54850 protein | -0.97 | **4.99E-03** | -0.07 | 9.51E-01 | 0.88 | **9.22E-03** |
| AT1G17430 | F28G4.9 | F28G4.9 protein | -0.97 | **1.43E-02** | -0.82 | 7.40E-02 | 0.14 | 8.28E-01 |
| AT3G60190 | At3g60190 | UMP-CMP kinase | -0.97 | **4.01E-02** | -0.02 | 9.84E-01 | 0.94 | **3.86E-02** |
| AT4G08480 | MEKK2 | Mitogen-activated protein kinase kinase kinase 9 | -0.95 | **6.03E-03** | -0.42 | 8.28E-02 | 0.52 | 1.83E-01 |
| AT4G31500 | CYP83B1 | Cytochrome P450 83B1 | -0.94 | **4.68E-02** | 0.65 | **8.27E-06** | 1.58 | **3.31E-05** |
| AT5G10610 | CYP81K1 | Cytochrome P450, family 81, subfamily K, polypeptide 1 | -0.94 | **3.16E-02** | -0.20 | 5.45E-01 | 0.72 | 9.94E-02 |
| AT1G22890 | PROSCOOP1 | Serine rich endogenous peptide 14 | -0.94 | **2.24E-03** | -0.02 | 9.94E-01 | 0.91 | **2.67E-03** |
| AT5G66210 | At5g66210 | AT5G66210 protein | -0.94 | **2.51E-02** | 0.19 | 5.89E-01 | 1.11 | **1.57E-03** |
| AT2G37940 | IPCS2 | Phosphatidylinositol:ceramide inositolphosphotransferase 2 | -0.93 | **4.46E-03** | -0.31 | 1.23E-01 | 0.61 | 8.09E-02 |
| AT4G19990 | FRS1 | Protein FAR1-RELATED SEQUENCE | -0.93 | **8.88E-03** | -0.26 | 5.80E-01 | 0.66 | 7.52E-02 |
| AT1G66920 | At1g66920 | Protein kinase superfamily protein | -0.93 | **4.51E-03** | 0.87 | **1.18E-04** | 1.79 | **5.29E-09** |
| AT5G28830 | At5g28830 | Calcium-binding EF hand family protein | -0.93 | **2.61E-02** | 0.11 | 8.77E-01 | 1.03 | **1.25E-02** |
| AT1G22070 | At1g22070 | Uncharacterized protein | -0.93 | **2.77E-02** | 0.13 | 7.70E-01 | 1.04 | **9.39E-03** |
| AT1G25390 | LRK10L-1.1 | LEAF RUST 10 DISEASE-RESISTANCE LOCUS RECEPTOR-LIKE PROTEIN KINASE-like 1.1 | -0.92 | **1.04E-02** | -0.13 | 7.29E-01 | 0.77 | **2.99E-02** |
| AT3G29400 | EXO70E1 | Exocyst subunit Exo70 family protein | -0.91 | **1.81E-02** | -0.13 | 7.48E-01 | 0.77 | **4.49E-02** |
| AT2G31870 | TEJ | poly | -0.91 | **5.75E-03** | -0.27 | 5.93E-01 | 0.62 | 6.11E-02 |
| AT1G76700 | ATJ10 | Chaperone protein dnaJ 10 | -0.90 | **2.66E-02** | -0.14 | 8.26E-01 | 0.75 | 7.03E-02 |
| AT1G56510 | ADR2 | Disease resistance protein ADR2 | -0.89 | **1.51E-02** | -0.20 | 6.02E-01 | 0.68 | **1.63E-02** |
| AT2G21500 | At2g21500 | Uncharacterized protein At2g21500 | -0.89 | **3.95E-02** | 0.23 | 5.49E-01 | 1.10 | **2.04E-03** |
| AT1G55120 | FRUCT5 | Beta-fructofuranosidase 5 | -0.89 | **3.64E-02** | -0.05 | 9.65E-01 | 0.82 | 5.35E-02 |
| AT3G53270 | At3g53270 | Small nuclear RNA activating complex | -0.88 | **8.52E-03** | -0.29 | 4.76E-01 | 0.57 | 1.26E-01 |
| AT5G62570 | CBP60A | Calmodulin binding protein-like protein | -0.88 | **4.56E-03** | -0.12 | 8.12E-01 | 0.74 | **9.89E-03** |
| AT3G49530 | NAC062 | NAC domain containing protein 62 | -0.87 | **3.00E-02** | 0.21 | 6.06E-01 | 1.06 | **2.51E-03** |
| AT1G14790 | RDR1 | RNA-dependent RNA polymerase 1 | -0.86 | **4.70E-03** | -0.07 | 8.71E-01 | 0.78 | **8.41E-03** |
| AT5G46490 | At5g46490 | Disease resistance protein | -0.86 | **1.85E-02** | -0.08 | 8.72E-01 | 0.76 | **1.84E-02** |
| AT3G23170 | PRP | At3g23170 | -0.85 | **2.23E-02** | 0.39 | 5.21E-01 | 1.23 | **4.07E-04** |
| AT1G52290 | PERK15 | non-specific serine/threonine protein kinase | -0.85 | **1.66E-02** | -0.45 | **6.17E-03** | 0.38 | 3.16E-01 |
| AT5G46260 | MPL12.4 | ADP-ribosyl cyclase/cyclic ADP-ribose hydrolase | -0.85 | **2.65E-02** | -0.05 | 9.65E-01 | 0.78 | **1.65E-02** |
| AT4G27740 | At4g27740 | Protein yippee-like At4g27740 | -0.83 | **3.30E-02** | 0.24 | 7.25E-01 | 1.06 | **1.10E-02** |
| AT1G28380 | NSL1 | MACPF domain-containing protein NSL1 | -0.82 | **1.53E-02** | -0.14 | 6.86E-01 | 0.67 | 5.69E-02 |
| AT1G56520 | F13N6.13 | Disease resistance protein | -0.82 | **2.76E-02** | -0.17 | 6.21E-01 | 0.64 | 7.77E-02 |
| AT5G66675 | At5g66675 | Uncharacterized protein | -0.82 | **4.58E-02** | -0.03 | 9.89E-01 | 0.77 | **1.61E-02** |
| AT4G34460 | At4g34460 | At4g34460 | -0.81 | **2.86E-02** | -0.02 | 1.00E+00 | 0.77 | 5.35E-02 |
| AT5G61530 | At5g61530 | Small G protein family protein / RhoGAP family protein | -0.81 | **1.90E-02** | 0.13 | 7.73E-01 | 0.93 | **7.46E-03** |
| AT5G45410 | ATNHR2A | AT5G45410 protein | -0.81 | **2.24E-02** | -0.26 | 5.64E-01 | 0.53 | 2.31E-01 |
| AT3G46510 | PUB13 | U-box domain-containing protein 13 | -0.80 | **4.18E-02** | 0.11 | 7.25E-01 | 0.90 | **9.74E-03** |
| AT1G17420 | At1g17420 | Lipoxygenase | -0.80 | **3.29E-02** | 0.00 | 1.00E+00 | 0.79 | **1.93E-02** |
| AT3G19190 | ATG2 | Autophagy-related protein 2 | -0.80 | **1.53E-02** | -0.02 | 9.76E-01 | 0.76 | **1.35E-02** |
| AT1G28280 | MVQ1 | VQ motif-containing protein | -0.79 | **3.75E-02** | -0.14 | 6.24E-01 | 0.63 | 8.34E-02 |
| AT2G01670 | NUDT17 | Nudix hydrolase 17, mitochondrial | -0.79 | **2.45E-02** | -0.12 | 7.62E-01 | 0.66 | 6.13E-02 |
| AT4G23260 | CRK18 | Cysteine-rich RLK | -0.79 | **1.39E-02** | 0.22 | 4.76E-01 | 0.99 | **5.39E-05** |
| AT1G69440 | AGO7 | Protein argonaute 7 | -0.78 | **4.58E-02** | -0.17 | 6.23E-01 | 0.60 | 1.31E-01 |
| AT5G56260 | At5g56260 | 4-hydroxy-4-methyl-2-oxoglutarate aldolase | -0.78 | **1.57E-02** | -0.23 | 4.56E-01 | 0.53 | 1.15E-01 |
| AT1G12990 | F3F19.2 | F3F19.2 protein | -0.78 | **1.23E-02** | -0.15 | 6.41E-01 | 0.61 | **4.12E-02** |
| AT1G75020 | LPAT4 | Probable 1-acyl-sn-glycerol-3-phosphate acyltransferase 4 | -0.76 | **3.64E-02** | -0.21 | 5.96E-01 | 0.53 | 1.66E-01 |
| AT5G44580 | PROSCOOP1 | Serine rich endogenous peptide 10 | -0.76 | **2.13E-02** | 0.07 | 8.72E-01 | 0.82 | **4.00E-03** |
| AT3G59080 | F17J16_130 | AT3g59080/F17J16_130 | -0.72 | **4.10E-02** | -0.17 | 7.12E-01 | 0.54 | 1.31E-01 |
| AT1G10920 | LOV1 | NB-ARC domain-containing disease resistance protein | -0.72 | **4.92E-02** | -0.37 | 2.46E-01 | 0.34 | 3.68E-01 |
| AT2G44500 | At2g44500 | O-fucosyltransferase family protein | -0.72 | **4.86E-02** | 0.47 | **4.28E-02** | 1.18 | **5.23E-07** |
| AT1G78920 | AVPL1 | Pyrophosphate-energized membrane proton pump 2 | -0.70 | **2.49E-02** | -0.15 | 6.11E-01 | 0.54 | 8.72E-02 |
| AT1G79380 | RGLG4 | E3 ubiquitin-protein ligase RGLG4 | -0.70 | **3.27E-02** | -0.33 | **4.09E-02** | 0.36 | 2.98E-01 |
| AT1G60490 | VPS34 | Phosphatidylinositol 3-kinase VPS34 | -0.65 | **3.78E-02** | -0.21 | 9.99E-01 | 0.44 | 3.88E-01 |
| AT4G12010 | DSC1 | Disease resistance-like protein DSC1 | -0.64 | **4.80E-02** | -0.43 | **3.61E-02** | 0.20 | 5.87E-01 |
